# The transcriptional landscape of *Arabidopsis thaliana* pattern-triggered immunity

**DOI:** 10.1101/2020.11.30.404566

**Authors:** Marta Bjornson, Priya Pimprikar, Thorsten Nürnberger, Cyril Zipfel

## Abstract

Plants initiate immunity upon recognition of a wide array of self and non-self molecular patterns. Whether plants tune their immune outputs to patterns of different biological origins or of different biochemical nature remains mostly unclear. Here, we performed a detailed early time-series transcriptomics analysis in *Arabidopsis thaliana*, revealing that the response to diverse patterns is remarkably congruent. Early transcriptional reprogramming is dominated by a plant general stress response (GSR), which is essential for pattern-induced immunity. The definition of ‘core immunity response’ genes common and specific to pattern response in addition revealed the function of previously uncharacterized GLUTAMATE RECEPTOR-LIKE calcium-permeable channels in immunity. This study thus illustrates general and unique properties of early immune transcriptional reprogramming and uncovered important components of plant immunity.

**One Sentence Summary:** Time-resolved transcriptomics reveals new properties of pattern-triggered immunity and function of calcium-permeable channels.

## Main Text

Plants are challenged by a wide variety of potentially pathogenic organisms; their health relies on their ability to recognize and respond to this plethora of challenges. This recognition is partly accomplished through cell surface-localized pattern recognition receptors (PRRs), which recognize pathogen-associated molecular patterns (PAMPs) or host-derived damage-associated molecular patterns (DAMPs), leading to pattern-triggered immunity (PTI) (*1*). While a wide variety of PRRs with an equivalent variety of cognate ligands have been identified in various plant species (*2*), it is still unclear to what extent plants discriminate among patterns from different source organism, of different chemical nature, or that are recognized by different PRR classes. Notably, while a few studies have compared transcriptional responses (as a proxy of a dynamic large immune cellular output) triggered by two or three patterns together (*3–5*), or used meta analyses to compare responses (*6, 7*), these studies were limited in scale or utilized different experimental conditions, which hinders meaningful comparisons.

To ascertain the timing and degree of discrimination among pattern-triggered transcriptional responses, we selected a panel of seven patterns with known PRRs, representing a variety of source organism, chemical composition, and recognition mechanisms. This included bacterial flg22 (a 22-amino acid epitope derived from bacterial flagellin) recognized by the leucine-rich repeat receptor kinase (LRR-RK) FLAGELLIN SENSING 2 (FLS2) (*8*), elf18 (an 18-amino acid epitope derived from bacterial elongation factor Tu) recognized by the LRR-RK EF-TU RECEPTOR (EFR) (*7*), Pep1 (a 23-amino acid peptide potentially released as DAMP upon cellular damage) recognized by the LRR-RKs PEP1-RECEPTOR (PEPR1) and PEPR2 (*9–11*), nlp20 (a 20-amino acid peptide derived from bacterial, oomycete, and fungal NECROSIS AND ETHYLENE-INDUCING PEPTIDE 1-LIKE PROTEINS) recognized by the LRR-receptor protein RECEPTOR-LIKE PROTEIN 23 (RLP23) (*12*), chitooctaose (CO8, an octamer fragment of fungal cell walls) recognized by the LysM-RKs LYSM-CONTAINING RECEPTOR KINASE 4 (LYK4) and LYK5 (*13*), 3-OH-FA (a bacterial hydroxylated fatty acid) recognized by the S-lectin-RK LIPOOLIGOSACCHARIDE-SPECIFIC REDUCED ELICITATION (LORE) (*14, 15*), and oligogalacturonides (OGs, derived from the plant cell wall) proposed to be recognized by the epidermal growth factor receptor-like-RK WALL-ASSOCIATED KINASE 1 (WAK1) (*16*). Both Pep1 and OGs are considered DAMPs, while the other patterns are PAMPs. Each pattern was applied in four replicate experiments to two-week-old *Arabidopsis thaliana* (hereafter Arabidopsis) seedlings grown in liquid culture, at concentrations either previously used in transcriptomics studies or shown to be saturating for upstream signaling responses (*7, 14, 17–19*). Each pattern was applied to Col-0 wild-type (WT) or cognate receptor mutant seedlings grown in liquid culture and seedlings were flash frozen for RNA extraction at 0, 5, 10, 30, 90, and 180 min post-treatment (Fig. 1A). Note that *wak1* mutants are not viable (*16*), and thus OG treatment was paired with a mock water treatment.

**Fig. 1.**
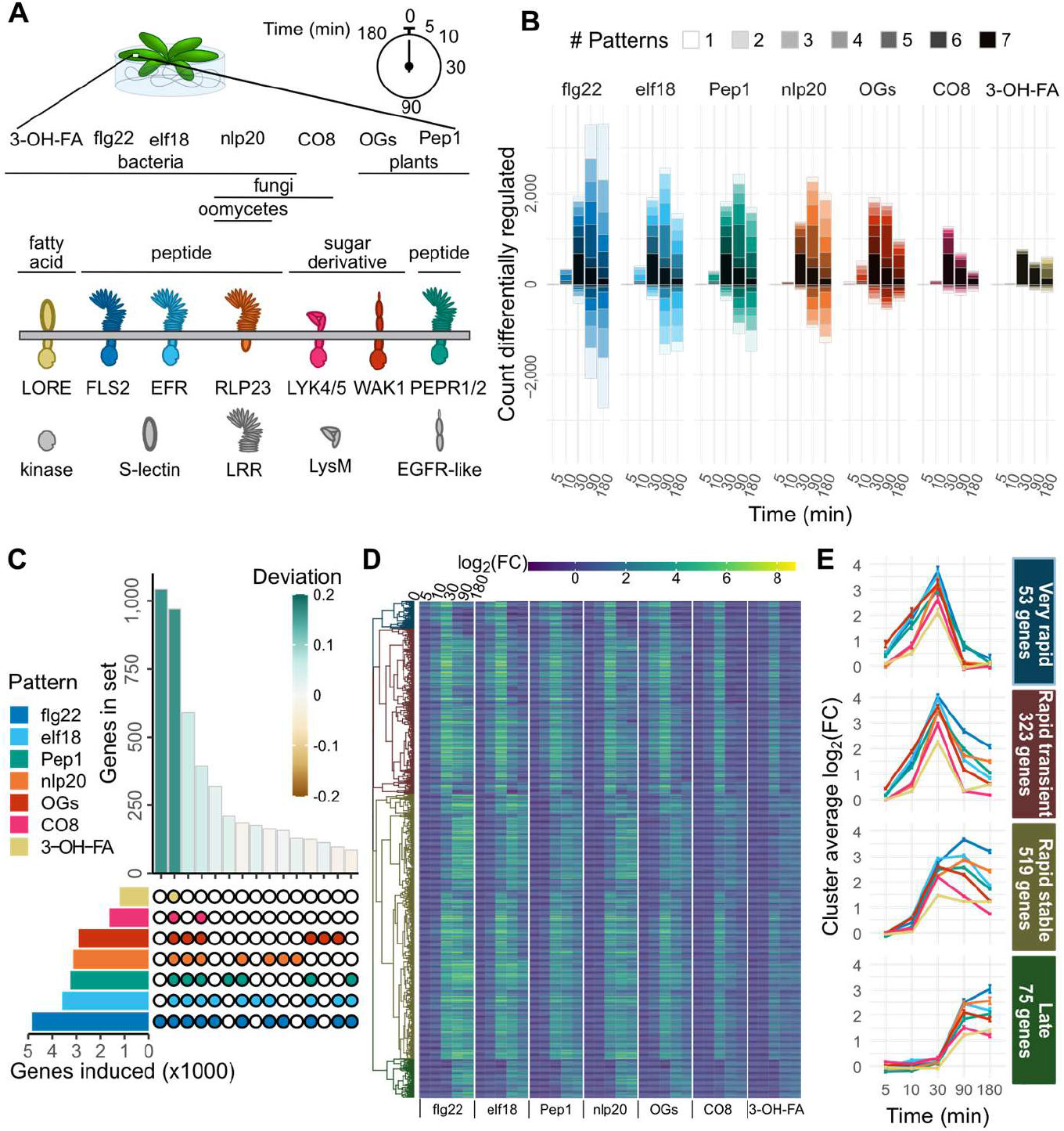
Rapid pattern-triggered transcriptional responses are largely common, with characteristic kinetics. (A) Arabidopsis seedlings were treated with a panel of patterns, and tissue harvested for RNA extraction at indicated times. (B) Genes up- or down-regulated (|log_2_(FC)|>1 and p_adj_<0.05) are shown for each time point within each pattern treatment (total height of bars). Bars are subdivided by the number of patterns affecting each gene set at that time, with darker colors representing more patterns co-regulating. (C) UpSet diagram showing the size of gene sets induced by each pattern (left, single gene list from all times combined) and the top 15 intersections (bottom right) by size (top right). Bars for set sizes are colored by deviation from size predicted by random mixing. (D) Heat map of expression of the genes commonly induced by all tested patterns. Genes are hierarchically clustered according to their behavior across all pattern/time combinations, and cut into four clusters. (E) Visualization of average log_2_(FC) patterns of the four clusters identified in (D), showing different approximate patterns of expression (time points spaced evenly to visualize early times). Error bars represent standard error of the mean.

Transcript abundance was assessed by RNA-seq and differentially expressed genes (DEGs) were identified by comparison with time 0 [log_2_(fold change, FC) >1, p_adj_<0.05], resulting in a total of 10,730 DEGs throughout the experiment (5,718 up-regulated; 5,012 down-regulated), with the strongest treatment being flg22 (8,451 DEGs; 4,816 up and 3,635 down) and the weakest being 3-OH-FA (1,633 DEGs; 1,246 up and 387 down; Tables S1 & S2; Fig. 1B). One selection criterium for treatments chosen here were saturation of upstream signaling outputs (*e.g*. ROS, Ca^2+^ influx), but it cannot be ruled out that higher concentrations of ‘weaker’ patterns would match responses observed here for ‘stronger’ patterns. Treatments in this study were also selected to match previously published transcriptomics experiments – indeed, log_2_(FC) expression values were similar to those published with single patterns (*3, 6, 7*), supporting the experimental and analysis setups used here (Fig. S1A, B). Principal component analysis (PCA) of DEGs revealed strong responses at 30, 90, and 180 min in WT plants that are absent in receptor mutant or mock controls (Figure S1C). Any genes behaving similarly in WT and controls were removed from further analysis. Similar to the PCA, correlation analysis implicated time post treatment as the main factor determining transcriptome response; WT samples became highly correlated at later time points (Fig. S1D; Pearson correlation at 5 min, 0.08; at 10 min, 0.49; at 30 min, 0.89; at 90 min, 0.80; at 180 min, 0.71).

We then collected the set of DEGs up- or down-regulated by each pattern at each time point, and subdivided these sets by the number of patterns similarly affecting each gene (Fig. 1B). This revealed a large set of DEGs induced by all tested patterns (n=970; Table S3; Fig. 1B, darkest bar segment). Furthermore, with the exception of flg22, no pattern induced or repressed a large number of genes uniquely (Figure S2; Table S4). To ascertain whether there exist sets induced specifically by pattern subclasses (e.g. by PRR type, pattern origin, etc.), we identified DEGs induced or repressed by all possible combinations of patterns (Fig. S3), and determined the extent to which the relative sizes of these sets departed from that of a random assortment of genes among patterns (deviation) (*20*). To avoid potential effects of accelerated or delayed induction, we collected all DEGs induced by a pattern in this experiment, into one representative set. As expected, this confirmed that the largest two sets were DEGs induced uniquely by flg22 (n=1,041) or commonly by all tested patterns (Fig. 1C; Fig. S3). Both of these sets were larger than would be expected by chance (deviation 0.16 for each). The next two largest sets comprised DEGs induced by at least five of the tested patterns – indeed the treatment of CO8 and 3-OH-FA in this experiment were relatively weaker than other patterns (Fig. 1B), suggesting that DEGs in these sets may also be induced by all patterns under specific conditions. Remarkably, none of the pattern subsets we identified *a priori* induced unique sets of DEGs much larger or smaller than would be expected by chance (Fig. S3). Taken together, these results suggest that gene induction within the first three hours mostly constitutes a general pattern-triggered response (against ‘non-self or ‘damaged-self), rather than being pattern- or pattern-subclass-specific.

To explore the set of ~1,000 DEGs up-regulated commonly by all treatments, we first hierarchically clustered these genes according to their log_2_(FC) values for each pattern/time combination (Fig. 1D). This revealed four clusters with characteristic expression patterns, described here as ‘Very rapid’, ‘Rapid transient’, ‘Rapid stable’, and ‘Late’ (Fig. 1E). Interestingly, though all tested patterns induced all DEGs and the overall expression patterns were similar, some differences in timing of gene induction could be observed. Among the ‘Very rapid’ and ‘Rapid’ sets OGs, flg22, elf18 and Pep1 induced gene expression already at 5 min, only detectable in response to nlp20, 3-OH-FA and CO8 after 10 min. This partially correlated with the total number of DEGs up-regulated (Fig. 1B), suggesting a potential relationship between amplitude and rapidity of transcriptional response, similar to that observed in some earlier steps of PTI signaling (*21, 22*). Of note, differences in diffusion cannot be excluded as contributing to this observation. A similar analysis of down-regulated DEGs revealed no similar congruence in pattern response – indeed, most sets had similar sizes to those expected by chance (deviation −0.03 – 0.11). There are approximately 100 DEGs down-regulated by all tested patterns (Table S5). Although this set was not significantly larger or smaller than expected by chance, we nevertheless clustered these genes to identify characteristic expression patterns, finding differences in kinetics similar to up-regulated genes (Fig. S4). Taken together, these results show that expression patterns in response to pattern perception are dominated by a small number of pattern-specific responses, and a large set of commonly-induced genes.

In order to investigate transcriptional regulators controlling this response, we expanded this analysis from the genes up-regulated by all patterns to the entire dataset. As timing was the dominant effect in pattern-induced transcriptional patterns (Fig. 1; Fig. S1), we grouped the up-regulated DEGs by the time at which they first became induced, regardless of the inducing pattern, as previously done in response to other stimuli (*23*). GO term enrichment of these five gene sets supports progressive waves of transcriptional response (Fig. 2A). A *cis*-element enrichment analysis revealed enrichment of binding sites for a large number of WRKY transcription factors (TFs) in the promoters of DEGs first induced at 10-30 min post-elicitation (Fig. 2B). This is in line with the established roles of many WRKY TFs in PTI (*24*). In contrast, among genes first induced at 5 or 10 min post-elicitation, there is enrichment in the binding sites for CALMODULIN-BINDING TRANSCRIPTIONAL ACTIVATORs (CAMTAs; Fig. 2B). TFs of the CAMTA family bind the core element vCGCGb, and are the major transcriptional regulators of the plant general stress response (GSR) – a rapid and transient induction of a core set of genes in response to a wide variety of stimuli (*25–27*). Given the congruence of pattern-induced gene sets, and the presence of CAMTA binding sites in promoters of rapidly up-regulated DEGs, we sought to ascertain the degree to which pattern-induced genes are also affected by varied abiotic stresses. To do this, we utilized the published AtGenExpress dataset of Arabidopsis seedling response to cold, drought, genotoxic stress, heat, osmotic stress, salt, UVB irradiation, or wounding (*28*). We then classified each of the DEGs up-regulated in this study according to (i) the time at which it is first induced, (ii) the number of patterns that induce it throughout the experiment, and (iii) the number of abiotic stresses tested in the AtGenExpress experiment that induce it within 3 h. Plotting each DEG according to these criteria, with the color of the point determined by the maximum log_2_(FC) observed in this study, revealed that rapidly induced genes tend to be strongly induced by all tested patterns, and induced by most tested abiotic stresses (Fig. 2C). This analysis extended the observation of a common set of genes induced by all tested patterns to the conclusion that the rapid transcriptional response to pattern perception is dominated by the GSR. As such, our analysis of transcriptional responses indicated that plant cells mostly respond to ‘stress’.

**Fig. 2.**
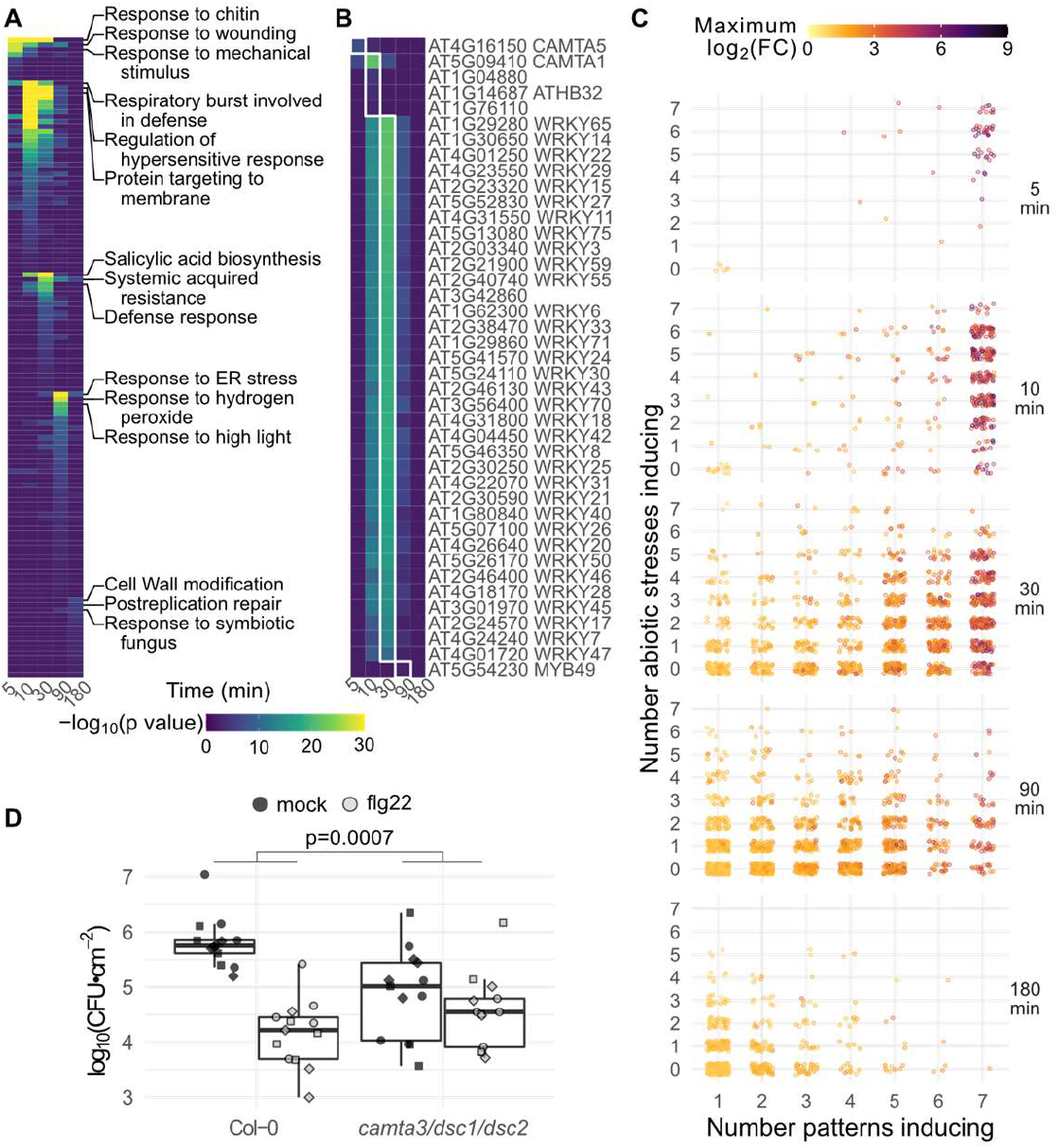
Pattern-triggered transcriptional responses act in time-resolved waves, with the first wave constituting a general stress response important for immune activation. (A) GO term and (B) *cis*-element enrichment analysis of induced genes, categorized according the time point at which they first passed induction threshold, regardless of which pattern caused induction. The top three GO terms for each time point are indicated. (C) Distribution of up-regulated genes. Each gene induced in this study was plotted according to the time it is first induced (panels from top to bottom), the number of tested patterns which induce it (x axis) and the number of abiotic stresses in the AtGenExpress dataset which also induce it within the first three hours (y axis). The color of each dot indicates the maximum log_2_(FC) observed in this study. (D) Box-and-beeswarm plots of flg22-induced resistance to Pto infection. Box plots center on the median, with box extending to the first and third quartile, and whiskers extending to the lesser value of the furthest point or 1.5x the interquartile range. Data were obtained from three independent experiments (point shapes), n=4 per genotype/treatment combination in each experiment. Data were analyzed in R: Two-way ANOVA with experiment as a blocking factor, and p value reports the interaction between treatment and genotype.

A similar analysis of down-regulated DEGs revealed mostly later responses than for up-regulated DEGs, with notably no down-regulated DEGs identified at 5 min (p<0.05). Comparison with gene repression under abiotic stress treatment did not reveal a trend like the GSR; though, interestingly, the most strongly affected genes do tend to be down-regulated commonly by most or all tested patterns (Fig. S5). Finally, while relatively few GO terms or TF binding sites were enriched in down-regulated genes found, many enriched GO terms were associated with growth hormones and response to light, consistent with previous reports that pattern treatment impedes photosynthesis (*29, 30*).

We next sought to test whether the GSR is required for PTI. CAMTA3 is the primary member of the CAMTA family in inducing the GSR (*26*). The genetic analysis of a role of CAMTA3 in PTI is however confounded by the autoimmune phenotype of *camta3* loss-of-function mutants, due at least in part to activation of the two nucleotide-binding leucine-rich repeat receptor proteins (NLRs) DOMINANT SUPPRESSOR OF CAMTA3 1 (DSC1) and DSC2 (*31*). We thus utilized the *camta3/dsc1/dsc2* triple mutant; while WT plants were able to mount an effective flg22-induced resistance to the bacterium *Pseudomonas syringae* pv. tomato DC3000 (Pto), this effect was almost completely lost in the GSR-deficient *camta3/dsc1/dsc2* (p=0.0007, Fig. 2D), consistent with similar results obtained with the dominant-negative *camta3D* allele (*32*). Interestingly, basal susceptibility to Pto was also significantly reduced in *camta3/dsc1/dsc2* compared to WT (p=0.0008, Data S1), in contrast to *camta3D* but in line with studies showing a negative role for CAMTA3 in salicylic acid-mediated immunity regardless of DSC1/2 (*33–35*).

Beyond highlighting the importance of the GSR in PTI, our comparison with AtGenExpress (extended to selected abiotic stress RNA-seq studies) (*36–38*) further identified DEGs up-regulated commonly by all tested patterns, but not by abiotic stresses. Notably, among these 39 ‘core immunity response’ (*CIR*) genes (Table S6), the most strongly up-regulated gene encodes GLUTAMATE RECEPTOR 2.9 (GLR2.9), and *GLR2.7* is also among the *CIR* set. *GLR2.7* and *2.9* are closely related and are present in a tandem repeat on the genome with *GLR2.8* (*39*), which is similarly induced by all tested patterns (Table S3). GLRs are Ca^2+^-permeable channels of which Arabidopsis GLR3 clade members, for example, are key for wound-responsive signaling (*40–42*). In contrast, GLR2 clade members – to which GLR2.7, 2.8 and 2.9 belong – are poorly characterized. Notably, previous pharmacological studies showed that GLRs contribute to pattern-induced Ca^2+^ influx in Arabidopsis (*43*), but the identity of relevant GLRs is still unknown. Given the high sequence similarity between *GLR2.7, 2.8* and 2.9, as well as their chromosomal clustering, we generated a *glr2.7/2.8/2.9* triple mutant using CRISPR-Cas9 in both Col-0 WT and a genetically encoded YELLOW CHAMELEON 3.6 (YC3.6) indicator line. In both backgrounds, this resulted in a large deletion in the *GLR2.7-2.9* genomic region (Fig. S6). Interestingly, the increase of [Ca^2+^]_cyt_ triggered by flg22, elf18 and Pep1 was approximately 25 % reduced in *glr2.7/2.8/2.9* relative to the YC3.6 parental line (Fig. 3A; Fig. S7A). In line with this reduced immune output, *glr2.7/2.8/2.9* plants (in both WT Col-0 and YC3.6 backgrounds) were more susceptible to Pto infection by infiltration, to a similar degree as the immune-deficient *bak1-5* mutant (Fig. 3C) (*44*). Notably, consistent with the specific regulation of *GLR2.7*and *2.9* by pattern perception, but not by abiotic stresses, *glr2.7/2.8/2.9* plants were not impaired in salt-induced [Ca^2+^]_cyt_ increase (Fig. 3B; Fig. S7B). Altogether, these results implicate the GLR2.7/2.8/2.9 clade of GLRs in PTI.

**Fig. 3.**
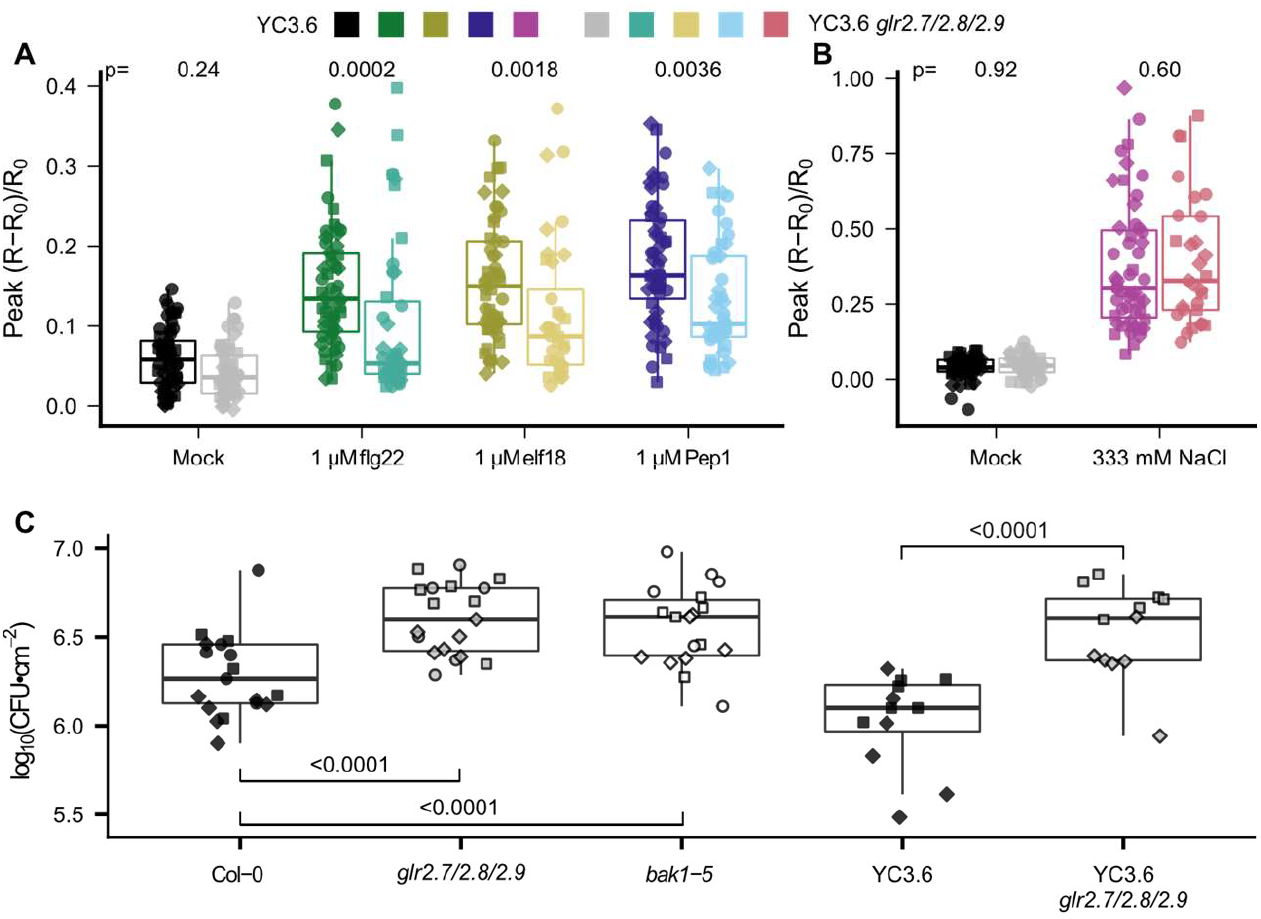
A *glr2.7/2.8/2.9* triple mutant is compromised in pattern-induced Ca^2+^ influx and bacterial disease resistance. (A, B) Parent (darker shades) or *glr2.7/2.8/2.9* (lighter shades) YC3.6 reporter lines were assayed for response to a variety of patterns and salt (NaCl) treatment; peak Ca^2+^ signal reported by YC3.6 within 25 min (patterns) or 1 min (salt) is shown. Each point represents one seedling and different shapes represent 3 independent experiments, n=10-20 for each experiment/line/treatment combination. (C) Parent and *glr2.7/2.8/2*.9 mutants in Col-0 and YC3.6 background were assayed for bacterial susceptibility, alongside the hypersusceptible *bak1-5* mutant. Colony forming units (CFU) were counted two days post infiltration. Each point represents one infected plant and different shapes represent 3 independent experiments, n=5-7 for each experiment/line/treatment combination. Box plots center on the median, with box extending to the first and third quartile, and whiskers extending to the lesser value of the furthest point or 1.5x the inter-quartile range. Statistical tests were performed in R: ANOVA with experiment as a blocking factor, on square root of peak normalized Ca^2+^ response or log_10_(CFU). Post-hoc tests were performed using the emmeans package in R: In (A) and (B) *glr2.7/2.8/2.9* was compared to parent under each treatment, and in (C) (left), each genotype was compared to Col-0 (dunnettx method) and (right) YC3.6 *glr2.7/2.8/2.9* was compared to YC3.6.

We recently reported that Ca^2+^-permeable channels from another family, OSCA1.3 and 1.7, contribute to pattern-induced stomatal immunity (*45*). In contrast, *glr2.7/2.8/2.9* was not compromised in pattern-induced stomatal closure (Fig. S7C), nor was this mutant more susceptible to Pto WT or a coronatine-deficient mutant upon surface-inoculation by spraying (Fig. S7D,E). *GLR2.7/2.8/2.9* are not strongly expressed prior to elicitation, and unlike OSCA1.3 and CNGC2/4 – calcium-permeable channels previously shown to play roles in PTI – they do not show strong preference for/against stomatal expression (Fig. S8). Also, the previously reported role of CNGC2/4 is only apparent under specific external [Ca^2+^] conditions (*46, 47*), indicating that additional calcium-permeable channels must be involved in PTI during normal conditions. These findings substantiate the emerging concept that multiple channels belonging to distinct Arabidopsis families (e.g. CNGCs, OSCAs, GLRs) contribute to the overall pattern-induced calcium response observed at the whole plant level.

The *CIR* gene set includes several other genes associated with immunity (Table S6) (*48–51*). We have here shown the utility of this transcriptomic dataset in identifying signaling and regulatory components of general stress and immune responses in Arabidopsis. The future characterization of other *CIR* genes with yet uncharacterized functions or unknown roles in immunity may thus reveal additional PTI players, and for understanding of how the plant transitions from the rapid general stress response to later immunity-specific responses.

## Acknowledgments

We thank Pingtao Ding for sharing protocols and material related to Tn-5 tagmentation; Stefanie Ranf for providing the *sd1-29* mutant and 3-OH-FA pattern prior to publication; Gary Stacey for providing the *lyk4/5* mutant; Carlos J. S. Moreira for assistance genotyping the *glr2.7/2.8/2.9* CRISPR line; and past and present members of the Zipfel laboratory for helpful discussions.

## Funding

This research was supported by the Gatsby Charitable Foundation, the University of Zurich, the European Research Council under the grant agreements no. 309858 and 773153 (grants ‘PHOSPHOinnATE’ and ‘IMMUNO-PEPTALK’ to CZ), and the Swiss National Science Foundation (grant agreement no. 31003A_182625 to CZ). MB was partially supported by the European Union’s Horizon 2020 Research and Innovation Program under Marie Skłodowska-Curie Actions (grant agreement no.703954).

## Author contributions

CZ, TN, and MB conceived and designed the experiments. CZ and MB obtained funding. MB and PP performed the experiments and analyzed the data. TN contributed conceptually to the study and also provided reagents. MB and CZ wrote the manuscript with feedback from all authors.

## Competing interests

The authors declare no competing interests.

## Data and materials availability

The RNA-seq datasets generated and analyzed in the current study have been deposited in the ArrayExpress database at EMBL-EBI (www. ebi. ac. uk/arrayexpress) under accession number E-MTAB-9694.

## Supplementary Materials

## Materials and Methods

### Arabidopsis growth conditions

For *in vitro* culture Arabidopsis seeds were surface-sterilized, stratified 3-5 days at 4 °C, then plated on full-strength MS medium, 1 % sucrose 0.8 % agar. Plates were placed at 22 °C, 16 h/8 h light/dark. After four days, germinated seedlings were transferred to liquid culture. For RNA-seq, seedlings were placed, two-per-well, in 24-well plates with 1 mL of MS media lacking agar, and plates were sealed with porous tape. For seedling Ca^2+^ measurements, seedlings were transferred, 30-50 per plate, to sterile 9 cm petri dishes containing ca. 25 mL MS media lacking agar, and plates were sealed with porous tape.

For soil growth Arabidopsis seeds were lightly surface-sterilized, stratified 3-5 days, and planted on soil. Plants were grown for four-to-five weeks at 20 °C, 60 % humidity, 10 h/14 h light/dark before assays were performed.

Lines used in this project include Col-0 used as WT control, *fls2c* (SAIL_691_C04) (*52*), *efr-1* (SALK_044334) (*7*), *pepr1-1/2-1* (SALK_059281/SALK_036564) (*10*), *rlp23-1* (SALK_034225) (*12*), *lyk4/5* (WiscDsLox297300_01C/SALK_131911c, seeds obtained from Gary Stacey) (*13*), *sd1-29* (*lore*, SAIL_857_E06, seeds obtained from Stefanie Ranf) (*15*), *bak1-5* (BAK1^C408Y^) (*44*), *camta3/dsc1/dsc2* (SALK_001152/SAIL_49_C05/FLAG014A11, seeds obtained from Morten Petersen) (*NB*: while the FLAG collection was generated in the Ws-2 background, containing a mutated *FLS2*, the *camta3/dsc1/dsc2* line contains a Col-0-version, functional *FLS2* gene) (*31*), and YC3.6 (obtained from Myriam Charpentier). The *glr2.7/2.8/2.9* lines generated in this study are described in Fig. S6.

### RNA-seq treatment

Each plate contained an equal number of wells of Col-0 wild type and PRR mutant control, with the exception of a single plate for combined OG/mock treatment. After nine days growth in liquid MS medium, sealing tape was removed from plates, media removed from wells, and replaced with 0.6 mL liquid MS per well. The following day, when seedlings were 14 days post-stratification, 400 μL of 2.5x pattern solution was added to each well. Two wells, for a total of four seedlings, were harvested for each genotype/treatment/time combination. Final pattern concentrations were 1 μM fg22 (*17, 52*) (Scilight-Peptide), 1 μM elf18 (*7*) (Scilight-Peptide), 1 μM Pep1 (*53*) (Scilight-Peptide), 1 μM nlp20 (*12*) (provided by Thorsten Nürnberger), 100 μg/mL OGs DP10/15 (*3, 54*) (elicityl GAT114), 1 μM CO8 (*18*) (IsoSep 57/12-001), and 1 μM 3-OH-FA (*14*) (provided by Stefanie Ranf).

### Tissue harvest, library preparation, and sequencing

Samples were collected and libraries prepared using a combination of published high-throughput protocols (*55–58*). Briefly, two wells per genotype/treatment/time combination were pooled at each of 0, 5, 10, 30, 90, or 180 min following treatment. Seedlings were blotted dry and flash-frozen in liquid nitrogen. Tissue was pulverized while frozen via two one-minute pulses in a BioRad TissueLyser, and divided in half for library preparation. Divided powder was further disrupted for one minute, prior to addition of extraction buffer, and disrupted in buffer for a further two one-minute pulses. Samples were spun down and lysate collected and incubated with biotin-oligo-dT and streptavidin magnetic beads. The full set of RNA washes and elution was performed twice, with DNAse I treatment in-between, to minimize rRNA and gDNA contamination. cDNA synthesis was performed as described, with the exception that only 2 μL of DNA Pol I was used. Serapure-cleaned dscDNA was quantified via SYBR-green based plate assay and normalized to 2 ng/μL for tagmentation (*59, 60*). Tagmentation was performed in 5 μL reactions containing 0.2 μL Tn-5 transposase, and the entire reaction used as template for PCR (*57*). PCR was performed using in-house primers to add 5’ and 3’ tags and the NEBnext 2x polymerase mix, amplifying for 10 cycles. Libraries were again Serapure cleaned, SYBR quantified, and normalized to 0.5 μM for pooling and sequencing. Pooled libraries were run on 2-3 flowcells of a NextSeq500, and pooling adjusted after each run to maximize overall read density per sample.

### Read mapping and differential expression analysis

Read data was analyzed using FastQC, trimmed (*61*), and mapped to the Arabidopsis TAIR10 genome via TopHat2 (*62, 63*). Mapped reads were assigned to genes, and differential expression analysis performed using DESeq2 (*64*). Prior to differential expression analysis, a total of 17/336 libraries were removed from later analysis, primarily for poor sequencing leading to few mapped reads. For each sample, differential expression was determined relative to the same genotype-treatment combination at time 0. To account for time and mechanical stress, for WT samples, genes were removed if also differentially expressed in PRR mutant controls, with the exception of OG-treated samples, which were filtered based on differential gene expression in mock-treated WT. For data exploration (e.g. PCA, correlation, GO term and *cis*-element enrichment) a relatively loose cutoff of |log_2_(FC)|>1, p_adj_<0.1 was used to obtain a broad landscape of DEGs. For analyses in which specific genes of interest would be analyzed (*e.g*. CIR gene set), a more stringent cutoff of |log_2_(FC)|>1, p_adj_<0.05 was used. Data manipulation was done in R (*65, 66*), using functions from the tidyverse (*67*).

### Exploratory data analysis

Principal component analysis was performed using the prcomp function in R and sample correlation was determined via the Pearson method, using the cor function in R. Visualization of genes induced by various combinations of patterns was done via user-modified adaptations of the UpSetR and SuperExactTest R packages (*68, 69*), and deviation was calculated as described (*20*). Expression of the core set of genes up- or down-regulated by pattern treatment was clustered using the hclust function, with extra functionality from the dendextend package in R (*70*).

Gene induction specific to individual pattern treatments was determined using a modification of tissue-specific gene expression assignment (*71, 72*). Briefly, normalized pseudocount data were first filtered to genes significantly upregulated (p<0.1, log_2_(FC)>1) in at least one condition. Filtered pseudocounts were next averaged across all replicates, then summed across all time points for each pattern. For each gene and each pattern the fraction of total counts for that gene attributed to that pattern was calculated (specificity measure, SPM). Data were finally filtered to those genes with SPM>0.33 for at least one pattern (approximately 1/3 total reads in experiment attributable to one pattern).

### GO term and *cis*-element enrichment

GO term enrichment was performed using the library TopGO in R, using GO terms obtained from TAIR, searching for enrichment in each gene set relative to the complete set of genes detected in this experiment, and determining enriched GO terms using the weight01 method with the Fisher statistic (*73*). *cis*-element enrichment analysis was performed using AME, part of the MEME suite (*74*), using a published library of TF binding sites found via DAPseq (*75*).

### Comparison with AtGenExpress abiotic stress microarray data analysis

As the AtGenExpress experiment was performed using the ATH1 microarray, we first restricted induced genes to those present on the array. Abiotic stress microarray data was obtained from http://jsp.weigelworld.org/AtGenExpress/resources/ in 2017 and analyzed using limma (*NB*: data are no longer hosted here, but CEL files can be downloaded through https://www.arabidopsis.org/portals/expression/microarray/ATGenExpress.jsp) (*28, 76*). We did not consider the oxidative stress treatment for filtering pattern-responsive genes, as most patterns induce production of reactive oxygen species. To facilitate comparisons with this study’s RNA-seq data, only time points from the first three hours were considered, and comparisons for differential expression were first made between each treatment and time 0, then between each treatment and mock at the same time, considering only genes that were differentially expressed [log_2_(FC)>1, p_adj_<0.05] under both criteria.

*CIR* genes were selected according to the following criteria: (i) significantly induced in at least one time by all seven patterns tested here (ii) not significantly induced at any time point by any of the selected stresses in the AtGenExpress Dataset (iii) Uniquely targeted by at least one probe in the ATH1 microarray (iv) not significantly induced in selected abiotic stress experiments (3 hr proteotoxic stress, 4 h darkness, 4 h flooding, 3 h 50, 150, or 200 mM NaCl) assayed using RNA-seq (*36–38*). This resulted in a set of 40 DEGs. Among these, one highly upregulated gene, *AT3G32090*, is a suspected pseudogene with strong homology to *WRKY40* in one region. All of the reads assigned to *AT3G32090* mapped to only this region. As *WRKY40* is both highly expressed and strongly upregulated by pattern treatment, we suspected these reads were mistakenly assigned to *AT3G32090*, and removed it from the *CIR* set.

### Measurement of intracellular Ca^2+^ concentration in seedlings

After five days growth in liquid MS medium, sealing tape was removed from plates and seedlings rinsed in sterile water and transferred one-per-well to black 96-well plates containing 150 μL sterile water. Seedlings were gently pressed to ensure the majority of the seedling was submerged, and plates were incubated in the dark under bench conditions overnight. The following day, when seedlings were 11 days post-stratification, plates were imaged in a Tecan SPARK microplate reader at two conditions: excitation 440 nM, emission 480 nM (CFP) and excitation 440 nM emission 530 nM (YFP). Pattern treatment was performed through addition of 38 μL of 5x solution injected after 5 min visualization by the microplate reader. The focal plane for fluorescence measurements was set to a single point in the center of each well, and moved up 0.5 mm post-injection to accommodate increased volume in wells. Despite this adjustment, overall fluorescence intensity and thus ratio was frequently altered post-injection, as seedlings did not uniformly fill well. Due to this change, and the generally slow pattern response, we normalized all subsequent fluorescence ratios to the first ratio measured post-injection (R_0_), as (R-R_0_)/R_0_. Wells were manually rejected if preinjection fluorescence was not stable or vastly different than R_0_. Salt (NaCl) treatment was performed similar to pattern treatment, with the following changes: to accommodate the faster response, injection and imaging was performed on a well-by-well basis rather than across a subsection of the plate. Due to the faster response, the first measurement postinjection already reflects the beginning of plant response - R_0_ was thus defined as pretreatment fluorescence ratio, though this resulted in more noise in the final data.

As some silencing was observed both in parent YC3.6 lines and YC3.6 *glr2.7/2.8/2.9* lines, only seedlings with visible fluorescence at 5 d were transferred to liquid culture, and following treatment, only seedlings (wells) with pre-treatment fluorescence in both wavelengths greater than 3x that of a non-fluorescent Col-0 control were considered. Total seedlings imaged were as follows: YC3.6 mock: 56, YC3.6 flg22: 54, YC3.6 elf18: 52, YC3.6 Pep1: 55, YC3.6 *glr2.7/2.8/2.9* mock: 48, YC3.6 *glr2.7/2.8/2.9* flg22: 43, YC3.6 *glr2.7/2.8/2.9* elf18: 36, YC3.6 *glr2.7/2.8/2.9* Pep1: 43, YC3.6 mock (NaCl): 56, YC3.6 NaCl: 51, *glr2.7/2.8/2.9* mock (NaCl): 38, YC3.6 *glr2.7/2.8/2.9* NaCl: 29.

### Bacterial infection assays

For all infection assays, Arabidopsis plants were treated when four-to five-week-old, and bacteria grown overnight in Kings B medium liquid culture, refreshed via a 1-2 h subculture in the morning, spun down and resuspended in 10 mM MgCl_2_. For induced resistance (*52*), three leaves from each plant were infiltrated with either 1 μM flg22 or water in the morning. The following morning, selected leaves were re-infiltrated with *Pseudomonas syringae* pv. tomato DC3000 (Pto) expressing luciferase (*77*) at OD_600_=0.0002 or ~1×10^5^ colony-forming units (CFU)/mL. Plants were covered and infection allowed to proceed for two days. For infiltration infection assays, infection was performed similarly with the following differences: WT Pto was used rather than the luciferase-expressing strain; trays were incubated uncovered; and there was no mock or pattern pretreatment. For spray infection, Pto was diluted to OD_600_=0.2 or ~1×10^8^ CFU/mL in MgCl_2_, Silwet L-77 added to 0.04 %, and plants sprayed to surface saturation (~4 mL per plant). For all infection assays, after approximately 48 h leaf discs were collected (infiltration: two from each infiltrated leaf; spray: 6 from 6 separate leaves), ground in 10 mM MgCl_2_, and serial dilutions from 1×10^-1^ to 1×10^-5^ plated to count CFU.

Following infection, log_10_(CFU) follow an approximately normal distribution. ANOVA was performed using the glm and anova functions in R, and post-hoc tests via emmeans package (*78*). Sample numbers are as follows: for induced resistance n=12 plants for all genotype/treatment combinations. For infiltration infection total plants counted were Col-0: 17, Col-0 *glr2.7/2.8/2.9*: 19, *bak1-5*: 18, YC3.6: 12, YC3.6 *glr2.7/2.8/2.9*: 12. For spray infection n=18 for all genotype/treatment combinations.

### Stomatal aperture measurements

For each experiment, three leaf discs were taken from each of 6 plants per line. The three leaf discs were floated one-per-well in 100 μM stomatal opening buffer (10 mM MES-KOH pH 6.15, 50 mM KCl, 10 μM CaCl_2_, 0.01 % Tween-20) in white 96-well plates for 2 h in the growth chamber. Subsequently, one leaf disc from each plant was treated with 5 μM flg22, 10 μM ABA, or mock through addition of stock solution to stomatal opening buffer. Leaf discs were incubated 2-3 h further, then imaged on a Leica DMR microscope and photographed with the equipped Leica DFC320 camera. Stomata length and width were annotated in ImageJ. The experiment was repeated twice. Total number of stomata counted per genotype/treatment combination are as follows: Col-0 mock: 581; Col-0 flg22: 529; Col-0 ABA: 519; *glr2.7/2.8/2.9* mock: 461; *glr2.7/2.8/2.9* flg22: 503; *glr2.7/2.8/2.9* ABA: 426; *bak1-5* mock: 567; *bak1-5* flg22: 607; *bak1-5* ABA: 719.

Stomatal aperture (width/length) followed an approximately square normal distribution. ANOVA was performed on square-root transformed ratios using the glm and anova functions in R, and post-hoc tests via emmeans package (*78*).

### Data and code availability

The RNA-seq datasets generated and analyzed in the current study have been deposited in the ArrayExpress database at EMBL-EBI (www.ebi.ac.uk/arrayexpress) under accession number E-MTAB-9694. Markdowns documenting the steps in filtering, visualizing, and analyzing the data in all figures and tables are available in Data S1. Data S1 also contains raw data for Figures 2, 3, and S7.

### Tissue expression patterns of genes encoding calcium channels implicated in PTI

Tissue-specific expression datasets containing aerial (rosette) tissue were selected in Genevestigator, comprising datasets from (*79–82*).

**Fig. S1.**
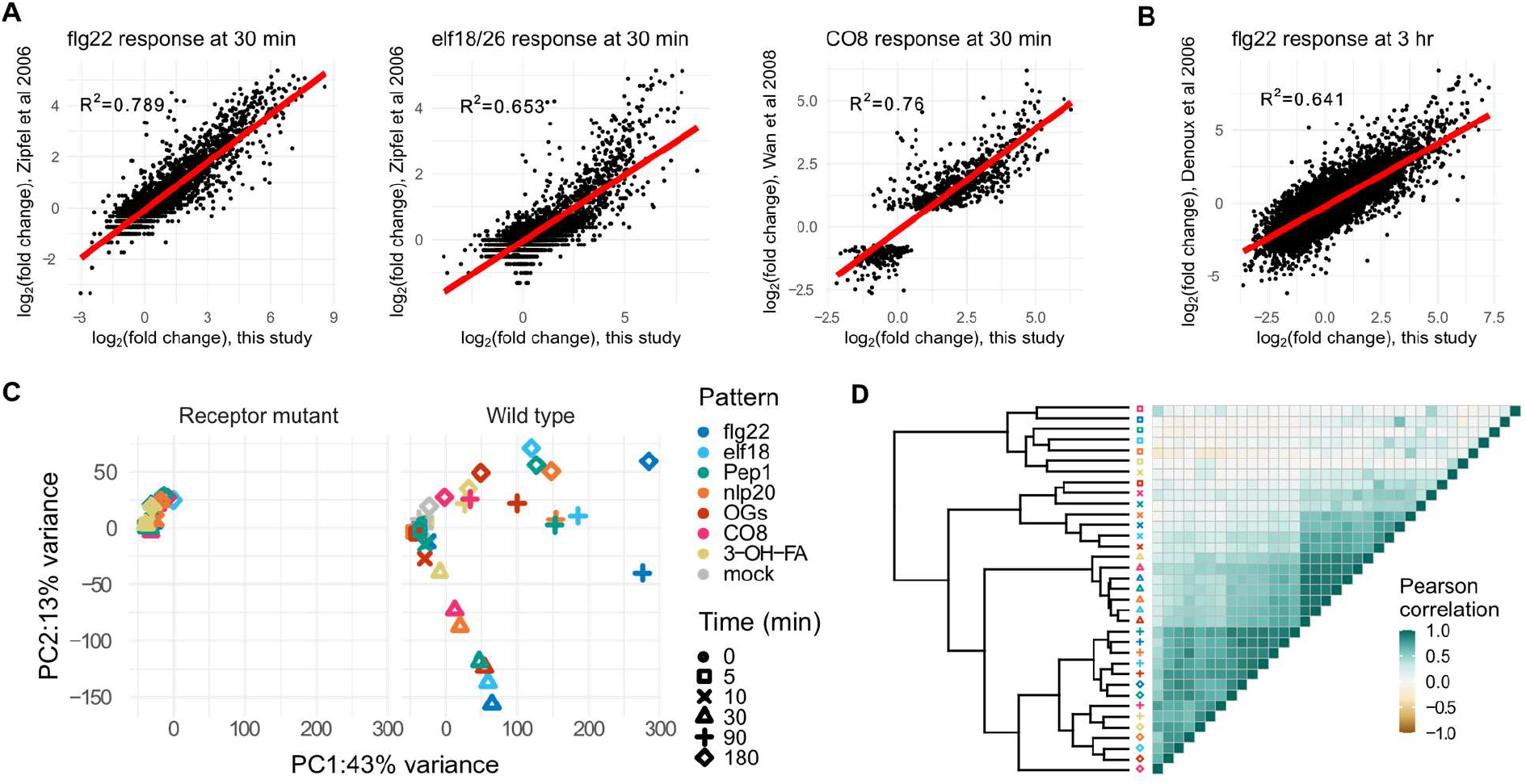
Quality control and exploratory analysis of RNA-seq data. Expression changes in this study at (A) 30 min and (B) 3 h are plotted against previously published results for flg22, elf18/26, and chitooctaose (CO8). Linear correlation shown in red, with R^2^ (linear regression) shown on each plot. (C) PCA analysis of log_2_(FC) of differentially expressed genes, showing (left) minimal changes in receptor-mutant treated plants, mostly corresponding with later time points, and rays of response (right) corresponding with plants at 30, 90, or 180 min post-treatment. (D) Pearson correlation heatmap of DESeq2-calculated log_2_(FC) showing clustering largely by time point, with the strongest correlations at 30 min.

**Fig. S2.**
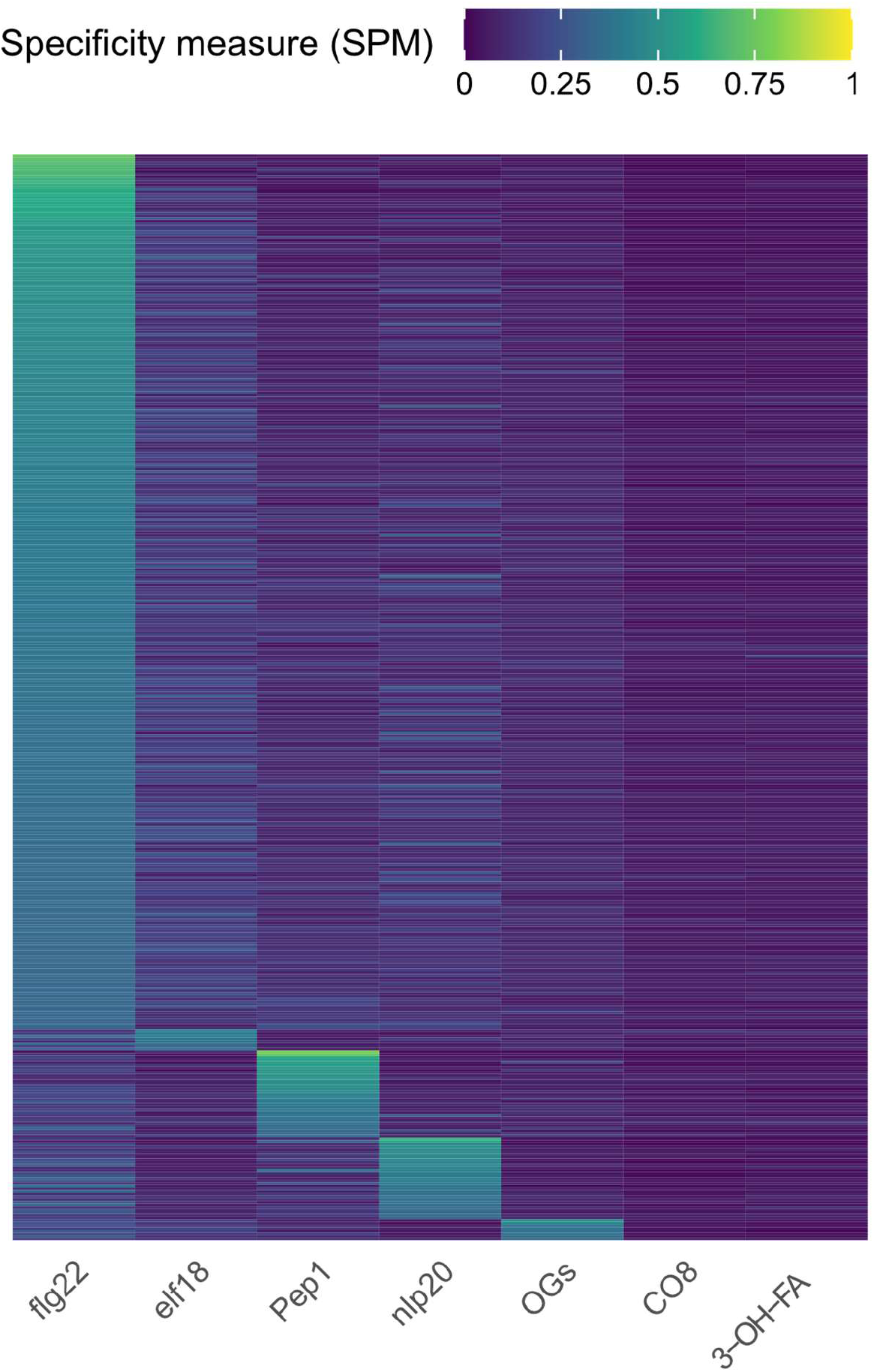
There is little specificity in pattern-induced genes. Among induced genes, for each pattern a specificity measure (expression in response to pattern/total expression in experiment) was calculated, and genes with at least one SPM>0.33 (one pattern treatment responsible for approximately 1/3 total expression in study, n=412) were gathered. flg22 is the only pattern treatment with a large number of pattern-selective genes expressed (flg22: 332, elf18: 8, Pep1: 33, nlp20:31, OGs: 8, CO8 and 3-OH-FA:0)

**Fig. S3.**
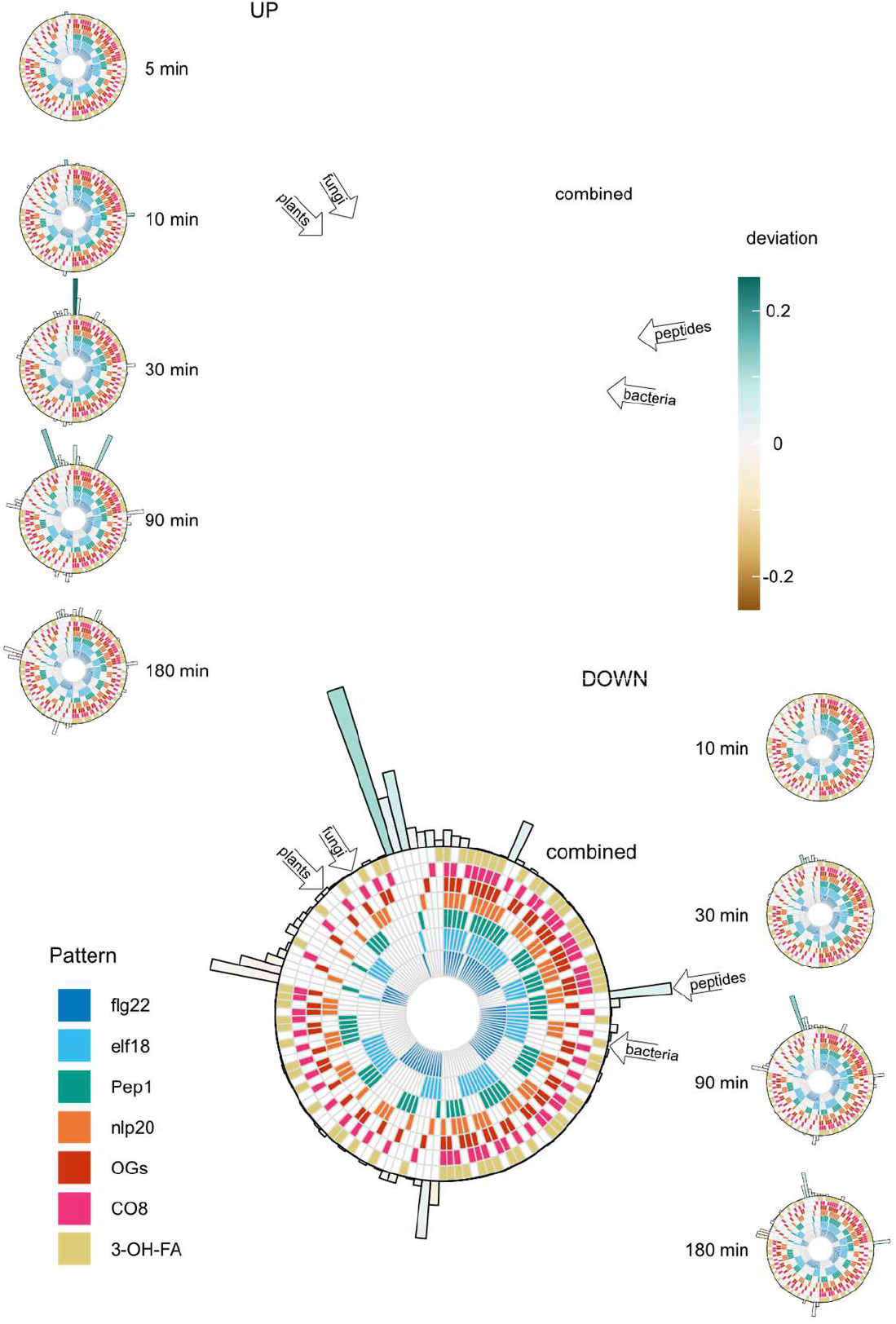
Complete complement of set sizes among collapsed pattern-induced and pattern-repressed gene sets. Each circular ‘track’ represents one pattern treatment; when filled the pattern in question alters the expression of the gene set shown at the perimeter. Gene set size is shown via bar height of bars surrounding pattern tracks, and bar color shows deviation: indicating whether the set size is larger or smaller than would be expected by chance. Large diagrams show the overall set complement of genes induced or repressed by patterns taking all time points into account, whereas smaller diagrams to the left and right are specific for the complement of genes induced or repressed at the indicated time point. No genes were significantly repressed at five minutes posttreatment. Selected pattern subset of *a priori* interest are highlighted through open arrows on large combined plots; none has deviation far from 0.

**Fig. S4.**
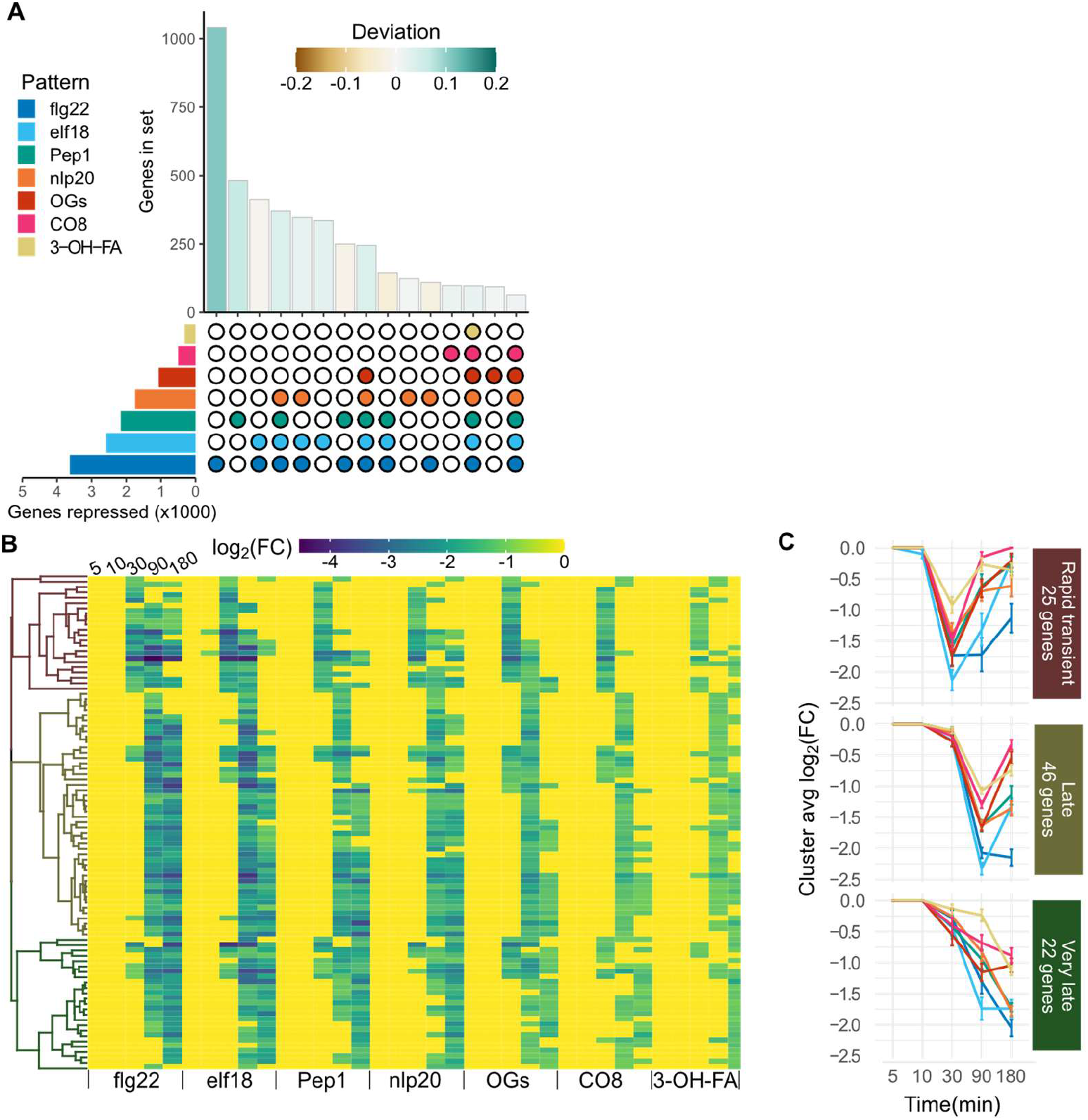
Pattern-responsive genes tend to be repressed by single patterns, though there does exist a core set of 93 genes repressed by all tested patterns with at least three patterns of expression. A single set of genes repressed [log_2_(FC)<−1, p<0.05] by each pattern treatment was found through combining the lists genes repressed at each time. (A) UpSet diagram showing the size of ‘collapsed’ gene sets repressed by each pattern (left) and the top 15 intersections (bottom right) by size (top right), colored by deviation from set size predicted by random mixing. (B) Heat map of expression of the 93 genes repressed by all tested patterns. Genes are hierarchically clustered according to their behavior across all pattern/time combinations, and cut into three clusters. (C) Visualization of average log_2_(FC) patterns of the three clusters identified in (B), showing different patterns of expression.

**Fig. S5.**
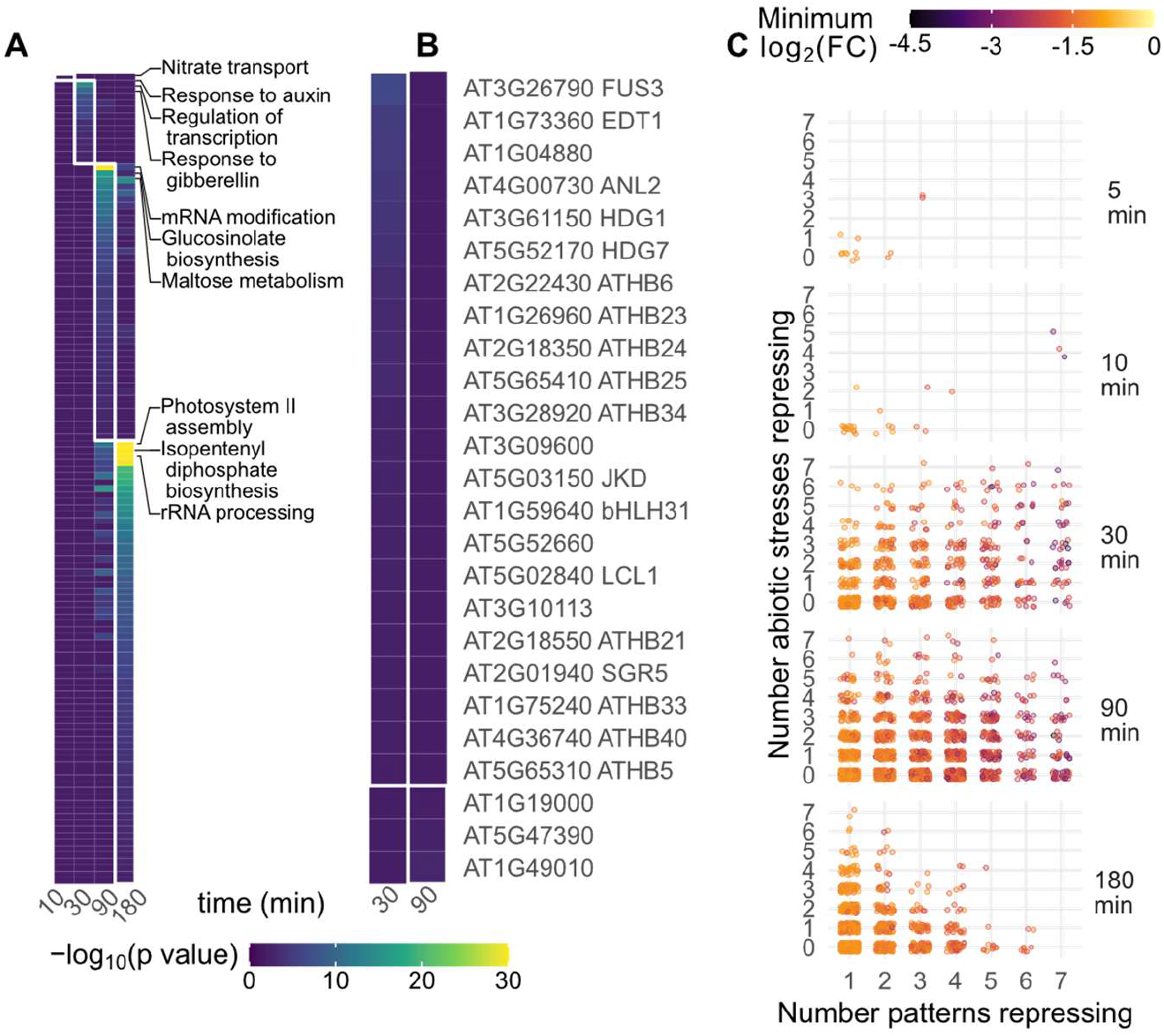
Pattern-triggered transcriptional repression acts in time-resolved waves. (A) GO term and (B) *cis*-element enrichment analysis of repressed genes, categorized according to the time point at which they first passed significance threshold, regardless of which pattern caused repression. The top three GO terms for each time point are indicated. (C) Distribution of repressed genes. Each gene repressed in this study was plotted according to the time it is first repressed (panels from top to bottom), the number of tested patterns which repress it (x axis) and the number of abiotic stresses in the AtGenExpress dataset which also repress it within the first 3 h (y axis). The color of each dot indicates the most negative log_2_(FC) observed in this study.

**Fig. S6.**
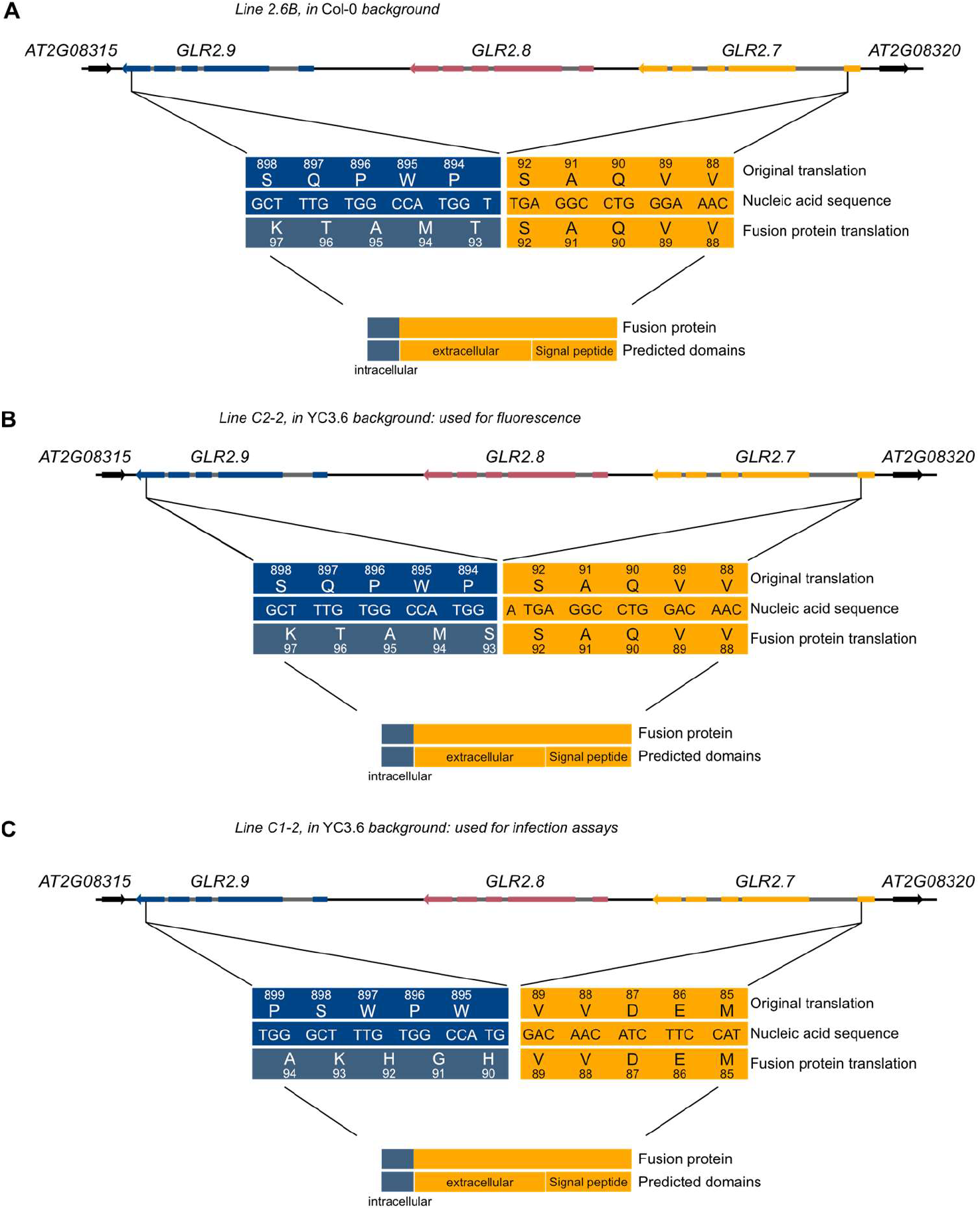
CRISPR deletes the majority of the *GLR2.7/2.8/2.9* genomic region in assayed lines. Schematic of the *GLR2.7/2.8/2.9* genomic region, with deletions in (A) Col-0, (B) and (C) YC3.6 background. In each case, a ‘fusion protein’ may be transcribed, consisting of approximately 90 (*92, 92, 89*) amino acids of GLR2.7, fused to approximately 12 (*12, 13, 12*) nonsense amino acids from the *GLR2.9* genomic region. The potential fusion protein does not encode any transmembrane domains. GLR exons are represented by colored boxes, introns by grey boxes, and intergenic regions by black lines. Neighboring genes shown in black. Arrows represent direction of transcription.

**Fig. S7.**
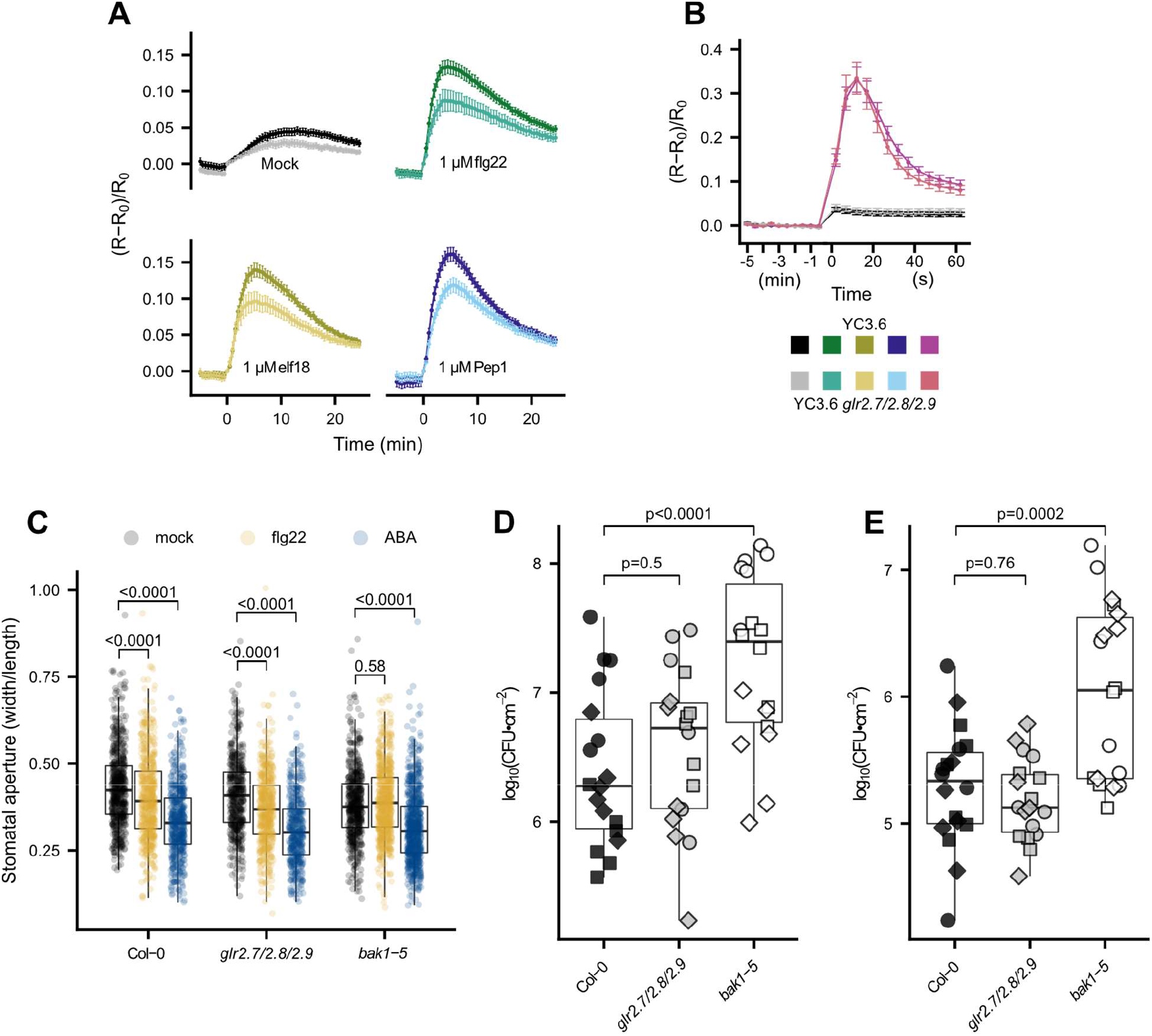
Characterization of *glr2.7/2.8/2.9* lines. (A, B) Increase in intracellular Ca^2+^ concentration in response to pattern treatment. Shown are mean corrected YFP/CFP ratio within (A) 25 min or (B) 1 min post-treatment (timepoint 0). Data were collected every 30 s (A) or 5 s (B), and corresponding peak values are shown in Figure 3. (C) Stomatal aperture of WT, *glr2.7/2.8/2.9*, or flg22-hyporesponsive *bak1-5* plants treated with water, 5 μM flg22, or 10 μM ABA. Each point represents one stoma, and plot represents stomata from a total of 12 plants assayed over 5 experiments (n=36-178 stomata per genotype/treatment/experiment). Statistical tests were performed in R, two-way ANOVA blocking by experiment. Post-hoc tests were performed using the emmeans package in R: within each genotype, stomatal aperture was compared with mock treatment with dunnettx multiple testing correction. In spray infection assays *glr2.7/2.8/2.9* are not more susceptible to (D) WT Pto DC3000, or (E) Pto *COR^-^*, deficient in the stomata-opening toxin coronatine. Bacteria were harvested from leaf discs two days post-inoculation; each point represents one plant, and shapes represent three independent experiments (n=6 plants per genotype/treatment/experiment). Statistics were performed in R: one-way ANOVA blocking by experiment followed by dunnettx multiple comparison to Col-0 performed using the emmeans package. Box plots center on the median, with box extending to the first and third quartile, and whiskers extending to the lesser value of the furthest point or 1.5x the inter-quartile range.

**Fig. S8.**
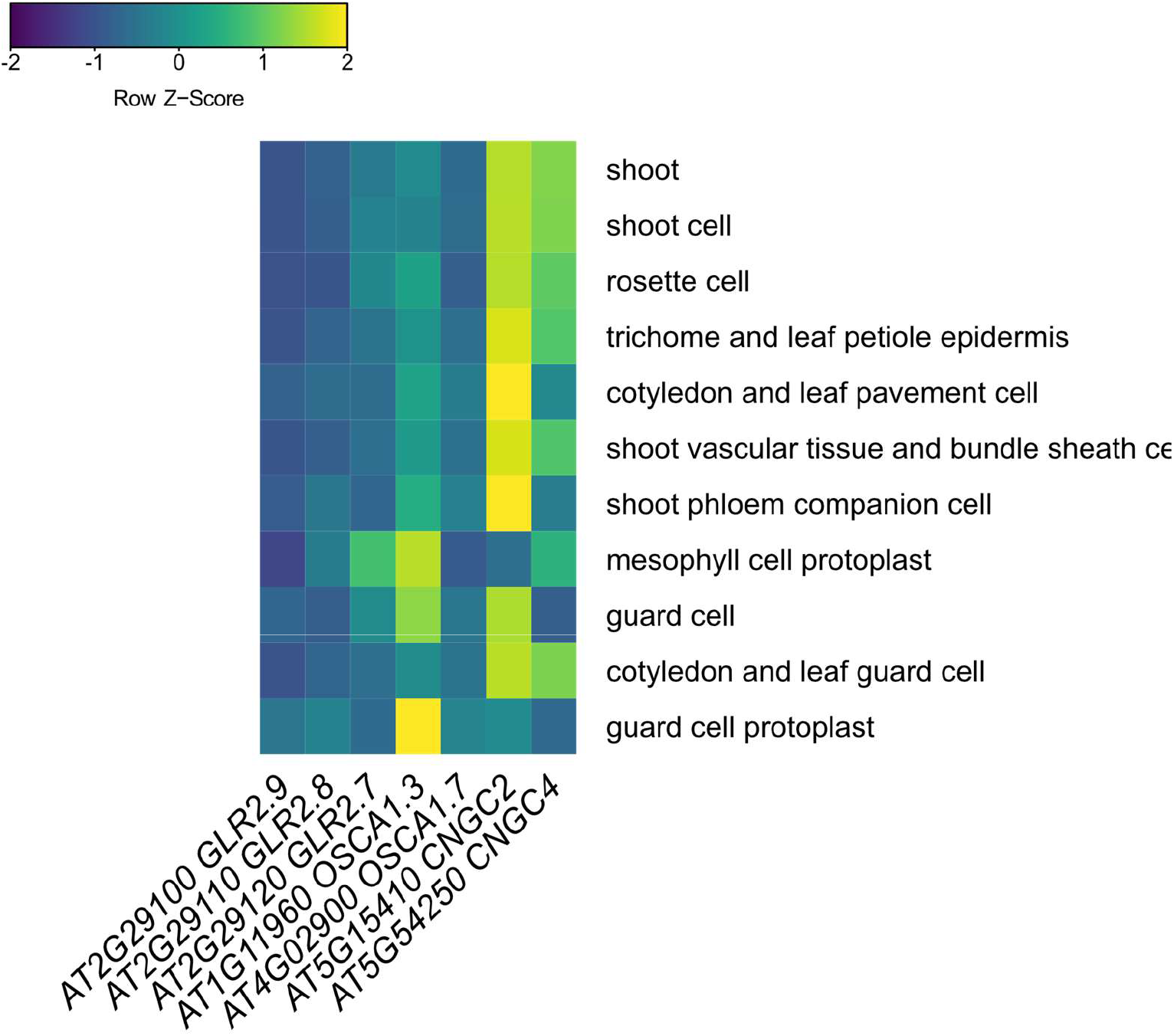
Leaf tissue expression patterns of genes encoding calcium-permeable channels implicated in PTI. Data collected from Genevestigator, and scaled by each experiment.

**Table S6.**
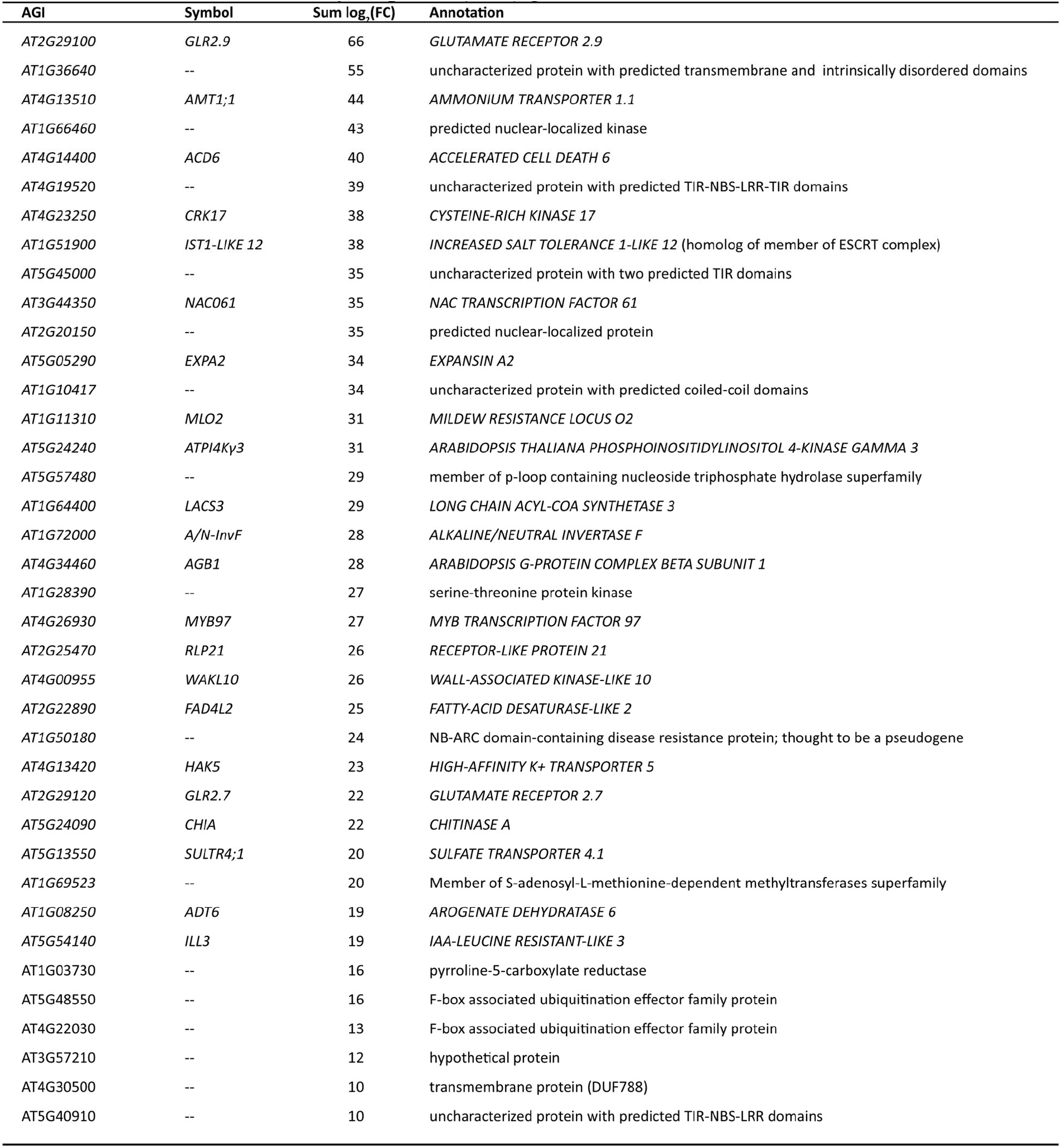
List of core immunity response (CIR) genes

Table S1. (separate file) Log_2_ fold change values for each gene calculated by DESeq for each condition relative to time 0.

Table S2. (separate file) P values for differential expression calculated by DESeq2 for each condition relative to time 0.

Table S3. (separate file) Genes upregulated commmonly by all tested patterns. Additional information provided: Common name and functional description when available (TAIR10). Cluster: which cluster gene belongs to in this study’s clustering analysis (Figure 1). Number of treatments among selected abiotic stress treatments performed via RNAseq (AbioRNAseqNumber) and in the AtGenExpress dataset (AtGenExpress Number) that also upregulate this gene. For AtGenExpress, data additionally provided if gene not present or not uniquely targeted on ATH1 array. base_counts: some low-expressed genes may gain a high log2(FC) from small variation in counts – this metric shows the average counts for this gene across all genotypes/treatments at time 0 to allow confidence in degree induction observed.

Table S4. (separate file) Specificity measure (SPM) values for genes found to be significantly induced selectively by one of the tested patterns.

Table S5. (separate file) Genes downregulated commonly by all tested patterns. Additional information provided: Common name and functional description where available (TAIR10). Cluster: which cluster gene belongs to in this study’s clustering analysis (Figure S4).Number of treatments among selected abiotic stress treatments performed via RNAseq (AbioRNAseqNumber) and in the AtGenExpress dataset (AtGenExpress Number) which also downregulate this gene. For AtGenExpress, data additionally provided if gene not present or not uniquely targeted on ATH1 array.

Data S1. (separate file) Rmarkdown files documenting steps of data analysis and visualization (.pdf)

## Notes

### Competing Interest Statement

The authors have declared no competing interest.

### Summary of Updates

Addition of previously missing supplemental files.

