## Supplemental Data S1 for "The transcriptional landscape of *Arabidopsis thaliana* pattern-triggered immunity"

### Figure1

Marta Bjornson

```
knitr::opts_chunk$set(echo = TRUE)
library(tidyverse)
library(cowplot)
library(rlang)
library(ggforce) # for theme_no_axes
require(cowplot) # for plot_grid
library(ggpubr) # for as_ggplot
library(dendextend)
library(DESeq2)
library(bioDist) # for spearman.dist
library(viridis)
library(scales) # for commas in axis text Panel B
```

This R Markdown documents the sections of Figure 1

Panel 1A does not include any R calculations

### PanelB

Figure 1b: counts up and down by time. First step is making a table as used as input for UpSet: https://www.ncbi.nlm.nih.gov/pmc/articles/PMC4720993/ (https://www.ncbi.nlm.nih.gov/pmc/articles/PMC4720993/)

```
rm(list=ls())
home <- "D:/OneDrive/Norwich_Z/Overall/CleanedData/wtonly/"
#Palette modified from https://personal.sron.nl/~pault/
definedvibrant <- c("0"="white", "flg22"="#0077BB", "elf18"="#33BBEE", "nlp20"="#EE7733", "Pep1"="#00998
8", "OGs"="#CC3311", "CO8"="#EE3377", "3-OH-FA"="#DDCC77", "mock"="#BBBBBB")

#Gathering lists of genes generated by DESeq2: |L2FC|>1, p<0.05, generated by comparison to time0 and fi
ltered to those responding specifically in wt and not receptor mutant.
SingleTimeTable <- function(time, direction){
  flg <- read.csv(paste0(home, "flg22_", time, direction,".csv"), header=F)
  elf <- read.csv(paste0(home, "elf18_", time, direction,".csv"), header=F)
  pep <- read.csv(paste0(home, "Pep1_", time, direction,".csv"), header=F)
  nlp <- read.csv(paste0(home, "nlp20_", time, direction,".csv"), header=F)
  OGs <- read.csv(paste0(home, "OGs_", time, direction,".csv"), header=F)
  CO8 <- read.csv(paste0(home, "ch8_", time, direction,".csv"), header=F)
  LPS <- read.csv(paste0(home, "LPS_", time, direction,".csv"), header=F)

  totalgenes <- flg %>% full_join(elf, by="V1") %>% full_join(pep, by="V1") %>% full_join(nlp, by="V1")
 %>% full_join(OGs, by="V1") %>% full_join(CO8, by="V1") %>% full_join(LPS, by="V1") %>% filter(V1 != "n
ull")

  timetable <- data.frame(ATG=totalgenes$V1, flg=0, elf=0, pep=0, nlp=0, OGs=0, CO8=0, LPS=0)
  timetable$flg[timetable$ATG %in% flg$V1] <- 1
  timetable$elf[timetable$ATG %in% elf$V1] <- 1
  timetable$pep[timetable$ATG %in% pep$V1] <- 1
  timetable$nlp[timetable$ATG %in% nlp$V1] <- 1
  timetable$OGs[timetable$ATG %in% OGs$V1] <- 1
  timetable$CO8[timetable$ATG %in% CO8$V1] <- 1
  timetable$LPS[timetable$ATG %in% LPS$V1] <- 1

  timetable <- column_to_rownames(timetable,var="ATG")
  return(timetable)
}

up005 <- SingleTimeTable("005","up")
up010 <- SingleTimeTable("010","up")
up030 <- SingleTimeTable("030","up")
up090 <- SingleTimeTable("090","up")
up180 <- SingleTimeTable("180","up")

#no genes significantly downregulated at 5 minutes
#down005 <- SingleTimeTable("005","down")
down010 <- SingleTimeTable("010","down")
down030 <- SingleTimeTable("030","down")
down090 <- SingleTimeTable("090","down")
down180 <- SingleTimeTable("180","down")
```

From the UpSet table, find the number of elicitors commonly affecting each gene up- or down-regulated by each elicitor at each time

```
#generating a data frame that will hold information on degree and cardinality of all sets for each time
Direction <- c(rep("up",245), rep("down",245))
Time <- rep(c(rep("005",49),rep("010",49), rep("030", 49),rep("090", 49),rep("180",49)),2)
Elicitor <- rep(c(rep("flg22",7), rep("elf18",7), rep("Pep1",7), rep("nlp20",7), rep("OGs",7), rep("CO8"
,7), rep("3-OH-FA",7)),10)
Degree <- rep(seq(1:7),70)
Cardinality <- rep(0,490)
bar <- data.frame(Direction, Time, Elicitor, Degree, Cardinality)

#Filling in that table
FillInBar <- function(timetable, direction, time, pass){
  bar <- pass
  elf <- timetable[timetable$elf==1,]
  for(i in 1:7){
    bar$Cardinality[bar$Elicitor== "elf18" & bar$Time== time & bar$Degree==i & bar$Direction== directio
n] <- length(rowSums(elf)[rowSums(elf)==i])
  }

  flg <- timetable[timetable$flg==1,]
  for(i in 1:7){
    bar$Cardinality[bar$Elicitor== "flg22" & bar$Time== time & bar$Degree==i & bar$Direction== directio
n] <- length(rowSums(flg)[rowSums(flg)==i])
  }

  pep <- timetable[timetable$pep==1,]
  for(i in 1:7){
    bar$Cardinality[bar$Elicitor== "Pep1" & bar$Time== time & bar$Degree==i & bar$Direction== direction]
<- length(rowSums(pep)[rowSums(pep)==i])
  }


  nlp <- timetable[timetable$nlp==1,]
  for(i in 1:7){
    bar$Cardinality[bar$Elicitor== "nlp20" & bar$Time== time & bar$Degree==i & bar$Direction== directio
n] <- length(rowSums(nlp)[rowSums(nlp)==i])
  }

  OGs <- timetable[timetable$OGs==1,]
  for(i in 1:7){
    bar$Cardinality[bar$Elicitor== "OGs" & bar$Time== time & bar$Degree==i & bar$Direction== direction]
 <- length(rowSums(OGs)[rowSums(OGs)==i])
  }


  CO8 <- timetable[timetable$CO8==1,]
  for(i in 1:7){
    bar$Cardinality[bar$Elicitor== "CO8" & bar$Time== time & bar$Degree==i & bar$Direction== direction]
 <- length(rowSums(CO8)[rowSums(CO8)==i])
  }


  LPS <- timetable[timetable$LPS==1,]
  for(i in 1:7){
    bar$Cardinality[bar$Elicitor== "3-OH-FA" & bar$Time== time & bar$Degree==i & bar$Direction== directi
on] <- length(rowSums(LPS)[rowSums(LPS)==i])
  }
  return(bar)

}
bar <- FillInBar(up005, "up","005",bar)
bar <- FillInBar(up010, "up","010",bar)
bar <- FillInBar(up030, "up","030",bar)
bar <- FillInBar(up090, "up","090",bar)
```

```
bar <- FillInBar(up180, "up","180",bar)
#No genes significantly downregulated at 5 minutes
#bar <- FillInBar("down","005",bar)
bar <- FillInBar(down010, "down","010",bar)
bar <- FillInBar(down030, "down","030",bar)
bar <- FillInBar(down090, "down","090",bar)
bar <- FillInBar(down180, "down","180",bar)

#Make the downregulated DEG bars extend downwards
bar$Cardinality[bar$Direction=="down"] <- -bar$Cardinality[bar$Direction=="down"]

#Some parameter wrangling for better graphing
bar$Time <- as.character(bar$Time)
bar$Time[bar$Time=="005"] <- "5"
bar$Time[bar$Time=="010"] <- "10"
bar$Time[bar$Time=="030"] <- "30"
bar$Time[bar$Time=="090"] <- "90"
bar$Time[bar$Time=="180"] <- "180"
bar$Time <- factor(bar$Time, levels=c("5","10","30","90","180"))
bar$Degree <- as.factor(bar$Degree)
bar$Elicitor <- factor(bar$Elicitor, levels=c("flg22","elf18","Pep1","nlp20","OGs","CO8","3-OH-FA"))
```

Making Panel B

```
PanelB <- ggplot(bar, aes(x=Time, y=Cardinality, group=Degree)) +
  geom_bar(aes(fill=Elicitor), stat="identity", position=position_stack()) +scale_fill_manual(values=def
inedvibrant, guide="none") +
  geom_bar(aes(alpha=Degree), fill="black", stat="identity", position=position_stack()) +
  geom_bar(aes(alpha=as.factor(8-as.numeric(Degree))), fill="white", stat="identity", position=position_
stack(), color="grey", size=0.05)  +
  scale_alpha_manual(values=c(0, 0, 0, 0, 0.3, 0.5, 0.9))+
  guides(alpha=guide_legend(ncol=7))+
  theme_minimal(base_size=6) +
  theme(axis.text.x=element_text(angle=70, hjust=1, size=5),
        axis.text.y=element_text(angle=90, hjust=0.5, size=5),
        strip.text=element_text(size=6),
        panel.background = element_blank(), plot.background = element_blank(), legend.background = eleme
nt_blank(), legend.box.background = element_blank(), panel.border = element_blank(),
        legend.box=NULL, legend.key.size=unit(0.2, "cm"), legend.position = "top", legend.text=element_t
ext(size=5), legend.title = element_text(size=7),
        axis.title=element_text(size=7)) +
  labs(y= "Count differentially regulated", x="Time (min)", alpha="# Patterns") +
  scale_y_continuous(label=comma, limits=c(-3600, 3600)) +
  facet_grid(.~Elicitor)

ggsave("PanelB.pdf", PanelB, height=6, width=6.55, units="cm", bg="transparent")
```

### PanelC

A lot of work to re-make the plots that can be made by UpSetR, adding color for elicitor sets and information on deviation, as available in the online tool version. First define all functions: extract all intersections (of all degrees). Code modified from stack overflow post by useR: https://stackoverflow.com/questions/47083311/r-extract-sets-in-arbitrary-intersection-from-a-data-frame (https://stackoverflow.com/questions/47083311/r-extract-sets-in-arbitrary-intersection-from-a-data-frame) First define all functions

```
#This function takes a table in UpsetFormat, and makes it into a long table for inter_sets. Could probab
ly combine, but so much going on in that function already I'd rather not
lengthen <- function(upsettable){
  longtable <- upsettable %>% gather(-ATG, key="elicitor",value="value")
  longtable$TF <- FALSE
  longtable$TF[longtable$value==1] <- TRUE

  colnames(longtable) <- c("ATG","group","numeric","value")
  longtable$numeric <- NULL
  return(longtable)
}


#master function to get set sizes and deviation: takes long table format with T/F for gene induced by el
icitor or not
inter_sets = function(time, direction){
  #this function parses a text string (A&B&!C) and finds the number of rows that actually meet those cri
teria: taken from StackOverflow with slight alteration. It's within inter_sets so I don't have to pass t
he 'long' table
  filter_sets = function(filter_expr){
    long %>%
      spread(group, value) %>%
      filter(!!parse_quosure(filter_expr)) %>%
      nrow()
  }

  #This function I wrote: it uses the definition of Deviation from UpSet (expected size of overlap given
the size of all sets participating and the size of all sets not participating). I checked it against val
ues calculated online and it seemed to be performing as expected. It's within inter_sets so I don't have
to pass the 'setsizes' data frame
  calculate_deviation = function(elicitors){
      use <- colnames(setsizes)[colnames(setsizes) %in% elicitors]
      yes <- setsizes[,use]
      yes <- yes/n
      yes <- prod(yes)

      dont <- colnames(setsizes)[!colnames(setsizes) %in% elicitors]
      if(length(dont) >0){
        no <- setsizes[,dont]
        no <- no/n
        no <- 1-no
        no <- prod(no)
      }else{
        no <- 1
      }

      product <- prod(yes, no)

      cardinality <- as.numeric(elicitors['value'])/n

      deviation <- cardinality - product
      deviation
  }



  upset <- SingleTimeTable(time, direction) %>% rownames_to_column("ATG")
  long <- lengthen(upset)

  (setsizes <- data.frame('flg'=nrow(upset[upset$flg==1,]),
                          'CO8'=nrow(upset[upset$CO8==1,]),
                          'elf'=nrow(upset[upset$elf==1,]),
                          'LPS'=nrow(upset[upset$LPS==1,]),
```

```
                          'nlp'=nrow(upset[upset$nlp==1,]),
                          'OGs'=nrow(upset[upset$OGs==1,]),
                          'pep'=nrow(upset[upset$pep==1,]))) #number induced by each individually regard
less of overlaps
  n <- nrow(upset) #total numberinduced

  #this is the second part of the code taken from StackOverflow - this generates all possible combinatio
ns of A&B&C, A&B&!C, A&!B&C, etc, as strings, then reduces them to non-redundant set
  combins = unique(long$group) %>%
    c(paste0("!", .)) %>%
    combn(length(.)/2) %>%
    t() %>%
    as.data.frame() %>%
    filter(apply(., 1, function(x) length(unique(gsub("!", "", x))) == ncol(.) & !(length(grep("!", x))
 %in% c(0, ncol(.))))) %>%
    unite("expressions", names(.), sep = " & ")
  #I added this, it is the all elicitor set
    combins <- rbind(combins, paste(unique(long$group), sep = ' & ', collapse = ' & '))
  #I had added this for an earlier version, it's the no elicitor set. Could be useful again depending on
what I want to look at
    #combins <- rbind(combins, paste(paste0("!",unique(long$group)), sep = ' & ', collapse = ' & '))

  #Here add the size of any interaction to the string describing that interaction
  combins$value = sapply(combins$expressions, filter_sets)
  combins$value <- as.numeric(combins$value)

  #For my code need elicitors each in own column
  separated <- combins %>% separate(expressions, into = c('a','b','c','d','e','f','g'), sep=' & ')

  # Calculate deviation, given binary presence/absence of elicitor and size of overlap set
  separated$deviation <- apply(separated, 1, calculate_deviation)
  #Make recombined column with set name all together
  recombined <- separated %>% unite(expression, a,b,c,d,e,f,g)

  return(recombined)
}

#for a given elicitor, make a binary presence/absence column
elicitor_columns=function(deviationdata, elicitor){
  deviationdata$new <- 1
  deviationdata$new[grep(paste0("!",elicitor),deviationdata$expression)] <- 0
  colnames(deviationdata)[colnames(deviationdata)=="new"] <- elicitor
  return(deviationdata)
}
```

Then generate data: here just for all times collapsed together, upregulated genes

```
#Again, the 'collapsed' list of upregulated genes was made externally by collecting all genes induced at
any time by any elicitor, in wt but not receptor mutants or mock
doit <- inter_sets("Collapsed","up") #get all intersections, deviations
```

```
## Warning: `parse_quosure()` is deprecated as of rlang 0.2.0.
## Please use `parse_quo()` instead.
## This warning is displayed once per session.
```

```
doit <- elicitor_columns(doit, "flg") #then make a column for each elicitor: 1 if included in set, 0 if
 not
doit <- elicitor_columns(doit, "elf")
doit <- elicitor_columns(doit, "nlp")
doit <- elicitor_columns(doit, "pep")
doit <- elicitor_columns(doit, "OGs")
doit <- elicitor_columns(doit, "CO8")
doit <- elicitor_columns(doit, "LPS")
doit$degree <- rowSums(doit[4:10])#calculate the degree
doit <- arrange(doit, desc(value))
doit$expression <- factor(doit$expression, ordered=T)#this should allow me to control the arrangment of
 the x axis but it doesn't work
#This was for the Extended Data figure setup: keep it because it's still useful for controlling the arra
ngement of the x axis
doit$start_angle <- seq(0,2*pi-(2*pi/127), length.out=127)
doit$end_angle <- seq(2*pi/127, 2*pi, length.out=127)
#this is mostly just for making the fills different colors, there's probably a better way to do it
doit$flg[doit$flg==1] <- "flg22"
doit$elf[doit$elf==1] <- "elf18"
doit$nlp[doit$nlp==1] <- "nlp20"
doit$pep[doit$pep==1] <- "Pep1"
doit$OGs[doit$OGs==1] <- "OGs"
doit$CO8[doit$CO8==1] <- "CO8"
doit$LPS[doit$LPS==1] <- "3-OH-FA"
```

And finally make the plot.

```
#make plots for vertical set sizes bars
vertical <- ggplot(doit[1:15,], aes(x=as.factor(start_angle), y=value, fill=deviation))+
  geom_bar(stat="identity", color="grey", size=0.15)+
  theme_classic(base_size=7)+
  theme(axis.title.x=element_blank(), axis.text.x=element_blank(), axis.text.y=element_text(angle=90, hjust=0.5), legend.position="none",
        rect=element_rect(fill="transparent",color=NA), plot.background = element_blank(), panel.background = element_blank())+
  scale_y_continuous(label=comma, expand=c(0,0), limits=c(0, 1100), name="Genes in set")+
  scale_fill_distiller(palette="BrBG", direction=1, limits=c(-0.2, 0.2))

vert <- ggplot(doit[1:15,], aes(x=as.factor(start_angle), y=value, fill=deviation))+
  geom_bar(stat="identity")+theme_minimal(base_size=5)+
  scale_fill_distiller(palette="BrBG", direction=1, limits=c(-0.2, 0.2), guide=guide_colorbar(barheight=0.3))+
  theme(legend.position = "top", legend.key.size=unit(0.25,"cm"))+labs(fill=expression("Deviation\n"))
deviat <- as_ggplot(get_legend(vert))


#make plot for horizontal elicitor sizes bars
elicitorsizes <- SingleTimeTable("collapsed", "up")
setsizes <- data.frame('flg22'=nrow(elicitorsizes[elicitorsizes$flg==1,]), 'CO8'=nrow(elicitorsizes[elicitorsizes$CO8==1,]), 'elf18'=nrow(elicitorsizes[elicitorsizes$elf==1,]),'3-OH-FA'=nrow(elicitorsizes[elicitorsizes$LPS==1,]), 'nlp20'=nrow(elicitorsizes[elicitorsizes$nlp==1,]), 'OGs'=nrow(elicitorsizes[elicitorsizes$OGs==1,]), 'Pep1'=nrow(elicitorsizes[elicitorsizes$pep==1,])) #number induced by each individually regardless of overlaps
setsizes <- setsizes %>% pivot_longer(everything(),names_to="elicitor", values_to="ngenes")
setsizes$elicitor <- gsub("X3.OH.FA","3-OH-FA",setsizes$elicitor)
setsizes$elicitor <- factor(setsizes$elicitor,
                            levels=c("flg22","elf18","Pep1","nlp20","OGs","CO8","3-OH-FA"))
horizontal <- ggplot(setsizes, aes(x=ngenes/1000, y=reorder(elicitor, -ngenes), fill=elicitor))+
  geom_bar(stat="identity")+
  theme_classic(base_size=7)+
  theme(axis.title.y=element_blank(), axis.text.y = element_blank(), legend.position="none",
        axis.line.y=element_blank(), axis.ticks.y=element_blank(),panel.background=element_rect(fill="transparent"),
        plot.background=element_rect(fill="transparent"), plot.margin = margin(0, 0, 0,0, "pt"))+
  scale_x_reverse(expand=c(0,0), limit=c(5,0), name="Genes induced (x1000)")+
  scale_fill_manual(values=definedvibrant)

#make a simple version, just to grab the legend
horiz <- ggplot(setsizes, aes(x=ngenes/1000, y=reorder(elicitor, -ngenes), fill=elicitor))+
  geom_bar(stat="identity")+labs(fill="Pattern")+
  scale_fill_manual(values=definedvibrant)+
  theme_minimal(base_size=7)+
  theme(legend.key.size=unit(0.25,"cm"))
elic <- get_legend(horiz)

#make plots for which elicitor sets dots
doit_long <- doit %>% pivot_longer(c(flg, elf, nlp,pep,OGs, CO8, LPS), names_to="elicitor", values_to="present")
doit_long$elicitor <- factor(doit_long$elicitor, levels=c("flg","elf","pep","nlp","OGs","CO8","LPS"))
dots <- ggplot(doit_long[1:105,], aes(x=start_angle, y=elicitor, fill=as.factor(present)))+
  geom_point(shape=21, size=1.5)+
  scale_fill_manual(values=definedvibrant)+
  theme_no_axes()+
  theme(panel.background = element_blank(), plot.background = element_blank(), panel.border=element_blank(),
        legend.position="none", plot.margin = margin(0, 0, 0,0, "pt"))


#Combine all the bits
upset <- ggdraw()+
```

```
  draw_plot(vertical,x=0.23, y=0.38, width=0.8, height=0.62)+
  draw_plot(horizontal, x=0.04, y=0.005, width=0.36, height=0.385)+
  draw_plot(dots, x=0.38, y=0.09, height=0.3, width=0.625)+
  draw_plot(deviat, x=0.65, y=0.85, height=0.1, width=0.2)+
  draw_plot(elic, x=0.03, y=0.55, height=0.3, width=0.15)

suppressWarnings(ggsave(upset, filename = "PanelC.pdf", height=6, width=4, units="cm", bg="transparent"
))
```

### Panel D

Clustering the core 1000 PAMP-induced genes based on L2FC in all elicitor/time combinations. First load in all data, and then reduce to the genes upregulated commonly by all elicitors

```
#get UpSet table for all times collapsed, one set of genes for each elicitor
up_all <- SingleTimeTable("Collapsed", "up")

#step back up a level, return to DESeq results objects to calculate L2FC fresh
home="D:/OneDrive/Norwich_Z/Overall/CleanedData/"

AllL2FC <- function(TrtTime){
  filepath=paste0(home,"ResultsObjects/",TrtTime,"_results")
  load(filepath)
  resdf <- as.data.frame(res)
  allFCs <- subset(resdf, select = "log2FoldChange")
  allFCs$log2FoldChange <- as.numeric(allFCs$log2FoldChange)
  names(allFCs) <- TrtTime
  return(allFCs)
}

PAMPTime <- c("Col_LPS_005","Col_LPS_010","Col_LPS_030","Col_LPS_090","Col_LPS_180","Col_ch8_005","Col_c
h8_010","Col_ch8_030","Col_ch8_090","Col_ch8_180","Col_elf18_005","Col_elf18_010","Col_elf18_030","Col_e
lf18_090","Col_elf18_180","Col_flg22_005","Col_flg22_010","Col_flg22_030","Col_flg22_090","Col_flg22_18
0","Col_nlp20_005","Col_nlp20_010","Col_nlp20_030","Col_nlp20_090","Col_nlp20_180","Col_OGs_005","Col_OG
s_010","Col_OGs_030","Col_OGs_090","Col_OGs_180","Col_Pep1_005","Col_Pep1_010","Col_Pep1_030","Col_Pep1_
090","Col_Pep1_180")
log2FClist <- lapply(PAMPTime, AllL2FC)
log2FCdf <- do.call("cbind", log2FClist)
nrow(log2FCdf)
```

```
## [1] 26397
```

```
up_all <- up_all %>% rownames_to_column("ATG") %>% filter(flg==1 & elf==1 & pep==1 & nlp==1 & OGs==1 & C
O8==1 & LPS==1) %>% select("ATG")

log2FC <- log2FCdf[rownames(log2FCdf) %in% up_all$ATG,]

nrow(log2FC)
```

```
## [1] 970
```

Cluster by spearman (no scaling/centering)

```
D <- spearman.dist(as.matrix(log2FC))
#x <- x %>% rownames_to_column(var="ATG")
Gene_column <- log2FC %>% rownames_to_column("ATG")

uptree <- hclust(D)
```

After exploratory data analysis, chose 4 clusters

```
#First make dendrogram
num = 4

clusters <- cutree(uptree, k=num)
cluster.df <- as.data.frame(clusters) %>% rownames_to_column("ATG")
#Re-ordering dendrogram by speed of response
updend <- as.dendrogram(uptree) %>% rotate(c(cluster.df$ATG[cluster.df$clusters==2], cluster.df$ATG[clus
ter.df$clusters==1], cluster.df$ATG[cluster.df$clusters==3], cluster.df$ATG[cluster.df$clusters==4]))
gene_order <- updend %>% labels
updatedorder <- c("#225522","#666633", "#663333","#07415A") #colors for dendrogram, modified from Paul T
ol dark qualitative
updend <- updend %>% set("branches_k_color", k=num, value=updatedorder)%>% set("branches_lwd", 0.15)

#arrange data for heatmap
longL2FC <- Gene_column %>% pivot_longer(-ATG, names_to="sample", values_to="L2FC") %>% separate(sample,
into=c("Genotype","elicitor","time"))
#fix elicitor names (will change in Affinity Design later anyway)
longL2FC$elicitor <- gsub("LPS","3-OH-FA",longL2FC$elicitor)
longL2FC$elicitor <- gsub("ch8","CO8",longL2FC$elicitor)
longL2FC$elicitor <- factor(longL2FC$elicitor, levels=c("flg22","elf18","Pep1","nlp20","OGs","CO8","3-OH
-FA"))
x_labs <- c("|   ","","flg22","","   |",
            "","","elf18","","   |",
            "","","Pep1","","   |",
            "","","nlp20","","   |",
            "","","OGs","","   |",
            "","","CO8","","   |",
            "","","3-OH-FA","","   | ")

#print approximate heatmap here, so I know what it will look like
p1 <- ggplot(longL2FC, aes(x=interaction(time, elicitor), y=factor(ATG, levels=gene_order), fill=L2FC))+
  geom_tile()+scale_fill_viridis()+
  scale_x_discrete(labels=x_labs)+
  theme_minimal(base_size=8)+
  theme(axis.ticks=element_blank(), axis.title = element_blank(), axis.text.y=element_blank())

#and tree plot
p2 <- ggplot(updend, horiz=T, labels=F)+
  theme(plot.margin = margin(0, 0, 0,0, "pt") )+
  scale_x_continuous(expand=c(0.01,0.01))+scale_y_reverse(expand=c(0,0)) +
  theme(rect=element_rect(fill="transparent", color=NA),
        panel.border=element_blank())
```

```
## Scale for 'y' is already present. Adding another scale for 'y', which will
## replace the existing scale.
```

Now make final plots, merge, and save

```
heatmap <- ggplot(longL2FC, aes(x=interaction(time, elicitor), y=factor(ATG, levels=gene_order), fill=L2
FC))+
  geom_tile()+
  scale_fill_viridis_c(guide=guide_colorbar(barheight=unit(0.1, "cm")))+
  scale_x_discrete(labels=x_labs, expand=c(0,0))+ scale_y_discrete(expand=c(0,0))+
  theme_minimal(base_size=5)+
  theme(axis.ticks=element_blank(), axis.title = element_blank(),
        axis.text.y=element_blank(), rect=element_rect(fill="transparent", color=NA),
        panel.border=element_blank(), legend.position="none")
simpleheat <- ggplot(longL2FC, aes(x=interaction(time, elicitor), y=ATG, fill=L2FC))+
  geom_tile()+scale_fill_viridis(guide=guide_colorbar(barheight=unit(0.1, "cm")))+
  theme_minimal(base_size=4)+
  theme(rect=element_rect(fill="transparent", color=NA),
        panel.border=element_blank(), legend.position="top")
heatlegend <- get_legend(simpleheat)

#and tree plot
tree <- ggplot(updend, horiz=T, labels=F)+
  theme(plot.margin = margin(0, 0, 0,0, "pt") )+
  scale_x_continuous(expand=c(0.01,0.01))+scale_y_reverse(expand=c(0.01,0)) +
  theme(rect=element_rect(fill="transparent", color=NA),
        panel.border=element_blank())
```

```
## Scale for 'y' is already present. Adding another scale for 'y', which will
## replace the existing scale.
```

```
combo <- ggdraw()+
  draw_plot(tree, x=0,width=0.115,y=-0.0469,height=0.972)+
  draw_plot(heatmap, x=0.10, width=0.915, y=0, height=0.93)+
  draw_plot(heatlegend, x=0.3, width=0.6, y=0.9, height=0.1)

ggsave(combo, filename="PanelD.pdf", height=6.5, width=5, units="cm", bg="transparent")
```

### Panel E

Plot average expression each cluster

```
se <- function(x) sqrt(var(x)/length(x))

clusters.df <- data.frame(cluster=clusters, ATG=names(clusters))

addcluster <- merge(clusters.df, longL2FC, by="ATG")

Percluster <- addcluster %>% group_by(cluster, elicitor, time) %>% summarize(mean=mean(L2FC), SE=se(L2F
C))
```

```
## `summarise()` regrouping output by 'cluster', 'elicitor' (override with `.groups` argument)
```

```
Percluster$cluster <- factor(Percluster$cluster, levels=c("4","3","1","2"))

labels <- c("1"="Rapid stable\n519 genes", "2"="Late\n75 genes","3"="Rapid transient\n323 genes", "4"="V
ery rapid\n53 genes")

clusterplot <- ggplot(Percluster, aes(x=as.factor(as.numeric(time)), y=mean, group=elicitor,color=elicit
or))+
  geom_line(size=0.4) +
  geom_errorbar(aes(ymin=mean-SE, ymax=mean+SE), size=0.3, width=0.12)+
  scale_color_manual(values=definedvibrant)+
  theme_minimal(base_size=7)+
  theme(rect=element_rect("transparent",color=NA), legend.position="none", strip.text=element_text(size=
6))+
  facet_grid(cluster~., labeller=labeller(cluster=labels))+
  labs(x="Time (min)",y=expression("Cluster average log"[2]*"(FC)"))

ggsave(clusterplot,filename="PanelE.pdf", height=6.8, width=3, units="cm", bg="transparent")
```

### Figure2

Marta Bjornson

```
knitr::opts_chunk$set(echo = TRUE)
library(tidyverse)
library(GO.db)
library(topGO)
library(org.At.tair.db)
library(cowplot)
library(ggpubr) #for ggarrange
library(viridis)
library(readxl)
library(emmeans)
library(ggthemes)
```

### Panel A

Collect genes according to when first induced, in wt only (JIT, https://www.pnas.org/content/115/25/6494 (https://www.pnas.org/content/115/25/6494)), get the enriched GO terms in each set, arrange them by greatest p value, reconnect and plot. First get all genes signficantly (p<0.1) induced, in wt only. Same as for making Supplementary Table 3, but looser cutoff

```
rm(list=ls())
home <- "D:/OneDrive/Norwich_Z/Overall/CleanedData/"

#get list of genes significantly upregulated specifically in wt at each time
wt_sig_up <- function(wt, mut){
  #first find the genes upregulatd in mutant
  mut.file=paste0(home,"ResultsObjects/",mut,"_results")
  load(mut.file)
  mut.df <- as.data.frame(res)

  mut.up <- mut.df %>% filter(padj<0.1)  %>%  filter(log2FoldChange > 1) %>% rownames_to_column("ATG")

  #then the genes upregulated in  wt, extra step filter out the ones also upregulated in control
  wt.file=paste0(home,"ResultsObjects/",wt,"_results")
  load(wt.file)
  wt.df <- as.data.frame(res)

  wt.sig <- wt.df %>% rownames_to_column("ATG") %>%
    filter(padj<0.1)  %>%
    filter(log2FoldChange > 1) %>%
    filter(!ATG %in% mut.up$ATG)

  #reduce to L2FC and return
  wt.FCs <- subset(wt.sig, select = c("ATG","log2FoldChange"))
  wt.FCs$log2FoldChange <- as.numeric(wt.FCs$log2FoldChange)
  names(wt.FCs) <- c("ATG", wt)
  return(wt.FCs)
}

#list of treatment results objects
PAMPTime <- c("Col_LPS_005","Col_LPS_010","Col_LPS_030","Col_LPS_090","Col_LPS_180",
              "Col_ch8_005","Col_ch8_010","Col_ch8_030","Col_ch8_090","Col_ch8_180",
              "Col_elf18_005","Col_elf18_010","Col_elf18_030","Col_elf18_090","Col_elf18_180",
              "Col_flg22_005","Col_flg22_010","Col_flg22_030","Col_flg22_090","Col_flg22_180",
              "Col_nlp20_005","Col_nlp20_010","Col_nlp20_030","Col_nlp20_090","Col_nlp20_180",
              "Col_OGs_005","Col_OGs_010","Col_OGs_030","Col_OGs_090","Col_OGs_180",
              "Col_Pep1_005","Col_Pep1_010","Col_Pep1_030","Col_Pep1_090","Col_Pep1_180")

#corresponding(!) list of control results objects
mutPAMPTime <- c("sd1-29_LPS_005","sd1-29_LPS_010","sd1-29_LPS_030","sd1-29_LPS_090","sd1-29_LPS_180",
                 "lyk4_5_ch8_005","lyk4_5_ch8_010","lyk4_5_ch8_030","lyk4_5_ch8_090","lyk4_5_ch8_180",
                 "efr-1_elf18_005","efr-1_elf18_010","efr-1_elf18_030","efr-1_elf18_090","efr-1_elf18_18
0",
                 "fls2c_flg22_005","fls2c_flg22_010","fls2c_flg22_030","fls2c_flg22_090","fls2c_flg22_18
0",
                 "rlp23-1_nlp20_005","rlp23-1_nlp20_010","rlp23-1_nlp20_030","rlp23-1_nlp20_090","rlp23-
1_nlp20_180",
                 "Col_mock_005","Col_mock_010","Col_mock_030","Col_mock_090","Col_mock_180",
                 "pepr1_2_Pep1_005","pepr1_2_Pep1_010","pepr1_2_Pep1_030","pepr1_2_Pep1_090","pepr1_2_Pe
p1_180")

#Run the function on each treatment/control together
log2FClist <- mapply(wt_sig_up, PAMPTime, mutPAMPTime,SIMPLIFY = F)
#combine into one data frame, and set log2FC to 0 for treatment/time combinations not significantly indu
ced
log2FCdf <- Reduce(function(x,y) merge(x, y, all=T), log2FClist)
log2FCdf[is.na(log2FCdf)] <- 0
names(log2FCdf) <- gsub("LPS", "3-OH-FA", names(log2FCdf))
names(log2FCdf) <- gsub("ch8", "CO8", names(log2FCdf))

nrow(log2FCdf)
```

```
## [1] 5876
```

Cut into five lists according to when first induced, get enriched GO terms in each

```
#First load up data and organize by time at which genes first induced
long2FCdf <- gather(log2FCdf, -ATG, key=sample, value=l2FC) %>% separate(sample, into=c("Geno","TRT","Ti
me"), sep="_")
PAMPup <- long2FCdf[long2FCdf$l2FC > 0,]
upfirst <- PAMPup %>% group_by(ATG) %>% summarise(FirstInduced = min(Time))



#Then calculate enriched GO terms. For comparison each set enrichment, use all genes detected in this st
udy
geneUniverse <- read.csv(paste0(home, "wtonly/Analysis-MMclustering/All_detected.csv"), header=F)
geneUniverse <- as.factor(as.vector(geneUniverse$V1))

MakeGOTable <- function (x,cid){
  clusters=unique(cid)
  K=length(clusters)
  for(i in 1:K){
    geneGalaxy <- x[cid==clusters[i]]
    if(length(geneGalaxy)!=1){
    geneList <- factor(as.integer(geneUniverse %in% geneGalaxy))
    names(geneList) <- geneUniverse
    GOdata <- new("topGOdata", ontology = "BP", allGenes = geneList, nodeSize = 10, annot = annFUN.org,
 mapping="org.At.tair.db")
    #so many options for different tests...
    resultTopGO.weight01 <- runTest(GOdata, algorithm = "weight01", statistic = "Fisher" )
    fill <- GenTable(GOdata, Uncorrected_p = resultTopGO.weight01, topNodes = 200)
    fill$Uncorrected_p[fill$Uncorrected_p=="< 1e-30"] <- 1e-30
    fill$Uncorrected_p <- as.numeric(fill$Uncorrected_p)
    write.csv(fill, file=paste0("GO_time",clusters[i], "up.csv"))
    }
  }
}

MakeGOTable(upfirst$ATG, upfirst$FirstInduced)
```

Could probably merge files in R, but easier since I’ve saved anyway to just pull in the external files, merging as I go

```
GetAndMerge <- function(big, new){
  new.df <- read.csv(paste0("GO_time", new,".csv"), header=T)
  new.df <- new.df[,c("Term", "Uncorrected_p")]
  #Optional add multiple testing correction here
  new.df <- new.df[new.df$Uncorrected_p<0.01,]
  bigger <- merge(big, new.df, by="Term", all=T)
  names(bigger) <- c(names(big), new)
  return(bigger)
}
GO <- read.csv("GO_time005up.csv", header=T)
GO <- GO[,c("Term", "Uncorrected_p")]
GO <- GO[GO$Uncorrected_p<0.01,]
names(GO) <- c("Term","005up")

GO <- GetAndMerge(GO, "010up")
GO <- GetAndMerge(GO, "030up")
GO <- GetAndMerge(GO, "090up")
GO <- GetAndMerge(GO, "180up")

write.csv(GO, file="GOup.csv")
```

Again, my environment is really crowded, just clear it out and bring in the file just made

```
GOsort <- GO %>% column_to_rownames("Term")
colnames(GOsort) <- gsub("up","min",colnames(GOsort))

#to make it graph-able replace all ""NA" (this GO term was not enriched) with 0.1 (The cutoff for being
 called enriched)
GOsort[is.na(GOsort)] <- 0.01

GOsortlog <- -log10(GOsort)
#arbritrary cutoff to get rid of GO terms only weakly enriched in one condition
GOsortlog <- GOsortlog %>% rownames_to_column("GOterm") %>% mutate(allp=rowSums(.[2:6])) %>% filter(allp
> 10.5)

#split again to arrange GO terms in each set according to behaviour across all sets, assign GO term enri
ched in more than one time to the time in which is most enriched
GO005 <- GOsortlog[max.col(GOsortlog[2:6], ties.method="first")==1,] %>% arrange(desc(allp)) %>% arrange
(desc(`005min`)) %>% column_to_rownames("GOterm")
GO010 <- GOsortlog[max.col(GOsortlog[2:6], ties.method="first")==2,] %>% arrange(desc(allp)) %>% arrange
(desc(`010min`)) %>% column_to_rownames("GOterm")
GO030 <- GOsortlog[max.col(GOsortlog[2:6], ties.method="first")==3,] %>% arrange(desc(allp)) %>% arrange
(desc(`030min`)) %>% column_to_rownames("GOterm")
GO090 <- GOsortlog[max.col(GOsortlog[2:6], ties.method="first")==4,] %>% arrange(desc(allp)) %>% arrange
(desc(`090min`)) %>% column_to_rownames("GOterm")
GO180 <- GOsortlog[max.col(GOsortlog[2:6], ties.method="first")==5,] %>% arrange(desc(allp)) %>% arrange
(desc(`180min`)) %>% column_to_rownames("GOterm")

#recombine the tables for each time, in order
GOarranged <- rbind(GO005, GO010, GO030, GO090, GO180)
#make nicer labels
colnames(GOarranged)[colnames(GOarranged)=="005min"] <- "5"
colnames(GOarranged)[colnames(GOarranged)=="010min"] <- "10"
colnames(GOarranged)[colnames(GOarranged)=="030min"] <- "30"
colnames(GOarranged)[colnames(GOarranged)=="090min"] <- "90"
colnames(GOarranged)[colnames(GOarranged)=="180min"] <- "180"

#pivot wider to plot with ggplot
ggGOarranged <- GOarranged[1:5] %>% rownames_to_column("GOterm") %>% pivot_longer(-"GOterm",names_to="ti
me",values_to = "p")
```

```
PanelA <- ggplot(ggGOarranged, aes(x=as.factor(as.numeric(time)), y=factor(GOterm, levels=rev(unique(GOt
erm))), fill=p)) +
  theme_minimal(base_size=7) + geom_tile()+scale_fill_viridis_c(limits=c(0,30), guide=guide_colorbar(bar
width=unit(1.5, "cm"), barheight=unit(0.2, "cm")))+
  theme(axis.text.y=element_blank(), axis.ticks.y=element_blank(), rect=element_rect(fill="transparent",
color=NA),
        panel.grid.major=element_blank(), legend.position="top", axis.text.x=element_text(angle=45, hjus
t=1, vjust=1),
        plot.margin=unit(c(0.25,0,0.25,-0.25), "cm"))+
  labs(x="time", y="", fill=expression("-log"[10]*"(p value)"))

#Just keep Panel A for a bit, but in the meantime save a separate plot that is the same but shows all th
e GO terms, will use it to fill in top3 at each time in main figure
GOguide <- ggplot(ggGOarranged, aes(x=as.factor(as.numeric(time)), y=factor(GOterm, levels=rev(unique(GO
term))), fill=p)) +
  theme_minimal(base_size=7) + geom_tile()+scale_fill_viridis_c(limits=c(0,30), guide=guide_colorbar(bar
width=unit(1.5, "cm"), barheight=unit(0.2, "cm")))+
  theme(axis.ticks.y=element_blank(), rect=element_rect(fill="transparent",color=NA), panel.grid.major=e
lement_blank(), legend.position="top", axis.text.x=element_text(angle=45, hjust=1, vjust=1))+
  labs(x="time", y="", fill=expression("-log"[10]*"(p value)"))
ggsave("GOguide.pdf", height=15, width=8)
```

### Panel B

To make a similar figure for cis element enrichment, need same data but didn't (can't?) do cis element enrichment in R. So save gene lists based on time induced

```
write.csv(upfirst[upfirst$FirstInduced=='005',], file="genes_005up.csv")
write.csv(upfirst[upfirst$FirstInduced=='010',], file="genes_010up.csv")
write.csv(upfirst[upfirst$FirstInduced=='030',], file="genes_030up.csv")
write.csv(upfirst[upfirst$FirstInduced=='090',], file="genes_090up.csv")
write.csv(upfirst[upfirst$FirstInduced=='180',], file="genes_180up.csv")
```

These .csv files were run through python scripts to get the 1000bp promoter of each (from TAIR database, GetPromotersFasta.py), then ame (http://meme-suite.org/doc/ame.html (http://meme-suite.org/doc/ame.html)) to get enriched cis elements for each set. The ame output was processed with a second python script: Extract_motif_p_fromAME.py, to get cis element enrichment (average p value) relative to 3 random sets of detected genes. Final output was one file per time

```
GetAndMerge <- function(big, new){
  new.df <- read.csv(paste0("D:/OneDrive/Norwich_Z/Overall/CleanedData/wtonly/Analysis-cis/Figure1_JIT_A
ME/", new,".csv"), header=T)
  new.df <- new.df[,c("X1", "X2")]
  names(new.df) <- c("TF","corrected_p")
  #Optional add multiple testing correction here
  new.df <- new.df[new.df$corrected_p<0.01,]
  bigger <- merge(big, new.df, by="TF", all=T)
  names(bigger) <- c(names(big), new)
  return(bigger)
}
up <- read.csv("D:/OneDrive/Norwich_Z/Overall/CleanedData/wtonly/Analysis-cis/Figure1_JIT_AME/up_005.cs
v", header=T)
up <- up[,c("X1", "X2")]
names(up) <- c("TF","corrected_p")
up <- up[up$corrected_p<0.01,]
names(up) <- c("TF","up_005")

up <- GetAndMerge(up, "up_010")
up <- GetAndMerge(up, "up_030")
up <- GetAndMerge(up, "up_090")
up <- GetAndMerge(up, "up_180")

up <- up %>% group_by(TF) %>% summarise(`005min`=mean(up_005), `010min`=mean(up_010),`030min`=mean(up_03
0),`090min`=mean(up_090),`180min`=mean(up_180))
```

```
## `summarise()` ungrouping output (override with `.groups` argument)
```

```
write.csv(up, file="cis_up.csv")
```

```
cistable <- read.csv("cis_up.csv")[2:7]
colnames(cistable) <- gsub("X","",colnames(cistable))
cistable[is.na(cistable)] <- 0.01
cistable <- column_to_rownames(cistable, "TF")
cistablelog <- -log10(cistable)
cistablelog <- cistablelog[rowSums(cistablelog)>10.5,]
cistablelog <- cistablelog %>% rownames_to_column("TF") %>% mutate(allp=rowSums(.[2:length(cistablelo
g)]))

cis005 <- cistablelog[max.col(cistablelog[2:length(cistable)], ties.method="first")==1,] %>% arrange(des
c(allp)) %>% arrange(desc(`005min`)) %>% column_to_rownames("TF")
cis010 <- cistablelog[max.col(cistablelog[2:length(cistable)], ties.method="first")==2,] %>% arrange(des
c(allp)) %>% arrange(desc(`010min`)) %>% column_to_rownames("TF")
cis030 <- cistablelog[max.col(cistablelog[2:length(cistable)], ties.method="first")==3,] %>% arrange(des
c(allp)) %>% arrange(desc(`030min`)) %>% column_to_rownames("TF")
cis090 <- cistablelog[max.col(cistablelog[2:length(cistable)], ties.method="first")==4,] %>% arrange(des
c(allp)) %>% arrange(desc(`090min`)) %>% column_to_rownames("TF")
cis180 <- cistablelog[max.col(cistablelog[2:length(cistable)], ties.method="first")==5,] %>% arrange(des
c(allp)) %>% arrange(desc(`180min`)) %>% column_to_rownames("TF")

cisarranged <- rbind(cis005, cis010, cis030, cis090, cis180)
colnames(cisarranged) <- c("5","10","30","90","180","allp")

ggcisarranged <- cisarranged[1:5] %>% rownames_to_column("cis") %>% pivot_longer(-"cis",names_to="time",
values_to = "p")
```

```
PanelB <- ggplot(ggcisarranged, aes(x=as.factor(as.numeric(time)), y=factor(cis, levels=rev(unique(ci
s))), fill=p)) +
  theme_minimal(base_size=7) + geom_tile()+scale_fill_viridis_c(limits=c(0,30), guide=guide_colorbar(bar
width=unit(1.5, "cm"), barheight=unit(0.2, "cm")))+
  theme(rect=element_rect(fill="transparent",color=NA), panel.grid.major=element_blank(),
        legend.position="top", axis.text.x=element_text(angle=45, hjust=1, vjust=1) , plot.margin=unit(c
(0.25,-0.25,0.25,0), "cm"))+
  scale_y_discrete(position="right")+
  labs(x="time (min)", y="", fill=expression("-log"[10]*"(p value)"))
```

```
spacer <- ggplot(ggcisarranged, aes(x=time, y=cis))+
  theme_nothing()+
  theme(rect=element_rect(fill="transparent",color=NA), axis.text=element_blank(), plot.margin=unit(c(0.
25,0,0.25,0), "cm"))

PanelAB <- ggarrange(PanelA, spacer, PanelB, widths = c(1.25,1.75,2), ncol=3, common.legend=T)
ggsave(PanelAB, filename="PanelAB.pdf",width=5, height=9, units="cm", bg="transparent")
```

### Panel C

Genes induced by number abio and number elicitors, by time

```
rm(list=ls())
home <- "D:/OneDrive/Norwich_Z/Overall/CleanedData/wtonly/"
#load up the new spreadsheet (as of 2020-04ish) with abio stress done by probe
FullInfo <- read.csv(paste0(home,"AllGenesInfo.csv"), header=T)
#remove genes not induced by elicitors, or where abiotic stress information isn't clean
TimeAbio <- FullInfo[!is.na(FullInfo$NumberElicitors),]
TimeAbio <- TimeAbio[TimeAbio$AtGenExpressNumber != "NotOnArray" & TimeAbio$AtGenExpressNumber != "Probe
HitsMultipleGenes",]
TimeAbio$NumberElicitors <- as.factor(TimeAbio$NumberElicitors)
```

Timebox with induction

```
Elicnumber <- FullInfo[!is.na(FullInfo$NumberElicitors),]
Elicnumber$NumberElicitors <- as.factor(Elicnumber$NumberElicitors)

TimeAbio <- Elicnumber[Elicnumber$AtGenExpressNumber != "NotOnArray" & Elicnumber$AtGenExpressNumber !=
"ProbeHitsMultipleGenes",]

TimeAbio <- TimeAbio[,2:length(TimeAbio)]
maxinduce <- TimeAbio %>% gather(-ATG, -AtGenExpressNumber, -NumberElicitors, -FirstInduced, key=conditi
on, value=L2FC) %>% group_by(ATG, AtGenExpressNumber, NumberElicitors, FirstInduced) %>% summarize(great
est=max(L2FC))
```

```
## `summarise()` regrouping output by 'ATG', 'AtGenExpressNumber', 'NumberElicitors' (override with `.gr
oups` argument)
```

```
PanelC <- ggplot(maxinduce, aes(x=NumberElicitors , y=AtGenExpressNumber)) +
  geom_jitter(height=0.25, width=0.25, size=0.5, aes(color=greatest, fill=greatest), shape=21,stroke=0.1
2) +
  theme_minimal(base_size=8) +
  theme(panel.background = element_blank(), plot.background = element_blank(), legend.background = eleme
nt_blank(), legend.box.background = element_blank(), panel.border = element_blank(), legend.box=NULL, le
gend.position = "top", strip.text.y=element_text(angle=0), legend.title=element_text(size=7), legend.tex
t=element_text(size=6)) +
  labs(x="Number patterns inducing", y="Number abiotic stresses inducing", color=bquote('Max '*~log[2]*
'(FC)')) +
  facet_grid(FirstInduced~.) +
  scale_color_viridis_c(option="inferno", direction=-1, limits=c(0, 9), breaks=c(0,3,6,9), guide=guide_c
olorbar(barwidth=unit(1.5, "cm"), barheight=unit(0.1, "cm")))+
  scale_fill_viridis_c(option="inferno", direction=-1, limits=c(0, 9), breaks=c(0,3,6,9), alpha=0.3, gui
de=F)

ggsave("PanelC.pdf", PanelC, height=13, width=5, units="cm", bg="transparent")
```

### Panel D

Making IR graphs to show immunity affected First gather and display raw data

```
#Would like to update camta3 result with (less impressive) results I'm getting at UZH. Pick my favorite
 days with similar levels of infection and IR in Col-0
rm(list=ls())
home <- "D:/OneDrive/Norwich_U/DATA- IR assays/"

A <- read_excel(paste0(home,"2018-11-13test1.xlsx"))
B <- read_excel(paste0(home,"2019-02-22.xlsx"))
C <- read_excel(paste0(home,"2019-05-07.xlsx"))

A$Exp <- "A"
B$Exp <- "B"
C$Exp <- "C"

cdd <- rbind(A,B,C)

#2019-05-07 used two chamber and two seed stocks, just keep one
cdd$Geno[cdd$Geno=="17_camta3/dsc1/dsc2.here"] <- "camta3/dsc1/dsc2"
cdd$Geno[cdd$Geno=="17_Col-0"] <- "Col-0"
cdd$Geno[cdd$Geno=="Col"] <- "Col-0"
cdd$TRT[cdd$TRT=="flg"] <- "flg22"

cdd <- cdd[cdd$Geno=="Col-0" | cdd$Geno=="camta3/dsc1/dsc2",]
cdd$Geno <- factor(cdd$Geno, levels=c("Col-0", "camta3/dsc1/dsc2"))
cdd$TRT <- factor(cdd$TRT, levels=c("mock","flg22"))
mylabels <- c("Col-0", expression(italic("camta3/dsc1/dsc2")))

print(cdd[,c(1, 3:6, 8:9, 12:13)], n=Inf, width=Inf)
```

```
## # A tibble: 52 x 9
##    Exp   Geno             TRT   dilution count `drop volume` discs `CFU/cm2`
##    <chr> <fct>            <fct>    <dbl> <dbl>         <dbl> <dbl>     <dbl>
##  1 A     Col-0            mock         4    62            15     6 10964007.
##  2 A     Col-0            mock         3    40            15     6   707355.
##  3 A     Col-0            mock         3    32            15     6   565884.
##  4 A     Col-0            mock         3    80            15     6  1414711.
##  5 A     Col-0            mock         3    13            15     6   229890.
##  6 A     Col-0            flg22        3    15            15     6   265258.
##  7 A     Col-0            flg22        1    63            15     3    22282.
##  8 A     Col-0            flg22        2    13            15     3    45978.
##  9 A     Col-0            flg22        1    80            15     3    28294.
## 10 A     Col-0            flg22        1    14            15     3     4951.
## 11 A     camta3/dsc1/dsc2 mock         1    26            15     3     9196.
## 12 A     camta3/dsc1/dsc2 mock         1    61            15     6    10787.
## 13 A     camta3/dsc1/dsc2 mock         2    74            15     6   130861.
## 14 A     camta3/dsc1/dsc2 mock         3    31            15     6   548200.
## 15 A     camta3/dsc1/dsc2 mock         2    39            15     6    68967.
## 16 A     camta3/dsc1/dsc2 flg22        1   117            15     2    62070.
## 17 A     camta3/dsc1/dsc2 flg22        2    10            15     3    35368.
## 18 A     camta3/dsc1/dsc2 flg22        2    10            15     3    35368.
## 19 A     camta3/dsc1/dsc2 flg22        1    37            15     6     6543.
## 20 A     camta3/dsc1/dsc2 flg22        1    46            15     6     8135.
## 21 B     Col-0            mock         3    97            20     6  1286502.
## 22 B     Col-0            mock         3    53            20     6   702934.
## 23 B     Col-0            mock         3    31            20     6   411150.
## 24 B     Col-0            mock         3    19            20     6   251995.
## 25 B     Col-0            flg22        1    36            20     6     4775.
## 26 B     Col-0            flg22        2    18            20     6    23873.
## 27 B     Col-0            flg22        2    11            20     6    14589.
## 28 B     Col-0            flg22        1    70            20     6     9284.
## 29 B     camta3/dsc1/dsc2 mock         3    84            20     3  2228169.
## 30 B     camta3/dsc1/dsc2 mock         2    39            20     3   103451.
## 31 B     camta3/dsc1/dsc2 mock         1    34            20     3     9019.
## 32 B     camta3/dsc1/dsc2 mock         1    14            20     3     3714.
## 33 B     camta3/dsc1/dsc2 flg22        1    81            20     2    32229.
## 34 B     camta3/dsc1/dsc2 flg22        2    52            20     3   137934.
## 35 B     camta3/dsc1/dsc2 flg22        3    56            20     3  1485446.
## 36 B     camta3/dsc1/dsc2 flg22        1    25            20     3     6631.
## 37 C     camta3/dsc1/dsc2 mock         3    10            20     6   132629.
## 38 C     Col-0            mock         3    40            20     6   530516.
## 39 C     camta3/dsc1/dsc2 flg22        2    77            20     6   102124.
## 40 C     Col-0            flg22        2    27            20     6    35810.
## 41 C     camta3/dsc1/dsc2 mock         3    24            20     6   318310.
## 42 C     Col-0            mock         3    38            20     6   503991.
## 43 C     camta3/dsc1/dsc2 flg22        2    43            20     6    57031.
## 44 C     Col-0            flg22        1   125            20     6    16579.
## 45 C     camta3/dsc1/dsc2 mock         2    48            20     6    63662.
## 46 C     Col-0            mock         2   118            20     6   156502.
## 47 C     camta3/dsc1/dsc2 flg22        2    23            20     6    30505.
## 48 C     Col-0            flg22        1    25            20     6     3316.
## 49 C     camta3/dsc1/dsc2 mock         3    14            20     4   278521.
## 50 C     Col-0            mock         3    34            20     4   676409.
## 51 C     camta3/dsc1/dsc2 flg22        1    26            20     4     5173.
## 52 C     Col-0            flg22        1     5            20     4      995.
##      log
##    <dbl>
##  1  7.04
##  2  5.85
##  3  5.75
##  4  6.15
##  5  5.36
##  6  5.42
```

```
##  7  4.35
##  8  4.66
##  9  4.45
## 10  3.69
## 11  3.96
## 12  4.03
## 13  5.12
## 14  5.74
## 15  4.84
## 16  4.79
## 17  4.55
## 18  4.55
## 19  3.82
## 20  3.91
## 21  6.11
## 22  5.85
## 23  5.61
## 24  5.40
## 25  3.68
## 26  4.38
## 27  4.16
## 28  3.97
## 29  6.35
## 30  5.01
## 31  3.96
## 32  3.57
## 33  4.51
## 34  5.14
## 35  6.17
## 36  3.82
## 37  5.12
## 38  5.72
## 39  5.01
## 40  4.55
## 41  5.50
## 42  5.70
## 43  4.76
## 44  4.22
## 45  4.80
## 46  5.19
## 47  4.48
## 48  3.52
## 49  5.44
## 50  5.83
## 51  3.71
## 52  3.00
```

Then make the plot

```
PanelD <- ggplot(cdd, aes(x=Geno , y=log)) +
  geom_boxplot(aes(group=interaction(Geno, TRT)), position=position_dodge(width=0.9), width=0.85, outlie
r.shape=NA, size=0.3) +
  geom_point(position=position_jitterdodge(dodge.width = 0.9, jitter.width=0.5), aes(fill=TRT, group=int
eraction(Geno, TRT), shape=Exp), size=0.8, alpha=0.7, stroke=0.3)+
  theme_minimal(base_size=7) +
  theme(panel.background = element_blank(), plot.background = element_blank(), legend.background = eleme
nt_blank(), legend.box.background = element_blank(), panel.border = element_blank(), legend.box=NULL, le
gend.key.size=unit(0.15, "cm"), legend.position="top", legend.title=element_blank()) +
  labs(x="", y=expression("log"[10]*"(CFU\U2022"*cm^-2*')'), fill="") +
  scale_fill_manual(values=c("black", "grey")) +
  scale_x_discrete(labels=mylabels)+
  scale_shape_manual(values=c(21, 22, 23, 24), guide=F)

ggsave("PanelD.pdf", PanelD, height=4, width=5, units="cm", bg="transparent")
```

and finally the stats

```
ir.lm <- lm(log~Geno*TRT+Exp, data=cdd)
plot(ir.lm, 1:2)
```

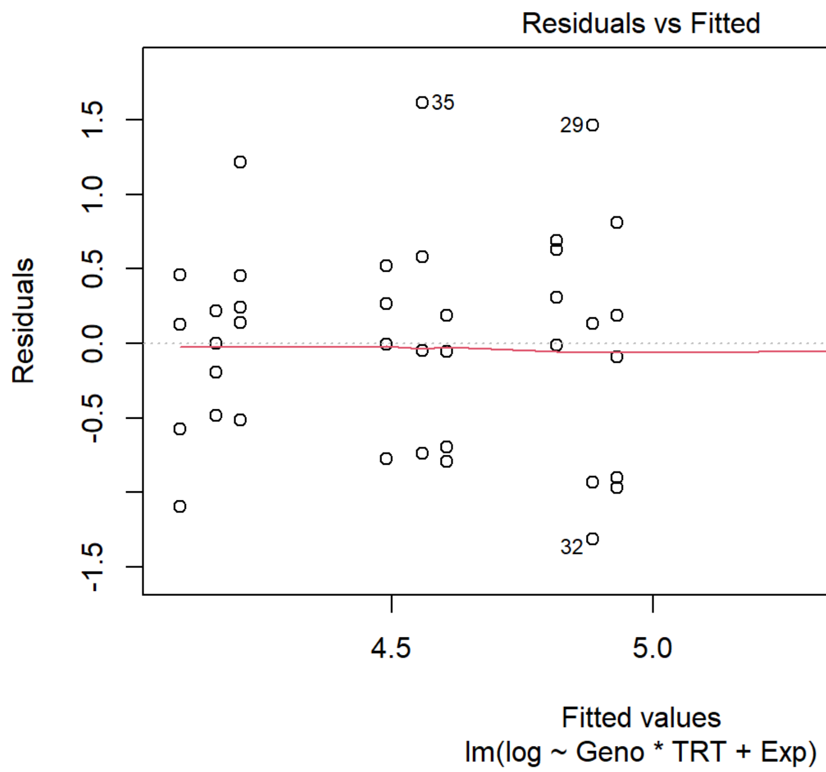
Residuals vs Fitted
Residuals
Fitted values
lm(log ~ Geno * TRT + Exp)
35
29
32
1.5
1.0
0.5
0.0
-0.5
-1.5
4.5
5.0
5.5

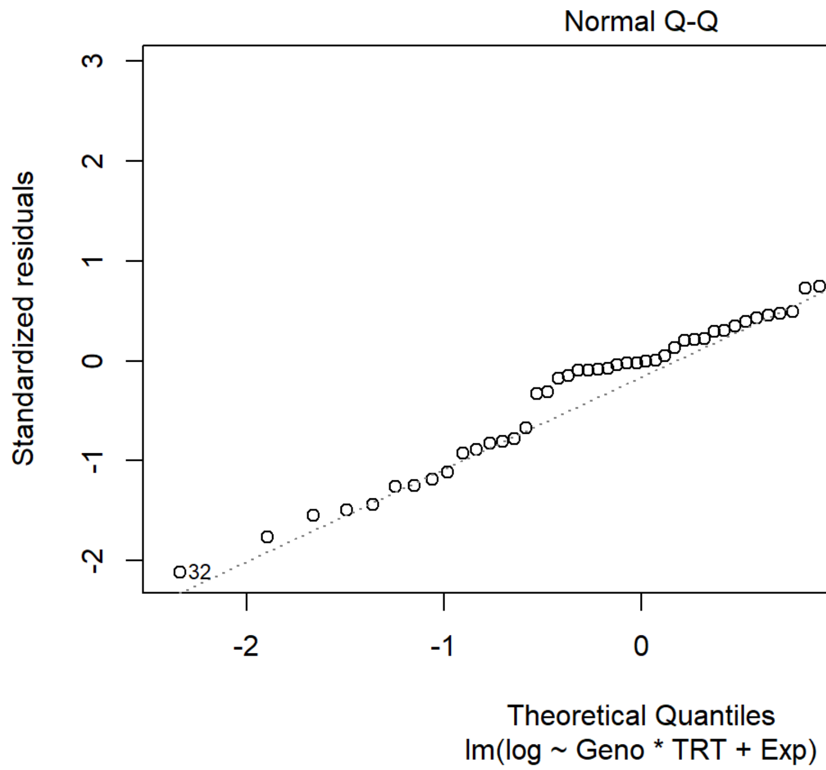
Normal Q-Q
Standardized residuals
Theoretical Quantiles
lm(log ~ Geno * TRT + Exp)
35
29
32
3
2
1
0
-1
-2
-2
-1
0
1
2

```
anova(ir.lm)
```

```
## Analysis of Variance Table
##
## Response: log
##           Df  Sum Sq Mean Sq F value    Pr(>F)
## Geno       1  0.9330  0.9330  2.1275 0.1514754
## TRT        1 12.7501 12.7501 29.0718 2.335e-06 ***
## Exp        2  0.1194  0.0597  0.1362 0.8730503
## Geno:TRT   1  5.7457  5.7457 13.1010 0.0007321 ***
## Residuals 46 20.1743  0.4386
## ---
## Signif. codes:  0 '***' 0.001 '**' 0.01 '*' 0.05 '.' 0.1 ' ' 1
```

```
#get effect of genotype in each treatment
twostep <- emmeans(ir.lm, ~ Geno | TRT)

  twostep.3 <- contrast(twostep, "dunnett")
  twostep.3
```

```
## TRT = mock:
##  contrast                     estimate   SE df t.ratio p.value
##  (camta3/dsc1/dsc2) - (Col-0)   -0.933 0.26 46 -3.591  0.0008
##
## TRT = flg22:
##  contrast                     estimate   SE df t.ratio p.value
##  (camta3/dsc1/dsc2) - (Col-0)    0.397 0.26 46  1.528  0.1334
##
## Results are averaged over the levels of: Exp
```

```
  #is effect of treatment the same in each genotype
  TRTeffect <-contrast(twostep, interaction = "pairwise")#interaction selected doesn't matter here, only two levels for each
  Int.effect <- contrast(TRTeffect, "dunnett", by = NULL)
  summary(Int.effect)
```

```
##  contrast
##  ((Col-0) - (camta3/dsc1/dsc2) flg22) - ((Col-0) - (camta3/dsc1/dsc2) mock)
##  estimate    SE df t.ratio p.value
##     -1.33 0.367 46 -3.620  0.0007
##
## Results are averaged over the levels of: Exp
```

```
#get effect of treatment in each genotype
twostep <- emmeans(ir.lm, ~ TRT | Geno)
twostep.3 <- contrast(twostep, "dunnett")
twostep.3
```

```
## Geno = Col-0:
##  contrast     estimate   SE df t.ratio p.value
##  flg22 - mock   -1.655 0.26 46 -6.372  <.0001
##
## Geno = camta3/dsc1/dsc2:
##  contrast     estimate   SE df t.ratio p.value
##  flg22 - mock   -0.326 0.26 46 -1.253  0.2165
##
## Results are averaged over the levels of: Exp
```

### Figure 3

Marta Bjornson

```
knitr::opts_chunk$set(echo = TRUE)
library(tidyverse)
library(readxl)
library(emmeans)
library(cowplot)
```

### Panel A

Panel A is peak (R0-R)/R0. Same data time course in extended data figure 6. First define functions for reading in data

```
rm(list=ls())
home <- "D:/OneDrive/Norwich_U/Data-Ca/"
muted <- c("black", "#BBBBBB", "#117733", "#44AA99", "#999933", "#DDCC77",  "#332288", "#88CCEE", "#AA44
99", "#CC6677")
smallmuted <- c("black", "#BBBBBB", "#AA4499", "#CC6677")
#These two functions read in treatment data, from an excel file. Set up the excel file in the orientatio
n in which the plate(s) went into the camera EXCEPT if there are two plates, stack them vertically not h
orizontally. This is critical! There is no error-checking in the code to ensure that a well lines up wit
h the appropriate region of interest.
makelist <- function(filename, sheet){
  worksheet <- sheet
  variable <- read_excel(filename, sheet = worksheet)
  variablelist <- variable[1,]
  for(i in 2:nrow(variable)){
    variablelist <- c(variablelist, variable[i,])
  }
  variablevector <- unlist(variablelist)
  return(variablevector)
}

add_treatments <- function(filename, inheritedros){
  looper <- excel_sheets(filename)
  for(i in 1:length(looper)){
    column <- looper[i]
    columnlist <- makelist(filename, column)
    inheritedros$new <- unlist(columnlist)
    colnames(inheritedros)[colnames(inheritedros)=="new"] <- column
  }
  return(inheritedros)
}

ReadTecan <- function(datafile, planfile, treatment, conc, exp){
  base_cfp1 <- read_excel(datafile, range="B82:CU92") %>% mutate_if(is.character, funs(recode(.,"OVER" =
0))) %>%
    pivot_longer(-c("Time [s]", "Temp. [°C]"), names_to="Well", values_to="CFP") %>% na.omit() %>% selec
t(-"Temp. [°C]")
  base_yfp1 <- read_excel(datafile, range="B95:CU105")%>% mutate_if(is.character, funs(recode(.,"OVER" =
0))) %>%
    pivot_longer(-c("Time [s]", "Temp. [°C]"), names_to="Well", values_to="YFP") %>% na.omit() %>% selec
t(-"Temp. [°C]")
  base_cfp1$`Time [s]` <- base_cfp1$`Time [s]` - 300
  base_yfp1$`Time [s]` <- base_yfp1$`Time [s]` - 300

  cfp1 <- read_excel(datafile, range="B161:CU211") %>% mutate_if(is.character, funs(recode(.,"OVER" = 0
))) %>%
    pivot_longer(-c("Time [s]", "Temp. [°C]"), names_to="Well", values_to="CFP") %>% na.omit() %>% selec
t(-"Temp. [°C]")
  yfp1 <- read_excel(datafile, range="B214:CU264") %>% mutate_if(is.character, funs(recode(.,"OVER" = 0
))) %>%
    pivot_longer(-c("Time [s]", "Temp. [°C]"), names_to="Well", values_to="YFP") %>% na.omit() %>% selec
t(-"Temp. [°C]")

  #combine into one data frame
  yfp <- rbind(base_yfp1, yfp1)
  cfp <- rbind(base_cfp1, cfp1)

  #get ratios
  fluor <- merge(cfp, yfp, by=c("Time [s]", "Well"))
  fluor$`Time [s]` <- round(fluor$`Time [s]`)
  fluor$`Time (min)` <- round(fluor$`Time [s]`/60, 1)
  fluor$ratio <- fluor$YFP/fluor$CFP

  fill <- data.frame(filler=rep(0,96))
```

```
  trt <- add_treatments(planfile,fill)

  fluor <- merge(fluor, trt[,2:ncol(trt)], by="Well")
  fluor$Row <- substring(fluor$Well, 1, 1)
  fluor$Column <- substring(fluor$Well, 2, nchar(as.character(fluor$Well))) %>% as.numeric()

  fluor$Trt <- paste0(conc,treatment)
  fluor$Date <- exp
  return(fluor)
}

ReadSalt <- function(datafile, planfile, treatment, conc, exp){
  base_cfp1 <- read_excel(datafile, range="B83:CU93") %>% mutate_if(is.character, funs(recode(.,"OVER" =
0))) %>%
    pivot_longer(-c("Time [s]", "Temp. [°C]"), names_to="Well", values_to="CFP") %>% na.omit() %>% selec
t(-"Temp. [°C]")
  base_yfp1 <- read_excel(datafile, range="B96:CU106")%>% mutate_if(is.character, funs(recode(.,"OVER" =
0))) %>%
    pivot_longer(-c("Time [s]", "Temp. [°C]"), names_to="Well", values_to="YFP") %>% na.omit() %>% selec
t(-"Temp. [°C]")
  base_cfp1$`Time [s]` <- base_cfp1$`Time [s]` - 300
  base_yfp1$`Time [s]` <- base_yfp1$`Time [s]` - 300

  cfp1 <- read_excel(datafile, range="B161:CU174") %>% mutate_if(is.character, funs(recode(.,"OVER" = 0
))) %>%
    pivot_longer(-c("Time [s]", "Temp. [°C]"), names_to="Well", values_to="CFP") %>% na.omit() %>% selec
t(-"Temp. [°C]")
  yfp1 <- read_excel(datafile, range="B177:CU190") %>% mutate_if(is.character, funs(recode(.,"OVER" = 0
))) %>%
    pivot_longer(-c("Time [s]", "Temp. [°C]"), names_to="Well", values_to="YFP") %>% na.omit() %>% selec
t(-"Temp. [°C]")

  #combine into one data frame
  yfp <- rbind(base_yfp1, yfp1)
  cfp <- rbind(base_cfp1, cfp1)

  #get ratios
  fluor <- merge(cfp, yfp, by=c("Time [s]", "Well"))
  fluor$`Time [s]` <- round(fluor$`Time [s]`)
  fluor$`Time (min)` <- round(fluor$`Time [s]`/60, 1)
  fluor$ratio <- fluor$YFP/fluor$CFP

  fill <- data.frame(filler=rep(0,96))
  trt <- add_treatments(planfile,fill)

  fluor <- merge(fluor, trt[,2:ncol(trt)], by="Well")
  fluor$Row <- substring(fluor$Well, 1, 1)
  fluor$Column <- substring(fluor$Well, 2, nchar(as.character(fluor$Well))) %>% as.numeric()

  fluor$Trt <- paste0(conc,treatment)
  fluor$Date <- exp
  return(fluor)
}
```

Then get all the runs. For each date had a few wells with bad data - large jump in ratio after injection, or fluorescence values with sudden spike in the middle of reading. Found these through looking through each day independently, and removed here.

```
Aflg <- ReadTecan(paste0(home,"2020-09-01_pn4b_1uMflg22.xlsx"), paste0(home,"2020-09-01_pn3.xlsx"), "flg
22", "1 \U00B5M ", "A")
```

```
## Warning: `funs()` is deprecated as of dplyr 0.8.0.
## Please use a list of either functions or lambdas:
##
##   # Simple named list:
##   list(mean = mean, median = median)
##
##   # Auto named with `tibble::lst()`:
##   tibble::lst(mean, median)
##
##   # Using lambdas
##   list(~ mean(., trim = .2), ~ median(., na.rm = TRUE))
## This warning is displayed once every 8 hours.
## Call `lifecycle::last_warnings()` to see where this warning was generated.
```

```
#Remove wells: check individual day for each one
Aflg <- Aflg %>% filter(!Well %in% c("H7", "A8"))

Bflg <- ReadTecan(paste0(home,"2020-09-08_p2a_1uMflg22.xlsx"), paste0(home,"2020-09-08_plan.xlsx"), "flg
22", "1 \U00B5M ", "B")
Bflg <- Bflg %>% filter(!Well %in% c("C4","D5","A3"))

Cflg <- ReadTecan(paste0(home,"2020-09-15_p1b_1uMflg22.xlsx"),paste0(home,"2020-09-15_plan.xlsx"), "flg2
2","1 \U00B5M ", "C")
Cflg <- Cflg %>% filter(!Well %in% c("E7","F8"))


Amock <- ReadTecan(paste0(home,"2020-09-01_pn3b_mock.xlsx"), paste0(home,"2020-09-01_pn3.xlsx"), "Mock",
"", "A")
#Remove wells: check individual day for each one
Amock <- Amock %>% filter(!Well %in% c())

Bmock <- ReadTecan(paste0(home,"2020-09-08_p1a_mock.xlsx"), paste0(home,"2020-09-08_plan.xlsx"), "Mock",
"", "B")
Bmock <- Bmock %>% filter(!Well  %in%  c("B4"))

Cmock <- ReadTecan(paste0(home,"2020-09-15_p1a_mock.xlsx"),paste0(home,"2020-09-15_plan.xlsx"),"Mock",
"", "C")
Cmock <- Cmock %>% filter(!Well %in% c("A6","C4","G2"))

Aelf <- ReadTecan(paste0(home,"2020-09-01_po4b_1uMelf18.xlsx"), paste0(home,"2020-09-01_pn3.xlsx"), "elf
18", "1 \U00B5M ", "A")
#Remove wells: check individual day for each one
Aelf <- Aelf %>% filter(!Well %in% c("A9"))

Belf <- ReadTecan(paste0(home,"2020-09-08_p2b_1uMelf18.xlsx"), paste0(home,"2020-09-08_plan.xlsx"), "elf
18", "1 \U00B5M ", "B")
```

```
## Warning: Problem with `mutate()` input `F11`.
## i Unreplaced values treated as NA as .x is not compatible. Please specify replacements exhaustively o
r supply .default
## i Input `F11` is `recode(F11, OVER = 0)`.
```

```
## Warning: Unreplaced values treated as NA as .x is not compatible. Please specify
## replacements exhaustively or supply .default
```

```
## Warning: Problem with `mutate()` input `F11`.
## i Unreplaced values treated as NA as .x is not compatible. Please specify replacements exhaustively o
r supply .default
## i Input `F11` is `recode(F11, OVER = 0)`.
```

```
## Warning: Unreplaced values treated as NA as .x is not compatible. Please specify
## replacements exhaustively or supply .default
```

```
Belf <- Belf %>% filter(!Well %in% c("C10","F11","B8","F7","G10"))

Celf <- ReadTecan(paste0(home,"2020-09-15_p4a_1uMelf18.xlsx"),paste0(home,"2020-09-15_plan.xlsx"), "elf1
8","1 \U00B5M ", "C")
```

```
## Warning: Problem with `mutate()` input `G2`.
## i Unreplaced values treated as NA as .x is not compatible. Please specify replacements exhaustively o
r supply .default
## i Input `G2` is `recode(G2, OVER = 0)`.

## Warning: Unreplaced values treated as NA as .x is not compatible. Please specify replacements exhaust
ively or supply .default
```

```
## Warning: Problem with `mutate()` input `G2`.
## i Unreplaced values treated as NA as .x is not compatible. Please specify replacements exhaustively o
r supply .default
## i Input `G2` is `recode(G2, OVER = 0)`.
```

```
## Warning: Unreplaced values treated as NA as .x is not compatible. Please specify
## replacements exhaustively or supply .default
```

```
Celf <- Celf %>% filter(!Well %in% c("G2","A3","C5","F5"))

Apep <- ReadTecan(paste0(home,"2020-09-01_po4a_1uMpep1.xlsx"), paste0(home,"2020-09-01_pn3.xlsx"), "Pep
1", "1 \U00B5M ", "A")
#Remove wells: check individual day for each one
Apep <- Apep %>% filter(!Well %in% c("A8"))

Bpep <- ReadTecan(paste0(home,"2020-09-08_p1b_1uMPep1.xlsx"), paste0(home,"2020-09-08_plan.xlsx"), "Pep
1", "1 \U00B5M ", "B")
Bpep <- Bpep %>% filter(!Well %in% c("A10", "A7", "G7", "G9"))

Cpep <- ReadTecan(paste0(home,"2020-09-15_p2a_1uMPep1_MovedByElectrician.xlsx"),paste0(home,"2020-09-15_
plan.xlsx"), "Pep1","1 \U00B5M ", "C")
Cpep <- Cpep %>% filter(!Well %in% c("B9","D10"))
Cpep$Column <- Cpep$Column+6
```

Bind all data together, then find and remove wells (by date) where baseYFP is not above 3x level of non-fluorescent Col-0 control, to deal with some silencing. Next normalize ratios to first value post-injection, combine genotypes (tested seedlings from 2-3 different parent plants, all parents homozygous) and clean up names.

```
fluor <- rbind(Amock, Bmock, Cmock, Aflg,  Bflg, Cflg, Aelf, Belf, Celf, Apep, Bpep, Cpep)
Conc <- "1 \U00B5M "
Trt <- "flg etc"

#for now getting one Col-0 fluorescence value across all exp. In theory (and from seeing many of these)
 it shouldn't change much
Col_YFP <- fluor %>% filter(`Time (min)`<0) %>% filter(Genotype=="Col-0") %>% group_by(Well) %>% summari
ze(wellmean = mean(YFP)) %>% ungroup() %>% summarize(mean=mean(wellmean))
```

```
## `summarise()` ungrouping output (override with `.groups` argument)
```

```
Col_CFP <- fluor %>% filter(`Time (min)`<0) %>% filter(Genotype=="Col-0") %>% group_by(Well) %>% summari
ze(wellmean = mean(CFP)) %>% ungroup() %>% summarize(mean=mean(wellmean))
```

```
## `summarise()` ungrouping output (override with `.groups` argument)
```

```
#select only if THREE times Col-0 mean. (previously 5x, don't think I need to be that stringent)
fluor_judge <- fluor %>% filter(`Time (min)`<0)  %>% group_by(Date, Well, Trt, Genotype) %>% summarize(meanYFP=mean(YFP), meanCFP=mean(CFP)) %>% mutate(YFP_Judgement = if_else(meanYFP > Col_YFP*3, "Yes", "No")) %>% mutate(CFP_Judgement = if_else(meanCFP > Col_CFP*3,"Yes", "No")) %>% mutate(Judgement = if_else(YFP_Judgement =="Yes" & CFP_Judgement=="Yes", "Yes", "No"))
```

```
## `summarise()` regrouping output by 'Date', 'Well', 'Trt' (override with `.groups` argument)
```

```
fluor <-  merge(fluor, fluor_judge[,c("Date", "Well", "Judgement", "Trt")], by=c("Date", "Well", "Trt"))

time0norm <- function(time, ratio){
  ratio.df <- data.frame(`Time (min)`=time, "ratio"=ratio)
  colnames(ratio.df) <- c("Time (min)","ratio")
  pretreat <- ratio.df$ratio[ratio.df$`Time (min)` == 0]
  ratio.df$difference <- (ratio.df$ratio-pretreat)/pretreat
  return(ratio.df$difference)
}

norm_0 <- fluor %>% group_by(Date, Well, Trt, Genotype,Judgement) %>% arrange(`Time (min)`) %>% mutate(difference=time0norm(`Time (min)`, ratio))

filtered <- norm_0[norm_0$Judgement=="Yes",]

filtered <- filtered %>% ungroup() %>% add_column(Geno = "filler") %>% mutate(Geno = if_else(Genotype == "YC3.6 A" | Genotype == "YC3.6 B" | Genotype == "YC3.6 C", "YC3.6", Genotype)) %>% mutate(Geno = if_else(Genotype =="C2-2C" | Genotype == "C2-2D" , "glr2.7/2.8/2.9", Geno))

filtered$Geno <- factor(filtered$Geno, levels=c("YC3.6","glr2.7/2.8/2.9"))
filtered$Trt <- factor(filtered$Trt, levels=c("Mock","1 µM flg22","1 µM elf18","1 µM Pep1"))
```

Plot peaks

```
peak <- filtered %>% filter(`Time (min)` > 0) %>% group_by(Date, Geno, Trt, Well) %>% summarize(peak=max(difference)) %>% ungroup()
```

```
## `summarise()` regrouping output by 'Date', 'Geno', 'Trt' (override with `.groups` argument)
```

```
#unicode for mu
smallabels <- c("Mock","1 \U00B5M flg22","1 \U00B5M elf18","1 \U00B5M Pep1")

PanelA <- ggplot(peak, aes(x=Trt, y=peak, color=interaction(Geno, Trt)))+
  geom_boxplot(outlier.color=NA, size=0.3)+
  geom_point(position=position_jitterdodge(jitter.width=0.6), aes(fill=interaction(Geno, Trt), shape=Date, group=interaction(Geno, Trt)), size=1, alpha=0.8) +
  labs(x="",y=expression("Peak (R-R"[0]*")/R"[0]), title="", fill="", color="") +
  scale_x_discrete(labels=smallabels)+
  scale_fill_manual(values=muted)+
  scale_color_manual(values=muted)+scale_shape_manual(values=c(21,22,23,24), guide="none")+
  theme_cowplot()+
  theme(legend.position = "none", panel.background = element_blank(), plot.background = element_blank(),
        panel.border = element_blank(),
        axis.line=element_line(size=0.5), axis.ticks=element_line(size=0.2),
        axis.text = element_text(size=6), axis.title = element_text(size=7),
        plot.margin=unit(c(-0.6, 0, -0.35, 0), "cm"))

ggsave(PanelA, filename = "PanelA.pdf", height=4, width=7.5, units="cm",bg="transparent")
```

And statistics: test effect of genotype within each treatment

```
peak %>% group_by(Geno, Trt) %>% count()
```

```
## # A tibble: 8 x 3
## # Groups:   Geno, Trt [8]
##   Geno           Trt            n
##   <fct>          <fct>      <int>
## 1 YC3.6          Mock          56
## 2 YC3.6          1 µM flg22    54
## 3 YC3.6          1 µM elf18    52
## 4 YC3.6          1 µM Pep1     55
## 5 glr2.7/2.8/2.9 Mock          48
## 6 glr2.7/2.8/2.9 1 µM flg22    43
## 7 glr2.7/2.8/2.9 1 µM elf18    36
## 8 glr2.7/2.8/2.9 1 µM Pep1     43
```

```
peak.lm <- lm(sqrt(peak) ~ Geno*Trt+Date, data=peak)
```

```
## Warning in sqrt(peak): NaNs produced
```

```
plot(peak.lm, 1:2)
```

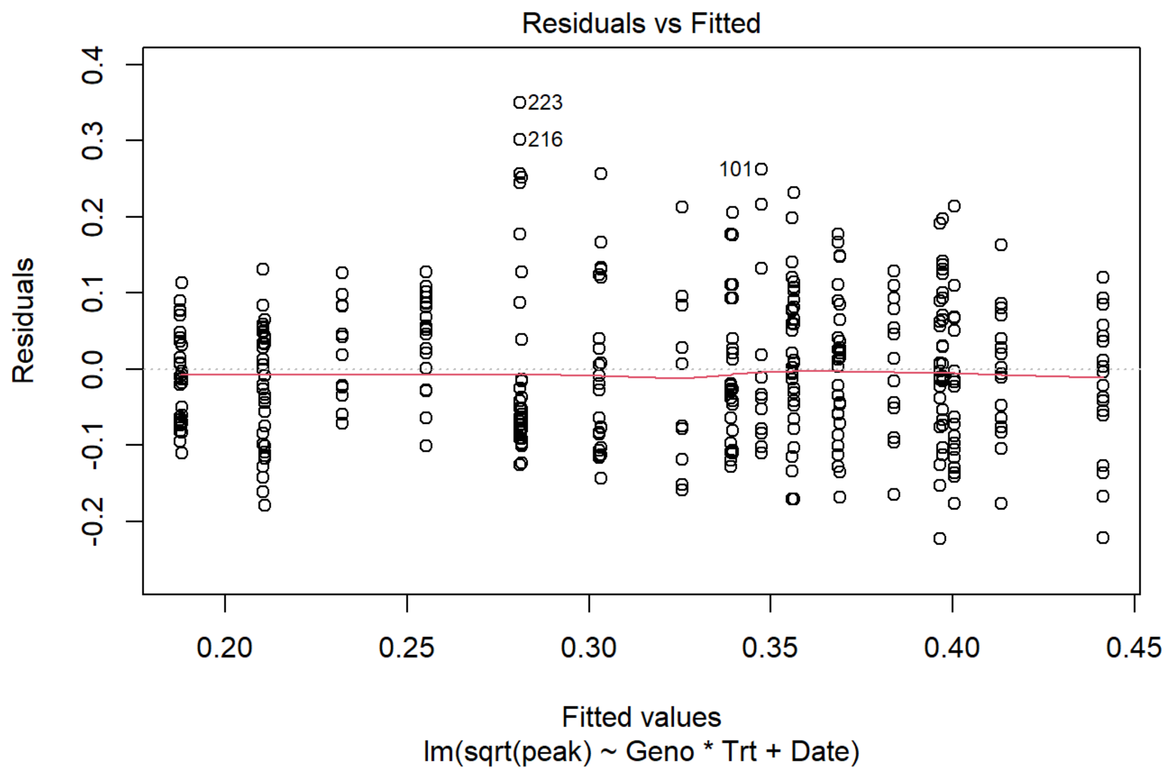


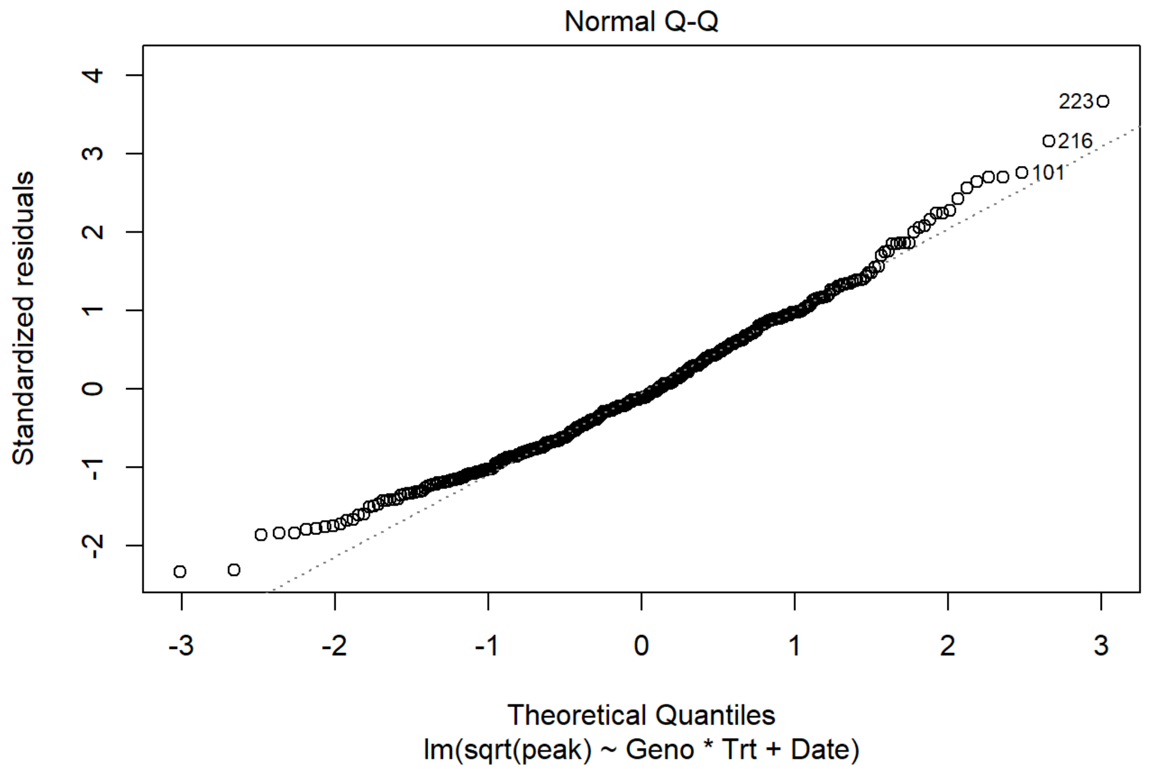


```
anova(peak.lm)
```

```
## Analysis of Variance Table
##
## Response: sqrt(peak)
##            Df Sum Sq Mean Sq F value    Pr(>F)
## Geno        1 0.3265 0.32648 35.0591 7.247e-09 ***
## Trt         3 1.6609 0.55364 59.4523 < 2.2e-16 ***
## Date        2 0.1686 0.08430  9.0528 0.0001448 ***
## Geno:Trt    3 0.0379 0.01264  1.3572 0.2555999
## Residuals 374 3.4828 0.00931
## ---
## Signif. codes:  0 '***' 0.001 '**' 0.01 '*' 0.05 '.' 0.1 ' ' 1
```

```
#could do in one step
#emmeans(peak.lm, specs=pairwise ~ Geno | Trt)

#Set up contrast: looking at effect of genotype in each treatment
peak.em <- emmeans(peak.lm,  ~ Geno | Trt)

#What is the effect of genotype IN each treatment?
contrast(peak.em, interaction = "pairwise")#interaction selected doesn't matter here, only two levels fo
r each
```

```
## Note: Use 'contrast(regrid(object), ...)' to obtain contrasts of back-transformed estimates
```

```
## Trt = Mock:
##  Geno_pairwise            estimate     SE  df t.ratio p.value
##  YC3.6 - (glr2.7/2.8/2.9)   0.0229 0.0193 374 1.184   0.2371
##
## Trt = 1 µM flg22:
##  Geno_pairwise            estimate     SE  df t.ratio p.value
##  YC3.6 - (glr2.7/2.8/2.9)   0.0750 0.0198 374 3.794   0.0002
##
## Trt = 1 µM elf18:
##  Geno_pairwise            estimate     SE  df t.ratio p.value
##  YC3.6 - (glr2.7/2.8/2.9)   0.0658 0.0209 374 3.144   0.0018
##
## Trt = 1 µM Pep1:
##  Geno_pairwise            estimate     SE  df t.ratio p.value
##  YC3.6 - (glr2.7/2.8/2.9)   0.0575 0.0197 374 2.926   0.0036
##
## Results are averaged over the levels of: Date
## Note: contrasts are still on the sqrt scale
```

### Panel B

The same for salt

```
setwd(home)
Amock <- ReadSalt("2020-09-15_p3a_FastMock.xlsx", "2020-09-15_plan.xlsx", "Mock", "", "A")
#Remove wells: check individual day for each one
Amock <- Amock %>% filter(!Well %in% c())

Asalt <- ReadSalt("2020-09-15_p3b_333mMNaCl.xlsx", "2020-09-15_plan.xlsx", "NaCl", "333 mM ", "A")
#Remove wells: check individual day for each one
Asalt <- Asalt %>% filter(!Well %in% c("A9","B7", "B9", "B11","E8","F11","G9","H11")) %>% mutate(Column=
Column-6)

Bmock <- ReadSalt("2020-10-02_p1a_mock.xlsx", "2020-10-02_plan.xlsx", "Mock", "", "B")
#Remove wells: check individual day for each one
Bmock <- Bmock %>% filter(!Well %in% c("E2")) %>% mutate(Column=Column+6)

Bsalt <- ReadSalt("2020-10-02_p1b_333mMNaCl.xlsx", "2020-10-02_plan.xlsx", "NaCl", "333 mM ", "B")
#Remove wells: check individual day for each one
Bsalt <- Bsalt %>% filter(!Well %in% c("B7", "B11","E9","F11"))

Cmock <- ReadSalt("2020-10-02_p2a_mock.xlsx", "2020-10-02_plan.xlsx", "Mock", "", "C")
#Remove wells: check individual day for each one
Cmock <- Cmock %>% filter(!Well %in% c("F4","H3")) %>% mutate(Column=Column+12)

Csalt <- ReadSalt("2020-10-02_p2b_333mMNaCl.xlsx", "2020-10-02_plan.xlsx", "NaCl", "333 mM ", "C")
#Remove wells: check individual day for each one
Csalt <- Csalt %>% filter(!Well %in% c("B11")) %>% mutate(Column=Column+6)
```

Bind all data together, then find and remove wells (by date) where baseYFP is not above 3x level of non-fluorescent Col-0 control, to deal with some silencing. Next normalize ratios to pretreatment values, combine genotypes (tested seedlings from 2-3 different parent plants, all parents homozygous) and clean up names.

```
fluor <- rbind(Amock, Asalt, Bmock, Bsalt, Cmock, Csalt)
Conc <- paste0("333 mM ")
Trt <- "NaCl, and Mock"

#for now getting one Col-0 fluorescence value across all exp. In theory (and from seeing many of these)
 it shouldn't change much
Col_YFP <- fluor %>% filter(`Time (min)`<0) %>% filter(Genotype=="Col-0") %>% group_by(Well) %>% summari
ze(wellmean = mean(YFP)) %>% ungroup() %>% summarize(mean=mean(wellmean))
```

```
## `summarise()` ungrouping output (override with `.groups` argument)
```

```
Col_CFP <- fluor %>% filter(`Time (min)`<0) %>% filter(Genotype=="Col-0") %>% group_by(Well) %>% summari
ze(wellmean = mean(CFP)) %>% ungroup() %>% summarize(mean=mean(wellmean))
```

```
## `summarise()` ungrouping output (override with `.groups` argument)
```

```
#select only if THREE times Col-0 mean. (previously 5x, don't think I need to be that stringent)
fluor_judge <- fluor %>% filter(`Time (min)`<0)  %>% group_by(Date, Well, Trt, Genotype) %>% summarize(m
eanYFP=mean(YFP), meanCFP=mean(CFP)) %>% mutate(YFP_Judgement = if_else(meanYFP > Col_YFP*3, "Yes", "No"
)) %>% mutate(CFP_Judgement = if_else(meanCFP > Col_CFP*3,"Yes", "No")) %>% mutate(Judgement = if_else(Y
FP_Judgement =="Yes" & CFP_Judgement=="Yes", "Yes", "No"))
```

```
## `summarise()` regrouping output by 'Date', 'Well', 'Trt' (override with `.groups` argument)
```

```
fluor <-  merge(fluor, fluor_judge[,c("Date", "Well", "Judgement", "Trt")], by=c("Date", "Well", "Trt"))

pretreatnorm <- function(time, ratio){
  ratio.df <- data.frame(`Time (min)`=time, "ratio"=ratio)
  colnames(ratio.df) <- c("Time (min)","ratio")
  pretreat <- mean(ratio.df$ratio[ratio.df$`Time (min)`<0])
  ratio.df$difference <- (ratio.df$ratio-pretreat)/pretreat
  return(ratio.df$difference)
}

norm_0 <- fluor %>% group_by(Date, Well, Trt, Genotype,Judgement) %>% arrange(`Time (min)`) %>% mutate(d
ifference=pretreatnorm(`Time (min)`, ratio))

filtered <- norm_0[norm_0$Judgement=="Yes",]

filtered <- filtered %>% ungroup() %>% add_column(Geno = "filler") %>% mutate(Geno = if_else(Genotype ==
"YC3.6 A" | Genotype == "YC3.6 B" | Genotype == "YC3.6 C", "YC3.6", Genotype)) %>% mutate(Geno = if_else
(Genotype =="C2-2C" | Genotype == "C2-2D" , "glr2.7/2.8/2.9", Geno))

filtered$Geno <- factor(filtered$Geno, levels=c("YC3.6","glr2.7/2.8/2.9"))
filtered$Trt <- factor(filtered$Trt, levels=c("Mock","333 mM NaCl"))
```

Plot peaks

```
peak <- filtered %>% filter(`Time (min)` > -0.5) %>% group_by(Date, Geno, Trt, Well) %>% summarize(peak=
max(difference)) %>% ungroup() %>% na.omit()
```

```
## `summarise()` regrouping output by 'Date', 'Geno', 'Trt' (override with `.groups` argument)
```

```
peak %>% group_by(Geno, Trt) %>% summarize(n=length(peak))
```

```
## `summarise()` regrouping output by 'Geno' (override with `.groups` argument)
```

```
## # A tibble: 4 x 3
## # Groups:   Geno [2]
##   Geno           Trt             n
##   <fct>          <fct>       <int>
## 1 YC3.6          Mock           56
## 2 YC3.6          333 mM NaCl    51
## 3 glr2.7/2.8/2.9 Mock           38
## 4 glr2.7/2.8/2.9 333 mM NaCl    29
```

```
PanelB <- ggplot(peak, aes(x=Trt, y=peak, color=interaction(Geno, Trt)))+
  geom_boxplot(outlier.color=NA, size=0.3)+
  geom_point(position=position_jitterdodge(jitter.width=0.6), aes(fill=interaction(Geno, Trt), shape=Dat
e, group=interaction(Geno, Trt)), size=1, alpha=0.8) +
  labs(x="",y=expression("Peak (R-R"[0]*")/R"[0]), title="", fill="", color="") +
  scale_fill_manual(values=smallmuted)+scale_color_manual(values=smallmuted)+scale_shape_manual(values=c
(21,22,23,24), guide="none")+
  theme_cowplot()+
  theme(legend.position = "none", panel.background = element_blank(), plot.background = element_blank(),
        panel.border = element_blank(),
        axis.line=element_line(size=0.5), axis.ticks=element_line(size=0.2),
        axis.text = element_text(size=6), axis.title = element_text(size=7),
        plot.margin=unit(c(-0.6, 0, -0.35, 0), "cm"))



ggsave(PanelB, filename = "PanelB.pdf", height=4, width=4.5, units="cm",bg="transparent")
```

And statistics: test effect of genotype within each treatment

```
peak %>% group_by(Geno, Trt) %>% count()
```

```
## # A tibble: 4 x 3
## # Groups:   Geno, Trt [4]
##   Geno           Trt              n
##   <fct>          <fct>        <int>
## 1 YC3.6          Mock            56
## 2 YC3.6          333 mM NaCl     51
## 3 glr2.7/2.8/2.9 Mock            38
## 4 glr2.7/2.8/2.9 333 mM NaCl     29
```

```
peak.lm <- lm(sqrt(peak) ~ Geno*Trt+Date, data=peak)
```

```
## Warning in sqrt(peak): NaNs produced
```

```
plot(peak.lm, 1:2)
```

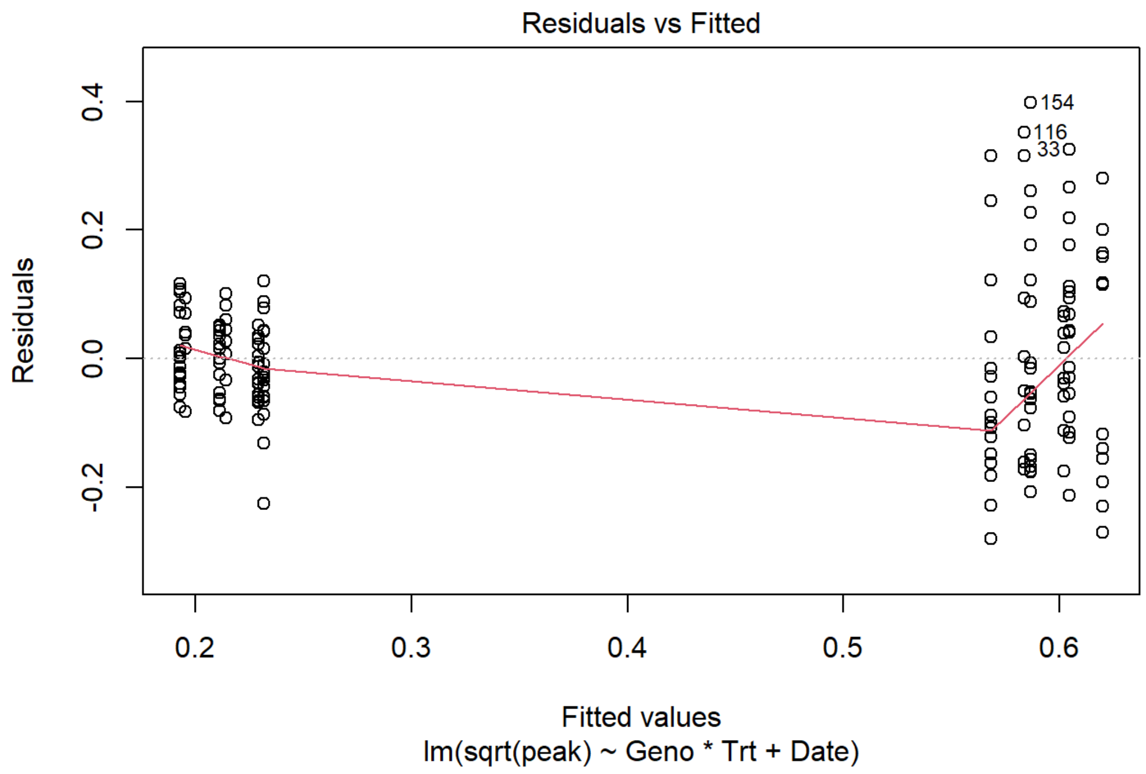


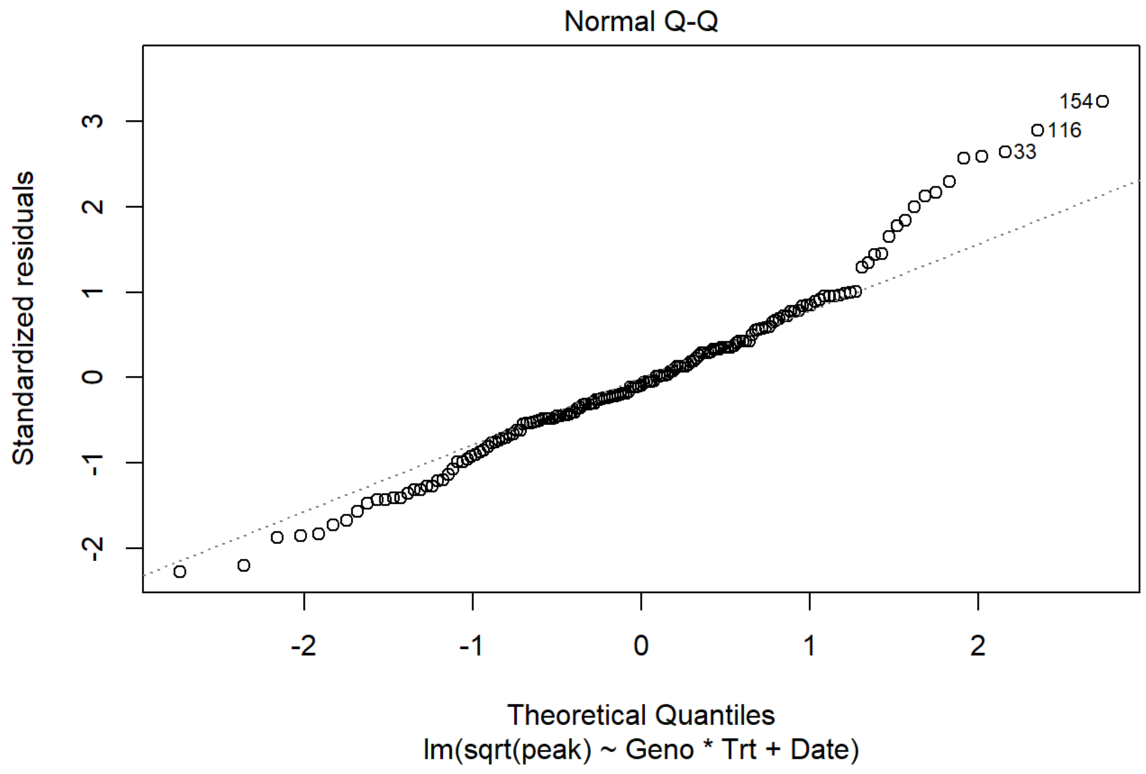


```
anova(peak.lm)
```

```
## Analysis of Variance Table
##
## Response: sqrt(peak)
##            Df Sum Sq Mean Sq  F value Pr(>F)
## Geno        1 0.0000  0.0000   0.0021 0.9637
## Trt         1 5.8577  5.8577 375.7212 <2e-16 ***
## Date        2 0.0350  0.0175   1.1214 0.3284
## Geno:Trt    1 0.0016  0.0016   0.0998 0.7524
## Residuals 157 2.4477  0.0156
## ---
## Signif. codes:  0 '***' 0.001 '**' 0.01 '*' 0.05 '.' 0.1 ' ' 1
```

```
#could do in one step
#emmeans(peak.lm, specs=pairwise ~ Geno | Trt)

#Set up contrast: looking at effect of genotype in each treatment
peak.em <- emmeans(peak.lm,  ~ Geno | Trt)

#What is the effect of genotype IN each treatment?
contrast(peak.em, interaction = "pairwise")#interaction selected doesn't matter here, only two levels fo
r each
```

```
## Note: Use 'contrast(regrid(object), ...)' to obtain contrasts of back-transformed estimates
```

```
## Trt = Mock:
##  Geno_pairwise            estimate     SE  df t.ratio p.value
##  YC3.6 - (glr2.7/2.8/2.9)  -0.0027 0.0283 157 -0.095  0.9243
##
## Trt = 333 mM NaCl:
##  Geno_pairwise            estimate     SE  df t.ratio p.value
##  YC3.6 - (glr2.7/2.8/2.9)  -0.0155 0.0291 157 -0.532  0.5954
##
## Results are averaged over the levels of: Date
## Note: contrasts are still on the sqrt scale
```

### Panel C

Infiltration infection experiments with glr CRISPR line. First clean up workspace and make required functions/palettes

```
rm(list=ls())
home <- "D:/OneDrive/Norwich_U/DATA- IR assays/"
monochrome <- c("black","grey","white", "black","grey")

#These two functions read in treatment data, from an excel file. Set up the excel file in the orientatio
n in which the plate(s) went into the camera EXCEPT if there are two plates, stack them vertically not h
orizontally
makelist <- function(filename, sheet){
  worksheet <- sheet
  print(worksheet)
  variable <- read_excel(filename, sheet = worksheet)
  variablelist <- variable[1,]
  for(i in 2:nrow(variable)){
    variablelist <- c(variablelist, variable[i,])
  }
  variablevector <- unlist(variablelist)
  return(variablevector)
}

add_treatments <- function(filename, inheritedros){
  looper <- excel_sheets(filename)
  for(i in 1:length(looper)){
    column <- looper[i]
    columnlist <- makelist(filename, column)
    inheritedros$new <- columnlist
    colnames(inheritedros)[colnames(inheritedros)=="new"] <- column
  }
  return(inheritedros)
}

CollectInfect <- function(date){
  #make empty data frame to hold treatment info
  empty <- data.frame(first=1:96)
  #this calls the two functions above to add one column for each sheet in treatment
  Filled <- add_treatments(paste0(home, date, "_guide.xlsx"), empty)
  Filled$LeafDiscs <- as.numeric(Filled$LeafDiscs)
  Filled <- Filled[Filled$LeafDiscs > 0 & Filled$Genotype != "none",]
  Filled$Date <- date
  #Get colony counts and merge with treatment information
  colonies <- read_excel(paste0(home,date,".xlsx")) %>% unite("Well", c("row", "column"), sep="") %>% se
lect("Plate","Well","original CFU", "drop volume","disc volume")
  Filled <- merge(colonies, Filled, by=c("Plate","Well"))
  #calculate CFU per cm2
  Filled$`CFU/cm2` <-  Filled$`original CFU`/Filled$`drop volume`*Filled$`disc volume`/(Filled$LeafDiscs
*0.2^2*pi)
  Filled$log <- log10(Filled$`CFU/cm2`)
  return(Filled)
}
```

Then bring in data

```
#Can't use function because two plates of data
#make empty data frame to hold treatment info
  empty <- data.frame(first=1:192)
#this calls the two functions above to add one column for each sheet in treatment
  Filled <- add_treatments(paste0(home, "2020-08-15", "_guide.xlsx"), empty)
```

```
## [1] "Genotype"
## [1] "Bacteria"
## [1] "LeafDiscs"
## [1] "Bugs"
## [1] "Well"
## [1] "Plate"
## [1] "Tray"
## [1] "Chamber"
```

```
  Filled$LeafDiscs <- as.numeric(Filled$LeafDiscs)
  Filled <- Filled[Filled$LeafDiscs > 0 & Filled$Genotype != "none",]
  Filled$Date <- "2020-08-15"
  #Get colony counts and merge with treatment information
  colonies <- read_excel(paste0(home,"2020-08-15",".xlsx")) %>% unite("Well", c("row", "column"), sep=""
) %>% select("Plate","Well","original CFU", "drop volume","disc volume")
  Filled <- merge(colonies, Filled, by=c("Plate","Well"))
  #calculate CFU per cm2
  Filled$`CFU/cm2` <-  Filled$`original CFU`/Filled$`drop volume`*Filled$`disc volume`/(Filled$LeafDiscs
*0.2^2*pi)
  Filled$log <- log10(Filled$`CFU/cm2`)
AB <- Filled
A <- AB[AB$Chamber=="17",]
A$Date <- "2020-08-15b"
B <- AB[AB$Chamber=="18",]

C <- CollectInfect("2020-08-21")
```

```
## [1] "Genotype"
## [1] "Bacteria"
## [1] "LeafDiscs"
## [1] "Bugs"
## [1] "Well"
## [1] "Plate"
## [1] "Tray"
## [1] "Chamber"
```

```
All <- rbind(A, B, C) %>% filter(Genotype != "pad4-1") %>% filter(Bacteria=="DC3000")
All$Genotype <- factor(All$Genotype, levels=c("Col-0", "glr CRISPR 2.6B", "bak1-5", "YC3.6","C1-2"))

print(All[,c(13, 7, 3, 9, 14:15)])
```

```
##            Date        Genotype original CFU LeafDiscs   CFU/cm2      log
## 1    2020-08-15b           YC3.6       138000         6 1830281.8 6.262518
## 2    2020-08-15b           YC3.6       126000         6 1671126.9 6.223009
## 3    2020-08-15b           YC3.6        79000         6 1047770.0 6.020266
## 4    2020-08-15b           YC3.6        95000         6 1259976.6 6.100362
## 5    2020-08-15b           YC3.6        68000         3 1803756.0 6.256178
## 6    2020-08-15b           YC3.6        95000         6 1259976.6 6.100362
## 13   2020-08-15b           C1-2       300000         6 3978873.6 6.599760
## 14   2020-08-15b           C1-2       350000         6 4642019.2 6.666707
## 15   2020-08-15b           C1-2       400000         6 5305164.8 6.724699
## 16   2020-08-15b           C1-2       540000         6 7161972.4 6.855033
## 17   2020-08-15b           C1-2       490000         6 6498826.8 6.812835
## 18   2020-08-15b           C1-2       390000         6 5172535.7 6.713703
## 37   2020-08-15b          bak1-5       310000         6 4111502.7 6.614001
## 42   2020-08-15b          bak1-5       400000         6 5305164.8 6.724699
## 44   2020-08-15b          bak1-5       350000         6 4642019.2 6.666707
## 53   2020-08-15b glr CRISPR 2.6B       460000         6 6100939.5 6.785397
## 55   2020-08-15b glr CRISPR 2.6B       580000         6 7692488.9 6.886067
## 56   2020-08-15b glr CRISPR 2.6B       370000         6 4907277.4 6.690841
## 57   2020-08-15b glr CRISPR 2.6B       510000         6 6764085.1 6.830209
## 61   2020-08-15b           Col-0       123000         3 3262676.3 6.513574
## 66   2020-08-15b           Col-0       227000         6 3010681.0 6.478665
## 77   2020-08-15b          bak1-5       142000         6 1883333.5 6.274927
## 79   2020-08-15b          bak1-5       217000         6 2878051.9 6.459099
## 81   2020-08-15b          bak1-5       330000         6 4376760.9 6.641153
## 85   2020-08-15b           Col-0       112000         6 1485446.1 6.171857
## 90   2020-08-15b           Col-0        83000         6 1100821.7 6.041717
## 92   2020-08-15b           Col-0       158000         6 2095540.1 6.321296
## 101  2020-08-15b glr CRISPR 2.6B       380000         6 5039906.5 6.702422
## 103  2020-08-15b glr CRISPR 2.6B       440000         6 5835681.2 6.766092
## 105  2020-08-15b glr CRISPR 2.6B       168000         6 2228169.2 6.347948
## 26    2020-08-15          bak1-5       490000         6 6498826.8 6.812835
## 28    2020-08-15          bak1-5       430000         6 5703052.1 6.756107
## 35    2020-08-15          bak1-5       720000         6 9549296.6 6.979971
## 39    2020-08-15           Col-0       189000         6 2506690.4 6.399101
## 46    2020-08-15           Col-0       217000         6 2878051.9 6.459099
## 48    2020-08-15           Col-0       570000         6 7559859.8 6.878514
## 51    2020-08-15 glr CRISPR 2.6B       610000         6 8090376.3 6.907969
## 52    2020-08-15 glr CRISPR 2.6B       450000         6 5968310.4 6.775851
## 59    2020-08-15 glr CRISPR 2.6B       450000         6 5968310.4 6.775851
## 74    2020-08-15           Col-0       101000         6 1339554.1 6.126960
## 76    2020-08-15           Col-0       139000         6 1843544.8 6.265654
## 83    2020-08-15           Col-0       196000         6 2599530.7 6.414895
## 87    2020-08-15 glr CRISPR 2.6B       146000         6 1936385.1 6.286992
## 94    2020-08-15 glr CRISPR 2.6B       240000         6 3183098.9 6.502850
## 96    2020-08-15 glr CRISPR 2.6B       177000         6 2347535.4 6.370612
## 111   2020-08-15          bak1-5       540000         6 7161972.4 6.855033
## 118   2020-08-15          bak1-5        97000         6 1286502.5 6.109411
## 120   2020-08-15          bak1-5       213000         6 2825000.2 6.451018
## 210   2020-08-21 glr CRISPR 2.6B       300000         6 3978873.6 6.599760
## 410   2020-08-21 glr CRISPR 2.6B       185000         6 2453638.7 6.389811
## 510   2020-08-21          bak1-5       184000         6 2440375.8 6.387457
## 710   2020-08-21          bak1-5       171000         6 2267957.9 6.355635
## 910   2020-08-21          bak1-5       181000         6 2400587.1 6.380317
## 1110  2020-08-21 glr CRISPR 2.6B       195000         6 2586267.8 6.412673
## 131   2020-08-21 glr CRISPR 2.6B       254000         6 3368779.6 6.527473
## 151   2020-08-21           Col-0       110000         6 1458920.3 6.164032
## 181   2020-08-21 glr CRISPR 2.6B       240000         6 3183098.9 6.502850
## 201   2020-08-21 glr CRISPR 2.6B       204000         6 2705634.0 6.432269
## 221   2020-08-21           Col-0       218000         6 2891314.8 6.461095
## 241   2020-08-21           Col-0       105000         6 1392605.8 6.143828
## 291   2020-08-21           Col-0        60000         6  795774.7 5.900790
## 311   2020-08-21           Col-0        95000         6 1259976.6 6.100362
```

```
## 331   2020-08-21           Col-0        80000         6 1061033.0 6.025729
## 391   2020-08-21          bak1-5       202000         6 2679108.2 6.427990
## 461   2020-08-21          bak1-5       320000         6 4244131.8 6.627789
## 481   2020-08-21          bak1-5       310000         6 4111502.7 6.614001
## 491   2020-08-21           YC3.6        51000         6  676408.5 5.830209
## 501   2020-08-21            C1-2       187000         6 2480164.5 6.394480
## 511   2020-08-21           YC3.6       107000         6 1419131.6 6.152023
## 521   2020-08-21            C1-2       177000         6 2347535.4 6.370612
## 531   2020-08-21            C1-2       174000         6 2307746.7 6.363188
## 541   2020-08-21           YC3.6        23000         6  305047.0 5.484367
## 551   2020-08-21            C1-2       169000         6 2241432.1 6.350526
## 561   2020-08-21           YC3.6       158000         6 2095540.1 6.321296
## 571   2020-08-21            C1-2        66000         6  875352.2 5.942183
## 581   2020-08-21           YC3.6        78000         6 1034507.1 6.014733
## 591   2020-08-21            C1-2       310000         6 4111502.7 6.614001
## 601   2020-08-21           YC3.6        31000         6  411150.3 5.614001
```

plot

```
newlabs <- c("Col-0", expression(italic("glr2.7/2.8/2.9")), expression(italic("bak1-5")), "YC3.6", expre
ssion("YC3.6\n "*italic("glr2.7/2.8/2.9")))
PanelC <- ggplot(All, aes(x=Genotype, y=log, group=Genotype))+
  geom_boxplot(outlier.color=NA, size=0.3, width=0.7) +
  geom_point(position=position_jitter(height=0, width=0.2), aes(shape=as.factor(Date), fill=Genotype), s
ize=1, alpha=0.8, lwd=0.1) +
  labs(x="",y=expression("log"[10]*"(CFU\U2022"*cm^-2*')'), title="") +
  theme_cowplot() +
  scale_fill_manual(values=monochrome, labels=newlabs) +
  scale_x_discrete(labels=newlabs)+
  ylim(c(5.45, 7))+
  scale_shape_manual(values=c(21,22,23,24))+
  theme(legend.position = "none", panel.background = element_blank(), plot.background = element_blank(),
        panel.border = element_blank(), strip.text.x = element_blank(),
        axis.text.x = element_text(size=6, hjust=0.5, vjust=0.5),
        axis.text.y = element_text(size=6),
        axis.title = element_text(size=7),
        axis.line=element_line(size=0.3), axis.ticks=element_line(size=0.5),
        plot.margin=unit(c(-0.5, 0, 0, 0), "cm"))
```

```
## Warning: Duplicated aesthetics after name standardisation: size
```

```
suppressWarnings(ggsave(PanelC, file="PanelC.pdf", height=4, width=12, units="cm", bg="transparent"))
```

and statistics

```
All %>% group_by(Genotype) %>% count()
```

```
## # A tibble: 5 x 2
## # Groups:   Genotype [5]
##   Genotype            n
##   <fct>           <int>
## 1 Col-0              17
## 2 glr CRISPR 2.6B    19
## 3 bak1-5             18
## 4 YC3.6              12
## 5 C1-2               12
```

```
Col <- All %>% filter(!Genotype %in% c("YC3.6","C1-2"))
Col$Genotype <- as.factor(Col$Genotype) %>% relevel("glr CRISPR 2.6B") %>% relevel("Col-0")

Col.lm <- lm(log~Genotype+Date/Tray, data=Col[Col$log >0,])
plot(Col.lm, 1:2)
```

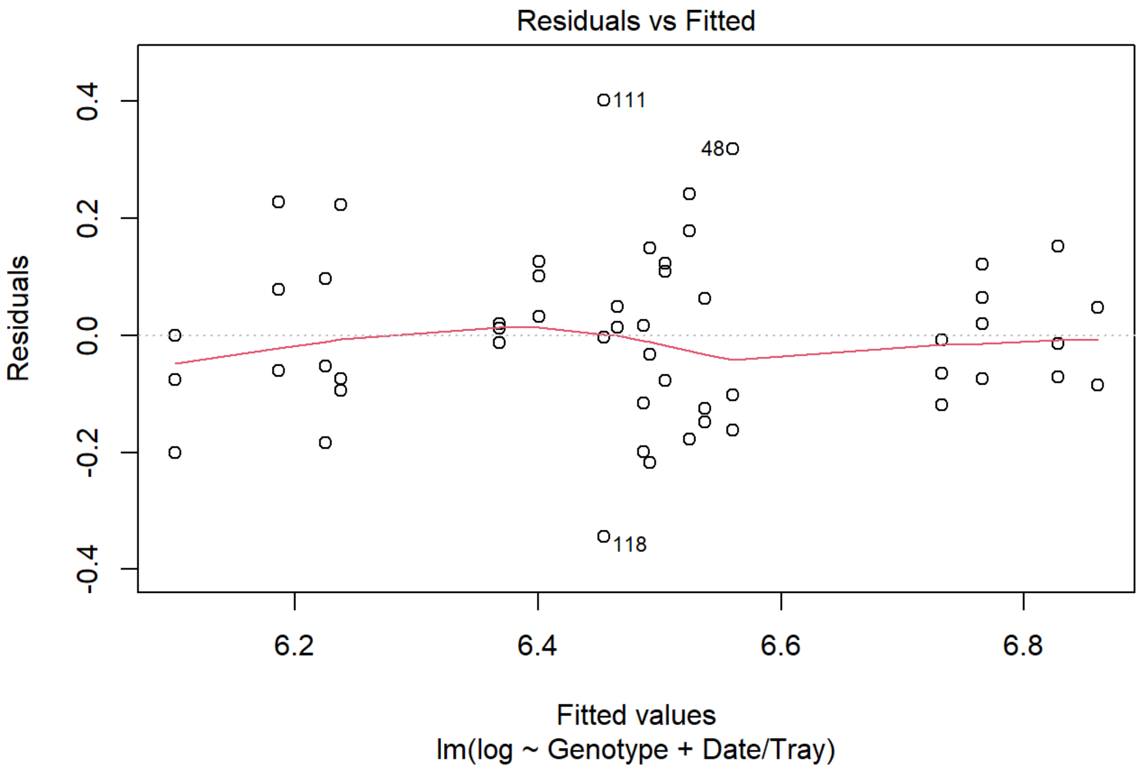


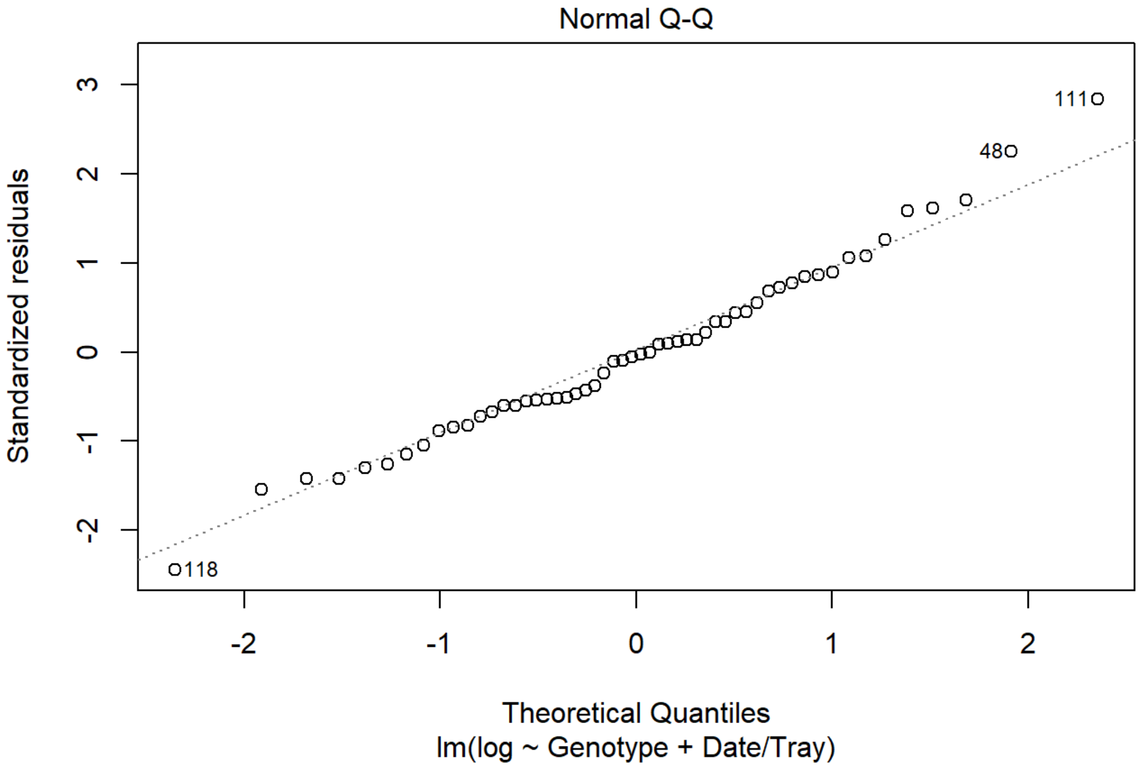


```
anova(Col.lm)
```

```
## Analysis of Variance Table
##
## Response: log
##           Df  Sum Sq Mean Sq F value    Pr(>F)
## Genotype   2 1.05290 0.52645 22.5759 1.475e-07 ***
## Date       2 0.43863 0.21932  9.4051 0.0003764 ***
## Date:Tray  3 0.97020 0.32340 13.8686 1.424e-06 ***
## Residuals 46 1.07267 0.02332
## ---
## Signif. codes:  0 '***' 0.001 '**' 0.01 '*' 0.05 '.' 0.1 ' ' 1
```

```
#compare all genotypes to each Col-0
emmeans(Col.lm, dunnett ~ Genotype)
```

```
## NOTE: A nesting structure was detected in the fitted model:
##     Tray %in% Date
```

```
## $emmeans
##  Genotype        emmean     SE df lower.CL upper.CL
##  Col-0             6.30 0.0371 46     6.22     6.37
##  glr CRISPR 2.6B   6.60 0.0351 46     6.53     6.67
##  bak1-5            6.56 0.0360 46     6.49     6.64
##
## Results are averaged over the levels of: Tray, Date
## Confidence level used: 0.95
##
## $contrasts
##  contrast                  estimate     SE df t.ratio p.value
##  glr CRISPR 2.6B - (Col-0)    0.300 0.0512 46 5.850   <.0001
##  (bak1-5) - (Col-0)           0.267 0.0517 46 5.162   <.0001
##
## Results are averaged over the levels of: Tray, Date
## P value adjustment: dunnettx method for 2 tests
```

```
YC <- All %>% filter(Genotype %in% c("YC3.6","C1-2"))
YC$Genotype <- as.factor(YC$Genotype)  %>% relevel("YC3.6")
YC.lm <- lm(log~Genotype+Date, data=YC[YC$log >0,])
plot(YC.lm, 1:2)
```

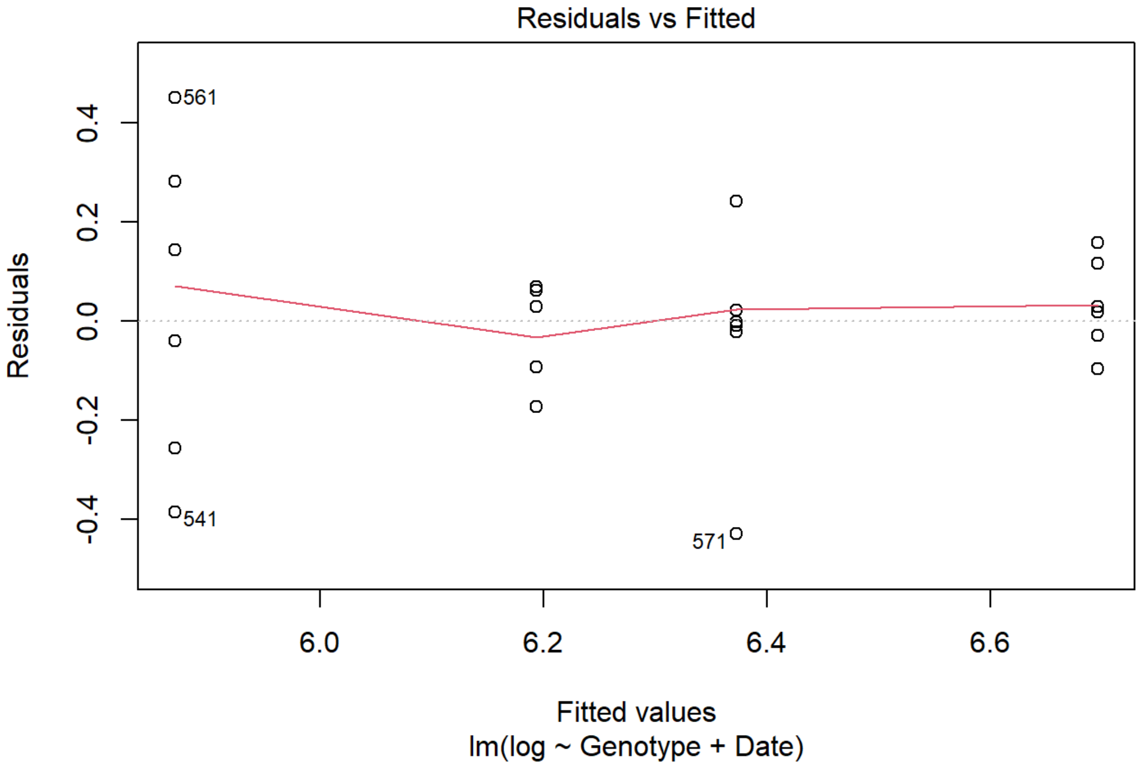
Residuals vs Fitted
Residuals
Fitted values
lm(log ~ Genotype + Date)
0.4
0.2
0.0
-0.2
-0.4
6.0
6.2
6.4
6.6
561
541
571

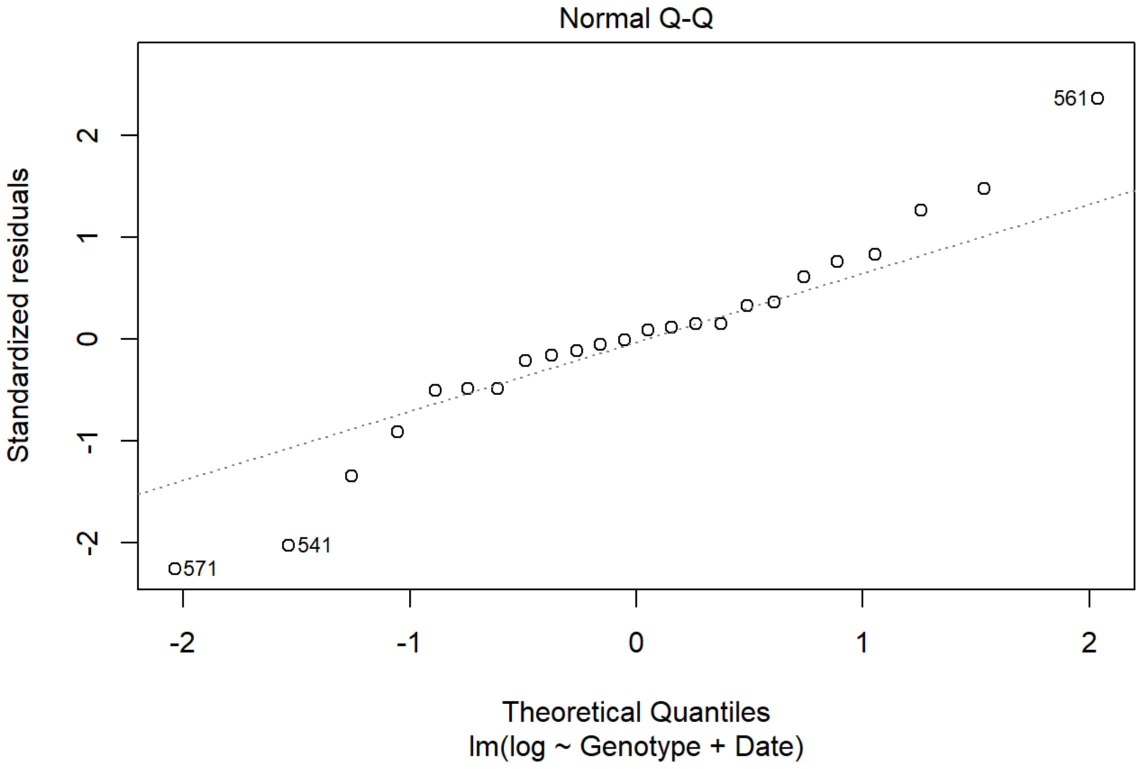
Normal Q-Q
Standardized residuals
Theoretical Quantiles
lm(log ~ Genotype + Date)
2
1
0
-1
-2
561
541
571

```
anova(YC.lm)
```

```
## Analysis of Variance Table
##
## Response: log
##           Df  Sum Sq Mean Sq F value    Pr(>F)
## Genotype   1 1.51423 1.51423  36.481   5.4e-06 ***
## Date       1 0.62850 0.62850  15.142 0.0008423 ***
## Residuals 21 0.87166 0.04151
## ---
## Signif. codes:  0 '***' 0.001 '**' 0.01 '*' 0.05 '.' 0.1 ' ' 1
```

```
#This is meaningless, as there are only two genotypes, include just for consistency with stats above
emmeans(YC.lm, dunnett ~ Genotype)
```

```
## $emmeans
##  Genotype emmean     SE df lower.CL upper.CL
##  YC3.6      6.03 0.0588 21     5.91     6.15
##  C1-2       6.53 0.0588 21     6.41     6.66
##
## Results are averaged over the levels of: Date
## Confidence level used: 0.95
##
## $contrasts
##  contrast       estimate     SE df t.ratio p.value
##  (C1-2) - YC3.6    0.502 0.0832 21 6.040   <.0001
##
## Results are averaged over the levels of: Date
```

### Figure S1

Marta Bjornson

```
knitr::opts_chunk$set(echo = TRUE)
library(DESeq2)
library(tidyverse)
library(ggthemes)
library(cowplot)
library(dendextend)
```

### Panel A

Comparing L2FC calculated here with published data. First, Zipfel et al 2006: elf18 and flg22, 30 min

```
rm(list=ls())
RNAseq <- read.csv("D:/OneDrive/Norwich_Z/Overall/CleanedData/CleanedData_nofilter_ordered.csv", header=
T)

pub <- read.csv("D:/OneDrive/Norwich_Z/Overall/CleanedData/wtonly/Analysis-PublishedResponses/elf18_Zipf
el2006/Processed.csv", header=T)
#R reads one column as a factor, just change that
pub$Col_elf26_30 <- as.numeric(as.character(pub$Col_elf26_30))
```

```
## Warning: NAs introduced by coercion
```

```
all <- merge(RNAseq, pub, by="ATG")

m <- lm(Col_elf26_30~elf18_030, data=all)

elf18 <- ggplot(all, aes(x=elf18_030, y=Col_elf26_30))+
  geom_point(size=0.2)+geom_smooth(method="lm", formula=y~x, color="red")+
  labs(x=expression('log'[2]*'(fold change), this study'),y=expression('log'[2]*'(fold change), Zipfel e
t al 2006'), title="elf18/26 response at 30 min")+
  annotate(geom="text", label=bquote('R'^2*' = '*.(round(summary(m)$r.squared, 3))), x=0.8*min(all$elf18
_030, na.rm=T), y=0.8*max(all$Col_elf26_30, na.rm=T),hjust=0, size=2)+
  theme_minimal(base_size=6)
ggsave("elf18.pdf", elf18,height=4, width=4, units="cm")
```

```
## Warning: Removed 1 rows containing non-finite values (stat_smooth).
```

```
## Warning: Removed 1 rows containing missing values (geom_point).
```

```
## Warning in is.na(x): is.na() applied to non-(list or vector) of type 'language'
```

```
m <- lm(Col_flg_30~flg22_030, data=all)



flg22 <- ggplot(all, aes(x=flg22_030, y=Col_flg_30))+
  geom_point(size=0.2)+geom_smooth(method="lm", formula=y~x, color="red")+
  labs(x=expression('log'[2]*'(fold change), this study'),y=expression('log'[2]*'(fold change), Zipfel e
t al 2006'), title="flg22 response at 30 min")+
  annotate(geom="text", label=bquote('R'^2*' = '*.(round(summary(m)$r.squared, 3))), x=0.8*min(all$elf18
_030, na.rm=T), y=0.8*max(all$Col_elf26_30, na.rm=T),hjust=0, size=2)+
  theme_minimal(base_size=6)
ggsave("flg22.pdf", flg22, height=4, width=4, units="cm")
```

```
## Warning in is.na(x): is.na() applied to non-(list or vector) of type 'language'
```

Wan et al 2008: CO8, 30 min

```
rm(list=ls())
RNAseq <- read.csv("D:/OneDrive/Norwich_Z/Overall/CleanedData/CleanedData_nofilter_ordered.csv", header=
T)

pub <- read.csv("D:/OneDrive/Norwich_Z/Overall/CleanedData/wtonly/Analysis-PublishedResponses/CO8_Wan200
8/Processed.csv", header=T)

all <- merge(RNAseq, pub, by="ATG")

m <- lm(Wan_CO8_30min~CO8_030, data=all)

CO8 <- ggplot(all, aes(x=CO8_030, y=Wan_CO8_30min))+
  geom_point(size=0.2)+geom_smooth(method="lm", formula=y~x, color="red")+
  labs(x=expression('log'[2]*'(fold change), this study'),y=expression('log'[2]*'(fold change), Wan et a
l 2008'), title="CO8 response at 30 min")+
  annotate(geom="text", label=bquote('R'^2*' = '*.(round(summary(m)$r.squared, 3))), x=0.8*min(all$CO8_0
30, na.rm=T), y=0.8*max(all$Wan_CO8_30min, na.rm=T),hjust=0, size=2)+
  theme_minimal(base_size=6)
ggsave("CO8.pdf", CO8, height=4, width=4, units="cm")
```

```
## Warning in is.na(x): is.na() applied to non-(list or vector) of type 'language'
```

### Panel B

Denoux et al 2006: flg, 3 hr

```
rm(list=ls())
RNAseq <- read.csv("D:/OneDrive/Norwich_Z/Overall/CleanedData/CleanedData_nofilter_ordered.csv", header=
T)

pub <- read.csv("D:/OneDrive/Norwich_Z/Overall/CleanedData/wtonly/Analysis-PublishedResponses/OGandFLG_D
enoux2006/Processed.csv", header=T)

all <- merge(RNAseq, pub, by="ATG")

m <- lm(Flg22_3h~flg22_180, data=all)

flg <- ggplot(all, aes(x=flg22_180, y=Flg22_3h))+
  geom_point(size=0.2)+geom_smooth(method="lm", formula=y~x, color="red")+
  labs(x=expression('log'[2]*'(fold change), this study'),y=expression('log'[2]*'(fold change), Denoux e
t al 2006'), title="flg22 response at 3 h")+
  annotate(geom="text", label=bquote('R'^2*' = '*.(round(summary(m)$r.squared, 3))), x=0.8*min(all$flg22
_180, na.rm=T), y=0.8*max(all$Flg22_3h, na.rm=T),hjust=0, size=2)+
  theme_minimal(base_size=6)
ggsave("flg_3hr.pdf", flg, height=4, width=4, units="cm")
```

```
## Warning in is.na(x): is.na() applied to non-(list or vector) of type 'language'
```

### Panel C

PCA, based on counts but filtered for genes called differentially expressed. (not shown here but very similar when done with full counts). Start by loading in counts data (pseudocounts after variance-stabilizing transformation, from DESeq2)

```
rm(list=ls())
home="D:/OneDrive/Norwich_Z/Overall/CleanedData/"
definedvibrant <- c("0"="white", "flg22"="#0077BB", "elf18"="#33BBEE", "nlp20"="#EE7733", "Pep1"="#00998
8", "OGs"="#CC3311", "CO8"="#EE3377", "3-OH-FA"="#DDCC77", "mock"="#BBBBBB")

load(paste0(home, "cleaned_ddsT"))
counts <- counts(ddsT, normalized=T)
counts <- counts %>% as.data.frame() %>% rownames_to_column("ATG")

long <- counts %>% gather(-ATG, key="fullname", value="counts") # this step destroys information on re
p... fine for PCA
long <- long %>% separate(fullname, into=c('Genotype', 'elicitor', 'timerep'), sep="_")
long$rep <- substr(long$timerep, 4,4)
long$time <- substr(long$timerep,1,3)
long <- long %>% group_by(ATG, Genotype, elicitor, time) %>% summarize(avg = mean(counts))
```

```
## `summarise()` regrouping output by 'ATG', 'Genotype', 'elicitor' (override with `.groups` argument)
```

```
long$elicitor[long$elicitor=="3-OH10"] <- "3-OH-FA"
long$elicitor[long$elicitor=="chitooct"] <- "CO8"

long <- long %>% unite(fullname, Genotype, elicitor, time, sep="_")
```

Now filter for genes found differentially expresed, perform PCA and get data into format for ggplot2

```
wide <- spread(long, key=fullname, value=avg)

#Load in changed genes (~10,000)
changed <- read.csv(paste0(home,"wtonly/AllGenesChangedInfo.csv"))
wide <- wide[wide$ATG %in% changed$ATG,,]

wide <- wide %>% column_to_rownames("ATG") %>% t() %>% as.matrix()

pc <- prcomp(wide, scale.=T)
df_out <- as.data.frame(pc$x)
df_out$Genotype <- sapply( strsplit(as.character(row.names(df_out)), "_"), "[[", 1 )
df_out$Elicitor <- sapply( strsplit(as.character(row.names(df_out)), "_"), "[[", 2 )
df_out$Time <- sapply( strsplit(as.character(row.names(df_out)), "_"), "[[", 3 )

df_out$Elicitor <- factor(df_out$Elicitor, levels=c("flg22","elf18","Pep1","nlp20","OGs","CO8","3-OH-FA"
, "mock"))
std_dev <- pc$sdev
pr_var <- std_dev^2
prop_varex <- pr_var/sum(pr_var)
prop_varex <- prop_varex*100
prop_varex <- round(prop_varex)
df_out$response <- "Receptor mutant"
df_out$response[df_out$Genotype=="Col"] <- "Wild type"
```

and plot

```
PanelC <- ggplot(df_out, aes(x=PC1, y=-PC2, color=Elicitor, shape=Time)) +geom_point(size=0.7, stroke=1)
+
  scale_shape_manual(values=c(1,0,4,2,3,5)) +scale_color_manual(values=definedvibrant) +
  theme_minimal(base_size=7) +labs(x=paste0("PC1:", prop_varex[1],"% variance"), y=paste0("PC2:", prop_v
arex[2],"% variance"), color="Pattern")+
  facet_wrap(~response) +
  theme(rect=element_rect(fill="transparent"),legend.key.size=unit(0.12,"cm"), legend.spacing=unit(0.01,
"cm"))
ggsave(PanelC, filename="PanelC.pdf", height=4, width=8, units="cm", bg="transparent")
```

### Panel D

Correlation heatmap of treatment/timepoints with each other. First subset data above to only Col-0, greater than time 0 and all numeric, then calculate correlation

```
rm(list=ls())
home="D:/OneDrive/Norwich_Z/Overall/CleanedData/"

#From results objects, get gene lists p<0.05 L2FC >1, and combine into one large file (expect ~6500)
OnlySigl2FC <- function(TrtTime){
  filepath=paste0(home,"ResultsObjects/",TrtTime,"_results")
  load(filepath)
  resdf <- as.data.frame(res)
  nonsig <- subset(resdf, padj >0.05 | padj == "NA")
  smallchange <- subset(resdf, log2FoldChange <  1 & log2FoldChange > -1)
  onlysig <- resdf
  onlysig[rownames(onlysig) %in% rownames(nonsig),] <- "0"
  onlysig[rownames(onlysig) %in% rownames(smallchange),] <- "0"
  sigFCs <- subset(onlysig, select = "log2FoldChange")
  sigFCs$log2FoldChange <- as.numeric(sigFCs$log2FoldChange)
  names(sigFCs) <- TrtTime
  return(sigFCs)
}

PAMPTime <- c("Col_LPS_005","Col_LPS_010","Col_LPS_030","Col_LPS_090","Col_LPS_180","Col_ch8_005","Col_c
h8_010","Col_ch8_030","Col_ch8_090","Col_ch8_180","Col_elf18_005","Col_elf18_010","Col_elf18_030","Col_e
lf18_090","Col_elf18_180","Col_flg22_005","Col_flg22_010","Col_flg22_030","Col_flg22_090","Col_flg22_18
0","Col_nlp20_005","Col_nlp20_010","Col_nlp20_030","Col_nlp20_090","Col_nlp20_180","Col_OGs_005","Col_OG
s_010","Col_OGs_030","Col_OGs_090","Col_OGs_180","Col_Pep1_005","Col_Pep1_010","Col_Pep1_030","Col_Pep1_
090","Col_Pep1_180")
log2FClist <- lapply(PAMPTime, OnlySigl2FC)
log2FCdf <- do.call("cbind", log2FClist)
nrow(log2FCdf)
```

```
## [1] 26397
```

reduce to genes significantly upregulated in at least one time for any of the seven (expect ~10,000)

```
altered <- log2FCdf %>% rownames_to_column("ATG") %>% gather(-"ATG", key=Sample, value=L2FC) %>% separat
e(Sample, c("Geno","elicitor","time"), sep="_") %>% group_by(elicitor, ATG) %>% filter(sum(L2FC)!=0)  %
>% ungroup() %>% group_by(ATG, time) %>% add_count() %>% filter(n>0) %>% unite(Sample, c("elicitor","tim
e")) %>% spread(key=Sample, value=L2FC)


nrow(altered)
```

```
## [1] 10730
```

```
AllL2FC <- function(TrtTime){
  filepath=paste0(home,"ResultsObjects/",TrtTime,"_results")
  load(filepath)
  resdf <- as.data.frame(res)
  allFCs <- subset(resdf, select = "log2FoldChange")
  allFCs$log2FoldChange <- as.numeric(allFCs$log2FoldChange)
  names(allFCs) <- TrtTime
  return(allFCs)
}

PAMPTime <- c("Col_LPS_005","Col_LPS_010","Col_LPS_030","Col_LPS_090","Col_LPS_180","Col_ch8_005","Col_c
h8_010","Col_ch8_030","Col_ch8_090","Col_ch8_180","Col_elf18_005","Col_elf18_010","Col_elf18_030","Col_e
lf18_090","Col_elf18_180","Col_flg22_005","Col_flg22_010","Col_flg22_030","Col_flg22_090","Col_flg22_18
0","Col_nlp20_005","Col_nlp20_010","Col_nlp20_030","Col_nlp20_090","Col_nlp20_180","Col_OGs_005","Col_OG
s_010","Col_OGs_030","Col_OGs_090","Col_OGs_180","Col_Pep1_005","Col_Pep1_010","Col_Pep1_030","Col_Pep1_
090","Col_Pep1_180")
log2FClist <- lapply(PAMPTime, AllL2FC)
log2FCdf <- do.call("cbind", log2FClist)
nrow(log2FCdf)
```

```
## [1] 26397
```

```
log2FC <- log2FCdf[rownames(log2FCdf) %in% altered$ATG,]
```

Get correlation matrix for heatmap

```
cor.mat <- cor(log2FC)
cor.df <- as.data.frame(cor.mat)

#for text
longcor.df <- cor.df %>% rownames_to_column("First") %>% pivot_longer(-First, names_to="Second", values_
to="pearson") %>% filter(pearson != 1)
(longcor.df %>% filter(str_detect(First, "005")) %>% filter(str_detect(Second, "005")) %>% summarize(mea
n=mean(pearson)))
```

```
## # A tibble: 1 x 1
##     mean
##    <dbl>
## 1 0.0753
```

```
(longcor.df %>% filter(str_detect(First, "010")) %>% filter(str_detect(Second, "010")) %>% summarize(mea
n=mean(pearson)))
```

```
## # A tibble: 1 x 1
##    mean
##   <dbl>
## 1 0.490
```

```
(longcor.df %>% filter(str_detect(First, "030")) %>% filter(str_detect(Second, "030")) %>% summarize(mea
n=mean(pearson)))
```

```
## # A tibble: 1 x 1
##    mean
##   <dbl>
## 1 0.898
```

```
(longcor.df %>% filter(str_detect(First, "090")) %>% filter(str_detect(Second, "090")) %>% summarize(mea
n=mean(pearson)))
```

```
## # A tibble: 1 x 1
##    mean
##   <dbl>
## 1 0.804
```

```
(longcor.df %>% filter(str_detect(First, "180")) %>% filter(str_detect(Second, "180")) %>% summarize(mea
n=mean(pearson)))
```

```
## # A tibble: 1 x 1
##    mean
##   <dbl>
## 1 0.713
```

```
#I made the tree, looked at how i wanted the leaves arranged, and made this list.
leaf_order <- c("Col_ch8_180","Col_OGs_180","Col_elf18_180","Col_Pep1_180","Col_LPS_180","Col_LPS_090",
"Col_ch8_090","Col_nlp20_180","Col_flg22_180","Col_OGs_090","Col_elf18_090","Col_Pep1_090","Col_flg22_09
0","Col_nlp20_090","Col_OGs_030","Col_elf18_030","Col_Pep1_030","Col_nlp20_030","Col_flg22_030","Col_ch8
_030","Col_LPS_030","Col_OGs_010","Col_elf18_010","Col_flg22_010","Col_Pep1_010","Col_nlp20_010","Col_ch
8_010","Col_OGs_005","Col_LPS_010","Col_LPS_005","Col_Pep1_005","Col_elf18_005","Col_nlp20_005","Col_flg
22_005","Col_ch8_005")

#this puts the matrix in the order I want for the tree - it' important for the pivto_longer step
ordered <- cor.df[match(leaf_order, colnames(cor.df)), match(leaf_order, rownames(cor.df))]

#remove lower triangle
ordered[lower.tri(ordered)]<- NA

#get into the format ggplot2 likes
upper_long <- ordered %>% as.data.frame %>% rownames_to_column(var="sample1") %>% pivot_longer(-sample1,
names_to="sample2", values_to="pearson") %>% na.omit()
```

Get distance matrix for dendrogram

```
#make dendrogram from correlation matrix, not from original data
hc <- hclust(dist(cor.mat))
den <- as.dendrogram(hc)

ordered_den <- den %>% set("branches_lwd", 0.3) %>% rotate(leaf_order)

denplot <- as.ggdend(ordered_den)
```

Make a set of icons

```
definedvibrant <- c("0"="white", "flg22"="#0077BB", "elf18"="#33BBEE", "nlp20"="#009988", "Pep1"="#EE773
3", "OGs"="#CC3311", "ch8"="#EE3377", "LPS"="#DDCC77", "mock"="#BBBBBB")

icons <- data.frame(sample=leaf_order)
icons$split <- icons$sample
icons <- icons %>% separate(split, into=c("Genotype", "Elicitor", "Time"), sep="_")
```

```
map <- ggplot(upper_long, aes(x=factor(sample1, levels=leaf_order), y=factor(sample2, levels=leaf_orde
r), fill=pearson))+
  geom_tile(color="grey")+coord_fixed()+scale_fill_viridis_c(limits=c(-1,1))+
  theme(axis.text=element_blank(), axis.ticks=element_blank(), axis.title=element_blank(),
        panel.background = element_blank(), plot.background = element_blank(),
        legend.position="none",plot.margin = margin(0, 0, 0,0, "pt"))+
  scale_x_discrete(expand=c(0,0))+scale_y_discrete(expand=c(0,0))

simplemap <- ggplot(upper_long, aes(x=factor(sample1, levels=leaf_order), y=factor(sample2, levels=leaf_
order), fill=pearson))+
  geom_tile(color="grey")+coord_fixed()+scale_fill_viridis_c(limits=c(-1,1), guide=guide_colorbar(barwid
th=0.3, barheight=2))+
  theme(legend.key.size=unit(0.01,"cm"), legend.position="top")+
  theme_minimal(base_size=6)+
  labs(fill=expression("Pearson\ncorrelation"))



suppressWarnings(maplegend <- get_legend(simplemap))

iplot <- ggplot(icons, aes(x=1,y=factor(sample, levels=leaf_order),
                           shape=Time, color=Elicitor))+
  geom_point(size=0.3, stroke=0.4)+scale_shape_manual(values=c(0,4,2,3,5))+scale_color_manual(values=def
inedvibrant)+
  theme_classic()+scale_x_continuous(limits=c(1,1), expand=c(0,0))+scale_y_discrete(expand=c(0.01,0.01))
+
  theme(axis.text=element_blank(), axis.title=element_blank(), axis.line=element_blank(), axis.ticks = e
lement_blank(),
        legend.position="none", panel.background=element_rect(fill="transparent", color=NA), plot.backgr
ound=element_rect(fill="transparent", color=NA),
        panel.border=element_blank())

tree <- ggplot(denplot, horiz=T, labels=F)+
  theme(plot.margin = margin(0, 0, 0,0, "pt") )+
  scale_x_continuous(expand=c(0.01,0.01))+scale_y_reverse(expand=c(0,0)) +
  theme(rect=element_rect(fill="transparent", color=NA),
        panel.border=element_blank())
```

```
## Scale for 'y' is already present. Adding another scale for 'y', which will
## replace the existing scale.
```

```
corheatmap <- ggdraw()+
  draw_plot(tree, x=0.01, y=-0.1, width=0.3, height=1.05)+
  draw_plot(iplot, x=0.268, width=0.1, y=-0.03, height=1.025)+
  draw_plot(map, x=0.095, width=0.93, y=0.015, height=0.93)+
  draw_plot(maplegend, x=0.7, width=0.01, y=0.3, height=0.01)

suppressWarnings(ggsave(corheatmap, filename="PanelD.pdf", width=8, height=4, units="cm"))
```

### Figure S2 & Table S4

```
knitr::opts_chunk$set(echo = TRUE)
library(tidyverse)
```

```
## -- Attaching packages ---------------------------------------------------------------------------------------------------------------------- tidyverse 1.3.0 --
```

```
## v ggplot2 3.3.2     v purrr   0.3.4
## v tibble  3.0.3     v dplyr   1.0.2
## v tidyr   1.1.2     v stringr 1.4.0
## v readr   1.3.1     v forcats 0.5.0
```

```
## -- Conflicts ------------------------------------------------------------------------------------------------------------------- tidyverse_conflicts() --
## x dplyr::filter() masks stats::filter()
## x dplyr::lag()    masks stats::lag()
```

### Panel A

elicitor-specific gene expression, starting from counts

```
home="D:/OneDrive/Norwich_Z/Overall/CleanedData/"

#For counts, already have tidy format data, just read it in
ncounts <- read.csv(paste0(home,"wtonly/Cleaned_normalized_counts_long_unfiltered.csv"), header=T, row.names=1)
ncounts$rep <- as.factor(as.character(ncounts$rep))
ncounts$PAMP <- as.character(ncounts$PAMP)
ncounts$PAMP[ncounts$PAMP=="3-OH10"] <- "3-OH-FA"
ncounts$PAMP[ncounts$PAMP=="chitooct"] <- "CO8"
ncounts$FullName <- NULL
ncounts$PAMP <-  factor(ncounts$PAMP, levels=c("flg22","elf18","Pep1","nlp20","OGs","CO8","3-OH-FA","mock"))

#average all reps, then collapse all time points, get total counts induced by elicitor
ncounts <- ncounts %>% select(-Genotype) %>% filter(PAMP != "mock") %>% group_by(Gene, PAMP,Time) %>% summarize(meanCounts=mean(normCounts)) %>% ungroup() %>% group_by(Gene, PAMP) %>% summarize(TotalCounts=sum(meanCounts))
```

```
## `summarise()` regrouping output by 'Gene', 'PAMP' (override with `.groups` argument)
```

```
## `summarise()` regrouping output by 'Gene' (override with `.groups` argument)
```

```
#calculate specificity measure:  SPM
spm <- function(counts){
  total <- sum(counts)
  spm <- counts/total
  return(spm)
}

cpm <- ncounts %>% group_by(Gene) %>% mutate(PAMP=PAMP, spm=spm(TotalCounts))


#filter to genes also called significantly induced in at least one condition - get rid of effects from o
utliers in single reps, genes basically not expressed, etc
fc <- read.csv(paste0(home,"CleanedData_p0.1_l2FCabs1_ordered.csv"), header=T, row.names=1)

genesup <- fc %>% rownames_to_column("Gene") %>% pivot_longer(-Gene, names_to="FullName", values_to="log
2FC") %>% separate( FullName, into=c("PAMP", "Time"), sep="_") %>% mutate(log2FC=ifelse(log2FC<0,0, log2
FC)) %>% group_by(Gene, PAMP)  %>% summarize(TotalInduction=sum(log2FC)) %>% ungroup() %>% group_by(Gen
e) %>% filter(TotalInduction>0)
```

```
## `summarise()` regrouping output by 'Gene' (override with `.groups` argument)
```

```
filtered <- cpm[cpm$Gene %in% genesup$Gene,]
```

make filter for specificity, then get genes specifically enriched in each elicitor (one-by-one, then recombine)

```
EnrichedGenes <- filtered %>% group_by(Gene) %>% summarize(maxenrich=max(spm)) %>% filter(maxenrich>0.33
)
```

```
## `summarise()` ungrouping output (override with `.groups` argument)
```

```
filteredenriched <- filtered[filtered$Gene %in% EnrichedGenes$Gene,]

filteredwide <- filteredenriched %>% select(-TotalCounts) %>% pivot_wider(names_from=PAMP, values_from=s
pm)

#split to arrange genes in each elicitor according to behaviour across all elicitors, assign Genes to th
e elicitor in which they are most enriched
flg <- filteredwide[max.col(filteredwide[2:8], ties.method="last")==1,] %>% arrange(desc(`flg22`)) %>% c
olumn_to_rownames("Gene")
elf <- filteredwide[max.col(filteredwide[2:8], ties.method="last")==2,] %>% arrange(desc(`elf18`)) %>% c
olumn_to_rownames("Gene")
pep <- filteredwide[max.col(filteredwide[2:8], ties.method="last")==3,] %>% arrange(desc(`Pep1`)) %>% co
lumn_to_rownames("Gene")
nlp <- filteredwide[max.col(filteredwide[2:8], ties.method="last")==4,] %>% arrange(desc(`nlp20`)) %>% c
olumn_to_rownames("Gene")
OGs <- filteredwide[max.col(filteredwide[2:8], ties.method="last")==5,] %>% arrange(desc(`OGs`)) %>% col
umn_to_rownames("Gene")
CO8 <- filteredwide[max.col(filteredwide[2:8], ties.method="last")==6,] %>% arrange(desc(`CO8`)) %>% col
umn_to_rownames("Gene")
LPS <- filteredwide[max.col(filteredwide[2:8], ties.method="last")==7,] %>% arrange(desc(`3-OH-FA`)) %>%
column_to_rownames("Gene")

nrow(flg)
```

```
## [1] 332
```

```
nrow(elf)
```

```
## [1] 8
```

```
nrow(pep)
```

```
## [1] 33
```

```
nrow(nlp)
```

```
## [1] 31
```

```
nrow(OGs)
```

```
## [1] 8
```

```
nrow(CO8)
```

```
## [1] 0
```

```
nrow(LPS)
```

```
## [1] 0
```

```
#recombine the tables for each time, in order
filteredarranged <- rbind(flg, elf, pep, nlp, OGs, CO8, LPS)

write.csv(filteredarranged, file="TableS4.csv")

#pivot longer to plot with ggplot
ggfiltered <- filteredarranged %>% rownames_to_column("Gene") %>% pivot_longer(-"Gene",names_to="elicito
r",values_to = "spm")

ggfiltered$elicitor <-  factor(ggfiltered$elicitor, levels=c("flg22","elf18","Pep1","nlp20","OGs","CO8",
"3-OH-FA"))
```

```
PanelA <- ggplot(ggfiltered, aes(x=elicitor, y=factor(Gene, levels=rev(unique(Gene))), fill=spm)) + them
e_minimal(base_size=7) +
  geom_tile()+scale_fill_viridis_c(guide=guide_colorbar(barwidth=unit(1.5, "cm"), barheight=unit(0.2, "c
m")), limits=c(0,1))+
  theme(axis.text.y=element_blank(), axis.ticks.y=element_blank(), rect=element_rect(fill="transparent",
color=NA),
        panel.grid.major=element_blank(), legend.position="top", axis.text.x=element_text(angle=45, hjus
t=1, vjust=1),
        plot.margin=unit(c(0.25,0,0.25,-0.25), "cm"))+
  labs(x="", y="", fill="Specificity measure (SPM)")

ggsave("PanelA.pdf", height=10, width=6, units='cm')
```

### Figure S3

Marta Bjornson

```
library(extraDistr)
library(tidyverse)
library(rlang)
library(SuperExactTest)
library(RColorBrewer)
library(ggthemes)
library(ggforce)
require(scales)
require(cowplot)
library(ggpubr)
```

Finding degree, set size, and deviation just the same as Figure 1c. Here will plot not just largest 3 sets, but all possible sets, in a wheel as in https://www.nature.com/articles/srep16923 (https://www.nature.com/articles/srep16923) (note difference in how set size defined). First define key variables

```
rm(list=ls())

path_to_data <- "D:/OneDrive/Norwich_Z/Overall/CleanedData/wtonly/"
home <- "D:/OneDrive/Norwich_Z/Overall/CleanedData/"

definedvibrant <- c("0"="white", "flg22"="#0077BB", "elf18"="#33BBEE", "nlp20"="#009988", "Pep1"="#EE773
3", "OGs"="#CC3311", "CO8"="#EE3377", "3-OH-FA"="#DDCC77", "mock"="#BBBBBB")
```

Define functions

```
#Start from most basic data, but to interpret get it into UpSet format, modified function SingleTimeTabl
e from figure 1
SingleTimeTable <- function(time, direction){
  readtable <- function(elicitor){
    c <- read.csv(paste0(path_to_data, elicitor, "_", time, direction,".csv"), header=F)
    c$present <- 1
    return(c)
  }
  flg <- readtable("flg22")
  elf <- readtable("elf18")
  pep <- readtable("Pep1")
  nlp <- readtable("nlp20")
  OGs <- readtable("OGs")
  CO8 <- readtable("ch8")
  LPS <- readtable("LPS")

  timetable <- merge(flg, elf, by="V1", all=T)
  timetable <- merge(timetable, pep, by="V1", all=T)
  timetable <- merge(timetable, nlp, by="V1", all=T)
  timetable <- merge(timetable, OGs, by="V1", all=T)
  timetable <- merge(timetable, CO8, by="V1", all=T)
  timetable <- merge(timetable, LPS, by="V1", all=T)
  colnames(timetable) <- c("ATG","flg","elf","pep","nlp","OGs","CO8","LPS")
  timetable[is.na(timetable)] <- 0
  timetable <- column_to_rownames(timetable,var="ATG")
  timetable[timetable!=0] <- 1
  timetable <- rownames_to_column(timetable,var="ATG")
  return(timetable)
}

#This function takes a table in UpsetFormat, and makes it into a long table for inter_sets. Could probab
ly combine, but so much going on in that function already I'd rather not
lengthen <- function(upsettable){
  longtable <- upsettable %>% gather(-ATG, key="elicitor",value="value")
  longtable$TF <- FALSE
  longtable$TF[longtable$value==1] <- TRUE

  colnames(longtable) <- c("ATG","group","numeric","value")
  longtable$numeric <- NULL
  return(longtable)
}

#master function to get set sizes and deviation: takes long table format with T/F for gene induced by el
icitor or not
inter_sets = function(time, direction){
#this function parses a text string (A&B&!C) and finds the number of rows that actually meet those crite
ria: taken from StackOverflow with slight alteration
  filter_sets = function(filter_expr){
    long %>%
      spread(group, value) %>%
      filter(!!parse_quosure(filter_expr)) %>%
      nrow()
  }
#This function I wrote: it uses the definition of Deviation from UpSet (expected size of overlap given t
he size of all sets participating and the size of all sets not participating). I checked it against valu
es calculated online and it seemed to be performing as expected
calculate_deviation = function(elicitors){
    use <- colnames(setsizes)[colnames(setsizes) %in% elicitors]
    yes <- setsizes[,use]
    yes <- yes/n
    yes <- prod(yes)

    dont <- colnames(setsizes)[!colnames(setsizes) %in% elicitors]
```

```
    if(length(dont) >0){
      no <- setsizes[,dont]
      no <- no/n
      no <- 1-no
      no <- prod(no)
    }else{
      no <- 1
    }

    product <- prod(yes, no)

    cardinality <- as.numeric(elicitors['value'])/n

    deviation <- cardinality - product
    deviation
}

upset <- SingleTimeTable(time, direction)
long <- lengthen(upset)

(setsizes <- data.frame('flg'=nrow(upset[upset$flg==1,]), 'CO8'=nrow(upset[upset$CO8==1,]), 'elf'=nrow(u
pset[upset$elf==1,]),'LPS'=nrow(upset[upset$LPS==1,]), 'nlp'=nrow(upset[upset$nlp==1,]), 'OGs'=nrow(upse
t[upset$OGs==1,]), 'pep'=nrow(upset[upset$pep==1,]))) #number induced by each individually regardless of
overlaps
n <- nrow(upset) #total numberinduced

#this is the second part of the code taken from StackOverflow - this generates all possible combinations
of A&B&C, A&B&!C, A&!B&C, etc, as strings, then reduces them to non-redundant set
  combins = unique(long$group) %>%
    c(paste0("!", .)) %>%
    combn(length(.)/2) %>%
    t() %>%
    as.data.frame() %>%
    filter(apply(., 1, function(x) length(unique(gsub("!", "", x))) == ncol(.) & !(length(grep("!", x))
 %in% c(0, ncol(.))))) %>%
    unite("expressions", names(.), sep = " & ")
  #I added this, it is the all elicitor set
    combins <- rbind(combins, paste(unique(long$group), sep = ' & ', collapse = ' & '))
  #I had added this for an earlier version, it's the no elicitor set. Could be useful again depending on
what I want to look at
    #combins <- rbind(combins, paste(paste0("!",unique(long$group)), sep = ' & ', collapse = ' & '))

#Here add the size of any interaction to the string describing that interaction
  combins$value = sapply(combins$expressions, filter_sets)
  combins$value <- as.numeric(combins$value)

#For my code need elicitors each in own column
  separated <- combins %>% separate(expressions, into = c('a','b','c','d','e','f','g'), sep=' & ')

# Calculate deviation, given binary presence/absence of elicitor and size of overlap set
  separated$deviation <- apply(separated, 1, calculate_deviation)
#Make recombined column with set name all together
  recombined <- separated %>% unite(expression, a,b,c,d,e,f,g)

  return(recombined)
}

#for a given elicitor, make a binary presence/absence column
elicitor_columns=function(deviationdata, elicitor){
  deviationdata$new <- 1
  deviationdata$new[grep(paste0("!",elicitor),deviationdata$expression)] <- 0
  colnames(deviationdata)[colnames(deviationdata)=="new"] <- elicitor
  return(deviationdata)
}
```

Perform on all interesting datasets

```
#define a function which does all steps, just so I can call it multiple times for multiple irises with d
ocumentation of each
wrapper <- function(time, direction, palette, name){
  doit <- inter_sets(time,direction) #get all intersections, deviations
  doit <- elicitor_columns(doit, "flg")
  doit <- elicitor_columns(doit, "elf")
  doit <- elicitor_columns(doit, "nlp")
  doit <- elicitor_columns(doit, "pep")
  doit <- elicitor_columns(doit, "OGs")
  doit <- elicitor_columns(doit, "CO8")
  doit <- elicitor_columns(doit, "LPS")
#to make plots with ggforce, need start and stop agle of each "chunk" of pie, so first arrange as I lik
e, then calculate and add
  doit$degree <- rowSums(doit[4:10])
  doit <- arrange(doit, desc(degree))
  doit$start_angle <- seq(0,2*pi-(2*pi/127), length.out=127)
  doit$end_angle <- seq(2*pi/127, 2*pi, length.out=127)
#this is mostly just for making the fills different colors, there's probably a better way to do it
  doit$flg[doit$flg==1] <- "flg22"
  doit$elf[doit$elf==1] <- "elf18"
  doit$nlp[doit$nlp==1] <- "nlp20"
  doit$pep[doit$pep==1] <- "Pep1"
  doit$OGs[doit$OGs==1] <- "OGs"
  doit$CO8[doit$CO8==1] <- "CO8"
  doit$LPS[doit$LPS==1] <- "3-OH-FA"
  write.csv(doit, file =paste0(name,"_table.csv"))

  tracks <- ggplot(doit) + theme_no_axes() + coord_fixed() +
    geom_arc_bar(aes(x0 = 0, y0 = 0, r0 = 1, r = 1.7, amount=1, fill = as.factor(flg)),
                 stat = 'pie', color="#DDDDDD")  +
    geom_arc_bar(aes(x0 = 0, y0 = 0, r0 = 1.7, r = 2.3, amount = 1, fill = as.factor(elf)),
                 stat = 'pie', color="#DDDDDD")  +
    geom_arc_bar(aes(x0 = 0, y0 = 0, r0 = 2.3, r = 2.8, amount = 1, fill = as.factor(pep)),
                 stat = 'pie', color="#DDDDDD")  +
    geom_arc_bar(aes(x0 = 0, y0 = 0, r0 = 2.8, r = 3.25, amount = 1, fill = as.factor(nlp)),
                 stat = 'pie', color="#DDDDDD")  +
    geom_arc_bar(aes(x0 = 0, y0 = 0, r0 = 3.25, r = 3.65, amount = 1, fill = as.factor(OGs)),
                 stat = 'pie', color="#DDDDDD")  +
    geom_arc_bar(aes(x0 = 0, y0 = 0, r0 = 3.65, r = 4.05, amount = 1, fill = as.factor(CO8)),
                 stat = 'pie', color="#DDDDDD") +
    geom_arc_bar(aes(x0 = 0, y0 = 0, r0 = 4.05, r = 4.45, amount = 1, fill = as.factor(LPS)),
                 stat = 'pie', color="#DDDDDD") +
    scale_fill_manual(values=palette)+
    theme(plot.margin = margin(0, 0, 0,0, "pt"), panel.grid=element_blank(),
          panel.background=element_rect(fill="transparent"),panel.border=element_blank(),
          plot.background=element_rect(fill="transparent"), legend.position="none") +
    labs(x = NULL, y = NULL, legend=NULL)

#can have each iris on its own scale, that what the commented line would do, but for now keeping all on
 same scale
#offset=max(doit$value)
  offset=1000

  arcbars <- ggplot(doit) + theme_no_axes() +coord_fixed() +
    geom_arc_bar(aes(x0 = 0, y0 = 0, r0 = offset, r = value+offset, fill = deviation,
                     start=start_angle, end=end_angle)) +
    scale_y_continuous(limits=c(-2*offset, 2*offset), expand=c(0,0)) +
    scale_x_continuous(limits=c(-2*offset, 2*offset), expand=c(0,0)) +
    theme(legend.position=c(0.85, 0.98), legend.background = element_blank(),
          legend.box.background = element_blank(), legend.box=NULL, legend.direction="horizontal",
          panel.border=element_blank(), panel.background=element_rect(fill="transparent"),
          plot.background=element_rect(fill="transparent"), panel.grid = element_blank(),
          plot.margin = margin(0, 0, 0,0, "pt")) +
```

```
    scale_fill_distiller(palette="BrBG", direction=1, limits=c(-0.25, 0.25))



  result <-  ggdraw() +draw_plot(tracks, x=0.225, y=0.225, width=0.55, height=0.55) +draw_plot(arcbars,
 x=0, y=0, width=1, height=1)
  ggsave(paste0(name, "_plot.pdf"), plot=result, height=7, width=7)

}

wrapper("collapsed","up",definedvibrant,"2020-10-04/Allup")
```

```
## Warning in merge.data.frame(timetable, nlp, by = "V1", all = T): column names
## 'present.x', 'present.y' are duplicated in the result
```

```
## Warning in merge.data.frame(timetable, OGs, by = "V1", all = T): column names
## 'present.x', 'present.y' are duplicated in the result
```

```
## Warning in merge.data.frame(timetable, CO8, by = "V1", all = T): column names
## 'present.x', 'present.y', 'present.x', 'present.y' are duplicated in the result
```

```
## Warning in merge.data.frame(timetable, LPS, by = "V1", all = T): column names
## 'present.x', 'present.y', 'present.x', 'present.y' are duplicated in the result
```

```
## Warning: `parse_quosure()` is deprecated as of rlang 0.2.0.
## Please use `parse_quo()` instead.
## This warning is displayed once per session.
```

```
wrapper("005","up",definedvibrant,"2020-10-04/005up")
```

```
## Warning in merge.data.frame(timetable, nlp, by = "V1", all = T): column names
## 'present.x', 'present.y' are duplicated in the result
```

```
## Warning in merge.data.frame(timetable, OGs, by = "V1", all = T): column names
## 'present.x', 'present.y' are duplicated in the result
```

```
## Warning in merge.data.frame(timetable, CO8, by = "V1", all = T): column names
## 'present.x', 'present.y', 'present.x', 'present.y' are duplicated in the result
```

```
## Warning in merge.data.frame(timetable, LPS, by = "V1", all = T): column names
## 'present.x', 'present.y', 'present.x', 'present.y' are duplicated in the result
```

```
wrapper("010","up",definedvibrant,"2020-10-04/010up")
```

```
## Warning in merge.data.frame(timetable, nlp, by = "V1", all = T): column names
## 'present.x', 'present.y' are duplicated in the result
```

```
## Warning in merge.data.frame(timetable, OGs, by = "V1", all = T): column names
## 'present.x', 'present.y' are duplicated in the result
```

```
## Warning in merge.data.frame(timetable, CO8, by = "V1", all = T): column names
## 'present.x', 'present.y', 'present.x', 'present.y' are duplicated in the result
```

```
## Warning in merge.data.frame(timetable, LPS, by = "V1", all = T): column names
## 'present.x', 'present.y', 'present.x', 'present.y' are duplicated in the result
```

```
wrapper("030","up",definedvibrant,"2020-10-04/030up")
```

```
## Warning in merge.data.frame(timetable, nlp, by = "V1", all = T): column names
## 'present.x', 'present.y' are duplicated in the result
```

```
## Warning in merge.data.frame(timetable, OGs, by = "V1", all = T): column names
## 'present.x', 'present.y' are duplicated in the result
```

```
## Warning in merge.data.frame(timetable, CO8, by = "V1", all = T): column names
## 'present.x', 'present.y', 'present.x', 'present.y' are duplicated in the result
```

```
## Warning in merge.data.frame(timetable, LPS, by = "V1", all = T): column names
## 'present.x', 'present.y', 'present.x', 'present.y' are duplicated in the result
```

```
wrapper("090","up",definedvibrant,"2020-10-04/090up")
```

```
## Warning in merge.data.frame(timetable, nlp, by = "V1", all = T): column names
## 'present.x', 'present.y' are duplicated in the result
```

```
## Warning in merge.data.frame(timetable, OGs, by = "V1", all = T): column names
## 'present.x', 'present.y' are duplicated in the result
```

```
## Warning in merge.data.frame(timetable, CO8, by = "V1", all = T): column names
## 'present.x', 'present.y', 'present.x', 'present.y' are duplicated in the result
```

```
## Warning in merge.data.frame(timetable, LPS, by = "V1", all = T): column names
## 'present.x', 'present.y', 'present.x', 'present.y' are duplicated in the result
```

```
wrapper("180","up",definedvibrant,"2020-10-04/180up")
```

```
## Warning in merge.data.frame(timetable, nlp, by = "V1", all = T): column names
## 'present.x', 'present.y' are duplicated in the result
```

```
## Warning in merge.data.frame(timetable, OGs, by = "V1", all = T): column names
## 'present.x', 'present.y' are duplicated in the result
```

```
## Warning in merge.data.frame(timetable, CO8, by = "V1", all = T): column names
## 'present.x', 'present.y', 'present.x', 'present.y' are duplicated in the result
```

```
## Warning in merge.data.frame(timetable, LPS, by = "V1", all = T): column names
## 'present.x', 'present.y', 'present.x', 'present.y' are duplicated in the result
```

```
wrapper("collapsed","down",definedvibrant,"2020-10-04/Alldown")
```

```
## Warning in merge.data.frame(timetable, nlp, by = "V1", all = T): column names
## 'present.x', 'present.y' are duplicated in the result
```

```
## Warning in merge.data.frame(timetable, OGs, by = "V1", all = T): column names
## 'present.x', 'present.y' are duplicated in the result
```

```
## Warning in merge.data.frame(timetable, CO8, by = "V1", all = T): column names
## 'present.x', 'present.y', 'present.x', 'present.y' are duplicated in the result
```

```
## Warning in merge.data.frame(timetable, LPS, by = "V1", all = T): column names
## 'present.x', 'present.y', 'present.x', 'present.y' are duplicated in the result
```

```
wrapper("010","down",definedvibrant,"2020-10-04/010down")
```

```
## Warning in merge.data.frame(timetable, nlp, by = "V1", all = T): column names
## 'present.x', 'present.y' are duplicated in the result
```

```
## Warning in merge.data.frame(timetable, OGs, by = "V1", all = T): column names
## 'present.x', 'present.y' are duplicated in the result
```

```
## Warning in merge.data.frame(timetable, CO8, by = "V1", all = T): column names
## 'present.x', 'present.y', 'present.x', 'present.y' are duplicated in the result
```

```
## Warning in merge.data.frame(timetable, LPS, by = "V1", all = T): column names
## 'present.x', 'present.y', 'present.x', 'present.y' are duplicated in the result
```

```
wrapper("030","down",definedvibrant,"2020-10-04/030down")
```

```
## Warning in merge.data.frame(timetable, nlp, by = "V1", all = T): column names
## 'present.x', 'present.y' are duplicated in the result
```

```
## Warning in merge.data.frame(timetable, OGs, by = "V1", all = T): column names
## 'present.x', 'present.y' are duplicated in the result
```

```
## Warning in merge.data.frame(timetable, CO8, by = "V1", all = T): column names
## 'present.x', 'present.y', 'present.x', 'present.y' are duplicated in the result
```

```
## Warning in merge.data.frame(timetable, LPS, by = "V1", all = T): column names
## 'present.x', 'present.y', 'present.x', 'present.y' are duplicated in the result
```

```
wrapper("090","down",definedvibrant,"2020-10-04/090down")
```

```
## Warning in merge.data.frame(timetable, nlp, by = "V1", all = T): column names
## 'present.x', 'present.y' are duplicated in the result
```

```
## Warning in merge.data.frame(timetable, OGs, by = "V1", all = T): column names
## 'present.x', 'present.y' are duplicated in the result
```

```
## Warning in merge.data.frame(timetable, CO8, by = "V1", all = T): column names
## 'present.x', 'present.y', 'present.x', 'present.y' are duplicated in the result
```

```
## Warning in merge.data.frame(timetable, LPS, by = "V1", all = T): column names
## 'present.x', 'present.y', 'present.x', 'present.y' are duplicated in the result
```

```
wrapper("180","down",definedvibrant,"2020-10-04/180down")
```

```
## Warning in merge.data.frame(timetable, nlp, by = "V1", all = T): column names
## 'present.x', 'present.y' are duplicated in the result
```

```
## Warning in merge.data.frame(timetable, OGs, by = "V1", all = T): column names
## 'present.x', 'present.y' are duplicated in the result
```

```
## Warning in merge.data.frame(timetable, CO8, by = "V1", all = T): column names
## 'present.x', 'present.y', 'present.x', 'present.y' are duplicated in the result
```

```
## Warning in merge.data.frame(timetable, LPS, by = "V1", all = T): column names
## 'present.x', 'present.y', 'present.x', 'present.y' are duplicated in the result
```

### Figure S4

Marta Bjornson

07 April 2020

```
knitr::opts_chunk$set(echo = TRUE)
library(tidyverse)
library(cowplot)
library(ggpubr) # for as_ggplot
library(ggforce) #for theme_no_axes
library(dendextend)
library(viridis)
library(rlang) #for parse_quosure (deprecated, but parse_quo doesn't work the same and I haven't update
d)
library(bioDist)#for spearman distance
```

### Panel A

As Figure 1C Step one: extract all intersections (of all degrees). Trying a chunk of code from stack overflow by useR: https://stackoverflow.com/questions/47083311/r-extract-sets-in-arbitrary-intersection-from-a-data-frame (https://stackoverflow.com/questions/47083311/r-extract-sets-in-arbitrary-intersection-from-a-data-frame) First define all functions

```r
rm(list=ls())
path_to_data <- "D:/OneDrive/Norwich_Z/Overall/CleanedData/wtonly/"
home <- "D:/OneDrive/Norwich_Z/Overall/CleanedData/"
definedvibrant <- c("0"="white", "flg22"="#0077BB", "elf18"="#33BBEE", "Pep1"="#009988", "nlp20"="#EE7733", "OGs"="#CC3311", "CO8"="#EE3377", "3-OH-FA"="#DDCC77", "mock"="#BBBBBB")


#Start from most basic data, but to interpret get it into UpSet format, modified function SingleTimeTable from figure 1
SingleTimeTable <- function(time, direction){
  readtable <- function(elicitor){
    c <- read.csv(paste0(path_to_data, elicitor, "_", time, direction,".csv"), header=F)
    c$present <- 1
    return(c)
  }
  flg <- readtable("flg22")
  elf <- readtable("elf18")
  pep <- readtable("Pep1")
  nlp <- readtable("nlp20")
  OGs <- readtable("OGs")
  CO8 <- readtable("ch8")
  LPS <- readtable("LPS")

  timetable <- merge(flg, elf, by="V1", all=T)
  timetable <- merge(timetable, pep, by="V1", all=T)
  timetable <- merge(timetable, nlp, by="V1", all=T)
  timetable <- merge(timetable, OGs, by="V1", all=T)
  timetable <- merge(timetable, CO8, by="V1", all=T)
  timetable <- merge(timetable, LPS, by="V1", all=T)
  colnames(timetable) <- c("ATG","flg","elf","pep","nlp","OGs","CO8","LPS")
  timetable[is.na(timetable)] <- 0
  timetable <- column_to_rownames(timetable,var="ATG")
  timetable[timetable!=0] <- 1
  timetable <- rownames_to_column(timetable,var="ATG")
  return(timetable)
}

#This function takes a table in UpsetFormat, and makes it into a long table for inter_sets. Could probably combine, but so much going on in that function already I'd rather not
lengthen <- function(upsettable){
  longtable <- upsettable %>% gather(-ATG, key="elicitor",value="value")
  longtable$TF <- FALSE
  longtable$TF[longtable$value==1] <- TRUE

  colnames(longtable) <- c("ATG","group","numeric","value")
  longtable$numeric <- NULL
  return(longtable)
}

#master function to get set sizes and deviation: takes long table format with T/F for gene induced by elicitor or not
inter_sets = function(time, direction){
#this function parses a text string (A&B&!C) and finds the number of rows that actually meet those criteria: taken from StackOverflow with slight alteration
  filter_sets = function(filter_expr){
    long %>%
      spread(group, value) %>%
      filter(!!parse_quosure(filter_expr)) %>%
      nrow()
  }
#This function I wrote: it uses the definition of Deviation from UpSet (expected size of overlap given the size of all sets participating and the size of all sets not participating). I checked it against values calculated online and it seemed to be performing as expected
```

```
calculate_deviation = function(elicitors){
    use <- colnames(setsizes)[colnames(setsizes) %in% elicitors]
    yes <- setsizes[,use]
    yes <- yes/n
    yes <- prod(yes)

    dont <- colnames(setsizes)[!colnames(setsizes) %in% elicitors]
    if(length(dont) >0){
      no <- setsizes[,dont]
      no <- no/n
      no <- 1-no
      no <- prod(no)
    }else{
      no <- 1
    }

    product <- prod(yes, no)

    cardinality <- as.numeric(elicitors['value'])/n

    deviation <- cardinality - product
    deviation
}

upset <- SingleTimeTable(time, direction)
long <- lengthen(upset)

(setsizes <- data.frame('flg'=nrow(upset[upset$flg==1,]), 'CO8'=nrow(upset[upset$CO8==1,]), 'elf'=nrow(u
pset[upset$elf==1,]),'LPS'=nrow(upset[upset$LPS==1,]), 'nlp'=nrow(upset[upset$nlp==1,]), 'OGs'=nrow(upse
t[upset$OGs==1,]), 'pep'=nrow(upset[upset$pep==1,]))) #number induced by each individually regardless of
overlaps
n <- nrow(upset) #total numberinduced

#this is the second part of the code taken from StackOverflow - this generates all possible combinations
of A&B&C, A&B&!C, A&!B&C, etc, as strings, then reduces them to non-redundant set
  combins = unique(long$group) %>%
    c(paste0("!", .)) %>%
    combn(length(.)/2) %>%
    t() %>%
    as.data.frame() %>%
    filter(apply(., 1, function(x) length(unique(gsub("!", "", x))) == ncol(.) & !(length(grep("!", x))
 %in% c(0, ncol(.))))) %>%
    unite("expressions", names(.), sep = " & ")
  #I added this, it is the all elicitor set
    combins <- rbind(combins, paste(unique(long$group), sep = ' & ', collapse = ' & '))
  #I had added this for an earlier version, it's the no elicitor set. Could be useful again depending on
what I want to look at
    #combins <- rbind(combins, paste(paste0("!",unique(long$group)), sep = ' & ', collapse = ' & '))

#Here add the size of any interaction to the string describing that interaction
  combins$value = sapply(combins$expressions, filter_sets)
  combins$value <- as.numeric(combins$value)

#For my code need elicitors each in own column
  separated <- combins %>% separate(expressions, into = c('a','b','c','d','e','f','g'), sep=' & ')

# Calculate deviation, given binary presence/absence of elicitor and size of overlap set
  separated$deviation <- apply(separated, 1, calculate_deviation)
#Make recombined column with set name all together
  recombined <- separated %>% unite(expression, a,b,c,d,e,f,g)

  return(recombined)
}
```

```
#for a given elicitor, make a binary presence/absence column
elicitor_columns=function(deviationdata, elicitor){
  deviationdata$new <- 1
  deviationdata$new[grep(paste0("!",elicitor),deviationdata$expression)] <- 0
  colnames(deviationdata)[colnames(deviationdata)=="new"] <- elicitor
  return(deviationdata)
}
```

Then generate data: first for all times collapsed together

```
#start with upregulated genes
doit <- inter_sets("Collapsed","down") #get all intersections, deviations
```

```
## Warning in merge.data.frame(timetable, nlp, by = "V1", all = T): column names
## 'present.x', 'present.y' are duplicated in the result
```

```
## Warning in merge.data.frame(timetable, OGs, by = "V1", all = T): column names
## 'present.x', 'present.y' are duplicated in the result
```

```
## Warning in merge.data.frame(timetable, CO8, by = "V1", all = T): column names
## 'present.x', 'present.y', 'present.x', 'present.y' are duplicated in the result
```

```
## Warning in merge.data.frame(timetable, LPS, by = "V1", all = T): column names
## 'present.x', 'present.y', 'present.x', 'present.y' are duplicated in the result
```

```
## Warning: `parse_quosure()` is deprecated as of rlang 0.2.0.
## Please use `parse_quo()` instead.
## This warning is displayed once per session.
```

```
doit <- elicitor_columns(doit, "flg") #then make a column for each elicitor: 1 if included in set, 0 if
 not
doit <- elicitor_columns(doit, "elf")
doit <- elicitor_columns(doit, "nlp")
doit <- elicitor_columns(doit, "pep")
doit <- elicitor_columns(doit, "OGs")
doit <- elicitor_columns(doit, "CO8")
doit <- elicitor_columns(doit, "LPS")
doit$degree <- rowSums(doit[4:10])#calculate the degree (shouldn't need this?)
doit <- arrange(doit, desc(value))
doit$expression <- factor(doit$expression, ordered=T)#this should allow me to control the arrangment of
 the x axis but it doesn't work
#This was for the iris setup: keep it because it's still useful for controlling the arrangement of the x
axis
doit$start_angle <- seq(0,2*pi-(2*pi/127), length.out=127)
doit$end_angle <- seq(2*pi/127, 2*pi, length.out=127)
#this is mostly just for making the fills different colors, there's probably a better way to do it
doit$flg[doit$flg==1] <- "flg22"
doit$elf[doit$elf==1] <- "elf18"
doit$nlp[doit$nlp==1] <- "nlp20"
doit$pep[doit$pep==1] <- "Pep1"
doit$OGs[doit$OGs==1] <- "OGs"
doit$CO8[doit$CO8==1] <- "CO8"
doit$LPS[doit$LPS==1] <- "3-OH-FA"
```

```
#make plots for vertical set sizes bars
vertical <- ggplot(doit[1:15,], aes(x=as.factor(start_angle), y=value, fill=deviation))+
  geom_bar(stat="identity", color="grey", size=0.15)+
  theme_classic(base_size=6)+
  theme(axis.title.x=element_blank(), axis.text.x=element_blank(),legend.position="none",
        rect=element_rect(fill="transparent",color=NA), plot.background = element_blank())+
  scale_y_continuous(expand=c(0,0), limits=c(0, 1100), name="Genes in set")+
  scale_fill_distiller(palette="BrBG", direction=1, limits=c(-0.2, 0.2))

vert <- ggplot(doit[1:15,], aes(x=as.factor(start_angle), y=value, fill=deviation))+
  geom_bar(stat="identity")+theme_minimal(base_size=6)+
  scale_fill_distiller(palette="BrBG", direction=1, limits=c(-0.2, 0.2), guide=guide_colorbar(barheight=
0.3))+
  theme(legend.position = "top", legend.key.size=unit(0.25,"cm"))+labs(fill=expression("Deviation\n"))
deviat <- as_ggplot(get_legend(vert))


#make plot for horizontal elicitor sizes bars
elicitorsizes <- SingleTimeTable("collapsed", "down")
```

```
## Warning in merge.data.frame(timetable, nlp, by = "V1", all = T): column names
## 'present.x', 'present.y' are duplicated in the result
```

```
## Warning in merge.data.frame(timetable, OGs, by = "V1", all = T): column names
## 'present.x', 'present.y' are duplicated in the result
```

```
## Warning in merge.data.frame(timetable, CO8, by = "V1", all = T): column names
## 'present.x', 'present.y', 'present.x', 'present.y' are duplicated in the result
```

```
## Warning in merge.data.frame(timetable, LPS, by = "V1", all = T): column names
## 'present.x', 'present.y', 'present.x', 'present.y' are duplicated in the result
```

```
setsizes <- data.frame('flg22'=nrow(elicitorsizes[elicitorsizes$flg==1,]), 'CO8'=nrow(elicitorsizes[elic
itorsizes$CO8==1,]), 'elf18'=nrow(elicitorsizes[elicitorsizes$elf==1,]),'3-OH-FA'=nrow(elicitorsizes[eli
citorsizes$LPS==1,]), 'nlp20'=nrow(elicitorsizes[elicitorsizes$nlp==1,]), 'OGs'=nrow(elicitorsizes[elici
torsizes$OGs==1,]), 'Pep1'=nrow(elicitorsizes[elicitorsizes$pep==1,])) #number induced by each individua
lly regardless of overlaps
setsizes <- setsizes %>% pivot_longer(everything(),names_to="elicitor", values_to="ngenes")
setsizes$elicitor <- gsub("X3.OH.FA","3-OH-FA",setsizes$elicitor)
setsizes$elicitor <- factor(setsizes$elicitor,
                            levels=c("flg22","elf18","Pep1","nlp20","OGs","CO8","3-OH-FA"))
horizontal <- ggplot(setsizes, aes(x=ngenes/1000, y=reorder(elicitor, -ngenes), fill=elicitor))+
  geom_bar(stat="identity")+
  theme_classic(base_size=6)+
  theme(axis.title.y=element_blank(), axis.text.y = element_blank(), legend.position="none",
        axis.line.y=element_blank(), axis.ticks.y=element_blank(),panel.background=element_rect(fill="tr
ansparent"),
        plot.background=element_rect(fill="transparent"), plot.margin = margin(0, 0, 0,0, "pt"))+
  scale_x_reverse(expand=c(0,0), limit=c(5,0), name="Genes repressed (x1000)")+
  scale_fill_manual(values=definedvibrant)
#+scale_y_discrete(position="right", labels=c("   flg22","   elf18","   Pep1","   nlp20","    OGs","
 CO8","3-OH-FA"))

horiz <- ggplot(setsizes, aes(x=ngenes/1000, y=reorder(elicitor, -ngenes), fill=elicitor))+
  geom_bar(stat="identity")+labs(fill="Pattern")+
  scale_fill_manual(values=definedvibrant)+
  theme_minimal(base_size=6)+
  theme(legend.key.size=unit(0.25,"cm"))
elic <- get_legend(horiz)

#make plots for which elicitor sets dots
doit_long <- doit %>% pivot_longer(c(flg, elf, nlp,pep,OGs, CO8, LPS), names_to="elicitor", values_to="p
resent")
doit_long$elicitor <- factor(doit_long$elicitor, levels=c("flg","elf","pep","nlp","OGs","CO8","LPS"))
dots <- ggplot(doit_long[1:105,], aes(x=start_angle, y=elicitor, fill=as.factor(present)))+
  geom_point(shape=21, size=2)+
  scale_fill_manual(values=definedvibrant)+
  theme_no_axes()+
  theme(panel.background = element_blank(), plot.background = element_blank(), panel.border=element_blan
k(),
        legend.position="none", plot.margin = margin(0, 0, 0,0, "pt"))



#Combine all the bits
upset <- ggdraw()+
  draw_plot(vertical,x=0.235, y=0.38, width=0.78, height=0.62)+
  draw_plot(horizontal, x=0.02, y=0.005, width=0.34, height=0.385)+
  draw_plot(dots, x=0.355, y=0.075, height=0.32, width=0.645)+
  draw_plot(deviat, x=0.65, y=0.85, height=0.1, width=0.2)+
  draw_plot(elic, x=0.03, y=0.55, height=0.3, width=0.2)

suppressWarnings(ggsave(upset, filename = "PanelA.pdf", height=6, width=6, units="cm", bg="transparent"
))
```

### PanelB

Clustering of core downregulated genes

```
wt_sig_down <- function(wt, mut){
  #first find the genes upregulatd in mutant
  mut.file=paste0(home,"ResultsObjects/",mut,"_results")
  load(mut.file)
  mut.df <- as.data.frame(res)

  mut.up <- mut.df %>% filter(padj<0.05)  %>%  filter(log2FoldChange < -1) %>% rownames_to_column("ATG")

  #then the genes upregulated in  wt, extra step filter out the ones also upregulated in control
  wt.file=paste0(home,"ResultsObjects/",wt,"_results")
  load(wt.file)
  wt.df <- as.data.frame(res)

  wt.sig <- wt.df %>% rownames_to_column("ATG") %>%
    filter(padj<0.05)  %>%
    filter(log2FoldChange < -1) %>%
    filter(!ATG %in% mut.up$ATG)

  #reduce to L2FC and return
  wt.FCs <- subset(wt.sig, select = c("ATG","log2FoldChange"))
  wt.FCs$log2FoldChange <- as.numeric(wt.FCs$log2FoldChange)
  names(wt.FCs) <- c("ATG", wt)
  return(wt.FCs)
}

#list of treatment results objects
PAMPTime <- c("Col_LPS_005","Col_LPS_010","Col_LPS_030","Col_LPS_090","Col_LPS_180",
              "Col_ch8_005","Col_ch8_010","Col_ch8_030","Col_ch8_090","Col_ch8_180",
              "Col_elf18_005","Col_elf18_010","Col_elf18_030","Col_elf18_090","Col_elf18_180",
              "Col_flg22_005","Col_flg22_010","Col_flg22_030","Col_flg22_090","Col_flg22_180",
              "Col_nlp20_005","Col_nlp20_010","Col_nlp20_030","Col_nlp20_090","Col_nlp20_180",
              "Col_OGs_005","Col_OGs_010","Col_OGs_030","Col_OGs_090","Col_OGs_180",
              "Col_Pep1_005","Col_Pep1_010","Col_Pep1_030","Col_Pep1_090","Col_Pep1_180")

#corresponding(!) list of control results objects
mutPAMPTime <- c("sd1-29_LPS_005","sd1-29_LPS_010","sd1-29_LPS_030","sd1-29_LPS_090","sd1-29_LPS_180",
                 "lyk4_5_ch8_005","lyk4_5_ch8_010","lyk4_5_ch8_030","lyk4_5_ch8_090","lyk4_5_ch8_180",
                 "efr-1_elf18_005","efr-1_elf18_010","efr-1_elf18_030","efr-1_elf18_090","efr-1_elf18_18
0",
                 "fls2c_flg22_005","fls2c_flg22_010","fls2c_flg22_030","fls2c_flg22_090","fls2c_flg22_18
0",
                 "rlp23-1_nlp20_005","rlp23-1_nlp20_010","rlp23-1_nlp20_030","rlp23-1_nlp20_090","rlp23-
1_nlp20_180",
                 "Col_mock_005","Col_mock_010","Col_mock_030","Col_mock_090","Col_mock_180",
                 "pepr1_2_Pep1_005","pepr1_2_Pep1_010","pepr1_2_Pep1_030","pepr1_2_Pep1_090","pepr1_2_Pe
p1_180")


#Run the function on each treatment/control together
log2FClist <- mapply(wt_sig_down, PAMPTime, mutPAMPTime,SIMPLIFY = F)
#combine into one data frame, and set log2FC to 0 for treatment/time combinations not significantly indu
ced
log2FCdf <- Reduce(function(x,y) merge(x, y, all=T), log2FClist)
log2FCdf[is.na(log2FCdf)] <- 0
nrow(log2FCdf)
```

```
## [1] 5157
```

filter to set downregulated by all 7 elicitors

```
altered <- log2FCdf %>% gather(-"ATG", key=Sample, value=L2FC) %>% separate(Sample, c("Geno","elicitor",
"time"), sep="_") %>% group_by(elicitor, ATG) %>% filter(sum(L2FC)<0)  %>% ungroup() %>% group_by(ATG, t
ime) %>% add_count() %>% filter(n>6) %>% unite(Sample, c("elicitor","time")) %>% spread(key=Sample, valu
e=L2FC) %>% select(-c("Geno","n")) %>% column_to_rownames("ATG")

#clean up some nomenclature, order
names(altered) <- gsub("LPS", "3-OH-FA", names(altered))
names(altered) <- gsub("ch8", "CO8", names(altered))


nrow(altered)
```

```
## [1] 93
```

Cluster by spearman (no scaling/centering)

```
D <- spearman.dist(as.matrix(altered))

Gene_column <- altered %>% rownames_to_column("ATG")

uptree <- hclust(D)
```

After exploratory data analysis, chose 3 clusters

```
#First make dendrogram
num = 3

clusters <- cutree(uptree, k=num)
cluster.df <- as.data.frame(clusters) %>% rownames_to_column("ATG")
#Re-ordering dendrogram by speed of response
updend <- as.dendrogram(uptree) %>% rotate(c(cluster.df$ATG[cluster.df$clusters==2], cluster.df$ATG[clus
ter.df$clusters==1], cluster.df$ATG[cluster.df$clusters==3]))
gene_order <- updend %>% labels
updatedorder <- c("#225522","#666633", "#663333") #colors for dendrogram, modified from Paul Tol dark qu
alitative
updend <- updend %>% set("branches_k_color", k=num, value=updatedorder)%>% set("branches_lwd", 0.3)

#arrange data for heatmap
longL2FC <- Gene_column %>% pivot_longer(-ATG, names_to="sample", values_to="L2FC") %>% separate(sample,
into=c("elicitor","time"), sep="_")

longL2FC$elicitor <- factor(longL2FC$elicitor, levels=c("flg22","elf18","Pep1","nlp20","OGs","CO8","3-OH
-FA"))
x_labs <- c("|   ","","flg22","","   |",
            "","","elf18","","   |",
            "","","Pep1","","   |",
            "","","nlp20","","   |",
            "","","OGs","","   |",
            "","","CO8","","   |",
            "","","3-OH-FA","","   | ")
```

Now make final plots, merge, and save

```
heatmap <- ggplot(longL2FC, aes(x=interaction(time, elicitor), y=factor(ATG, levels=gene_order), fill=L2
FC))+
  geom_tile()+
  scale_fill_viridis_c(guide=guide_colorbar(barheight=unit(0.1, "cm")))+
  scale_x_discrete(labels=x_labs, expand=c(0,0))+ scale_y_discrete(expand=c(0,0))+
  theme_minimal(base_size=5)+
  theme(axis.ticks=element_blank(), axis.title = element_blank(),
        axis.text.y=element_blank(), rect=element_rect(fill="transparent", color=NA),
        panel.border=element_blank(), legend.position="none")
simpleheat <- ggplot(longL2FC, aes(x=interaction(time, elicitor), y=ATG, fill=L2FC))+
  geom_tile()+scale_fill_viridis(guide=guide_colorbar(barheight=unit(0.1, "cm")))+
  theme_minimal(base_size=4)+
  theme(rect=element_rect(fill="transparent", color=NA),
        panel.border=element_blank(), legend.position="top")
heatlegend <- get_legend(simpleheat)

#and tree plot
tree <- ggplot(updend, horiz=T, labels=F)+
  theme(plot.margin = margin(0, 0, 0,0, "pt") )+
  scale_x_continuous(expand=c(0.01,0.01))+scale_y_reverse(expand=c(0.01,0)) +
  theme(rect=element_rect(fill="transparent", color=NA),
        panel.border=element_blank())
```

```
## Scale for 'y' is already present. Adding another scale for 'y', which will
## replace the existing scale.
```

```
combo <- ggdraw()+
  draw_plot(tree, x=0,width=0.115,y=-0.0469,height=0.972)+
  draw_plot(heatmap, x=0.10, width=0.915, y=0, height=0.93)+
  draw_plot(heatlegend, x=0.3, width=0.6, y=0.9, height=0.1)

ggsave(combo, filename="PanelB.pdf", height=6.5, width=8.5, units="cm", bg="transparent")
```

##Panel C

Plot average expression each cluster

```
se <- function(x) sqrt(var(x)/length(x))

clusters.df <- data.frame(cluster=clusters, ATG=names(clusters))
addcluster <- merge(clusters.df, longL2FC, by="ATG")

Percluster <- addcluster %>% group_by(cluster, elicitor, time) %>% summarize(mean=mean(L2FC), SE=se(L2F
C))
```

```
## `summarise()` regrouping output by 'cluster', 'elicitor' (override with `.groups` argument)
```

```
Percluster$cluster <- factor(Percluster$cluster, levels=c("3","1","2"))

labels <- c("1"="Late\n46 genes", "3"="Rapid transient\n25 genes","2"="Very late\n22 genes")

clusterplot <- ggplot(Percluster, aes(x=as.factor(as.numeric(time)), y=mean, group=elicitor,color=elicit
or))+
  geom_line(size=0.4) +
  geom_errorbar(aes(ymin=mean-SE, ymax=mean+SE), size=0.3, width=0.12)+
  scale_color_manual(values=definedvibrant)+
  theme_minimal(base_size=7)+
  theme(rect=element_rect("transparent",color=NA), legend.position="none")+
  facet_grid(cluster~., labeller=labeller(cluster=labels))+
  labs(x="Time (min)",y=expression("Cluster avg log"[2]*"(FC)"), color="Elicitor")

ggsave(clusterplot,filename="PanelC.pdf", height=6.8, width=3.5, units="cm", bg="transparent")
```

### Figure S5

Marta Bjornson

```
knitr::opts_chunk$set(echo = TRUE)
library(tidyverse)
library(GO.db)
library(topGO)
library(org.At.tair.db)
library(cowplot)
library(ggpubr) #for ggarrange
library(viridis)
library(readxl)
```

### Panel A

As in Figure 2, divide genes by when first down-regulated, then find enriched GO terms in each subset

```
rm(list=ls())
home <- "D:/OneDrive/Norwich_Z/Overall/CleanedData/"

#First load up data and organize by time at which genes first induced
PAMPaltered <- read.csv(paste0(home, "CleanedData_p0.1_l2FCabs1_ordered.csv"),header=T)
PAMPaltered <- gather(PAMPaltered, -X, key=sample, value=l2FC)
PAMPaltered <- PAMPaltered %>% separate(sample, into=c("TRT","Time"), sep="_")
colnames(PAMPaltered)[colnames(PAMPaltered)=="X"] <- "ATG"

PAMPdown <- PAMPaltered[PAMPaltered$l2FC < 0,]
downfirst <- PAMPdown %>% group_by(ATG) %>% summarise(FirstInduced = min(Time))


#Then calculate enriched GO terms
geneUniverse <- read.csv(paste0(home, "wtonly/Analysis-MMclustering/All_detected.csv"), header=F)
geneUniverse <- as.factor(as.vector(geneUniverse$V1))

MakeGOTable <- function (x,cid){
  clusters=unique(cid)
  K=length(clusters)
  for(i in 1:K){
    geneGalaxy <- x[cid==clusters[i]]
    if(length(geneGalaxy)!=1){
    geneList <- factor(as.integer(geneUniverse %in% geneGalaxy))
    names(geneList) <- geneUniverse
    GOdata <- new("topGOdata", ontology = "BP", allGenes = geneList, nodeSize = 10, annot = annFUN.org,
 mapping="org.At.tair.db")
    #so many options for different tests...
    resultTopGO.weight01 <- runTest(GOdata, algorithm = "weight01", statistic = "Fisher" )
    fill <- GenTable(GOdata, Uncorrected_p = resultTopGO.weight01, topNodes = 200)
    fill$Uncorrected_p[fill$Uncorrected_p=="< 1e-30"] <- 1e-30
    fill$Uncorrected_p <- as.numeric(fill$Uncorrected_p)
    write.csv(fill, file=paste0("GO_time",clusters[i], "down.csv"))
    }
  }
}

MakeGOTable(downfirst$ATG, downfirst$FirstInduced)
```

```r
GetAndMerge <- function(big, new){
  new.df <- read.csv(paste0("GO_time", new,".csv"), header=T)
  new.df <- new.df[,c("Term", "Uncorrected_p")]
  #Optional add multiple testing correction here
  new.df <- new.df[new.df$Uncorrected_p<0.01,]
  bigger <- merge(big, new.df, by="Term", all=T)
  names(bigger) <- c(names(big), new)
  return(bigger)
}
down <- read.csv("GO_time005down.csv", header=T)
down <- down[,c("Term", "Uncorrected_p")]
down <- down[down$Uncorrected_p<0.01,]
names(down) <- c("Term","005down")

down <- GetAndMerge(down, "010down")
down <- GetAndMerge(down, "030down")
down <- GetAndMerge(down, "090down")
down <- GetAndMerge(down, "180down")

write.csv(down, file="GOdown.csv")
```

```r
rm(list=ls())

GOsort <- read.csv("GOdown.csv", header=T, row.names=2)
GOsort[1] <- NULL
colnames(GOsort) <- gsub("down","min",colnames(GOsort))
colnames(GOsort) <- gsub("X","",colnames(GOsort))
GOsort[is.na(GOsort)] <- 0.01

GOsortlog <- -log10(GOsort)
GOsortlog <- GOsortlog[rowSums(GOsortlog)>10.5,]
GOsortlog <- GOsortlog %>% rownames_to_column("ATG") %>% mutate(allp=rowSums(.[2:6]))


GO005 <- GOsortlog[max.col(GOsortlog[2:6], ties.method="first")==1,] %>% arrange(desc(allp)) %>% arrange
(desc(`005min`)) %>% column_to_rownames("ATG")
GO010 <- GOsortlog[max.col(GOsortlog[2:6], ties.method="first")==2,] %>% arrange(desc(allp)) %>% arrange
(desc(`010min`)) %>% column_to_rownames("ATG")
GO030 <- GOsortlog[max.col(GOsortlog[2:6], ties.method="first")==3,] %>% arrange(desc(allp)) %>% arrange
(desc(`030min`)) %>% column_to_rownames("ATG")
GO090 <- GOsortlog[max.col(GOsortlog[2:6], ties.method="first")==4,] %>% arrange(desc(allp)) %>% arrange
(desc(`090min`)) %>% column_to_rownames("ATG")
GO180 <- GOsortlog[max.col(GOsortlog[2:6], ties.method="first")==5,] %>% arrange(desc(allp)) %>% arrange
(desc(`180min`)) %>% column_to_rownames("ATG")

GOarranged <- rbind(GO005, GO010, GO030, GO090, GO180)
colnames(GOarranged)[colnames(GOarranged)=="005min"] <- "5"
colnames(GOarranged)[colnames(GOarranged)=="010min"] <- "10"
colnames(GOarranged)[colnames(GOarranged)=="030min"] <- "30"
colnames(GOarranged)[colnames(GOarranged)=="090min"] <- "90"
colnames(GOarranged)[colnames(GOarranged)=="180min"] <- "180"

ggGOarranged <- GOarranged[1:5] %>% rownames_to_column("GOterm") %>% pivot_longer(-"GOterm",names_to="ti
me",values_to = "p")
```

```
GOhm <- ggplot(ggGOarranged, aes(x=as.factor(as.numeric(time)), y=factor(GOterm, levels=rev(unique(GOter
m))), fill=p)) +
  theme_minimal(base_size=7) + geom_tile()+scale_fill_viridis_c(limits=c(0,30), guide=guide_colorbar(bar
width=unit(1.5, "cm"), barheight=unit(0.2, "cm")))+
  theme(axis.text.y=element_blank(), axis.ticks.y=element_blank(), rect=element_rect(fill="transparent",
color=NA), panel.grid.major=element_blank(), legend.position="top", axis.text.x=element_text(angle=45, h
just=1, vjust=1), plot.margin=unit(c(0.25,0,0.25,0), "cm"))+
  labs(x="time", y="", fill=expression("-log"[10]*"(p value)"))


GOguide <- ggplot(ggGOarranged, aes(x=as.factor(as.numeric(time)), y=factor(GOterm, levels=rev(unique(GO
term))), fill=p)) +
  theme_minimal(base_size=7) + geom_tile()+scale_fill_viridis_c(limits=c(0,30), guide=guide_colorbar(bar
width=unit(1.5, "cm"), barheight=unit(0.2, "cm")))+
  theme(axis.ticks.y=element_blank(), rect=element_rect(fill="transparent",color=NA), panel.grid.major=e
lement_blank(), legend.position="top", axis.text.x=element_text(angle=45, hjust=1, vjust=1))+
  labs(x="time", y="", fill=expression("-log"[10]*"(p value)"))
ggsave("GOguide_down.pdf", height=15, width=8)
```

### Panel B

To make a similar figure for cis element enrichment, need same data but this time save gene lists based on time induced

```
home <- "D:/OneDrive/Norwich_Z/Overall/CleanedData/"
#First load up data and organize by time at which genes first induced
PAMPaltered <- read.csv(paste0(home, "CleanedData_p0.1_l2FCabs1_ordered.csv"),header=T)
PAMPaltered <- gather(PAMPaltered, -X, key=sample, value=l2FC)
PAMPaltered <- PAMPaltered %>% separate(sample, into=c("TRT","Time"), sep="_")
colnames(PAMPaltered)[colnames(PAMPaltered)=="X"] <- "ATG"

PAMPdown <- PAMPaltered[PAMPaltered$l2FC < 0,]
downfirst <- PAMPdown %>% group_by(ATG) %>% summarise(FirstInduced = min(Time))
```

```
## `summarise()` ungrouping output (override with `.groups` argument)
```

```
write.csv(downfirst[downfirst$FirstInduced=='005',], file="genes_005down.csv")
write.csv(downfirst[downfirst$FirstInduced=='010',], file="genes_010down.csv")
write.csv(downfirst[downfirst$FirstInduced=='030',], file="genes_030down.csv")
write.csv(downfirst[downfirst$FirstInduced=='090',], file="genes_090down.csv")
write.csv(downfirst[downfirst$FirstInduced=='180',], file="genes_180down.csv")
```

These .csv files were run through GetPromotersFasta.py through cygwin and then ame, on cluster, to get enriched cis elements for each set. The ame output was processed with Extract_motif_p_fromAME.py, called through AMEparse.bash. This gave me one file per time (up and down)

```
GetAndMerge <- function(big, new){
  new.df <- read.csv(paste0("D:/OneDrive/Norwich_Z/Overall/CleanedData/wtonly/Analysis-cis/Figure1_JIT_A
ME/", new,".csv"), header=T)
  new.df <- new.df[,c("X1", "X2")]
  names(new.df) <- c("TF","corrected_p")
  #Optional add multiple testing correction here
  new.df <- new.df[new.df$corrected_p<0.01,]
  bigger <- merge(big, new.df, by="TF", all=T)
  names(bigger) <- c(names(big), new)
  return(bigger)
}

down <- read.csv("D:/OneDrive/Norwich_Z/Overall/CleanedData/wtonly/Analysis-cis/Figure1_JIT_AME/down_00
5.csv", header=T)
down <- down[,c("X1", "X2")]
names(down) <- c("TF","corrected_p")
down <- down[down$corrected_p<0.01,]
names(down) <- c("TF","down_005")

#down <- GetAndMerge(down, "down_010")
down <- GetAndMerge(down, "down_030")
down <- GetAndMerge(down, "down_090")
#down <- GetAndMerge(down, "down_180")


down <- down %>% group_by(TF) %>% summarise(`005min`=mean(down_005), `030min`=mean(down_030),`090min`=me
an(down_090))
```

```
## `summarise()` ungrouping output (override with `.groups` argument)
```

```
write.csv(down, file="cis_down.csv")
```

```
cistable <- read.csv("cis_down.csv")[,c(2,4,5)]
colnames(cistable) <- gsub("X","",colnames(cistable))
cistable[is.na(cistable)] <- 0.01
cistable <- column_to_rownames(cistable, "TF")
cistablelog <- -log10(cistable)
cistablelog <- cistablelog[rowSums(cistablelog)>4.1,]
cistablelog <- cistablelog %>% rownames_to_column("TF") %>% mutate(allp=rowSums(.[2:length(cistablelo
g)]))


cis030 <- cistablelog[max.col(cistablelog[2:length(cistable)], ties.method="first")==1,] %>% arrange(des
c(allp)) %>% arrange(desc(`030min`)) %>% column_to_rownames("TF")
cis090 <- cistablelog[max.col(cistablelog[2:length(cistable)], ties.method="first")==2,] %>% arrange(des
c(allp)) %>% arrange(desc(`090min`)) %>% column_to_rownames("TF")


cisarranged <- rbind(cis030, cis090)
colnames(cisarranged) <- c("30","90","allp")

ggcisarranged <- cisarranged[,1:2] %>% rownames_to_column("cis") %>% pivot_longer(-"cis",names_to="time"
,values_to = "p")
```

```
cishm <- ggplot(ggcisarranged, aes(x=as.factor(as.numeric(time)), y=factor(cis, levels=rev(unique(ci
s))), fill=p)) +
  theme_minimal(base_size=7) + geom_tile()+scale_fill_viridis_c(limits=c(0,30),guide=guide_colorbar(barw
idth=unit(1.5, "cm"), barheight=unit(0.2, "cm")))+
  theme(rect=element_rect(fill="transparent",color=NA), panel.grid.major=element_blank(),
        legend.position="top", axis.text.x=element_text(angle=45, hjust=1, vjust=1), plot.margin=unit(c(
0.25,-0.25,0.25,0), "cm"))+
  scale_y_discrete(position="right")+
  labs(x="time (min)", y="", fill=expression("-log"[10]*"(p value)"))
```

```
spacer <- ggplot(ggcisarranged, aes(x=time, y=cis))+
  theme_nothing()+
  theme(rect=element_rect(fill="transparent",color=NA), axis.text=element_blank(), plot.margin=unit(c(0.
25,0,0.25,0), "cm"))

hms <- ggarrange(GOhm, spacer, cishm, widths = c(1.25,1.75,2), ncol=3, common.legend=T)
ggsave(hms, filename="PanelAB.pdf",width=5, height=9, units="cm", bg="transparent")
```

### Panel C

Genes down by number abio and number elicitors, by time.

```
rm(list=ls())
home <- "D:/OneDrive/Norwich_Z/Overall/CleanedData/wtonly/"
#load up the new spreadsheet (as of 2019-04ish) with abio stress done by probe
FullInfo <- read.csv(paste0(home,"2020-11-23AllGenesInfo_DOWNfocus.csv"), header=T)
#remove genes not induced by elicitors, or where abiotic stress information isn't clean
TimeAbio <- FullInfo[!is.na(FullInfo$NumberElicitors),]
TimeAbio <- TimeAbio[TimeAbio$AtGenExpressNumber != "NotOnArray" & TimeAbio$AtGenExpressNumber != "Probe
HitsMultipleGenes",]
TimeAbio$NumberElicitors <- as.factor(TimeAbio$NumberElicitors)
```

timebox with induction

```
TimeAbio <- TimeAbio[,2:length(TimeAbio)]
maxrepress <- TimeAbio %>% gather(-ATG, -AtGenExpressNumber, -NumberElicitors, -FirstRepressed, key=cond
ition, value=L2FC) %>% group_by(ATG, AtGenExpressNumber, NumberElicitors, FirstRepressed) %>% summarize
(greatest=min(L2FC))
```

```
## `summarise()` regrouping output by 'ATG', 'AtGenExpressNumber', 'NumberElicitors' (override with `.gr
oups` argument)
```

```
box <- ggplot(maxrepress, aes(x=NumberElicitors , y=AtGenExpressNumber)) +
  geom_jitter(height=0.25, width=0.25, size=0.5, aes(color=greatest, fill=greatest), shape=21,stroke=0.1
2) +
  theme_minimal(base_size=7) +
  theme(panel.background = element_blank(), plot.background = element_blank(), legend.background = eleme
nt_blank(), legend.box.background = element_blank(), panel.border = element_blank(), legend.box=NULL, le
gend.position = "top", strip.text.y=element_text(angle=0), legend.title=element_text(size=4), legend.tex
t=element_text(size=6)) +
  labs(x="Number patterns repressing", y="Number abiotic stresses repressing", color=bquote('Min '*~log[
2]*'(FC)')) +
  facet_grid(FirstRepressed~.) +
  scale_color_viridis_c(option="inferno", direction=1, limits=c(-4.5,0), breaks=c(-4.5, -3, -1.5, 0), gu
ide=guide_colorbar(barwidth=unit(1.5, "cm"), barheight=unit(0.1, "cm")))+
  scale_fill_viridis_c(option="inferno", direction=1, limits=c(-4.5, 0), alpha=0.3, guide=F)



ggsave("PanelC.pdf", box, height=9, width=3.5, units="cm", bg="transparent")
```

### Figure S7

Marta Bjornson

```
knitr::opts_chunk$set(echo = TRUE)
library(tidyverse)
library(readxl)
library(emmeans)
library(gtools)#for mixedsort
library(cowplot)
```

### Panel A

Kinetics of the Calcium burst, where peak shown in Figure 3a. First establish functions for reading in data

```
rm(list=ls())
home <- "D:/OneDrive/Norwich_U/Data-Ca/"
muted <- c("black", "#BBBBBB", "#117733", "#44AA99", "#999933", "#DDCC77",  "#332288", "#88CCEE", "#AA44
99", "#CC6677")
smallmuted <- c("black", "#BBBBBB", "#AA4499", "#CC6677")
#These two functions read in treatment data, from an excel file. Set up the excel file in the orientatio
n in which the plate(s) went into the camera EXCEPT if there are two plates, stack them vertically not h
orizontally. This is critical! There is no error-checking in the code to ensure that a well lines up wit
h the appropriate region of interest.
makelist <- function(filename, sheet){
  worksheet <- sheet
  variable <- read_excel(filename, sheet = worksheet)
  variablelist <- variable[1,]
  for(i in 2:nrow(variable)){
    variablelist <- c(variablelist, variable[i,])
  }
  variablevector <- unlist(variablelist)
  return(variablevector)
}

add_treatments <- function(filename, inheritedros){
  looper <- excel_sheets(filename)
  for(i in 1:length(looper)){
    column <- looper[i]
    columnlist <- makelist(filename, column)
    inheritedros$new <- unlist(columnlist)
    colnames(inheritedros)[colnames(inheritedros)=="new"] <- column
  }
  return(inheritedros)
}

ReadTecan <- function(datafile, planfile, treatment, conc, exp){
  base_cfp1 <- read_excel(datafile, range="B82:CU92") %>% mutate_if(is.character, funs(recode(.,"OVER" =
0))) %>%
    pivot_longer(-c("Time [s]", "Temp. [°C]"), names_to="Well", values_to="CFP") %>% na.omit() %>% selec
t(-"Temp. [°C]")
  base_yfp1 <- read_excel(datafile, range="B95:CU105")%>% mutate_if(is.character, funs(recode(.,"OVER" =
0))) %>%
    pivot_longer(-c("Time [s]", "Temp. [°C]"), names_to="Well", values_to="YFP") %>% na.omit() %>% selec
t(-"Temp. [°C]")
  base_cfp1$`Time [s]` <- base_cfp1$`Time [s]` - 300
  base_yfp1$`Time [s]` <- base_yfp1$`Time [s]` - 300

  cfp1 <- read_excel(datafile, range="B161:CU211") %>% mutate_if(is.character, funs(recode(.,"OVER" = 0
))) %>%
    pivot_longer(-c("Time [s]", "Temp. [°C]"), names_to="Well", values_to="CFP") %>% na.omit() %>% selec
t(-"Temp. [°C]")
  yfp1 <- read_excel(datafile, range="B214:CU264") %>% mutate_if(is.character, funs(recode(.,"OVER" = 0
))) %>%
    pivot_longer(-c("Time [s]", "Temp. [°C]"), names_to="Well", values_to="YFP") %>% na.omit() %>% selec
t(-"Temp. [°C]")

  #combine into one data frame
  yfp <- rbind(base_yfp1, yfp1)
  cfp <- rbind(base_cfp1, cfp1)

  #get ratios
  fluor <- merge(cfp, yfp, by=c("Time [s]", "Well"))
  fluor$`Time [s]` <- round(fluor$`Time [s]`)
  fluor$`Time (min)` <- round(fluor$`Time [s]`/60, 1)
  fluor$ratio <- fluor$YFP/fluor$CFP
  fluor$Well <- factor(fluor$Well, levels=mixedsort(unique(fluor$Well)))
```

```
  fill <- data.frame(filler=rep(0,96))
  trt <- add_treatments(planfile,fill)

  fluor <- merge(fluor, trt[,2:ncol(trt)], by="Well")
  fluor$Row <- substring(fluor$Well, 1, 1)
  fluor$Column <- substring(fluor$Well, 2, nchar(as.character(fluor$Well))) %>% as.numeric()

  fluor$Trt <- paste0(conc,treatment)
  fluor$Date <- exp
  return(fluor)
}

ReadSalt <- function(datafile, planfile, treatment, conc, exp){
  base_cfp1 <- read_excel(datafile, range="B83:CU93") %>% mutate_if(is.character, funs(recode(.,"OVER" = 
0))) %>%
    pivot_longer(-c("Time [s]", "Temp. [°C]"), names_to="Well", values_to="CFP") %>% na.omit() %>% selec
t(-"Temp. [°C]")
  base_yfp1 <- read_excel(datafile, range="B96:CU106")%>% mutate_if(is.character, funs(recode(.,"OVER" = 
0))) %>%
    pivot_longer(-c("Time [s]", "Temp. [°C]"), names_to="Well", values_to="YFP") %>% na.omit() %>% selec
t(-"Temp. [°C]")
  base_cfp1$`Time [s]` <- base_cfp1$`Time [s]` - 300
  base_yfp1$`Time [s]` <- base_yfp1$`Time [s]` - 300

  cfp1 <- read_excel(datafile, range="B161:CU174") %>% mutate_if(is.character, funs(recode(.,"OVER" = 0
))) %>%
    pivot_longer(-c("Time [s]", "Temp. [°C]"), names_to="Well", values_to="CFP") %>% na.omit() %>% selec
t(-"Temp. [°C]")
  yfp1 <- read_excel(datafile, range="B177:CU190") %>% mutate_if(is.character, funs(recode(.,"OVER" = 0
))) %>%
    pivot_longer(-c("Time [s]", "Temp. [°C]"), names_to="Well", values_to="YFP") %>% na.omit() %>% selec
t(-"Temp. [°C]")

  #combine into one data frame
  yfp <- rbind(base_yfp1, yfp1)
  cfp <- rbind(base_cfp1, cfp1)

  #get ratios
  fluor <- merge(cfp, yfp, by=c("Time [s]", "Well"))
  fluor$`Time [s]` <- round(fluor$`Time [s]`)
  fluor$`Time (min)` <- round(fluor$`Time [s]`/60, 1)
  fluor$ratio <- fluor$YFP/fluor$CFP

  fill <- data.frame(filler=rep(0,96))
  trt <- add_treatments(planfile,fill)

  fluor <- merge(fluor, trt[,2:ncol(trt)], by="Well")
  fluor$Row <- substring(fluor$Well, 1, 1)
  fluor$Column <- substring(fluor$Well, 2, nchar(as.character(fluor$Well))) %>% as.numeric()

  fluor$Trt <- paste0(conc,treatment)
  fluor$Date <- exp
  return(fluor)
}
```

Read in data, all identical to Figure 3a

```
#A <- ReadTecan(datafile, planfile, trt, conc, exp)
#Techniclly, I have some 1uM flg22 on 2020-08-24 as well, but the plants aren't the same age/pushed dow
n/just take the three standard exp as I have for other elicitors
Aflg <- ReadTecan(paste0(home,"2020-09-01_pn4b_1uMflg22.xlsx"), paste0(home,"2020-09-01_pn3.xlsx"), "flg
22", "1 \U00B5M ", "A")
```

```
## Warning: `funs()` is deprecated as of dplyr 0.8.0.
## Please use a list of either functions or lambdas:
##
##   # Simple named list:
##   list(mean = mean, median = median)
##
##   # Auto named with `tibble::lst()`:
##   tibble::lst(mean, median)
##
##   # Using lambdas
##   list(~ mean(., trim = .2), ~ median(., na.rm = TRUE))
## This warning is displayed once every 8 hours.
## Call `lifecycle::last_warnings()` to see where this warning was generated.
```

```
#Remove wells: check individual day for each one
Aflg <- Aflg %>% filter(!Well %in% c("H7", "A8"))

Bflg <- ReadTecan(paste0(home,"2020-09-08_p2a_1uMflg22.xlsx"), paste0(home,"2020-09-08_plan.xlsx"), "flg
22", "1 \U00B5M ", "B")
Bflg <- Bflg %>% filter(!Well %in% c("C4","D5","A3"))

Cflg <- ReadTecan(paste0(home,"2020-09-15_p1b_1uMflg22.xlsx"),paste0(home,"2020-09-15_plan.xlsx"), "flg2
2","1 \U00B5M ", "C")
Cflg <- Cflg %>% filter(!Well %in% c("E7","F8"))


Amock <- ReadTecan(paste0(home,"2020-09-01_pn3b_mock.xlsx"), paste0(home,"2020-09-01_pn3.xlsx"), "Mock",
"", "A")
#Remove wells: check individual day for each one
Amock <- Amock %>% filter(!Well %in% c())

Bmock <- ReadTecan(paste0(home,"2020-09-08_p1a_mock.xlsx"), paste0(home,"2020-09-08_plan.xlsx"), "Mock",
"", "B")
Bmock <- Bmock %>% filter(!Well  %in%  c("B4"))

Cmock <- ReadTecan(paste0(home,"2020-09-15_p1a_mock.xlsx"),paste0(home,"2020-09-15_plan.xlsx"),"Mock",
"", "C")
Cmock <- Cmock %>% filter(!Well %in% c("A6","C4","G2"))

Aelf <- ReadTecan(paste0(home,"2020-09-01_po4b_1uMelf18.xlsx"), paste0(home,"2020-09-01_pn3.xlsx"), "elf
18", "1 \U00B5M ", "A")
#Remove wells: check individual day for each one
Aelf <- Aelf %>% filter(!Well %in% c("A9"))

Belf <- ReadTecan(paste0(home,"2020-09-08_p2b_1uMelf18.xlsx"), paste0(home,"2020-09-08_plan.xlsx"), "elf
18", "1 \U00B5M ", "B")
```

```
## Warning: Problem with `mutate()` input `F11`.
## i Unreplaced values treated as NA as .x is not compatible. Please specify replacements exhaustively o
r supply .default
## i Input `F11` is `recode(F11, OVER = 0)`.
```

```
## Warning: Unreplaced values treated as NA as .x is not compatible. Please specify
## replacements exhaustively or supply .default
```

```
## Warning: Problem with `mutate()` input `F11`.
## i Unreplaced values treated as NA as .x is not compatible. Please specify replacements exhaustively o
r supply .default
## i Input `F11` is `recode(F11, OVER = 0)`.
```

```
## Warning: Unreplaced values treated as NA as .x is not compatible. Please specify
## replacements exhaustively or supply .default
```

```
Belf <- Belf %>% filter(!Well %in% c("C10","F11","B8","F7","G10"))

Celf <- ReadTecan(paste0(home,"2020-09-15_p4a_1uMelf18.xlsx"),paste0(home,"2020-09-15_plan.xlsx"), "elf1
8","1 \U00B5M ", "C")
```

```
## Warning: Problem with `mutate()` input `G2`.
## i Unreplaced values treated as NA as .x is not compatible. Please specify replacements exhaustively o
r supply .default
## i Input `G2` is `recode(G2, OVER = 0)`.

## Warning: Unreplaced values treated as NA as .x is not compatible. Please specify replacements exhaust
ively or supply .default
```

```
## Warning: Problem with `mutate()` input `G2`.
## i Unreplaced values treated as NA as .x is not compatible. Please specify replacements exhaustively o
r supply .default
## i Input `G2` is `recode(G2, OVER = 0)`.
```

```
## Warning: Unreplaced values treated as NA as .x is not compatible. Please specify
## replacements exhaustively or supply .default
```

```
Celf <- Celf %>% filter(!Well %in% c("G2","A3","C5","F5"))

Apep <- ReadTecan(paste0(home,"2020-09-01_po4a_1uMpep1.xlsx"), paste0(home,"2020-09-01_pn3.xlsx"), "Pep
1", "1 \U00B5M ", "A")
#Remove wells: check individual day for each one
Apep <- Apep %>% filter(!Well %in% c("A8"))

Bpep <- ReadTecan(paste0(home,"2020-09-08_p1b_1uMPep1.xlsx"), paste0(home,"2020-09-08_plan.xlsx"), "Pep
1", "1 \U00B5M ", "B")
Bpep <- Bpep %>% filter(!Well %in% c("A10", "A7", "G7", "G9"))

Cpep <- ReadTecan(paste0(home,"2020-09-15_p2a_1uMPep1_MovedByElectrician.xlsx"),paste0(home,"2020-09-15_
plan.xlsx"), "Pep1","1 \U00B5M ", "C")
Cpep <- Cpep %>% filter(!Well %in% c("B9","D10"))
Cpep$Column <- Cpep$Column+6
```

Bind all data together, then find and remove wells (by date) where baseYFP is not above 3x level of non-fluorescent Col-0 control, to deal with some silencing. Next normalize ratios to first value post-injection, combine genotypes (tested seedlings from 2-3 different parent plants, all parents homozygous) and clean up names.

```
fluor <- rbind(Amock, Bmock, Cmock, Aflg,  Bflg, Cflg, Aelf, Belf, Celf, Apep, Bpep, Cpep)
Conc <- "1 \U00B5M "
Trt <- "flg etc"

#for now getting one Col-0 fluorescence value across all exp. In theory (and from seeing many of these)
 it shouldn't change much
Col_YFP <- fluor %>% filter(`Time (min)`<0) %>% filter(Genotype=="Col-0") %>% group_by(Well) %>% summari
ze(wellmean = mean(YFP)) %>% ungroup() %>% summarize(mean=mean(wellmean))
```

```
## `summarise()` ungrouping output (override with `.groups` argument)
```

```
Col_CFP <- fluor %>% filter(`Time (min)`<0) %>% filter(Genotype=="Col-0") %>% group_by(Well) %>% summari
ze(wellmean = mean(CFP)) %>% ungroup() %>% summarize(mean=mean(wellmean))
```

```
## `summarise()` ungrouping output (override with `.groups` argument)
```

```
#select only if THREE times Col-0 mean. (previously 5x, don't think I need to be that stringent)
fluor_judge <- fluor %>% filter(`Time (min)`<0)  %>% group_by(Date, Well, Trt, Genotype) %>% summarize(m
eanYFP=mean(YFP), meanCFP=mean(CFP)) %>% mutate(YFP_Judgement = if_else(meanYFP > Col_YFP*3, "Yes", "No"
)) %>% mutate(CFP_Judgement = if_else(meanCFP > Col_CFP*3,"Yes", "No")) %>% mutate(Judgement = if_else(Y
FP_Judgement =="Yes" & CFP_Judgement=="Yes", "Yes", "No"))
```

```
## `summarise()` regrouping output by 'Date', 'Well', 'Trt' (override with `.groups` argument)
```

```
fluor <-  merge(fluor, fluor_judge[,c("Date", "Well", "Judgement", "Trt")], by=c("Date", "Well", "Trt"))

time0norm <- function(time, ratio){
  ratio.df <- data.frame(`Time (min)`=time, "ratio"=ratio)
  colnames(ratio.df) <- c("Time (min)","ratio")
  pretreat <- ratio.df$ratio[ratio.df$`Time (min)` == 0]
  ratio.df$difference <- (ratio.df$ratio-pretreat)/pretreat
  return(ratio.df$difference)
}

norm_0 <- fluor %>% group_by(Date, Well, Trt, Genotype,Judgement) %>% arrange(`Time (min)`) %>% mutate(d
ifference=time0norm(`Time (min)`, ratio))

filtered <- norm_0[norm_0$Judgement=="Yes",]

filtered <- filtered %>% ungroup() %>% add_column(Geno = "filler") %>% mutate(Geno = if_else(Genotype ==
"YC3.6 A" | Genotype == "YC3.6 B" | Genotype == "YC3.6 C", "YC3.6", Genotype)) %>% mutate(Geno = if_else
(Genotype =="C2-2C" | Genotype == "C2-2D" , "glr2.7/2.8/2.9", Geno))

filtered$Geno <- factor(filtered$Geno, levels=c("YC3.6","glr2.7/2.8/2.9"))
filtered$Trt <- factor(filtered$Trt, levels=c("Mock","1 μM flg22","1 μM elf18","1 μM Pep1"))
```

```
se <- function(au){ifelse(length(au)>1, sd(au)/sqrt(sum(!is.na(au))), 0)}
alltogether <- filtered %>% group_by(Geno, Trt, `Time (min)`) %>% summarize(mean=mean(difference), sterr
= se(difference), test=length(difference))
```

```
## `summarise()` regrouping output by 'Geno', 'Trt' (override with `.groups` argument)
```

```
PanelA <- ggplot(alltogether, aes(x=`Time (min)`, y=mean, color=interaction(Geno, Trt))) +
  geom_line(size=0.3)+geom_point(size=0.1) +geom_errorbar(aes(ymin=mean-sterr, ymax=mean+sterr), size=0.
2) +
  labs(y=expression("(R-R"[0]*")/R"[0]), title="")+
  scale_color_manual(values=muted)+
  facet_wrap(~Trt)+
  theme_cowplot()+
  theme(legend.position = "none", panel.background = element_blank(), plot.background = element_blank(),
        panel.border = element_blank(), strip.text.x = element_text(size=6), strip.background = element_
blank(),
        axis.text = element_text(size=6), axis.title = element_text(size=7),
        plot.margin=unit(c(-0.8, 0, 0, 0), "cm"))

ggsave(PanelA, filename = "PanelA.pdf", height=6, width=6, units="cm",bg="transparent")
```

### Panel B

The same for salt

```
setwd(home)
Amock <- ReadSalt("2020-09-15_p3a_FastMock.xlsx", "2020-09-15_plan.xlsx", "Mock", "", "A")
#Remove wells: check individual day for each one
Amock <- Amock %>% filter(!Well %in% c())

Asalt <- ReadSalt("2020-09-15_p3b_333mMNaCl.xlsx", "2020-09-15_plan.xlsx", "NaCl", "333 mM ", "A")
#Remove wells: check individual day for each one
Asalt <- Asalt %>% filter(!Well %in% c("A9","B7", "B9", "B11","E8","F11","G9","H11")) %>% mutate(Column=
Column-6)

Bmock <- ReadSalt("2020-10-02_p1a_mock.xlsx", "2020-10-02_plan.xlsx", "Mock", "", "B")
#Remove wells: check individual day for each one
Bmock <- Bmock %>% filter(!Well %in% c("E2")) %>% mutate(Column=Column+6)

Bsalt <- ReadSalt("2020-10-02_p1b_333mMNaCl.xlsx", "2020-10-02_plan.xlsx", "NaCl", "333 mM ", "B")
#Remove wells: check individual day for each one
Bsalt <- Bsalt %>% filter(!Well %in% c("B7", "B11","E9","F11"))

Cmock <- ReadSalt("2020-10-02_p2a_mock.xlsx", "2020-10-02_plan.xlsx", "Mock", "", "C")
#Remove wells: check individual day for each one
Cmock <- Cmock %>% filter(!Well %in% c("F4","H3")) %>% mutate(Column=Column+12)

Csalt <- ReadSalt("2020-10-02_p2b_333mMNaCl.xlsx", "2020-10-02_plan.xlsx", "NaCl", "333 mM ", "C")
#Remove wells: check individual day for each one
Csalt <- Csalt %>% filter(!Well %in% c("B11")) %>% mutate(Column=Column+6)
```

Bind all data together, then find and remove wells (by date) where baseYFP is not above 3x level of non-fluorescent Col-0 control, to deal with some silencing. Next normalize ratios to pretreatment values, combine genotypes (tested seedlings from 2-3 different parent plants, all parents homozygous) and clean up names.

```
fluor <- rbind(Amock, Asalt, Bmock, Bsalt, Cmock, Csalt)
Conc <- paste0("333 mM ")
Trt <- "NaCl, and Mock"

#for now getting one Col-0 fluorescence value across all exp. In theory (and from seeing many of these)
 it shouldn't change much
Col_YFP <- fluor %>% filter(`Time (min)`<0) %>% filter(Genotype=="Col-0") %>% group_by(Well) %>% summari
ze(wellmean = mean(YFP)) %>% ungroup() %>% summarize(mean=mean(wellmean))
```

```
## `summarise()` ungrouping output (override with `.groups` argument)
```

```
Col_CFP <- fluor %>% filter(`Time (min)`<0) %>% filter(Genotype=="Col-0") %>% group_by(Well) %>% summari
ze(wellmean = mean(CFP)) %>% ungroup() %>% summarize(mean=mean(wellmean))
```

```
## `summarise()` ungrouping output (override with `.groups` argument)
```

```
#select only if THREE times Col-0 mean. (previously 5x, don't think I need to be that stringent)
fluor_judge <- fluor %>% filter(`Time (min)`<0)  %>% group_by(Date, Well, Trt, Genotype) %>% summarize(m
eanYFP=mean(YFP), meanCFP=mean(CFP)) %>% mutate(YFP_Judgement = if_else(meanYFP > Col_YFP*3, "Yes", "No"
)) %>% mutate(CFP_Judgement = if_else(meanCFP > Col_CFP*3,"Yes", "No")) %>% mutate(Judgement = if_else(Y
FP_Judgement =="Yes" & CFP_Judgement=="Yes", "Yes", "No"))
```

```
## `summarise()` regrouping output by 'Date', 'Well', 'Trt' (override with `.groups` argument)
```

```r
fluor <-  merge(fluor, fluor_judge[,c("Date", "Well", "Judgement", "Trt")], by=c("Date", "Well", "Trt"))

pretreatnorm <- function(time, ratio){
  ratio.df <- data.frame(`Time (min)`=time, "ratio"=ratio)
  colnames(ratio.df) <- c("Time (min)","ratio")
  pretreat <- mean(ratio.df$ratio[ratio.df$`Time (min)`<0])
  ratio.df$difference <- (ratio.df$ratio-pretreat)/pretreat
  return(ratio.df$difference)
}

norm_0 <- fluor %>% group_by(Date, Well, Trt, Genotype,Judgement) %>% arrange(`Time (min)`) %>% mutate(d
ifference=pretreatnorm(`Time (min)`, ratio))

filtered <- norm_0[norm_0$Judgement=="Yes",]

filtered <- filtered %>% ungroup() %>% add_column(Geno = "filler") %>% mutate(Geno = if_else(Genotype ==
"YC3.6 A" | Genotype == "YC3.6 B" | Genotype == "YC3.6 C", "YC3.6", Genotype)) %>% mutate(Geno = if_else
(Genotype =="C2-2C" | Genotype == "C2-2D" , "glr2.7/2.8/2.9", Geno))

filtered$Geno <- factor(filtered$Geno, levels=c("YC3.6","glr2.7/2.8/2.9"))
filtered$Trt <- factor(filtered$Trt, levels=c("Mock","333 mM NaCl"))
```

```r
filterpre <- filtered[filtered$`Time [s]`<0,]
filterpre$`Time [min]` <- filterpre$`Time [s]`/60

pre <- filterpre %>% group_by(Geno, Trt, `Time [min]`) %>% summarize(mean=mean(difference), sterr = se(d
ifference), test=length(difference))
```

```
## `summarise()` regrouping output by 'Geno', 'Trt' (override with `.groups` argument)
```

```r
PanelCleft <- ggplot(pre, aes(x=`Time [min]`, y=mean, color=interaction(Geno, Trt))) +
  geom_line(size=0.3)+geom_point(size=0.1) +geom_errorbar(aes(ymin=mean-sterr, ymax=mean+sterr), size=0.
2) +
  labs(y=expression("(R-R"[0]*")/R"[0]), x="", title="")+
  scale_color_manual(values=smallmuted)+
  theme_cowplot()+
  ylim(c(-0.02, 0.4))+
  theme(legend.position = "none", panel.background = element_blank(), plot.background = element_blank(),
        panel.border = element_blank(), strip.text.x = element_blank(),
        axis.text = element_text(size=6), axis.title = element_text(size=7),
        plot.margin = unit(c(-0.25, -0.5, 0, 0), "cm"))

filtertrt <- filtered[filtered$`Time [s]`>-1,]
filtertrt$`Time [s]` <- filtertrt$`Time [s]`+2
post <- filtertrt %>% group_by(Geno, Trt, `Time [s]`) %>% summarize(mean=mean(difference), sterr = se(di
fference), test=length(difference))
```

```
## `summarise()` regrouping output by 'Geno', 'Trt' (override with `.groups` argument)
```

```
PanelCright <- ggplot(post, aes(x=`Time [s]`, y=mean, color=interaction(Geno, Trt))) +
  geom_line(size=0.3)+geom_point(size=0.1) +geom_errorbar(aes(ymin=mean-sterr, ymax=mean+sterr), size=0.
2) +
  labs(y="", x="", title="")+
  scale_color_manual(values=smallmuted)+
  theme_cowplot()+
  ylim(c(-0.02, 0.4))+
  theme(legend.position = "none", panel.background = element_blank(), plot.background = element_blank(),
        panel.border = element_blank(), strip.text.x = element_blank(),
        axis.text = element_text(size=6), axis.title = element_text(size=7),
        axis.line.y=element_blank(), axis.ticks.y=element_blank(), axis.text.y=element_blank(),
        plot.margin = unit(c(-0.25, 0, 0, 0), "cm"))

PanelB <- plot_grid(PanelCleft, PanelCright, rel_widths = c(1.3, 2))
```

```
## Warning: Removed 10 row(s) containing missing values (geom_path).
```

```
## Warning: Removed 10 rows containing missing values (geom_point).
```

```
## Warning: Removed 13 row(s) containing missing values (geom_path).
```

```
## Warning: Removed 13 rows containing missing values (geom_point).
```

```
ggsave(PanelB, filename = "PanelB.pdf", height=4, width=4, units="cm",bg="transparent")
```

### Panel C

Visualize/analyse stomatal aperture

```r
rm(list=ls())
home <- "D:/OneDrive/Norwich_U/DATA-Stomata/"
mtt <- c("#000000","#DDAA33","#004488")

date <- "2020-09-16"
A <- read_excel(paste0(home, date, ".xlsx")) %>% mutate(Stomate=rep(1:(length(Length)/2), each=2)) %>% s
elect(-ImageCounter, -Angle) %>% pivot_wider(names_from="Parameter", values_from="Length") %>% mutate(ra
tio=W/L)
A$Date <- date
A <- A %>% filter(Plate != '4')

date <- "2020-09-18"
B <- read_excel(paste0(home, date, ".xlsx")) %>% mutate(Stomate=rep(1:(length(Length)/2), each=2)) %>% s
elect(-ImageCounter, -Angle) %>% pivot_wider(names_from="Parameter", values_from="Length") %>% mutate(ra
tio=W/L)
B$Date <- date

date <- "2020-09-25"
C <- read_excel(paste0(home, date, ".xlsx")) %>% mutate(Stomate=rep(1:(length(Length)/2), each=2)) %>% s
elect(-ImageCounter, -Angle) %>% pivot_wider(names_from="Parameter", values_from="Length") %>% mutate(ra
tio=W/L)
C$Treatment[C$Treatment=="flg"] <- "flg22"
C$Date <- date
#experimented today with a clear plate, didn't see good treatment differences in Col-0 so exclude
C <- C[C$Plate !="c",]

date <- "2020-09-26"
D <- read_excel(paste0(home, date, ".xlsx")) %>% mutate(Stomate=rep(1:(length(Length)/2), each=2)) %>% s
elect(-ImageCounter, -Angle) %>% pivot_wider(names_from="Parameter", values_from="Length") %>% mutate(ra
tio=W/L)
D$Treatment[D$Treatment=="flg"] <- "flg22"
D$Date <- date

date <- "2020-09-29"
E <- read_excel(paste0(home, date, ".xlsx")) %>% mutate(Stomate=rep(1:(length(Length)/2), each=2)) %>% s
elect(-ImageCounter, -Angle) %>% pivot_wider(names_from="Parameter", values_from="Length") %>% mutate(ra
tio=W/L)
E$Treatment[E$Treatment=="flg"] <- "flg22"
E$Date <- date

stom <- rbind(A, B, C, D, E)

stom$Treatment <- factor(stom$Treatment, levels=c("mock","flg22","ABA"))
stom$Genotype <- factor(stom$Genotype, levels=c("Col-0","glr","bak1-5"))

stomcount <- stom %>% group_by(Date, Plate, Genotype, Treatment) %>% summarize(count=length(ratio)) %>%
 group_by(Genotype, Treatment) %>% summarize(min=min(count), max=max(count))
```

```
## `summarise()` regrouping output by 'Date', 'Plate', 'Genotype' (override with `.groups` argument)
```

```
## `summarise()` regrouping output by 'Genotype' (override with `.groups` argument)
```

```r
stomcount
```

```
## # A tibble: 9 x 4
## # Groups:   Genotype [3]
##   Genotype Treatment   min   max
##   <fct>    <fct>     <int> <int>
## 1 Col-0    mock         49   178
## 2 Col-0    flg22        48   130
## 3 Col-0    ABA          56   122
## 4 glr      mock         45    99
## 5 glr      flg22        48   131
## 6 glr      ABA          47   103
## 7 bak1-5   mock         50   145
## 8 bak1-5   flg22        36   151
## 9 bak1-5   ABA          77   151
```

plot

```
#facet by experiment/plate, double check all reasonably similar
ggplot(stom, aes(x=Genotype, y=ratio, color=Treatment))+geom_boxplot(aes(group=interaction(Genotype, Tre
atment)), outlier.color=NA)+geom_point(aes(group=interaction(Genotype, Treatment)), position=position_ji
tterdodge(jitter.width=0.1), alpha=0.5)+facet_wrap(~Plate)
```

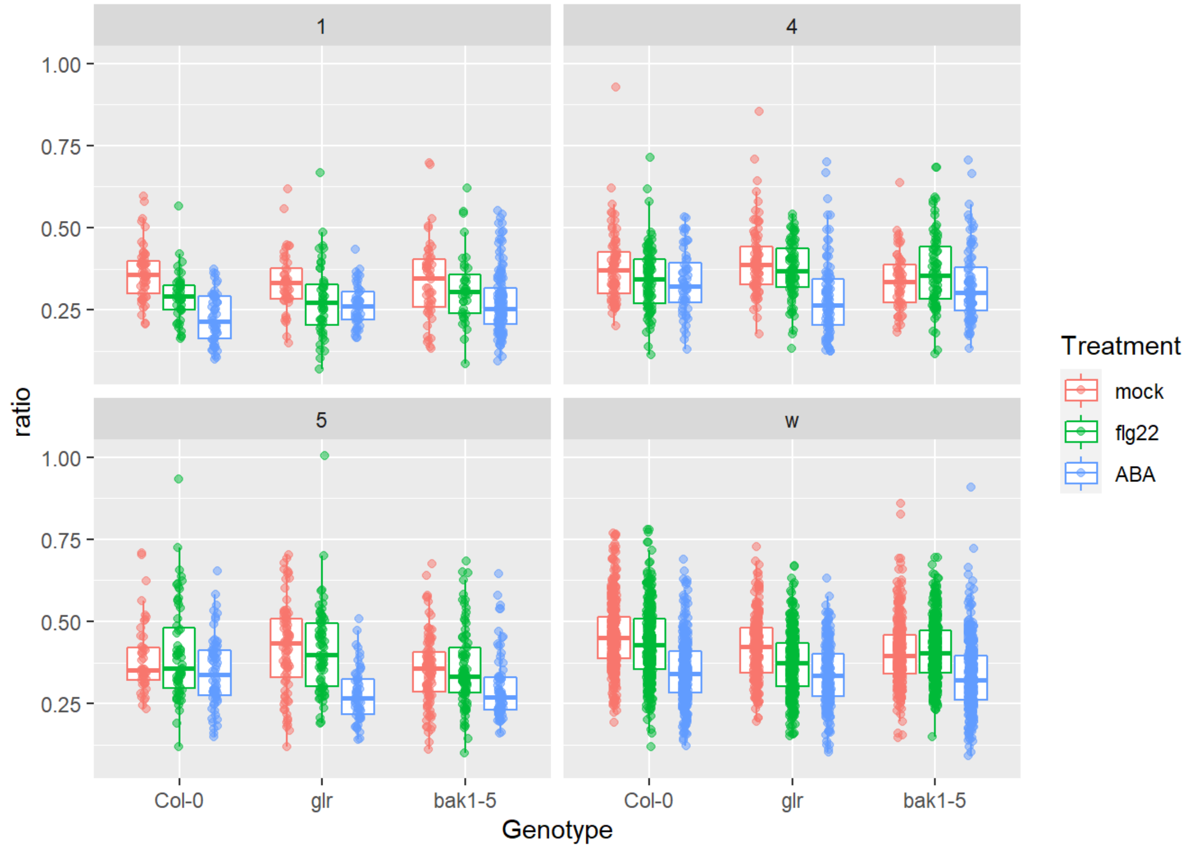

```
italabels <- c("Col-0", expression(italic("glr2.7/2.8/2.9")), expression(italic("bak1-5")))

PanelB <- ggplot(stom, aes(x=Genotype, y=ratio))+
  geom_boxplot(aes(group=interaction(Genotype, Treatment)), outlier.color=NA, size=0.25)+
  geom_point(aes(color=Treatment, group=interaction(Genotype, Treatment)), position=position_jitterdodge
(jitter.width=0.3), alpha=0.2, size=0.5)+
  scale_color_manual(values=mtt)+
  labs(x="",y="Stomatal aperture (width/length)")+
  theme_cowplot()+
  scale_x_discrete(labels=italabels)+
  theme(panel.background = element_blank(), plot.background = element_blank(), panel.border = element_bl
ank(),
        axis.text = element_text(size=6), axis.title = element_text(size=7), legend.box.margin = unit(c(
0,0,-0.3,0), "cm"),
        legend.text=element_text(size=6), legend.title=element_blank(), legend.position = "top",
        plot.margin=unit(c(0,0,-0.3, 0), "cm"))

ggsave(PanelB, filename="PanelB.pdf", height=6, width=6, units="cm", bg="transparent")
```

and statistics

```
#for this model, bulking both plants (two different wells) per genotype/treatment combination
stom.lm <- lm(ratio~Genotype*Treatment+Date/Plate, data=stom)
anova(stom.lm)
```

```
## Analysis of Variance Table
##
## Response: ratio
##                      Df Sum Sq Mean Sq  F value    Pr(>F)
## Genotype              2  0.921  0.4603  43.1727 < 2.2e-16 ***
## Treatment             2  6.645  3.3225 311.6484 < 2.2e-16 ***
## Date                  4  4.665  1.1663 109.4012 < 2.2e-16 ***
## Genotype:Treatment    4  0.416  0.1040   9.7508 7.458e-08 ***
## Date:Plate            1  0.059  0.0593   5.5668   0.01834 *
## Residuals          4898 52.218  0.0107
## ---
## Signif. codes:  0 '***' 0.001 '**' 0.01 '*' 0.05 '.' 0.1 ' ' 1
```

```
#effect of treatment in each genotype
stom.em <- emmeans(stom.lm, dunnett ~ Treatment | Genotype)
```

```
## NOTE: A nesting structure was detected in the fitted model:
##     Plate %in% Date
```

```
stom.em
```

```
## $emmeans
## Genotype = Col-0:
##  Treatment emmean      SE   df lower.CL upper.CL
##  mock       0.425 0.00433 4898    0.417    0.434
##  flg22      0.397 0.00453 4898    0.388    0.406
##  ABA        0.327 0.00456 4898    0.318    0.336
##
## Genotype = glr:
##  Treatment emmean      SE   df lower.CL upper.CL
##  mock       0.405 0.00486 4898    0.396    0.415
##  flg22      0.365 0.00464 4898    0.355    0.374
##  ABA        0.310 0.00505 4898    0.300    0.320
##
## Genotype = bak1-5:
##  Treatment emmean      SE   df lower.CL upper.CL
##  mock       0.376 0.00438 4898    0.367    0.384
##  flg22      0.381 0.00425 4898    0.373    0.389
##  ABA        0.319 0.00385 4898    0.311    0.326
##
## Results are averaged over the levels of: Plate, Date
## Confidence level used: 0.95
##
## $contrasts
## Genotype = Col-0:
##  contrast     estimate      SE   df t.ratio p.value
##  flg22 - mock -0.02824 0.00621 4898  -4.547 <.0001
##  ABA - mock   -0.09883 0.00626 4898 -15.792 <.0001
##
## Genotype = glr:
##  contrast     estimate      SE   df t.ratio p.value
##  flg22 - mock -0.04079 0.00667 4898  -6.119 <.0001
##  ABA - mock   -0.09494 0.00695 4898 -13.668 <.0001
##
## Genotype = bak1-5:
##  contrast     estimate      SE   df t.ratio p.value
##  flg22 - mock  0.00531 0.00604 4898   0.879 0.5818
##  ABA - mock   -0.05718 0.00583 4898  -9.803 <.0001
##
## Results are averaged over the levels of: Plate, Date
## P value adjustment: dunnettx method for 2 tests
```

### Panel D

DC3000 spray infection

```
rm(list=ls())
home <- "D:/OneDrive/Norwich_U/DATA- IR assays/"
monochrome <- c("black","grey","white", "black","grey")

#These two functions read in treatment data, from an excel file. Set up the excel file in the orientatio
n in which the plate(s) went into the camera EXCEPT if there are two plates, stack them vertically not h
orizontally
makelist <- function(filename, sheet){
  worksheet <- sheet
  variable <- read_excel(filename, sheet = worksheet)
  variablelist <- variable[1,]
  for(i in 2:nrow(variable)){
    variablelist <- c(variablelist, variable[i,])
  }
  variablevector <- unlist(variablelist)
  return(variablevector)
}

add_treatments <- function(filename, inheritedros){
  looper <- excel_sheets(filename)
  for(i in 1:length(looper)){
    column <- looper[i]
    columnlist <- makelist(filename, column)
    inheritedros$new <- columnlist
    colnames(inheritedros)[colnames(inheritedros)=="new"] <- column
  }
  return(inheritedros)
}

CollectInfect <- function(date){
  #make empty data frame to hold treatment info
  empty <- data.frame(first=1:96)
  #this calls the two functions above to add one column for each sheet in treatment
  Filled <- add_treatments(paste0(home, date, "_guide.xlsx"), empty)
  Filled$LeafDiscs <- as.numeric(Filled$LeafDiscs)
  Filled <- Filled[Filled$LeafDiscs > 0 & Filled$Genotype != "none",]
  Filled$Date <- date
  #Get colony counts and merge with treatment information
  colonies <- read_excel(paste0(home,date,".xlsx")) %>% unite("Well", c("row", "column"), sep="") %>% se
lect("Plate","Well","original CFU", "drop volume","disc volume")
  Filled <- merge(colonies, Filled, by=c("Plate","Well"))
  #calculate CFU per cm2
  Filled$`CFU/cm2` <-  Filled$`original CFU`/Filled$`drop volume`*Filled$`disc volume`/(Filled$LeafDiscs
*0.2^2*pi)
  Filled$log <- log10(Filled$`CFU/cm2`)
  return(Filled)
}
```

get data

```r
A <- CollectInfect("2020-08-19")
A <- A %>% select(-Bugs) %>% filter(Bacteria=="DC3000")

B <- CollectInfect("2020-09-17")
B <- B %>% select(-Type) %>% filter(Bacteria=="DC3000")

C <- CollectInfect("2020-09-29")
C <- C %>% filter(Bacteria=="DC3000")

All <- rbind(A, B, C) %>% filter(Genotype != "pad4-1")
All$Genotype <- as.factor(All$Genotype) %>% relevel("glr CRISPR 2.6B") %>% relevel("Col-0")
All$Bacteria <- as.factor(All$Bacteria)

print(All[,c(12, 7, 3, 13, 14)])
```

```
##          Date        Genotype original CFU     CFU/cm2      log
## 1  2020-08-19           Col-0       270000   3580986.2 6.554003
## 2  2020-08-19           Col-0      1360000  18037560.2 7.256178
## 3  2020-08-19           Col-0       320000   4244131.8 6.627789
## 4  2020-08-19          bak1-5     10400000 137934284.0 8.139672
## 5  2020-08-19          bak1-5      7900000 104777004.2 8.020266
## 6  2020-08-19          bak1-5      7000000  92840383.5 7.967737
## 7  2020-08-19           Col-0      1340000  17772302.0 7.249744
## 8  2020-08-19           Col-0       960000  12732395.4 7.104910
## 9  2020-08-19           Col-0      2900000  38462444.6 7.585037
## 10 2020-08-19          bak1-5      2300000  30504697.4 7.484367
## 11 2020-08-19          bak1-5      8900000 118039916.1 8.072029
## 12 2020-08-19          bak1-5      6600000  87535218.7 7.942183
## 13 2020-08-19 glr CRISPR 2.6B      2290000  30372068.3 7.482474
## 14 2020-08-19 glr CRISPR 2.6B       370000   4907277.4 6.690841
## 15 2020-08-19 glr CRISPR 2.6B        52000    689671.4 5.838642
## 16 2020-08-19 glr CRISPR 2.6B      2050000  27188969.4 7.434393
## 17 2020-08-19 glr CRISPR 2.6B      1340000  17772302.0 7.249744
## 18 2020-08-19 glr CRISPR 2.6B        94000   1246713.7 6.095767
## 19 2020-09-17          bak1-5       410000   5437793.9 6.735423
## 20 2020-09-17 glr CRISPR 2.6B       430000   5703052.1 6.756107
## 21 2020-09-17 glr CRISPR 2.6B      1080000  14323944.9 7.156063
## 22 2020-09-17           Col-0       146000   1936385.1 6.286992
## 23 2020-09-17          bak1-5      1660000  22016433.8 7.342747
## 24 2020-09-17           Col-0        36000    477464.8 5.678941
## 25 2020-09-17          bak1-5       580000   7692488.9 6.886067
## 26 2020-09-17           Col-0        44000    583568.1 5.766092
## 27 2020-09-17 glr CRISPR 2.6B       143000   1896596.4 6.277975
## 28 2020-09-17          bak1-5      2600000  34483571.0 7.537612
## 29 2020-09-17 glr CRISPR 2.6B       490000   6498826.8 6.812835
## 30 2020-09-17 glr CRISPR 2.6B       520000   6896714.2 6.838642
## 31 2020-09-17           Col-0        28000    371361.5 5.569797
## 32 2020-09-17          bak1-5      2100000  27852115.0 7.444858
## 33 2020-09-17           Col-0        75000    994718.4 5.997700
## 34 2020-09-17          bak1-5      2300000  30504697.4 7.484367
## 35 2020-09-17           Col-0        64000    848826.4 5.928819
## 36 2020-09-17 glr CRISPR 2.6B       210000   2785211.5 6.444858
## 37 2020-09-29           Col-0       139000   1843544.8 6.265654
## 38 2020-09-29           Col-0       165000   2188380.5 6.340123
## 39 2020-09-29           Col-0        91000   1206925.0 6.081680
## 40 2020-09-29 glr CRISPR 2.6B        13000    172417.9 5.236582
## 41 2020-09-29          bak1-5       104000   1379342.8 6.139672
## 42 2020-09-29 glr CRISPR 2.6B        99000   1313028.3 6.118274
## 43 2020-09-29          bak1-5       300000   3978873.6 6.599760
## 44 2020-09-29 glr CRISPR 2.6B        79000   1047770.0 6.020266
## 45 2020-09-29          bak1-5       360000   4774648.3 6.678941
## 46 2020-09-29           Col-0       112000   1485446.1 6.171857
## 47 2020-09-29           Col-0        54000    716197.2 5.855033
## 48 2020-09-29           Col-0       530000   7029343.3 6.846915
## 49 2020-09-29 glr CRISPR 2.6B       580000   7692488.9 6.886067
## 50 2020-09-29          bak1-5       780000  10345071.3 7.014733
## 51 2020-09-29 glr CRISPR 2.6B        58000    769248.9 5.886067
## 52 2020-09-29          bak1-5        74000    981455.5 5.991871
## 53 2020-09-29 glr CRISPR 2.6B       640000   8488263.6 6.928819
## 54 2020-09-29          bak1-5       550000   7294601.6 6.863002
```

plot it

```
mylabs <- c("Col-0", expression(italic("glr2.7/2.8/2.9")), expression(italic("bak1-5")))
PanelC <- ggplot(All, aes(x=Genotype, y=log, group=Genotype))+
  geom_boxplot(outlier.color=NA, size=0.25) +
  geom_point(position=position_jitter(height=0, width=0.2), aes(shape=as.factor(Date), fill=Genotype), s
ize=2, alpha=0.8) +
  labs(x="",y=expression("log"[10]*"(CFU\U2022"*cm^-2*')'), title="") +
  theme_cowplot() +
  scale_fill_manual(values=monochrome, labels=mylabs) +
  scale_x_discrete(labels=mylabs)+
  scale_shape_manual(values=c(21,22,23,24))+
  theme(legend.position = "none", panel.background = element_blank(), plot.background = element_blank(),
        panel.border = element_blank(), strip.text.x = element_blank(),
        axis.text.x = element_text(size=6, angle=45, hjust=1, vjust=1),
        axis.text.y = element_text(size=6), title=element_text(size=10),
        axis.title = element_text(size=7),
        plot.margin=unit(c(-0.25, 0, -0.25, 0), "cm"))
PanelC
```

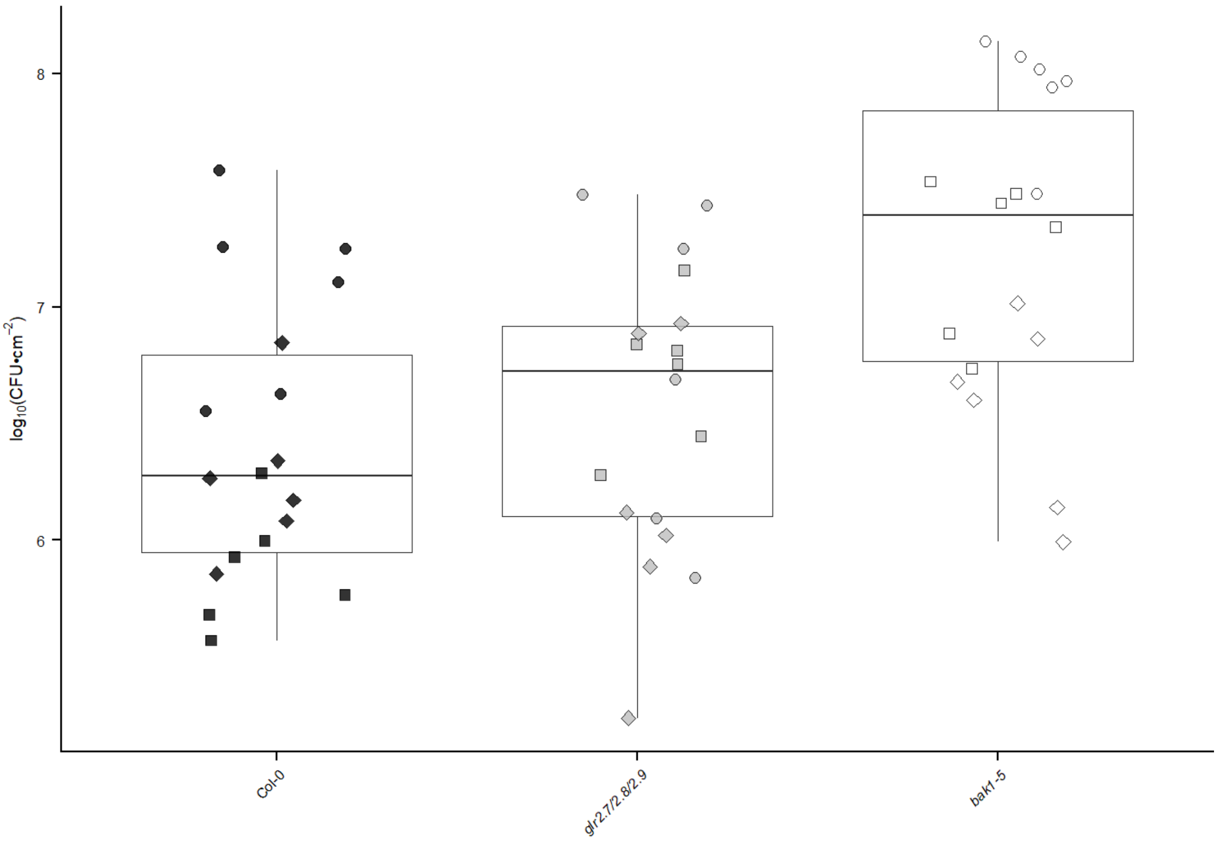


```
ggsave(PanelC, file="PanelC.pdf", height=7.2, width=4, units="cm", bg="transparent")
```

stats

```
All %>% group_by(Genotype) %>% count()
```

```
## # A tibble: 3 x 2
## # Groups:   Genotype [3]
##   Genotype          n
##   <fct>         <int>
## 1 Col-0            18
## 2 glr CRISPR 2.6B    18
## 3 bak1-5           18
```

```
DC3000.lm <- lm(log~Genotype+Date/Tray, data=All[All$log >0,])
anova(DC3000.lm)
```

```
## Analysis of Variance Table
##
## Response: log
##           Df  Sum Sq Mean Sq F value    Pr(>F)
## Genotype   2  7.1836  3.5918 14.9446 9.002e-06 ***
## Date       2  8.3377  4.1689 17.3456 2.142e-06 ***
## Date:Tray  1  0.1032  0.1032  0.4293    0.5154
## Residuals 48 11.5364  0.2403
## ---
## Signif. codes:  0 '***' 0.001 '**' 0.01 '*' 0.05 '.' 0.1 ' ' 1
```

```
#compare all genotypes to each other, independently for each pretreatment
DC3000.em <- emmeans(DC3000.lm, dunnett ~ Genotype)
DC3000.em
```

```
## $emmeans
##  Genotype        emmean SE df asymp.LCL asymp.UCL
##  Col-0           nonEst NA NA        NA        NA
##  glr CRISPR 2.6B nonEst NA NA        NA        NA
##  bak1-5          nonEst NA NA        NA        NA
##
## Results are averaged over the levels of: Date
## Confidence level used: 0.95
##
## $contrasts
##  contrast                  estimate    SE df t.ratio p.value
##  glr CRISPR 2.6B - (Col-0)    0.166 0.163 48 1.016   0.5002
##  (bak1-5) - (Col-0)           0.843 0.163 48 5.160   <.0001
##
## Results are averaged over the levels of: Date
## P value adjustment: dunnettx method for 2 tests
```

### Panel E

Cor- spray infection

```
AB <- CollectInfect("2020-08-13")
A <- AB %>% filter(Chamber=="17") %>% mutate(rep="A") %>% select(-Bugs)

B <- AB %>% filter(Chamber=="18") %>% mutate(rep="B") %>% select(-Bugs)

C <- CollectInfect("2020-10-01") %>% mutate(rep="C")

All <- rbind(A, B, C) %>% filter(Genotype != "pad4-1")
All$Genotype <- as.factor(All$Genotype) %>% relevel("glr CRISPR 2.6B") %>% relevel("Col-0")
All$Bacteria <- as.factor(All$Bacteria)

print(All[,c(15, 7, 3, 13, 14)])
```

```
##    rep        Genotype original CFU     CFU/cm2      log
## 1    A          bak1-5      1180000 15650236.07 7.194521
## 2    A          bak1-5        31000   411150.27 5.614001
## 3    A          bak1-5        15000   198943.68 5.298730
## 4    A          bak1-5        18800   249342.74 5.396797
## 5    A          bak1-5       790000 10477700.42 7.020266
## 6    A          bak1-5       206000  2732159.86 6.436506
## 7    A           Col-0        14400   190985.93 5.281001
## 8    A           Col-0        20300   269237.11 5.430135
## 9    A           Col-0       132000  1750704.37 6.243213
## 10   A           Col-0        18500   245363.87 5.389811
## 11   A           Col-0        29000   384624.45 5.585037
## 12   A           Col-0         1300    17241.79 4.236582
## 13   A glr CRISPR 2.6B         9300   123345.08 5.091122
## 14   A glr CRISPR 2.6B        28700   380645.57 5.580521
## 15   A glr CRISPR 2.6B         6200    82230.05 4.915031
## 16   A glr CRISPR 2.6B        25600   339530.55 5.530879
## 17   A glr CRISPR 2.6B        10100   133955.41 5.126960
## 18   A glr CRISPR 2.6B         7200    95492.97 4.979971
## 19   B          bak1-5        10000   132629.12 5.122639
## 20   B          bak1-5        17200   228122.09 5.358167
## 21   B          bak1-5        88000  1167136.25 6.067122
## 22   B          bak1-5        82000  1087558.78 6.036453
## 23   B          bak1-5       400000  5305164.77 6.724699
## 24   B          bak1-5        15200   201596.26 5.304482
## 25   B           Col-0        31000   411150.27 5.614001
## 26   B           Col-0         5600    74272.31 4.870827
## 27   B           Col-0         7400    98145.55 4.991871
## 28   B           Col-0         8400   111408.46 5.046918
## 29   B           Col-0        22000   291784.06 5.465062
## 30   B           Col-0        45000   596831.04 5.775851
## 31   B glr CRISPR 2.6B         4700    62335.69 4.794737
## 32   B glr CRISPR 2.6B         5800    76924.89 4.886067
## 33   B glr CRISPR 2.6B         6000    79577.47 4.900790
## 34   B glr CRISPR 2.6B        18700   248016.45 5.394480
## 35   B glr CRISPR 2.6B        17200   228122.09 5.358167
## 36   B glr CRISPR 2.6B        11800   156502.36 5.194521
## 37   C glr CRISPR 2.6B        34000   450939.01 5.654118
## 38   C          bak1-5        14500   192312.22 5.284007
## 39   C glr CRISPR 2.6B        16000   212206.59 5.326759
## 40   C          bak1-5       260000  3448357.10 6.537612
## 41   C          bak1-5       340000  4509390.05 6.654118
## 42   C glr CRISPR 2.6B         2900    38462.44 4.585037
## 43   C          bak1-5       440000  5835681.25 6.766092
## 44   C glr CRISPR 2.6B         9400   124671.37 5.095767
## 45   C          bak1-5        17000   225469.50 5.353088
## 46   C glr CRISPR 2.6B        10000   132629.12 5.122639
## 47   C          bak1-5       230000  3050469.74 6.484367
## 48   C glr CRISPR 2.6B        46000   610093.95 5.785397
## 49   C           Col-0        23000   305046.97 5.484367
## 50   C           Col-0         7900   104777.00 5.020266
## 51   C           Col-0        13800   183028.18 5.262518
## 52   C           Col-0         7000    92840.38 4.967737
## 53   C           Col-0         3200    42441.32 4.627789
## 54   C           Col-0        68000   901878.01 5.955148
```

plot it

```
mylabs <- c("Col-0", expression(italic("glr2.7/2.8/2.9")), expression(italic("bak1-5")))
PanelD <- ggplot(All, aes(x=Genotype, y=log, group=Genotype))+
  geom_boxplot(outlier.color=NA, size=0.25) +
  geom_point(position=position_jitter(height=0, width=0.2), aes(shape=as.factor(rep), fill=Genotype), si
ze=2, alpha=0.8) +
  labs(x="",y=expression("log"[10]*"(CFU\U2022"*cm^-2*')'), title="") +
  theme_cowplot() +
  scale_fill_manual(values=monochrome, labels=mylabs) +
  scale_x_discrete(labels=mylabs)+
  scale_shape_manual(values=c(21,22,23,24))+
  theme(legend.position = "none", panel.background = element_blank(), plot.background = element_blank(),
        panel.border = element_blank(), strip.text.x = element_blank(),
        axis.text.x = element_text(size=6, angle=45, hjust=1, vjust=1),
        axis.text.y = element_text(size=6), title=element_text(size=10),
        axis.title = element_text(size=7),
        plot.margin=unit(c(-0.25, 0, -0.25, 0), "cm"))
PanelD
```

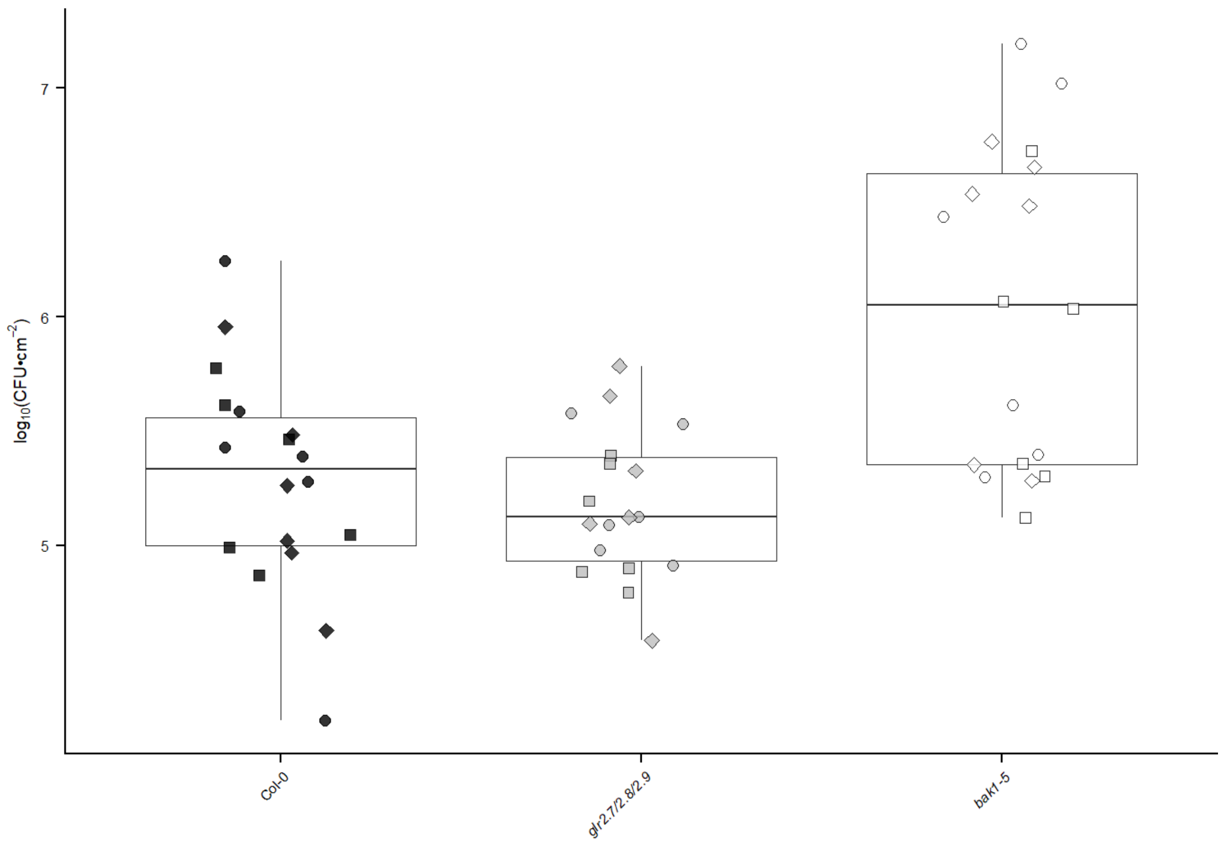


```
ggsave(PanelD, file="PanelD.pdf", height=7.2, width=4, units="cm", bg="transparent")
```

stats

```
All %>% group_by(Genotype) %>% count()
```

```
## # A tibble: 3 x 2
## # Groups:   Genotype [3]
##   Genotype          n
##   <fct>         <int>
## 1 Col-0            18
## 2 glr CRISPR 2.6B  18
## 3 bak1-5           18
```

```
Cor.lm <- lm(log~Genotype+rep, data=All[All$log >0,])
anova(Cor.lm)
```

```
## Analysis of Variance Table
##
## Response: log
##           Df  Sum Sq Mean Sq F value  Pr(>F)
## Genotype   2  7.7490  3.8745 13.9514 1.6e-05 ***
## rep        2  0.3958  0.1979  0.7126  0.4954
## Residuals 49 13.6079  0.2777
## ---
## Signif. codes:  0 '***' 0.001 '**' 0.01 '*' 0.05 '.' 0.1 ' ' 1
```

```
#compare all genotypes to each other, independently for each pretreatment
Cor.em <- emmeans(Cor.lm, dunnett ~ Genotype)
Cor.em
```

```
## $emmeans
##  Genotype        emmean    SE df lower.CL upper.CL
##  Col-0             5.29 0.124 49     5.04     5.54
##  glr CRISPR 2.6B   5.18 0.124 49     4.93     5.43
##  bak1-5            6.04 0.124 49     5.79     6.29
##
## Results are averaged over the levels of: rep
## Confidence level used: 0.95
##
## $contrasts
##  contrast                   estimate    SE df t.ratio p.value
##  glr CRISPR 2.6B - (Col-0)   -0.107 0.176 49 -0.609  0.7588
##  (bak1-5) - (Col-0)           0.745 0.176 49  4.240  0.0002
##
## Results are averaged over the levels of: rep
## P value adjustment: dunnettx method for 2 tests
```

### Figure S8

Marta Bjornson

```
knitr::opts_chunk$set(echo = TRUE)
library(tidyverse)
library(readxl)
library(gplots)# for heatmap.2
library(viridis)
```

### Panel A

Heatmap of avg expression in various tissues, from GeneVestigator

```
tissue <- read_excel("Tissue-specificExpression_R.xlsx") %>% column_to_rownames("...1") %>% as.matrix()
```

```
## New names:
## * `` -> ...1
```

```
pdf(file="FigureS7.pdf")
heatmap.2(tissue, Colv = F, Rowv=F, dendrogram = "none", density.info = "none", scale="row", trace="non
e", breaks=seq(-2,2, length.out=257), col=viridis(256), srtCol = 45, margins= c(10,20))
```

```
## Warning in heatmap.2(tissue, Colv = F, Rowv = F, dendrogram = "none",
## density.info = "none", : Using scale="row" or scale="column" when breaks
## arespecified can produce unpredictable results.Please consider using only one or
## the other.
```

```
dev.off()
```

```
## png
##   2
```

```
png(file="FigureS7.png")
heatmap.2(tissue, Colv = F, Rowv=F, dendrogram = "none", density.info = "none", scale="row", trace="non
e", breaks=seq(-2,2, length.out=257), col=viridis(256), srtCol = 45, margins= c(10,20))
```

```
## Warning in heatmap.2(tissue, Colv = F, Rowv = F, dendrogram = "none",
## density.info = "none", : Using scale="row" or scale="column" when breaks
## arespecified can produce unpredictable results.Please consider using only one or
## the other.
```

```
dev.off()
```

```
## png
##   2
```

### Tables S1, S2, S3, S5

Marta Bjornson

### Tables S1 & S2

L2FC and adjusted p for every gene in every condition.

```
#From results objects, get L2FC and p values for each gene, each treatment/genotype/time combination rel
ative to own time 0
Findl2FC <- function(TrtTime){
  filepath=paste0(home,"ResultsObjects/",TrtTime,"_results")
  load(filepath)
  res.df <- as.data.frame(res)
  FCs <- subset(res.df, select = "log2FoldChange")
  FCs$log2FoldChange <- as.numeric(FCs$log2FoldChange)
  names(FCs) <- TrtTime
  gsub("ch8","CO8", names(FCs))
  gsub("LPS","3-OH-FA", names(FCs))
  return(FCs)
}

Findp<- function(TrtTime){
  filepath=paste0(home,"ResultsObjects/",TrtTime,"_results")
  load(filepath)
  res.df <- as.data.frame(res)
  p <- subset(res.df, select = "padj")
  p$padj <- as.numeric(p$padj)
  names(p) <- TrtTime
  gsub("ch8","CO8", names(p))
  gsub("LPS","3-OH-FA", names(p))
  return(p)
}

PAMPTime <- c("Col_flg22_005","Col_flg22_010","Col_flg22_030","Col_flg22_090","Col_flg22_180",
              "fls2c_flg22_005","fls2c_flg22_010","fls2c_flg22_030","fls2c_flg22_090","fls2c_flg22_180",
              "Col_elf18_005","Col_elf18_010","Col_elf18_030","Col_elf18_090","Col_elf18_180",
              "efr-1_elf18_005","efr-1_elf18_010","efr-1_elf18_030","efr-1_elf18_090","efr-1_elf18_180",
              "Col_Pep1_005","Col_Pep1_010","Col_Pep1_030","Col_Pep1_090","Col_Pep1_180",
              "pepr1_2_Pep1_005","pepr1_2_Pep1_010","pepr1_2_Pep1_030","pepr1_2_Pep1_090","pepr1_2_Pep1_
180",
              "Col_nlp20_005","Col_nlp20_010","Col_nlp20_030","Col_nlp20_090","Col_nlp20_180",
              "rlp23-1_nlp20_005","rlp23-1_nlp20_010","rlp23-1_nlp20_030","rlp23-1_nlp20_090","rlp23-1_n
lp20_180",
              "Col_OGs_005","Col_OGs_010","Col_OGs_030","Col_OGs_090","Col_OGs_180",
              "Col_mock_005","Col_mock_010","Col_mock_030","Col_mock_090","Col_mock_180",
              "Col_ch8_005","Col_ch8_010","Col_ch8_030","Col_ch8_090","Col_ch8_180",
              "lyk4_5_ch8_005","lyk4_5_ch8_010","lyk4_5_ch8_030","lyk4_5_ch8_090","lyk4_5_ch8_180",
              "Col_LPS_005","Col_LPS_010","Col_LPS_030","Col_LPS_090","Col_LPS_180",
              "sd1-29_LPS_005","sd1-29_LPS_010","sd1-29_LPS_030","sd1-29_LPS_090","sd1-29_LPS_180")
log2FC.list <- lapply(PAMPTime, Findl2FC)
log2FC.df <- do.call("cbind", log2FC.list) %>% rownames_to_column("AGI")
nrow(log2FC.df)
```

```
## [1] 26397
```

```
p.list <- lapply(PAMPTime, Findp)
p.df <- do.call("cbind", p.list) %>% rownames_to_column("AGI")
nrow(p.df)
```

```
## [1] 26397
```

```
write_excel_csv(log2FC.df, "TableS1_L2FC.csv")
write_excel_csv(p.df, "TableS2_p.csv")
```

#Table S3

970 genes induced by all elicitors, with some extra information

```
rm(list=ls())
home <- "D:/OneDrive/Norwich_Z/Overall/CleanedData/"

#get list of genes significantly upregulated specifically in wt at each time
wt_sig_up <- function(wt, mut){
  #first find the genes upregulatd in mutant
  mut.file=paste0(home,"ResultsObjects/",mut,"_results")
  load(mut.file)
  mut.df <- as.data.frame(res)

  mut.up <- mut.df %>% filter(padj<0.05)  %>%  filter(log2FoldChange > 1) %>% rownames_to_column("ATG")

  #then the genes upregulated in  wt, extra step filter out the ones also upregulated in control
  wt.file=paste0(home,"ResultsObjects/",wt,"_results")
  load(wt.file)
  wt.df <- as.data.frame(res)

  wt.sig <- wt.df %>% rownames_to_column("ATG") %>%
    filter(padj<0.05)  %>%
    filter(log2FoldChange > 1) %>%
    filter(!ATG %in% mut.up$ATG)

  #reduce to L2FC and return
  wt.FCs <- subset(wt.sig, select = c("ATG","log2FoldChange"))
  wt.FCs$log2FoldChange <- as.numeric(wt.FCs$log2FoldChange)
  names(wt.FCs) <- c("ATG", wt)
  return(wt.FCs)
}

#list of treatment results objects
PAMPTime <- c("Col_LPS_005","Col_LPS_010","Col_LPS_030","Col_LPS_090","Col_LPS_180",
              "Col_ch8_005","Col_ch8_010","Col_ch8_030","Col_ch8_090","Col_ch8_180",
              "Col_elf18_005","Col_elf18_010","Col_elf18_030","Col_elf18_090","Col_elf18_180",
              "Col_flg22_005","Col_flg22_010","Col_flg22_030","Col_flg22_090","Col_flg22_180",
              "Col_nlp20_005","Col_nlp20_010","Col_nlp20_030","Col_nlp20_090","Col_nlp20_180",
              "Col_OGs_005","Col_OGs_010","Col_OGs_030","Col_OGs_090","Col_OGs_180",
              "Col_Pep1_005","Col_Pep1_010","Col_Pep1_030","Col_Pep1_090","Col_Pep1_180")

#corresponding(!) list of control results objects
mutPAMPTime <- c("sd1-29_LPS_005","sd1-29_LPS_010","sd1-29_LPS_030","sd1-29_LPS_090","sd1-29_LPS_180",
                 "lyk4_5_ch8_005","lyk4_5_ch8_010","lyk4_5_ch8_030","lyk4_5_ch8_090","lyk4_5_ch8_180",
                 "efr-1_elf18_005","efr-1_elf18_010","efr-1_elf18_030","efr-1_elf18_090","efr-1_elf18_18
0",
                 "fls2c_flg22_005","fls2c_flg22_010","fls2c_flg22_030","fls2c_flg22_090","fls2c_flg22_18
0",
                 "rlp23-1_nlp20_005","rlp23-1_nlp20_010","rlp23-1_nlp20_030","rlp23-1_nlp20_090","rlp23-
1_nlp20_180",
                 "Col_mock_005","Col_mock_010","Col_mock_030","Col_mock_090","Col_mock_180",
                 "pepr1_2_Pep1_005","pepr1_2_Pep1_010","pepr1_2_Pep1_030","pepr1_2_Pep1_090","pepr1_2_Pe
p1_180")

#Run the function on each treatment/control together
log2FClist <- mapply(wt_sig_up, PAMPTime, mutPAMPTime,SIMPLIFY = F)
#combine into one data frame, and set log2FC to 0 for treatment/time combinations not significantly indu
ced
log2FCdf <- Reduce(function(x,y) merge(x, y, all=T), log2FClist)
log2FCdf[is.na(log2FCdf)] <- 0
nrow(log2FCdf)
```

```
## [1] 5822
```

reduce to genes significantly upregulated in at least one time for all seven (expect 997)

```
AllPAMPs <- log2FCdf %>% gather(-"ATG", key=Sample, value=L2FC) %>% separate(Sample, c("Geno","elicitor"
,"time"), sep="_") %>% group_by(elicitor, ATG) %>% filter(sum(L2FC)>0)  %>% ungroup() %>% group_by(ATG,
 time) %>% add_count() %>% filter(n>6) %>% unite(Sample, c("elicitor","time")) %>% spread(key=Sample, va
lue=L2FC) %>% ungroup() %>% as.data.frame() %>% dplyr::select(-c("Geno","n"))

#clean up some nomenclature, order
names(AllPAMPs) <- gsub("LPS", "3-OH-FA", names(AllPAMPs))
names(AllPAMPs) <- gsub("ch8", "CO8", names(AllPAMPs))

AllPAMPs <- AllPAMPs[,c(1, 12:16, 7:11, 32:36, 22:31, 2:6, 17:21)]

nrow(AllPAMPs)
```

```
## [1] 970
```

Some genes only seem highly induced because they have few starting counts - add this information so it can be judged

```
load(paste0(home, "cleaned_ddsT"))

counts <- counts(ddsT, normalized=T)
onlytime0 <- counts %>% as.data.frame() %>% dplyr::select(matches("000"))
base_counts <- rowSums(onlytime0)/52
tomerge <- rownames_to_column(as.data.frame(base_counts), "ATG")
median(tomerge$base_counts)
```

```
## [1] 78.48919
```

```
mean(tomerge$base_counts)
```

```
## [1] 323.1574
```

```
AllPAMPs <- merge(tomerge, AllPAMPs, by="ATG", all.y=T)
```

Next: get information on abiotic stress response, starting with AtGenExpress: great treatments but microarray

```
#From normalized intensity values, calculate L2FC by probe for AtGen Express. Remove oxidative stress co
ndition
library(Biobase)
library(limma)
```

```
##
## Attaching package: 'limma'
```

```
## The following object is masked from 'package:DESeq2':
##
##     plotMA
```

```
## The following object is masked from 'package:BiocGenerics':
##
##     plotMA
```

```
data <- read.delim(paste0(home, "/wtonly/Analysis-AtGenExpress/AtGenExpress_expression_byprobe.txt"), he
ader=T, row.names=1)
eset<-new("ExpressionSet", exprs=as.matrix(data))
fordesign <- read.delim(paste0(home, "/wtonly/Analysis-AtGenExpress/AtGenExpress_expression_byprobe.txt"
), header=T, row.names=1, check.names=F)
f <- factor(colnames(fordesign))
design <- model.matrix(~0+f)
colnames(design) <- levels(f)
fit <- lmFit(eset, design)
Time0 <- makeContrasts(Cold_0.5_shoot-Control_0_shoot, Cold_1_shoot-Control_0_shoot, Cold_3_shoot-Contro
l_0_shoot,
                       Osmotic_0.5_shoot-Control_0_shoot, Osmotic_1_shoot-Control_0_shoot, Osmotic_3_sho
ot-Control_0_shoot,
                       Salt_0.5_shoot-Control_0_shoot, Salt_1_shoot-Control_0_shoot, Salt_3_shoot-Contro
l_0_shoot,
                       Drought_0.25_shoot-Control_0_shoot, Drought_0.5_shoot-Control_0_shoot, Drought_1_
shoot-Control_0_shoot, Drought_3_shoot-Control_0_shoot,
                       Genotoxic_0.5_shoot-Control_0_shoot, Genotoxic_1_shoot-Control_0_shoot, Genotoxic
_3_shoot-Control_0_shoot,
                       UVB_0.25_shoot-Control_0_shoot, UVB_0.5_shoot-Control_0_shoot, UVB_1_shoot-Contro
l_0_shoot, UVB_3_shoot-Control_0_shoot,
                       Wounding_0.25_shoot-Control_0_shoot,Wounding_0.5_shoot-Control_0_shoot, Wounding_
1_shoot-Control_0_shoot, Wounding_3_shoot-Control_0_shoot,
                       Heat_0.25_shoot-Control_0_shoot,Heat_0.5_shoot-Control_0_shoot, Heat_1_shoot-Cont
rol_0_shoot, Heat_3_shoot-Control_0_shoot,levels=design)
Time0fit <- contrasts.fit(fit, Time0)
Time0fit <- eBayes(Time0fit)

mock <- makeContrasts(Cold_0.5_shoot-Control_0.5_shoot, Cold_1_shoot-Control_1_shoot, Cold_3_shoot-Contr
ol_3_shoot,
                       Osmotic_0.5_shoot-Control_0.5_shoot, Osmotic_1_shoot-Control_1_shoot, Osmotic_3_s
hoot-Control_3_shoot,
                       Salt_0.5_shoot-Control_0.5_shoot, Salt_1_shoot-Control_1_shoot, Salt_3_shoot-Cont
rol_3_shoot,
                       Drought_0.25_shoot-Control_0.25_shoot, Drought_0.5_shoot-Control_0.5_shoot, Droug
ht_1_shoot-Control_1_shoot, Drought_3_shoot-Control_3_shoot,
                       Genotoxic_0.5_shoot-Control_0.5_shoot, Genotoxic_1_shoot-Control_1_shoot, Genotox
ic_3_shoot-Control_3_shoot,
                       UVB_0.25_shoot-Control_0.25_shoot, UVB_0.5_shoot-Control_0.5_shoot, UVB_1_shoot-C
ontrol_1_shoot, UVB_3_shoot-Control_3_shoot,
                       Wounding_0.25_shoot-Control_0.25_shoot,Wounding_0.5_shoot-Control_0.5_shoot, Woun
ding_1_shoot-Control_1_shoot, Wounding_3_shoot-Control_3_shoot,
                       Heat_0.25_shoot-Control_0.25_shoot,Heat_0.5_shoot-Control_0.5_shoot, Heat_1_shoot
-Control_1_shoot, Heat_3_shoot-Control_3_shoot,levels=design)
mockfit <- contrasts.fit(fit, mock)
mockfit <- eBayes(mockfit)

condition <- c("Cold_0-5_shoot", "Cold_1_shoot", "Cold_3_shoot",
                       "Osmotic_0-5_shoot", "Osmotic_1_shoot", "Osmotic_3_shoot",
                       "Salt_0-5_shoot", "Salt_1_shoot", "Salt_3_shoot",
                       "Drought_0-25_shoot", "Drought_0-5_shoot", "Drought_1_shoot", "Drought_3_shoot",
                       "Genotoxic_0-5_shoot", "Genotoxic_1_shoot", "Genotoxic_3_shoot",
                       "UVB_0-25_shoot", "UVB_0-5_shoot", "UVB_1_shoot", "UVB_3_shoot",
                       "Wounding_0-25_shoot","Wounding_0-5_shoot", "Wounding_1_shoot", "Wounding_3_shoo
t",
                       "Heat_0-25_shoot","Heat_0-5_shoot", "Heat_1_shoot", "Heat_3_shoot")

Time0table <- topTable(Time0fit, coef=1, adjust="BH", n=Inf)
Time0table <- Time0table[Time0table$logFC>1,]
Time0table <- Time0table[Time0table$adj.P.Val<0.05,]
Time0table$probe <- rownames(Time0table)
Time0table <- Time0table[,c("logFC", "probe")]
```

```
mocktable <- topTable(mockfit, coef=1, adjust="BH", n=Inf)
mocktable <- mocktable[mocktable$logFC>1,]
mocktable <- mocktable[mocktable$adj.P.Val<0.05,]
Time0table <- Time0table[Time0table$probe %in% rownames(mocktable),]
ShootAbio <- Time0table
colnames(ShootAbio)[1] <- condition[1]

for(i in 2:length(condition)){
  Time0table <- topTable(Time0fit, coef=i, adjust="BH", n=Inf)
  Time0table <- Time0table[Time0table$logFC>1,]
  Time0table <- Time0table[Time0table$adj.P.Val<0.05,]
  Time0table$probe <- rownames(Time0table)
  Time0table <- Time0table[,c("logFC", "probe")]
  colnames(Time0table) <- c(condition[i],"probe")

  mocktable <- topTable(mockfit, coef=i, adjust="BH", n=Inf)
  mocktable <- mocktable[mocktable$logFC>1,]
  mocktable <- mocktable[mocktable$adj.P.Val<0.05,]
  Time0table <- Time0table[Time0table$probe %in% rownames(mocktable),]
  ShootAbio <- merge(ShootAbio, Time0table, by="probe", all=T)
}
```

```
#now do the same for roots
fit <- lmFit(eset, design)
Time0 <- makeContrasts(Cold_0.5_root-Control_0_root, Cold_1_root-Control_0_root, Cold_3_root-Control_0_r
oot,
                       Osmotic_0.5_root-Control_0_root, Osmotic_1_root-Control_0_root, Osmotic_3_root-Co
ntrol_0_root,
                       Salt_0.5_root-Control_0_root, Salt_1_root-Control_0_root, Salt_3_root-Control_0_r
oot,
                       Drought_0.25_root-Control_0_root, Drought_0.5_root-Control_0_root, Drought_1_root
-Control_0_root, Drought_3_root-Control_0_root,
                       Genotoxic_0.5_root-Control_0_root, Genotoxic_1_root-Control_0_root, Genotoxic_3_r
oot-Control_0_root,
                       UVB_0.25_root-Control_0_root, UVB_0.5_root-Control_0_root, UVB_1_root-Control_0_r
oot, UVB_3_root-Control_0_root,
                       Wounding_0.25_root-Control_0_root,Wounding_0.5_root-Control_0_root, Wounding_1_ro
ot-Control_0_root, Wounding_3_root-Control_0_root,
                       Heat_0.25_root-Control_0_root,Heat_0.5_root-Control_0_root, Heat_1_root-Control_0
_root, Heat_3_root-Control_0_root,levels=design)
Time0fit <- contrasts.fit(fit, Time0)
Time0fit <- eBayes(Time0fit)

mock <- makeContrasts(Cold_0.5_root-Control_0.5_root, Cold_1_root-Control_1_root, Cold_3_root-Control_3_
root,
                       Osmotic_0.5_root-Control_0.5_root, Osmotic_1_root-Control_1_root, Osmotic_3_root-
Control_3_root,
                       Salt_0.5_root-Control_0.5_root, Salt_1_root-Control_1_root, Salt_3_root-Control_3
_root,
                       Drought_0.25_root-Control_0.25_root, Drought_0.5_root-Control_0.5_root, Drought_1
_root-Control_1_root, Drought_3_root-Control_3_root,
                       Genotoxic_0.5_root-Control_0.5_root, Genotoxic_1_root-Control_1_root, Genotoxic_3
_root-Control_3_root,
                       UVB_0.25_root-Control_0.25_root, UVB_0.5_root-Control_0.5_root, UVB_1_root-Contro
l_1_root, UVB_3_root-Control_3_root,
                       Wounding_0.25_root-Control_0.25_root,Wounding_0.5_root-Control_0.5_root, Wounding
_1_root-Control_1_root, Wounding_3_root-Control_3_root,
                       Heat_0.25_root-Control_0.25_root,Heat_0.5_root-Control_0.5_root, Heat_1_root-Cont
rol_1_root, Heat_3_root-Control_3_root,levels=design)
mockfit <- contrasts.fit(fit, mock)
mockfit <- eBayes(mockfit)

condition <- c("Cold_0-5_root", "Cold_1_root", "Cold_3_root",
                       "Osmotic_0-5_root", "Osmotic_1_root", "Osmotic_3_root",
                       "Salt_0-5_root", "Salt_1_root", "Salt_3_root",
                       "Drought_0-25_root", "Drought_0-5_root", "Drought_1_root", "Drought_3_root",
                       "Genotoxic_0-5_root", "Genotoxic_1_root", "Genotoxic_3_root",
                       "UVB_0-25_root", "UVB_0-5_root", "UVB_1_root", "UVB_3_root",
                       "Wounding_0-25_root","Wounding_0-5_root", "Wounding_1_root", "Wounding_3_root",
                       "Heat_0-25_root","Heat_0-5_root", "Heat_1_root", "Heat_3_root")

Time0table <- topTable(Time0fit, coef=1, adjust="BH", n=Inf)
Time0table <- Time0table[Time0table$logFC>1,]
Time0table <- Time0table[Time0table$adj.P.Val<0.05,]
Time0table$probe <- rownames(Time0table)
Time0table <- Time0table[,c("logFC", "probe")]

mocktable <- topTable(mockfit, coef=1, adjust="BH", n=Inf)
mocktable <- mocktable[mocktable$logFC>1,]
mocktable <- mocktable[mocktable$adj.P.Val<0.05,]
Time0table <- Time0table[Time0table$probe %in% rownames(mocktable),]
RootAbio <- Time0table
colnames(RootAbio)[1] <- condition[1]

for(i in 2:length(condition)){
```

```
  Time0table <- topTable(Time0fit, coef=i, adjust="BH", n=Inf)
  Time0table <- Time0table[Time0table$logFC>1,]
  Time0table <- Time0table[Time0table$adj.P.Val<0.05,]
  Time0table$probe <- rownames(Time0table)
  Time0table <- Time0table[,c("logFC", "probe")]
  colnames(Time0table) <- c(condition[i],"probe")

  mocktable <- topTable(mockfit, coef=i, adjust="BH", n=Inf)
  mocktable <- mocktable[mocktable$logFC>1,]
  mocktable <- mocktable[mocktable$adj.P.Val<0.05,]
  Time0table <- Time0table[Time0table$probe %in% rownames(mocktable),]
  RootAbio <- merge(RootAbio, Time0table, by="probe", all=T)
}


#remove oxidative, combine root and shoot, taking max
Shootlong <- ShootAbio %>% gather(-probe, key="Sample", value="L2FC") %>% separate("Sample", c("Stress",
"Time","Tissue"), sep="_")
Rootlong <- RootAbio %>% gather(-probe, key="Sample", value="L2FC") %>% separate("Sample", c("Stress","T
ime","Tissue"), sep="_")
Abiolong <- rbind(Shootlong, Rootlong)
Abiocombine <- Abiolong %>% group_by(probe, Stress, Time) %>% summarize(mL2FC = ifelse(all(is.na(L2FC)),
0, max(L2FC, na.rm=T))) %>% unite(Sample, c("Stress", "Time")) %>% spread(key=Sample, value=mL2FC)
```

```
## `summarise()` regrouping output by 'probe', 'Stress' (override with `.groups` argument)
```

```
nrow(RootAbio)
```

```
## [1] 4491
```

```
nrow(ShootAbio)
```

```
## [1] 4358
```

```
nrow(Abiocombine)
```

```
## [1] 6470
```

filter and output tables of genes meeting the following criteria: upregulated compared to time 0, upregulated compared to mock, log2fold change (absolute value) greater than one, and corrected p less than 0.05. Can then look at these like my DESeq2 data sets (upset, etc)

```
#probes to genes: if multiple probes target one gene, take max value. If one probe targets multiple gene
s, remove that probe and all genes
Abiolong <- Abiocombine %>% gather(-probe, key=sample, value=L2FC)

PROBES<- keys(ath1121501.db)
#select(ath1121501.db, c("244901_at","244902_at"), c("TAIR","SYMBOL", "GENENAME"))

#the one:many here doesn't always line up with the probe matches I downloaded from TAIR. since that file
said 2010 and this is Bioconductor which should be updated every six months, going to go with this
MERGEME <- select(ath1121501.db, PROBES, "TAIR")
```

```
## 'select()' returned 1:many mapping between keys and columns
```

```
colnames(MERGEME) <- c("probe","ATG")
#find probes that match to more than one gene
oneprobe_manygenes <- MERGEME %>% group_by(probe) %>% add_count() %>% filter(n>1)

Abioverylong <- merge(Abiolong, MERGEME, by="probe", all.x=T)
Abioverylong <- Abioverylong[!is.na(Abioverylong$ATG),]


#remove probes matching to more than one gene and average results from multiple probes that match to sam
e gene
one2one <- Abioverylong %>% group_by(probe, sample) %>% add_count() %>% filter(n<2) %>% ungroup() %>% gr
oup_by(ATG, sample) %>% summarize(mL2FC=mean(L2FC, na.rm=T))
```

```
## `summarise()` regrouping output by 'ATG' (override with `.groups` argument)
```

```
#get count of abiotic stresses which induce a gene (at any time)
NumberAbio <- one2one %>% separate(sample, c("stress","time"), sep="_") %>% group_by(stress, ATG) %>% fi
lter(sum(mL2FC)>0)  %>% ungroup() %>% group_by(ATG) %>% mutate(count = length(unique(stress))) %>% unite
(Sample, c("stress","time")) %>% ungroup() %>% group_by(ATG) %>% summarize(AtGenExpressNumber=mean(coun
t))
```

```
## `summarise()` ungrouping output (override with `.groups` argument)
```

```
#Add count Abio on to  all PAMPs
AllPAMPs_AtGen <- merge(NumberAbio, AllPAMPs, by="ATG", all.y=T) %>% mutate(AtGenExpressNumber=replace_n
a(AtGenExpressNumber,0))
AllPAMPs_AtGen$AtGenExpressNumber[!AllPAMPs_AtGen$ATG %in% MERGEME$ATG] <- "NotOnArray"
AllPAMPs_AtGen$AtGenExpressNumber[AllPAMPs_AtGen$ATG %in% oneprobe_manygenes$ATG] <- "ProbeHitsMultipleG
enes"
```

further information from the few RNAseq experiments I could find - for this files were assembled from different papers anyway, just going straight from that aggregate file

```
#by gene assign RNAseqAbio: if nothing "none", if 1 "one", if x-y "few", if  (expect ~30 "none" "none")
RNAseq <- read.csv(paste0(home,"/wtonly/Analysis-AtGenExpress/RNAseq_logFC.csv"), header=T)

RNAseqcount <- RNAseq %>% gather(-"ATG", key=Sample, value=L2FC) %>% group_by(ATG) %>% filter(!is.na(L2F
C)) %>% add_count() %>% summarize(AbioRNAseqNumber=mean(n))
```

```
## `summarise()` ungrouping output (override with `.groups` argument)
```

```
AllPAMPs_abio <- merge(RNAseqcount, AllPAMPs_AtGen, by="ATG", all.y=T) %>% mutate(AbioRNAseqNumber=repla
ce_na(AbioRNAseqNumber,0))
```

Add cluster information (Figure 1)

```
D <- AllPAMPs %>% column_to_rownames("ATG") %>% as.matrix() %>% spearman.dist()
#x <- x %>% rownames_to_column(var="ATG")

uptree <- hclust(D)

#After exploratory data analysis, chose 4 clusters
num = 4

cluster <- cutree(uptree, k=num)
cluster.df <- as.data.frame(cluster) %>% rownames_to_column("ATG")

AllPAMPs_prefinal <- merge(cluster.df, AllPAMPs_abio, by="ATG", all.y=T)
```

And finally get descriptions

```
library(org.At.tair.db)
GENES <- AllPAMPs_abio$ATG
info <- select(org.At.tair.db, as.character(GENES), c("TAIR", "SYMBOL","GENENAME"))
```

```
## 'select()' returned 1:many mapping between keys and columns
```

```
info <- info %>% group_by(TAIR) %>% summarize(Name = SYMBOL[1], Description = GENENAME[1])
```

```
## `summarise()` ungrouping output (override with `.groups` argument)
```

```
colnames(info) <- c("ATG","Name","Description")

AllPAMPs_final <- merge(info, AllPAMPs_prefinal, by="ATG",all.y=T)
```

Save

```
write_excel_csv(AllPAMPs_final, "TableS3.csv")
```

### Table S5

136 genes down-regulated by all elicitors, with some extra information, same as table 3 but downregulated

```
rm(list=ls())
home <- "D:/OneDrive/Norwich_Z/Overall/CleanedData/"

#get list of genes significantly upregulated specifically in wt at each time
wt_sig_down <- function(wt, mut){
  #first find the genes upregulatd in mutant
  mut.file=paste0(home,"ResultsObjects/",mut,"_results")
  load(mut.file)
  mut.df <- as.data.frame(res)

  mut.up <- mut.df %>% filter(padj<0.05)  %>%  filter(log2FoldChange < -1) %>% rownames_to_column("ATG")

  #then the genes upregulated in  wt, extra step filter out the ones also upregulated in control
  wt.file=paste0(home,"ResultsObjects/",wt,"_results")
  load(wt.file)
  wt.df <- as.data.frame(res)

  wt.sig <- wt.df %>% rownames_to_column("ATG") %>%
    filter(padj<0.05)  %>%
    filter(log2FoldChange < -1) %>%
    filter(!ATG %in% mut.up$ATG)

  #reduce to L2FC and return
  wt.FCs <- subset(wt.sig, select = c("ATG","log2FoldChange"))
  wt.FCs$log2FoldChange <- as.numeric(wt.FCs$log2FoldChange)
  names(wt.FCs) <- c("ATG", wt)
  return(wt.FCs)
}

#list of treatment results objects
PAMPTime <- c("Col_LPS_005","Col_LPS_010","Col_LPS_030","Col_LPS_090","Col_LPS_180",
              "Col_ch8_005","Col_ch8_010","Col_ch8_030","Col_ch8_090","Col_ch8_180",
              "Col_elf18_005","Col_elf18_010","Col_elf18_030","Col_elf18_090","Col_elf18_180",
              "Col_flg22_005","Col_flg22_010","Col_flg22_030","Col_flg22_090","Col_flg22_180",
              "Col_nlp20_005","Col_nlp20_010","Col_nlp20_030","Col_nlp20_090","Col_nlp20_180",
              "Col_OGs_005","Col_OGs_010","Col_OGs_030","Col_OGs_090","Col_OGs_180",
              "Col_Pep1_005","Col_Pep1_010","Col_Pep1_030","Col_Pep1_090","Col_Pep1_180")

#corresponding(!) list of control results objects
mutPAMPTime <- c("sd1-29_LPS_005","sd1-29_LPS_010","sd1-29_LPS_030","sd1-29_LPS_090","sd1-29_LPS_180",
                 "lyk4_5_ch8_005","lyk4_5_ch8_010","lyk4_5_ch8_030","lyk4_5_ch8_090","lyk4_5_ch8_180",
                 "efr-1_elf18_005","efr-1_elf18_010","efr-1_elf18_030","efr-1_elf18_090","efr-1_elf18_18
0",
                 "fls2c_flg22_005","fls2c_flg22_010","fls2c_flg22_030","fls2c_flg22_090","fls2c_flg22_18
0",
                 "rlp23-1_nlp20_005","rlp23-1_nlp20_010","rlp23-1_nlp20_030","rlp23-1_nlp20_090","rlp23-
1_nlp20_180",
                 "Col_mock_005","Col_mock_010","Col_mock_030","Col_mock_090","Col_mock_180",
                 "pepr1_2_Pep1_005","pepr1_2_Pep1_010","pepr1_2_Pep1_030","pepr1_2_Pep1_090","pepr1_2_Pe
p1_180")

#Run the function on each treatment/control together
log2FClist <- mapply(wt_sig_down, PAMPTime, mutPAMPTime,SIMPLIFY = F)
#combine into one data frame, and set log2FC to 0 for treatment/time combinations not significantly indu
ced
log2FCdf <- Reduce(function(x,y) merge(x, y, all=T), log2FClist)
log2FCdf[is.na(log2FCdf)] <- 0
nrow(log2FCdf)
```

```
## [1] 5157
```

reduce to genes significantly upregulated in at least one time for all seven (expect 997)

```
AllPAMPs <- log2FCdf %>% gather(-"ATG", key=Sample, value=L2FC) %>% separate(Sample, c("Geno","elicitor"
,"time"), sep="_") %>% group_by(elicitor, ATG) %>% filter(sum(L2FC)<0)  %>% ungroup() %>% group_by(ATG,
 time) %>% add_count() %>% filter(n>6) %>% unite(Sample, c("elicitor","time")) %>% spread(key=Sample, va
lue=L2FC) %>% ungroup() %>% as.data.frame() %>% dplyr::select(-c("Geno","n"))

#clean up some nomenclature, order
names(AllPAMPs) <- gsub("LPS", "3-OH-FA", names(AllPAMPs))
names(AllPAMPs) <- gsub("ch8", "CO8", names(AllPAMPs))

AllPAMPs <- AllPAMPs[,c(1, 12:16, 7:11, 32:36, 22:31, 2:6, 17:21)]

nrow(AllPAMPs)
```

```
## [1] 93
```

Some genes only seem highly induced because they have few starting counts - add this information so it can be judged

```
load(paste0(home, "cleaned_ddsT"))

counts <- counts(ddsT, normalized=T)
onlytime0 <- counts %>% as.data.frame() %>% dplyr::select(matches("000"))
base_counts <- rowSums(onlytime0)/52
tomerge <- rownames_to_column(as.data.frame(base_counts), "ATG")
median(tomerge$base_counts)
```

```
## [1] 78.48919
```

```
mean(tomerge$base_counts)
```

```
## [1] 323.1574
```

```
AllPAMPs <- merge(tomerge, AllPAMPs, by="ATG", all.y=T)
```

Next: get information on abiotic stress response, starting with AtGenExpress: great treatments but microarray

```
#From normalized intensity values, calculate L2FC by probe for AtGen Express. Remove oxidative stress co
ndition
library(Biobase)
library(limma)
data <- read.delim(paste0(home, "/wtonly/Analysis-AtGenExpress/AtGenExpress_expression_byprobe.txt"), he
ader=T, row.names=1)
eset<-new("ExpressionSet", exprs=as.matrix(data))
fordesign <- read.delim(paste0(home, "/wtonly/Analysis-AtGenExpress/AtGenExpress_expression_byprobe.txt"
), header=T, row.names=1, check.names=F)
f <- factor(colnames(fordesign))
design <- model.matrix(~0+f)
colnames(design) <- levels(f)
fit <- lmFit(eset, design)
Time0 <- makeContrasts(Cold_0.5_shoot-Control_0_shoot, Cold_1_shoot-Control_0_shoot, Cold_3_shoot-Contro
l_0_shoot,
                       Osmotic_0.5_shoot-Control_0_shoot, Osmotic_1_shoot-Control_0_shoot, Osmotic_3_sho
ot-Control_0_shoot,
                       Salt_0.5_shoot-Control_0_shoot, Salt_1_shoot-Control_0_shoot, Salt_3_shoot-Contro
l_0_shoot,
                       Drought_0.25_shoot-Control_0_shoot, Drought_0.5_shoot-Control_0_shoot, Drought_1_
shoot-Control_0_shoot, Drought_3_shoot-Control_0_shoot,
                       Genotoxic_0.5_shoot-Control_0_shoot, Genotoxic_1_shoot-Control_0_shoot, Genotoxic
_3_shoot-Control_0_shoot,
                       UVB_0.25_shoot-Control_0_shoot, UVB_0.5_shoot-Control_0_shoot, UVB_1_shoot-Contro
l_0_shoot, UVB_3_shoot-Control_0_shoot,
                       Wounding_0.25_shoot-Control_0_shoot,Wounding_0.5_shoot-Control_0_shoot, Wounding_
1_shoot-Control_0_shoot, Wounding_3_shoot-Control_0_shoot,
                       Heat_0.25_shoot-Control_0_shoot,Heat_0.5_shoot-Control_0_shoot, Heat_1_shoot-Cont
rol_0_shoot, Heat_3_shoot-Control_0_shoot,levels=design)
Time0fit <- contrasts.fit(fit, Time0)
Time0fit <- eBayes(Time0fit)

mock <- makeContrasts(Cold_0.5_shoot-Control_0.5_shoot, Cold_1_shoot-Control_1_shoot, Cold_3_shoot-Contr
ol_3_shoot,
                       Osmotic_0.5_shoot-Control_0.5_shoot, Osmotic_1_shoot-Control_1_shoot, Osmotic_3_s
hoot-Control_3_shoot,
                       Salt_0.5_shoot-Control_0.5_shoot, Salt_1_shoot-Control_1_shoot, Salt_3_shoot-Cont
rol_3_shoot,
                       Drought_0.25_shoot-Control_0.25_shoot, Drought_0.5_shoot-Control_0.5_shoot, Droug
ht_1_shoot-Control_1_shoot, Drought_3_shoot-Control_3_shoot,
                       Genotoxic_0.5_shoot-Control_0.5_shoot, Genotoxic_1_shoot-Control_1_shoot, Genotox
ic_3_shoot-Control_3_shoot,
                       UVB_0.25_shoot-Control_0.25_shoot, UVB_0.5_shoot-Control_0.5_shoot, UVB_1_shoot-C
ontrol_1_shoot, UVB_3_shoot-Control_3_shoot,
                       Wounding_0.25_shoot-Control_0.25_shoot,Wounding_0.5_shoot-Control_0.5_shoot, Woun
ding_1_shoot-Control_1_shoot, Wounding_3_shoot-Control_3_shoot,
                       Heat_0.25_shoot-Control_0.25_shoot,Heat_0.5_shoot-Control_0.5_shoot, Heat_1_shoot
-Control_1_shoot, Heat_3_shoot-Control_3_shoot,levels=design)
mockfit <- contrasts.fit(fit, mock)
mockfit <- eBayes(mockfit)

condition <- c("Cold_0-5_shoot", "Cold_1_shoot", "Cold_3_shoot",
                       "Osmotic_0-5_shoot", "Osmotic_1_shoot", "Osmotic_3_shoot",
                       "Salt_0-5_shoot", "Salt_1_shoot", "Salt_3_shoot",
                       "Drought_0-25_shoot", "Drought_0-5_shoot", "Drought_1_shoot", "Drought_3_shoot",
                       "Genotoxic_0-5_shoot", "Genotoxic_1_shoot", "Genotoxic_3_shoot",
                       "UVB_0-25_shoot", "UVB_0-5_shoot", "UVB_1_shoot", "UVB_3_shoot",
                       "Wounding_0-25_shoot","Wounding_0-5_shoot", "Wounding_1_shoot", "Wounding_3_shoo
t",
                       "Heat_0-25_shoot","Heat_0-5_shoot", "Heat_1_shoot", "Heat_3_shoot")

Time0table <- topTable(Time0fit, coef=1, adjust="BH", n=Inf)
Time0table <- Time0table[Time0table$logFC< -1,]
```

```
Time0table <- Time0table[Time0table$adj.P.Val<0.05,]
Time0table$probe <- rownames(Time0table)
Time0table <- Time0table[,c("logFC", "probe")]

mocktable <- topTable(mockfit, coef=1, adjust="BH", n=Inf)
mocktable <- mocktable[mocktable$logFC< -1,]
mocktable <- mocktable[mocktable$adj.P.Val<0.05,]
Time0table <- Time0table[Time0table$probe %in% rownames(mocktable),]
ShootAbio <- Time0table
colnames(ShootAbio)[1] <- condition[1]

for(i in 2:length(condition)){
  Time0table <- topTable(Time0fit, coef=i, adjust="BH", n=Inf)
  Time0table <- Time0table[Time0table$logFC< -1,]
  Time0table <- Time0table[Time0table$adj.P.Val<0.05,]
  Time0table$probe <- rownames(Time0table)
  Time0table <- Time0table[,c("logFC", "probe")]
  colnames(Time0table) <- c(condition[i],"probe")

  mocktable <- topTable(mockfit, coef=i, adjust="BH", n=Inf)
  mocktable <- mocktable[mocktable$logFC< -1,]
  mocktable <- mocktable[mocktable$adj.P.Val<0.05,]
  Time0table <- Time0table[Time0table$probe %in% rownames(mocktable),]
  ShootAbio <- merge(ShootAbio, Time0table, by="probe", all=T)
}
```

```
#now do the same for roots
fit <- lmFit(eset, design)
Time0 <- makeContrasts(Cold_0.5_root-Control_0_root, Cold_1_root-Control_0_root, Cold_3_root-Control_0_r
oot,
                       Osmotic_0.5_root-Control_0_root, Osmotic_1_root-Control_0_root, Osmotic_3_root-Co
ntrol_0_root,
                       Salt_0.5_root-Control_0_root, Salt_1_root-Control_0_root, Salt_3_root-Control_0_r
oot,
                       Drought_0.25_root-Control_0_root, Drought_0.5_root-Control_0_root, Drought_1_root
-Control_0_root, Drought_3_root-Control_0_root,
                       Genotoxic_0.5_root-Control_0_root, Genotoxic_1_root-Control_0_root, Genotoxic_3_r
oot-Control_0_root,
                       UVB_0.25_root-Control_0_root, UVB_0.5_root-Control_0_root, UVB_1_root-Control_0_r
oot, UVB_3_root-Control_0_root,
                       Wounding_0.25_root-Control_0_root,Wounding_0.5_root-Control_0_root, Wounding_1_ro
ot-Control_0_root, Wounding_3_root-Control_0_root,
                       Heat_0.25_root-Control_0_root,Heat_0.5_root-Control_0_root, Heat_1_root-Control_0
_root, Heat_3_root-Control_0_root,levels=design)
Time0fit <- contrasts.fit(fit, Time0)
Time0fit <- eBayes(Time0fit)

mock <- makeContrasts(Cold_0.5_root-Control_0.5_root, Cold_1_root-Control_1_root, Cold_3_root-Control_3_
root,
                       Osmotic_0.5_root-Control_0.5_root, Osmotic_1_root-Control_1_root, Osmotic_3_root-
Control_3_root,
                       Salt_0.5_root-Control_0.5_root, Salt_1_root-Control_1_root, Salt_3_root-Control_3
_root,
                       Drought_0.25_root-Control_0.25_root, Drought_0.5_root-Control_0.5_root, Drought_1
_root-Control_1_root, Drought_3_root-Control_3_root,
                       Genotoxic_0.5_root-Control_0.5_root, Genotoxic_1_root-Control_1_root, Genotoxic_3
_root-Control_3_root,
                       UVB_0.25_root-Control_0.25_root, UVB_0.5_root-Control_0.5_root, UVB_1_root-Contro
l_1_root, UVB_3_root-Control_3_root,
                       Wounding_0.25_root-Control_0.25_root,Wounding_0.5_root-Control_0.5_root, Wounding
_1_root-Control_1_root, Wounding_3_root-Control_3_root,
                       Heat_0.25_root-Control_0.25_root,Heat_0.5_root-Control_0.5_root, Heat_1_root-Cont
rol_1_root, Heat_3_root-Control_3_root,levels=design)
mockfit <- contrasts.fit(fit, mock)
mockfit <- eBayes(mockfit)

condition <- c("Cold_0-5_root", "Cold_1_root", "Cold_3_root",
                       "Osmotic_0-5_root", "Osmotic_1_root", "Osmotic_3_root",
                       "Salt_0-5_root", "Salt_1_root", "Salt_3_root",
                       "Drought_0-25_root", "Drought_0-5_root", "Drought_1_root", "Drought_3_root",
                       "Genotoxic_0-5_root", "Genotoxic_1_root", "Genotoxic_3_root",
                       "UVB_0-25_root", "UVB_0-5_root", "UVB_1_root", "UVB_3_root",
                       "Wounding_0-25_root","Wounding_0-5_root", "Wounding_1_root", "Wounding_3_root",
                       "Heat_0-25_root","Heat_0-5_root", "Heat_1_root", "Heat_3_root")

Time0table <- topTable(Time0fit, coef=1, adjust="BH", n=Inf)
Time0table <- Time0table[Time0table$logFC< -1,]
Time0table <- Time0table[Time0table$adj.P.Val<0.05,]
Time0table$probe <- rownames(Time0table)
Time0table <- Time0table[,c("logFC", "probe")]

mocktable <- topTable(mockfit, coef=1, adjust="BH", n=Inf)
mocktable <- mocktable[mocktable$logFC< -1,]
mocktable <- mocktable[mocktable$adj.P.Val<0.05,]
Time0table <- Time0table[Time0table$probe %in% rownames(mocktable),]
RootAbio <- Time0table
colnames(RootAbio)[1] <- condition[1]

for(i in 2:length(condition)){
```

```
  Time0table <- topTable(Time0fit, coef=i, adjust="BH", n=Inf)
  Time0table <- Time0table[Time0table$logFC< -1,]
  Time0table <- Time0table[Time0table$adj.P.Val<0.05,]
  Time0table$probe <- rownames(Time0table)
  Time0table <- Time0table[,c("logFC", "probe")]
  colnames(Time0table) <- c(condition[i],"probe")

  mocktable <- topTable(mockfit, coef=i, adjust="BH", n=Inf)
  mocktable <- mocktable[mocktable$logFC< -1,]
  mocktable <- mocktable[mocktable$adj.P.Val<0.05,]
  Time0table <- Time0table[Time0table$probe %in% rownames(mocktable),]
  RootAbio <- merge(RootAbio, Time0table, by="probe", all=T)
}


#remove oxidative, combine root and shoot, taking max
Shootlong <- ShootAbio %>% gather(-probe, key="Sample", value="L2FC") %>% separate("Sample", c("Stress",
"Time","Tissue"), sep="_")
Rootlong <- RootAbio %>% gather(-probe, key="Sample", value="L2FC") %>% separate("Sample", c("Stress","T
ime","Tissue"), sep="_")
Abiolong <- rbind(Shootlong, Rootlong)
Abiocombine <- Abiolong %>% group_by(probe, Stress, Time) %>% summarize(mL2FC = ifelse(all(is.na(L2FC)),
0, max(L2FC, na.rm=T))) %>% unite(Sample, c("Stress", "Time")) %>% spread(key=Sample, value=mL2FC)
```

```
## `summarise()` regrouping output by 'probe', 'Stress' (override with `.groups` argument)
```

```
nrow(RootAbio)
```

```
## [1] 3635
```

```
nrow(ShootAbio)
```

```
## [1] 3748
```

```
nrow(Abiocombine)
```

```
## [1] 6041
```

filter and output tables of genes meeting the following criteria: upregulated compared to time 0, upregulated compared to mock, log2fold change (absolute value) greater than one, and corrected p less than 0.05. Can then look at these like my DESeq2 data sets (upset, etc)

```
#probes to genes: if multiple probes target one gene, take max value. If one probe targets multiple gene
s, remove that probe and all genes
Abiolong <- Abiocombine %>% gather(-probe, key=sample, value=L2FC)

PROBES<- keys(ath1121501.db)
#select(ath1121501.db, c("244901_at","244902_at"), c("TAIR","SYMBOL", "GENENAME"))

#the one:many here doesn't always line up with the probe matches I downloaded from TAIR. since that file
said 2010 and this is Bioconductor which should be updated every six months, going to go with this
MERGEME <- select(ath1121501.db, PROBES, "TAIR")
```

```
## 'select()' returned 1:many mapping between keys and columns
```

```
colnames(MERGEME) <- c("probe","ATG")
#find probes that match to more than one gene
oneprobe_manygenes <- MERGEME %>% group_by(probe) %>% add_count() %>% filter(n>1)

Abioverylong <- merge(Abiolong, MERGEME, by="probe", all.x=T)
Abioverylong <- Abioverylong[!is.na(Abioverylong$ATG),]



#remove probes matching to more than one gene and average results from multiple probes that match to sam
e gene
one2one <- Abioverylong %>% group_by(probe, sample) %>% add_count() %>% filter(n<2) %>% ungroup() %>% gr
oup_by(ATG, sample) %>% summarize(mL2FC=mean(L2FC, na.rm=T))
```

```
## `summarise()` regrouping output by 'ATG' (override with `.groups` argument)
```

```
#get count of abiotic stresses which induce a gene (at any time)
NumberAbio <- one2one %>% separate(sample, c("stress","time"), sep="_") %>% group_by(stress, ATG) %>% fi
lter(sum(mL2FC)>0)  %>% ungroup() %>% group_by(ATG) %>% mutate(count = length(unique(stress))) %>% unite
(Sample, c("stress","time")) %>% ungroup() %>% group_by(ATG) %>% summarize(AtGenExpressNumber=mean(coun
t))
```

```
## `summarise()` ungrouping output (override with `.groups` argument)
```

```
#Add count Abio on to  all PAMPs
AllPAMPs_AtGen <- merge(NumberAbio, AllPAMPs, by="ATG", all.y=T) %>% mutate(AtGenExpressNumber=replace_n
a(AtGenExpressNumber,0))
AllPAMPs_AtGen$AtGenExpressNumber[!AllPAMPs_AtGen$ATG %in% MERGEME$ATG] <- "NotOnArray"
AllPAMPs_AtGen$AtGenExpressNumber[AllPAMPs_AtGen$ATG %in% oneprobe_manygenes$ATG] <- "ProbeHitsMultipleG
enes"
```

further information from the few RNAseq experiments I could find - for this files were assembled from different papers anyway, just going straight from that aggregate file

```
#by gene assign RNAseqAbio: if nothing "none", if 1 "one", if x-y "few", if  (expect ~30 "none" "none")
RNAseq <- read.csv(paste0(home,"/wtonly/Analysis-AtGenExpress/RNAseq_logFC.csv"), header=T)

RNAseqcount <- RNAseq %>% gather(-"ATG", key=Sample, value=L2FC) %>% group_by(ATG) %>% filter(!is.na(L2F
C)) %>% add_count() %>% summarize(AbioRNAseqNumber=mean(n))
```

```
## `summarise()` ungrouping output (override with `.groups` argument)
```

```
AllPAMPs_abio <- merge(RNAseqcount, AllPAMPs_AtGen, by="ATG", all.y=T) %>% mutate(AbioRNAseqNumber=repla
ce_na(AbioRNAseqNumber,0))
```

Add cluster information (Figure 1)

```
D <- AllPAMPs %>% column_to_rownames("ATG") %>% as.matrix() %>% spearman.dist()
#x <- x %>% rownames_to_column(var="ATG")

uptree <- hclust(D)

#After exploratory data analysis, chose 3 clusters
num = 3

cluster <- cutree(uptree, k=num)
cluster.df <- as.data.frame(cluster) %>% rownames_to_column("ATG")

AllPAMPs_prefinal <- merge(cluster.df, AllPAMPs_abio, by="ATG", all.y=T)
```

And finally get descriptions

```
library(org.At.tair.db)
GENES <- AllPAMPs_abio$ATG
info <- select(org.At.tair.db, as.character(GENES), c("TAIR", "SYMBOL","GENENAME"))
```

```
## 'select()' returned 1:many mapping between keys and columns
```

```
info <- info %>% group_by(TAIR) %>% summarize(Name = SYMBOL[1], Description = GENENAME[1])
```

```
## `summarise()` ungrouping output (override with `.groups` argument)
```

```
colnames(info) <- c("ATG","Name","Description")

AllPAMPs_final <- merge(info, AllPAMPs_prefinal, by="ATG",all.y=T)
```

Save

```
write_excel_csv(AllPAMPs_final, "TableS5.csv")
```

### Table S6

Marta Bjornson

```
knitr::opts_chunk$set(echo = TRUE)
library(tidyverse)
library(DESeq2)
#forAtGenExpress analysis

library(ath1121501.db)
library(AnnotationDbi)
library(affy)
library(annotate)

library(Biobase)
library(limma)
```

### Distilling the core immunity response gene set

```
rm(list=ls())
home <- "D:/OneDrive/Norwich_Z/Overall/CleanedData/"

#From results objects, get gene lists p<0.05 L2FC >1, and combine into one large file (expect ~6500)
OnlySigl2FC <- function(TrtTime){
  filepath=paste0(home,"ResultsObjects/",TrtTime,"_results")
  load(filepath)
  resdf <- as.data.frame(res)
  nonsig <- subset(resdf, padj >0.05 | padj == "NA")
  smallchange <- subset(resdf, log2FoldChange <  1 & log2FoldChange > -1)
  onlysig <- resdf
  onlysig[rownames(onlysig) %in% rownames(nonsig),] <- "0"
  onlysig[rownames(onlysig) %in% rownames(smallchange),] <- "0"
  sigFCs <- subset(onlysig, select = "log2FoldChange")
  sigFCs$log2FoldChange <- as.numeric(sigFCs$log2FoldChange)
  names(sigFCs) <- TrtTime
  return(sigFCs)
}

PAMPTime <- c("Col_LPS_005","Col_LPS_010","Col_LPS_030","Col_LPS_090","Col_LPS_180","Col_ch8_005","Col_c
h8_010","Col_ch8_030","Col_ch8_090","Col_ch8_180","Col_elf18_005","Col_elf18_010","Col_elf18_030","Col_e
lf18_090","Col_elf18_180","Col_flg22_005","Col_flg22_010","Col_flg22_030","Col_flg22_090","Col_flg22_18
0","Col_nlp20_005","Col_nlp20_010","Col_nlp20_030","Col_nlp20_090","Col_nlp20_180","Col_OGs_005","Col_OG
s_010","Col_OGs_030","Col_OGs_090","Col_OGs_180","Col_Pep1_005","Col_Pep1_010","Col_Pep1_030","Col_Pep1_
090","Col_Pep1_180")
log2FClist <- lapply(PAMPTime, OnlySigl2FC)
log2FCdf <- do.call("cbind", log2FClist)
nrow(log2FCdf)
```

```
## [1] 26397
```

reduce to genes significantly upregulated in at least one time for all seven (expect 997, MORE than Table S3 becuase this still includes genes induced also in mock/mutant lines)

```
AllPAMPs <- log2FCdf %>% rownames_to_column("ATG") %>% gather(-"ATG", key=Sample, value=L2FC) %>% separa
te(Sample, c("Geno","elicitor","time"), sep="_") %>% group_by(elicitor, ATG) %>% filter(sum(L2FC)>0)  %
>% ungroup() %>% group_by(ATG, time) %>% add_count() %>% filter(n>6) %>% unite(Sample, c("elicitor","tim
e")) %>% spread(key=Sample, value=L2FC)


nrow(AllPAMPs)
```

```
## [1] 997
```

mark genes upregulated (same criteria) in mock or mutant at any time

```
#get significant expression in mutants and mock
mutPAMPTime <- c("sd1-29_LPS_005","sd1-29_LPS_010","sd1-29_LPS_030","sd1-29_LPS_090","sd1-29_LPS_180","l
yk4_5_ch8_005","lyk4_5_ch8_010","lyk4_5_ch8_030","lyk4_5_ch8_090","lyk4_5_ch8_180","efr-1_elf18_005","ef
r-1_elf18_010","efr-1_elf18_030","efr-1_elf18_090","efr-1_elf18_180","fls2c_flg22_005","fls2c_flg22_010"
,"fls2c_flg22_030","fls2c_flg22_090","fls2c_flg22_180","rlp23-1_nlp20_005","rlp23-1_nlp20_010","rlp23-1_
nlp20_030","rlp23-1_nlp20_090","rlp23-1_nlp20_180","Col_mock_005","Col_mock_010","Col_mock_030","Col_moc
k_090","Col_mock_180","pepr1_2_Pep1_005","pepr1_2_Pep1_010","pepr1_2_Pep1_030","pepr1_2_Pep1_090","pepr1
_2_Pep1_180")
mutlog2FClist <- lapply(mutPAMPTime, OnlySigl2FC)
mutlog2FCdf <- do.call("cbind", mutlog2FClist)
nrow(mutlog2FCdf)
```

```
## [1] 26397
```

```
#reduce to genes which have a signficant upregulation in at least one treatment/time combination
upmut <- mutlog2FCdf[rowSums(mutlog2FCdf)>0.5,]
nrow(upmut)
```

```
## [1] 715
```

```
#controlledAllPAMPs <- AllPAMPs[!AllPAMPs$ATG %in% rownames(upmut),]
#nrow(controlledAllPAMPs)
#first surprise: this knocks out some that are in FullUp, which should have been made filtering out gene
s upregulated in mutant or mock. The difference is that was done by asking at each time is this also in
 mock/mutant. In the case of At1g01560, it's upregulated at many times by elf18 in wt, but also at one t
ime in efr-1 (to a small degree), so it was called up at all but 180 for FullUp, and included, but filte
red out here. Is this best? it does NOT explain my discrepancies

colnames(mutlog2FCdf) <- gsub("lyk4_5", "lyk4/5", colnames(mutlog2FCdf))
colnames(mutlog2FCdf) <- gsub("pepr1_2", "pepr1/2", colnames(mutlog2FCdf))
mmreport <- mutlog2FCdf %>% rownames_to_column("ATG") %>% gather(-"ATG", key=Sample, value=L2FC) %>% sep
arate(Sample, c("Geno","elicitor","time"), sep="_") %>% group_by(ATG) %>% summarize(TotalInductionMockMu
t=sum(L2FC))
```

```
## `summarise()` ungrouping output (override with `.groups` argument)
```

```
controlledAllPAMPs <- merge(mmreport, AllPAMPs, by="ATG", all.y=T)
```

Next: don’t FILTER, but give a measure of confidence based on abiotic stress, starting with AtGenExpress: great treatments but microarray

```
#From normalized intensity values, calculate L2FC by probe for AtGen Express. Remove oxidative stress co
ndition
library(Biobase)
library(limma)
data <- read.delim(paste0(home, "/wtonly/Analysis-AtGenExpress/AtGenExpress_expression_byprobe.txt"), he
ader=T, row.names=1)
eset<-new("ExpressionSet", exprs=as.matrix(data))
fordesign <- read.delim(paste0(home, "/wtonly/Analysis-AtGenExpress/AtGenExpress_expression_byprobe.txt"
), header=T, row.names=1, check.names=F)
f <- factor(colnames(fordesign))
design <- model.matrix(~0+f)
colnames(design) <- levels(f)
fit <- lmFit(eset, design)
Time0 <- makeContrasts(Cold_0.5_shoot-Control_0_shoot, Cold_1_shoot-Control_0_shoot, Cold_3_shoot-Contro
l_0_shoot,
                       Osmotic_0.5_shoot-Control_0_shoot, Osmotic_1_shoot-Control_0_shoot, Osmotic_3_sho
ot-Control_0_shoot,
                       Salt_0.5_shoot-Control_0_shoot, Salt_1_shoot-Control_0_shoot, Salt_3_shoot-Contro
l_0_shoot,
                       Drought_0.25_shoot-Control_0_shoot, Drought_0.5_shoot-Control_0_shoot, Drought_1_
shoot-Control_0_shoot, Drought_3_shoot-Control_0_shoot,
                       Genotoxic_0.5_shoot-Control_0_shoot, Genotoxic_1_shoot-Control_0_shoot, Genotoxic
_3_shoot-Control_0_shoot,
                       Oxidative_0.5_shoot-Control_0_shoot, Oxidative_1_shoot-Control_0_shoot, Oxidative
_3_shoot-Control_0_shoot,
                       UVB_0.25_shoot-Control_0_shoot, UVB_0.5_shoot-Control_0_shoot, UVB_1_shoot-Contro
l_0_shoot, UVB_3_shoot-Control_0_shoot,
                       Wounding_0.25_shoot-Control_0_shoot,Wounding_0.5_shoot-Control_0_shoot, Wounding_
1_shoot-Control_0_shoot, Wounding_3_shoot-Control_0_shoot,
                       Heat_0.25_shoot-Control_0_shoot,Heat_0.5_shoot-Control_0_shoot, Heat_1_shoot-Cont
rol_0_shoot, Heat_3_shoot-Control_0_shoot,levels=design)
Time0fit <- contrasts.fit(fit, Time0)
Time0fit <- eBayes(Time0fit)

mock <- makeContrasts(Cold_0.5_shoot-Control_0.5_shoot, Cold_1_shoot-Control_1_shoot, Cold_3_shoot-Contr
ol_3_shoot,
                       Osmotic_0.5_shoot-Control_0.5_shoot, Osmotic_1_shoot-Control_1_shoot, Osmotic_3_s
hoot-Control_3_shoot,
                       Salt_0.5_shoot-Control_0.5_shoot, Salt_1_shoot-Control_1_shoot, Salt_3_shoot-Cont
rol_3_shoot,
                       Drought_0.25_shoot-Control_0.25_shoot, Drought_0.5_shoot-Control_0.5_shoot, Droug
ht_1_shoot-Control_1_shoot, Drought_3_shoot-Control_3_shoot,
                       Genotoxic_0.5_shoot-Control_0.5_shoot, Genotoxic_1_shoot-Control_1_shoot, Genotox
ic_3_shoot-Control_3_shoot,
                       Oxidative_0.5_shoot-Control_0.5_shoot, Oxidative_1_shoot-Control_1_shoot, Oxidati
ve_3_shoot-Control_3_shoot,
                       UVB_0.25_shoot-Control_0.25_shoot, UVB_0.5_shoot-Control_0.5_shoot, UVB_1_shoot-C
ontrol_1_shoot, UVB_3_shoot-Control_3_shoot,
                       Wounding_0.25_shoot-Control_0.25_shoot,Wounding_0.5_shoot-Control_0.5_shoot, Woun
ding_1_shoot-Control_1_shoot, Wounding_3_shoot-Control_3_shoot,
                       Heat_0.25_shoot-Control_0.25_shoot,Heat_0.5_shoot-Control_0.5_shoot, Heat_1_shoot
-Control_1_shoot, Heat_3_shoot-Control_3_shoot,levels=design)
mockfit <- contrasts.fit(fit, mock)
mockfit <- eBayes(mockfit)

condition <- c("Cold_0-5_shoot", "Cold_1_shoot", "Cold_3_shoot",
                       "Osmotic_0-5_shoot", "Osmotic_1_shoot", "Osmotic_3_shoot",
                       "Salt_0-5_shoot", "Salt_1_shoot", "Salt_3_shoot",
                       "Drought_0-25_shoot", "Drought_0-5_shoot", "Drought_1_shoot", "Drought_3_shoot",
                       "Genotoxic_0-5_shoot", "Genotoxic_1_shoot", "Genotoxic_3_shoot",
                       "Oxidative_0-5_shoot", "Oxidative_1_shoot", "Oxidative_3_shoot",
                       "UVB_0-25_shoot", "UVB_0-5_shoot", "UVB_1_shoot", "UVB_3_shoot",
                       "Wounding_0-25_shoot","Wounding_0-5_shoot", "Wounding_1_shoot", "Wounding_3_shoo
```

```
t",
                       "Heat_0-25_shoot","Heat_0-5_shoot", "Heat_1_shoot", "Heat_3_shoot")

Time0table <- topTable(Time0fit, coef=1, adjust="BH", n=Inf)
Time0table <- Time0table[Time0table$logFC>1,]
Time0table <- Time0table[Time0table$adj.P.Val<0.05,]
Time0table$probe <- rownames(Time0table)
Time0table <- Time0table[,c("logFC", "probe")]

mocktable <- topTable(mockfit, coef=1, adjust="BH", n=Inf)
mocktable <- mocktable[mocktable$logFC>1,]
mocktable <- mocktable[mocktable$adj.P.Val<0.05,]
Time0table <- Time0table[Time0table$probe %in% rownames(mocktable),]
ShootAbio <- Time0table
colnames(ShootAbio)[1] <- condition[1]

for(i in 2:31){
  Time0table <- topTable(Time0fit, coef=i, adjust="BH", n=Inf)
  Time0table <- Time0table[Time0table$logFC>1,]
  Time0table <- Time0table[Time0table$adj.P.Val<0.05,]
  Time0table$probe <- rownames(Time0table)
  Time0table <- Time0table[,c("logFC", "probe")]
  colnames(Time0table) <- c(condition[i],"probe")

  mocktable <- topTable(mockfit, coef=i, adjust="BH", n=Inf)
  mocktable <- mocktable[mocktable$logFC>1,]
  mocktable <- mocktable[mocktable$adj.P.Val<0.05,]
  Time0table <- Time0table[Time0table$probe %in% rownames(mocktable),]
  ShootAbio <- merge(ShootAbio, Time0table, by="probe", all=T)
}
```

```
#now do the same for roots
fit <- lmFit(eset, design)
Time0 <- makeContrasts(Cold_0.5_root-Control_0_root, Cold_1_root-Control_0_root, Cold_3_root-Control_0_r
oot,
                       Osmotic_0.5_root-Control_0_root, Osmotic_1_root-Control_0_root, Osmotic_3_root-Co
ntrol_0_root,
                       Salt_0.5_root-Control_0_root, Salt_1_root-Control_0_root, Salt_3_root-Control_0_r
oot,
                       Drought_0.25_root-Control_0_root, Drought_0.5_root-Control_0_root, Drought_1_root
-Control_0_root, Drought_3_root-Control_0_root,
                       Genotoxic_0.5_root-Control_0_root, Genotoxic_1_root-Control_0_root, Genotoxic_3_r
oot-Control_0_root,
                       Oxidative_0.5_root-Control_0_root, Oxidative_1_root-Control_0_root, Oxidative_3_r
oot-Control_0_root,
                       UVB_0.25_root-Control_0_root, UVB_0.5_root-Control_0_root, UVB_1_root-Control_0_r
oot, UVB_3_root-Control_0_root,
                       Wounding_0.25_root-Control_0_root,Wounding_0.5_root-Control_0_root, Wounding_1_ro
ot-Control_0_root, Wounding_3_root-Control_0_root,
                       Heat_0.25_root-Control_0_root,Heat_0.5_root-Control_0_root, Heat_1_root-Control_0
_root, Heat_3_root-Control_0_root,levels=design)
Time0fit <- contrasts.fit(fit, Time0)
Time0fit <- eBayes(Time0fit)

mock <- makeContrasts(Cold_0.5_root-Control_0.5_root, Cold_1_root-Control_1_root, Cold_3_root-Control_3_
root,
                       Osmotic_0.5_root-Control_0.5_root, Osmotic_1_root-Control_1_root, Osmotic_3_root-
Control_3_root,
                       Salt_0.5_root-Control_0.5_root, Salt_1_root-Control_1_root, Salt_3_root-Control_3
_root,
                       Drought_0.25_root-Control_0.25_root, Drought_0.5_root-Control_0.5_root, Drought_1
_root-Control_1_root, Drought_3_root-Control_3_root,
                       Genotoxic_0.5_root-Control_0.5_root, Genotoxic_1_root-Control_1_root, Genotoxic_3
_root-Control_3_root,
                       Oxidative_0.5_root-Control_0.5_root, Oxidative_1_root-Control_1_root, Oxidative_3
_root-Control_3_root,
                       UVB_0.25_root-Control_0.25_root, UVB_0.5_root-Control_0.5_root, UVB_1_root-Contro
l_1_root, UVB_3_root-Control_3_root,
                       Wounding_0.25_root-Control_0.25_root,Wounding_0.5_root-Control_0.5_root, Wounding
_1_root-Control_1_root, Wounding_3_root-Control_3_root,
                       Heat_0.25_root-Control_0.25_root,Heat_0.5_root-Control_0.5_root, Heat_1_root-Cont
rol_1_root, Heat_3_root-Control_3_root,levels=design)
mockfit <- contrasts.fit(fit, mock)
mockfit <- eBayes(mockfit)

condition <- c("Cold_0-5_root", "Cold_1_root", "Cold_3_root",
                       "Osmotic_0-5_root", "Osmotic_1_root", "Osmotic_3_root",
                       "Salt_0-5_root", "Salt_1_root", "Salt_3_root",
                       "Drought_0-25_root", "Drought_0-5_root", "Drought_1_root", "Drought_3_root",
                       "Genotoxic_0-5_root", "Genotoxic_1_root", "Genotoxic_3_root",
                       "Oxidative_0-5_root", "Oxidative_1_root", "Oxidative_3_root",
                       "UVB_0-25_root", "UVB_0-5_root", "UVB_1_root", "UVB_3_root",
                       "Wounding_0-25_root","Wounding_0-5_root", "Wounding_1_root", "Wounding_3_root",
                       "Heat_0-25_root","Heat_0-5_root", "Heat_1_root", "Heat_3_root")

Time0table <- topTable(Time0fit, coef=1, adjust="BH", n=Inf)
Time0table <- Time0table[Time0table$logFC>1,]
Time0table <- Time0table[Time0table$adj.P.Val<0.05,]
Time0table$probe <- rownames(Time0table)
Time0table <- Time0table[,c("logFC", "probe")]

mocktable <- topTable(mockfit, coef=1, adjust="BH", n=Inf)
mocktable <- mocktable[mocktable$logFC>1,]
mocktable <- mocktable[mocktable$adj.P.Val<0.05,]
```

```
Time0table <- Time0table[Time0table$probe %in% rownames(mocktable),]
RootAbio <- Time0table
colnames(RootAbio)[1] <- condition[1]

for(i in 2:31){
  Time0table <- topTable(Time0fit, coef=i, adjust="BH", n=Inf)
  Time0table <- Time0table[Time0table$logFC>1,]
  Time0table <- Time0table[Time0table$adj.P.Val<0.05,]
  Time0table$probe <- rownames(Time0table)
  Time0table <- Time0table[,c("logFC", "probe")]
  colnames(Time0table) <- c(condition[i],"probe")

  mocktable <- topTable(mockfit, coef=i, adjust="BH", n=Inf)
  mocktable <- mocktable[mocktable$logFC>1,]
  mocktable <- mocktable[mocktable$adj.P.Val<0.05,]
  Time0table <- Time0table[Time0table$probe %in% rownames(mocktable),]
  RootAbio <- merge(RootAbio, Time0table, by="probe", all=T)
}


#remove oxidative, combine root and shoot, taking max
Shootlong <- ShootAbio %>% gather(-probe, key="Sample", value="L2FC") %>% separate("Sample", c("Stress",
"Time","Tissue"), sep="_")
Rootlong <- RootAbio %>% gather(-probe, key="Sample", value="L2FC") %>% separate("Sample", c("Stress","T
ime","Tissue"), sep="_")
Abiolong <- rbind(Shootlong, Rootlong)
Abiocombine <- Abiolong %>% filter(Stress != "Oxidative") %>% group_by(probe, Stress, Time) %>% summariz
e(mL2FC = ifelse(all(is.na(L2FC)), 0, max(L2FC, na.rm=T))) %>% unite(Sample, c("Stress", "Time")) %>% sp
read(key=Sample, value=mL2FC)
```

```
## `summarise()` regrouping output by 'probe', 'Stress' (override with `.groups` argument)
```

```
nrow(RootAbio)
```

```
## [1] 4499
```

```
nrow(ShootAbio)
```

```
## [1] 4363
```

```
nrow(Abiocombine)
```

```
## [1] 6481
```

filter and output tables of genes meeting the following criteria: upregulated compared to time 0, upregulated compared to mock, log2fold change (absolute value) greater than one, and corrected p less than 0.05. Can then look at these like my DESeq2 data sets (upset, etc)

```
#probes to genes: if multiple probes target one gene, take max value. If one probe targets multiple gene
s, remove that probe and all genes
Abiolong <- Abiocombine %>% gather(-probe, key=sample, value=L2FC)

library(ath1121501.db)
library(AnnotationDbi)
library(affy)
library(annotate)

PROBES<- keys(ath1121501.db)
#select(ath1121501.db, c("244901_at","244902_at"), c("TAIR","SYMBOL", "GENENAME"))

#the one:many here doesn't always line up with the probe matches I downloaded from TAIR. since that file
said 2010 and this is Bioconductor which should be updated every six months, going to go with this
MERGEME <- select(ath1121501.db, PROBES, "TAIR")
```

```
## 'select()' returned 1:many mapping between keys and columns
```

```
colnames(MERGEME) <- c("probe","ATG")
#find probes that match to more than one gene
oneprobe_manygenes <- MERGEME %>% group_by(probe) %>% add_count() %>% filter(n>1)

Abioverylong <- merge(Abiolong, MERGEME, by="probe", all.x=T)
Abioverylong <- Abioverylong[!is.na(Abioverylong$ATG),]


#remove probes matching to more than one gene and average results from multiple probes that match to sam
e gene
one2one <- Abioverylong %>% group_by(probe, sample) %>% add_count() %>% filter(n<2) %>% ungroup() %>% gr
oup_by(ATG, sample) %>% summarize(mL2FC=mean(L2FC, na.rm=T))
```

```
## `summarise()` regrouping output by 'ATG' (override with `.groups` argument)
```

```
#get count of abiotic stresses which induce a gene (at any time)
NumberAbio <- one2one %>% separate(sample, c("stress","time"), sep="_") %>% group_by(stress, ATG) %>% fi
lter(sum(mL2FC)>0)  %>% ungroup() %>% group_by(ATG) %>% mutate(count = length(unique(stress))) %>% unite
(Sample, c("stress","time")) %>% ungroup() %>% group_by(ATG) %>% summarize(AtGenExpressNumber=mean(coun
t))
```

```
## `summarise()` ungrouping output (override with `.groups` argument)
```

```
#Add count Abio on to controlled all PAMPs
CAP_AtGen <- merge(NumberAbio, controlledAllPAMPs, by="ATG", all.y=T)
CAP_AtGen$AtGenExpressNumber[is.na(CAP_AtGen$AtGenExpressNumber)] <- 0

#by gene assign AbioInduction: if nothing = "none", if one stress within three hours = "one", if 2-4 str
esses = "few", if 5-6 stresses "many" if 7 stresses "all". If 1-4 stress >3 hours, "later", if 5-7 stres
ses >3 hours, "manylater".(expect ~80 "none")
CAP_AtGen$AtGenExpressCategorical <- "none"
CAP_AtGen$AtGenExpressCategorical[!CAP_AtGen$ATG %in% MERGEME$ATG] <- "NotOnArray"
CAP_AtGen$AtGenExpressCategorical[CAP_AtGen$ATG %in% oneprobe_manygenes$ATG] <- "ProbeHitsMultipleGenes"
CAP_AtGen$AtGenExpressCategorical[CAP_AtGen$AtGenExpressNumber == 1] <- "one"
CAP_AtGen$AtGenExpressCategorical[CAP_AtGen$AtGenExpressNumber > 1] <- "few"
CAP_AtGen$AtGenExpressCategorical[CAP_AtGen$AtGenExpressNumber > 4] <- "many"
CAP_AtGen$AtGenExpressCategorical[CAP_AtGen$AtGenExpressNumber > 6] <- "all"
CAP_AtGen <- CAP_AtGen[,c(1,2,41,3, 4:40)]

nrow(CAP_AtGen[CAP_AtGen$AtGenExpressCategorical=="none",])
```

```
## [1] 75
```

further information from the few RNAseq experiments I could find - for this files were assembled from different papers anyway, just going straight from that aggregate file

```
#by gene assign RNAseqAbio: if nothing "none", if 1 "one", if x-y "few", if  (expect ~30 "none" "none")
RNAseq <- read.csv(paste0(home,"/wtonly/Analysis-AtGenExpress/RNAseq_logFC.csv"), header=T)

RNAseqcount <- RNAseq %>% gather(-"ATG", key=Sample, value=L2FC) %>% group_by(ATG) %>% filter(!is.na(L2F
C)) %>% add_count() %>% summarize(AbioRNAseqNumber=mean(n))
```

```
## `summarise()` ungrouping output (override with `.groups` argument)
```

```
CAP_abio <- merge(RNAseqcount, CAP_AtGen, by="ATG", all.y=T)
CAP_abio$AbioRNAseqNumber[is.na(CAP_abio$AbioRNAseqNumber)] <- 0

CAP_abio$AbioRNAseqCategorical <- "none"
CAP_abio$AbioRNAseqCategorical[CAP_abio$AbioRNAseqNumber == 1] <- "one"
CAP_abio$AbioRNAseqCategorical[CAP_abio$AbioRNAseqNumber > 1] <- "few"
CAP_abio$AbioRNAseqCategorical[CAP_abio$AbioRNAseqNumber > 3] <- "many"
CAP_abio<- CAP_abio[,c(1,2,3,4, 43, 5:42)]

nrow(CAP_abio[CAP_abio$AtGenExpressCategorical=="none" & CAP_abio$AbioRNAseqCategorical=="none",])
```

```
## [1] 43
```

get descriptions

```
library(org.At.tair.db)
GENES <- CAP_abio$ATG
info <- select(org.At.tair.db, as.character(GENES), c("TAIR", "SYMBOL","GENENAME"))
```

```
## 'select()' returned 1:many mapping between keys and columns
```

```
info <- info %>% group_by(TAIR) %>% summarize(Name = SYMBOL[1], Description = GENENAME[1])
```

```
## `summarise()` ungrouping output (override with `.groups` argument)
```

```
colnames(info) <- c("ATG","Name","Description")

CAP_description <- merge(info, CAP_abio, by="ATG",all.y=T)
```

```
OnlySigl2FCmm <- function(TrtTime){
  filepath=paste0(home,"vsMockMut/",TrtTime,"_results")
  load(filepath)
  resdf <- as.data.frame(res)
  nonsig <- subset(resdf, padj >0.05 | padj == "NA")
  smallchange <- subset(resdf, log2FoldChange <  1 & log2FoldChange > -1)
  onlysig <- resdf
  onlysig[rownames(onlysig) %in% rownames(nonsig),] <- "0"
  onlysig[rownames(onlysig) %in% rownames(smallchange),] <- "0"
  sigFCs <- subset(onlysig, select = "log2FoldChange")
  sigFCs$log2FoldChange <- as.numeric(sigFCs$log2FoldChange)
  names(sigFCs) <- TrtTime
  return(sigFCs)
}

PAMPTimemm <- c("LPS_005","LPS_010","LPS_030","LPS_090","LPS_180","CO8_005","CO8_010","CO8_030","CO8_09
0","CO8_180","elf18_005","elf18_010","elf18_030","elf18_090","elf18_180","flg22_005","flg22_010","flg22_
030","flg22_090","flg22_180","nlp20_005","nlp20_010","nlp20_030","nlp20_090","nlp20_180","OGs_005","OGs_
010","OGs_030","OGs_090","OGs_180","Pep1_005","Pep1_010","Pep1_030","Pep1_090","Pep1_180")
log2FClistmm <- lapply(PAMPTimemm, OnlySigl2FCmm)
log2FCdfmm <- do.call("cbind", log2FClistmm)
nrow(log2FCdfmm)
```

```
## [1] 26397
```

get number of genes induced when comparing to mock/PRR mutant (for each PAMP at each time), add that information to larger dataframe

```
AllPAMPsmm <- log2FCdfmm %>% rownames_to_column("ATG") %>% gather(-"ATG", key=Sample, value=L2FC) %>% se
parate(Sample, c("elicitor","time"), sep="_") %>% group_by(elicitor, ATG) %>% filter(sum(L2FC)>0)  %>% u
ngroup() %>% group_by(ATG, time) %>% add_count() %>% ungroup() %>% group_by(ATG) %>% summarize(number_vM
M=mean(n))
```

```
## `summarise()` ungrouping output (override with `.groups` argument)
```

```
nrow(AllPAMPsmm)
```

```
## [1] 5973
```

```
CAP_vMM <- merge(AllPAMPsmm, CAP_description, by="ATG", all.y=T)
```

#write.csv(CAP_description, file="newCIR.csv")

get counts at time 0, need to go in and manually check for genes which are not induced at time 0, possible they are induced only by genes misaligning from another highly expressed gene

```
load(paste0(home, "cleaned_ddsT"))

counts <- counts(ddsT, normalized=T)
onlytime0 <- counts %>% as.data.frame() %>% dplyr::select(matches("000"))
startingcounts <- rowSums(onlytime0)/52
tomerge <- rownames_to_column(as.data.frame(startingcounts), "ATG")
median(tomerge$startingcounts)
```

```
## [1] 78.48919
```

```
mean(tomerge$startingcounts)
```

```
## [1] 323.1574
```

```
CAP_final <- merge(tomerge, CAP_vMM, by="ATG", all.y=T)
```

```
categoricals <- CAP_final[,c("Description","ATG","Name", "number_vMM","n", "startingcounts","TotalInduct
ionMockMut", "AbioRNAseqNumber","AbioRNAseqCategorical", "AtGenExpressNumber", "AtGenExpressCategorical"
)]
L2FCs <- CAP_final[,c(13:ncol(CAP_final))]

total <- cbind(categoricals, L2FCs)

#filter: remove anything induced in any abiotic stress condition
filtered <- total[total$AbioRNAseqCategorical=="none" & total$AtGenExpressCategorical=="none",]

#Filter: remove anything induced in mock or mutant line treatments, UNLESS it is also induced in Col-0 r
elative to mock or mutant lines (might just pass threshold in mutant, but be strongly induced in wt)
filtered <- filtered[filtered$number_vMM > 6 | filtered$TotalInductionMockMut <1,]
```

```
CIR <- filtered %>% mutate(TotalL2FC=rowSums(.[,12:46])) %>% dplyr::select(ATG, TotalL2FC, everything())
%>% arrange(desc(TotalL2FC))

#One gene manually filtered: AT3G32090 appears to be a pseudogene with strong homology to WRKY40 in one
 small region: entirety of counts mapping to this gene map to only that region, and WRKY 40 is a highly
 expressed gene with strong induction. Most likely, this is just an artefact of a few mis-mapped reads.
 Several other genes had low basal expression, likely contributing to seemingly high induction. Manually
checked mapping of these genes and indeed some seemed influenced by nearby highly expressed genes, but n
one as obvious as AT3G32090

CIR <- CIR %>% filter(ATG != 'AT3G32090')

print(CIR[,1:2])
```

```
##           ATG TotalL2FC
## 1  AT2G29100 65.765687
## 2  AT1G36640 55.467887
## 3  AT4G13510 44.302075
## 4  AT1G66460 42.596469
## 5  AT4G14400 39.534441
## 6  AT4G19520 38.576384
## 7  AT4G23250 38.334736
## 8  AT1G51900 37.631973
## 9  AT5G45000 35.333783
## 10 AT3G44350 35.169245
## 11 AT2G20150 34.997193
## 12 AT5G05290 34.298679
## 13 AT1G10417 33.643695
## 14 AT1G11310 31.354798
## 15 AT5G24240 31.332791
## 16 AT5G57480 29.440758
## 17 AT1G64400 29.370068
## 18 AT1G72000 28.470024
## 19 AT4G34460 28.093459
## 20 AT1G28390 27.032083
## 21 AT4G26930 26.791642
## 22 AT2G25470 26.426990
## 23 AT4G00955 26.180352
## 24 AT2G22890 25.044793
## 25 AT1G50180 23.946975
## 26 AT4G13420 23.434112
## 27 AT2G29120 22.252845
## 28 AT5G24090 21.656334
## 29 AT5G13550 20.303378
## 30 AT1G69523 19.533964
## 31 AT1G08250 19.459335
## 32 AT5G54140 19.050969
## 33 AT1G03730 16.177294
## 34 AT5G48550 15.635742
## 35 AT4G22030 13.386265
## 36 AT3G57210 11.785214
## 37 AT4G30500  9.731193
## 38 AT5G40910  9.620492
```
